## Supplementary Figures for "Connectomic Analysis of the *Drosophila* Lateral Neuron Clock Cells Reveals the Synaptic Basis of Functional Pacemaker Classes"

Figure S1

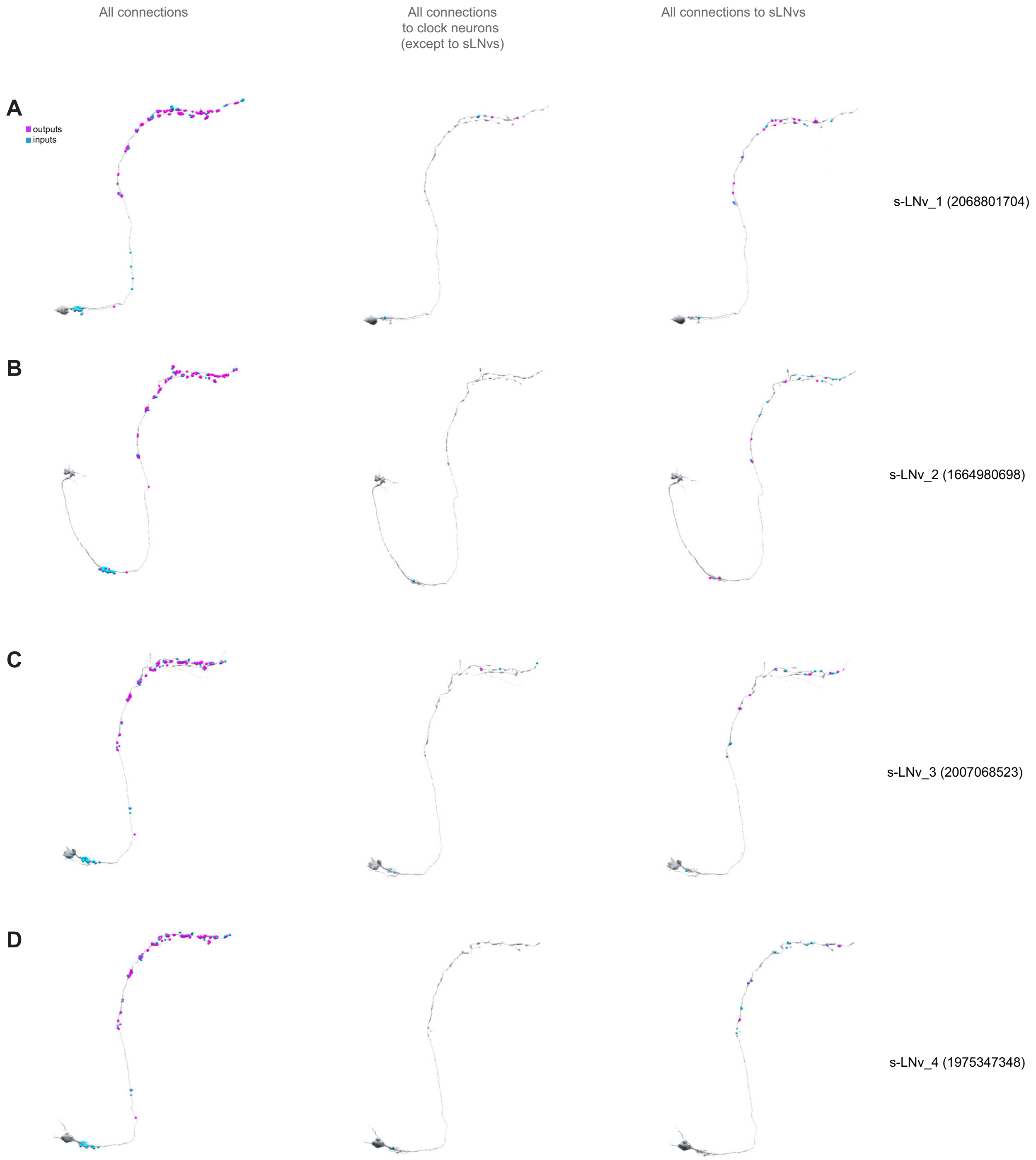

All connections
All connections to clock neurons (except to sLNvs)
All connections to sLNvs
A
outputs
inputs
s-LNv_1 (2068801704)
B
s-LNv_2 (1664980698)
C
s-LNv_3 (2007068523)
D
s-LNv_4 (1975347348)

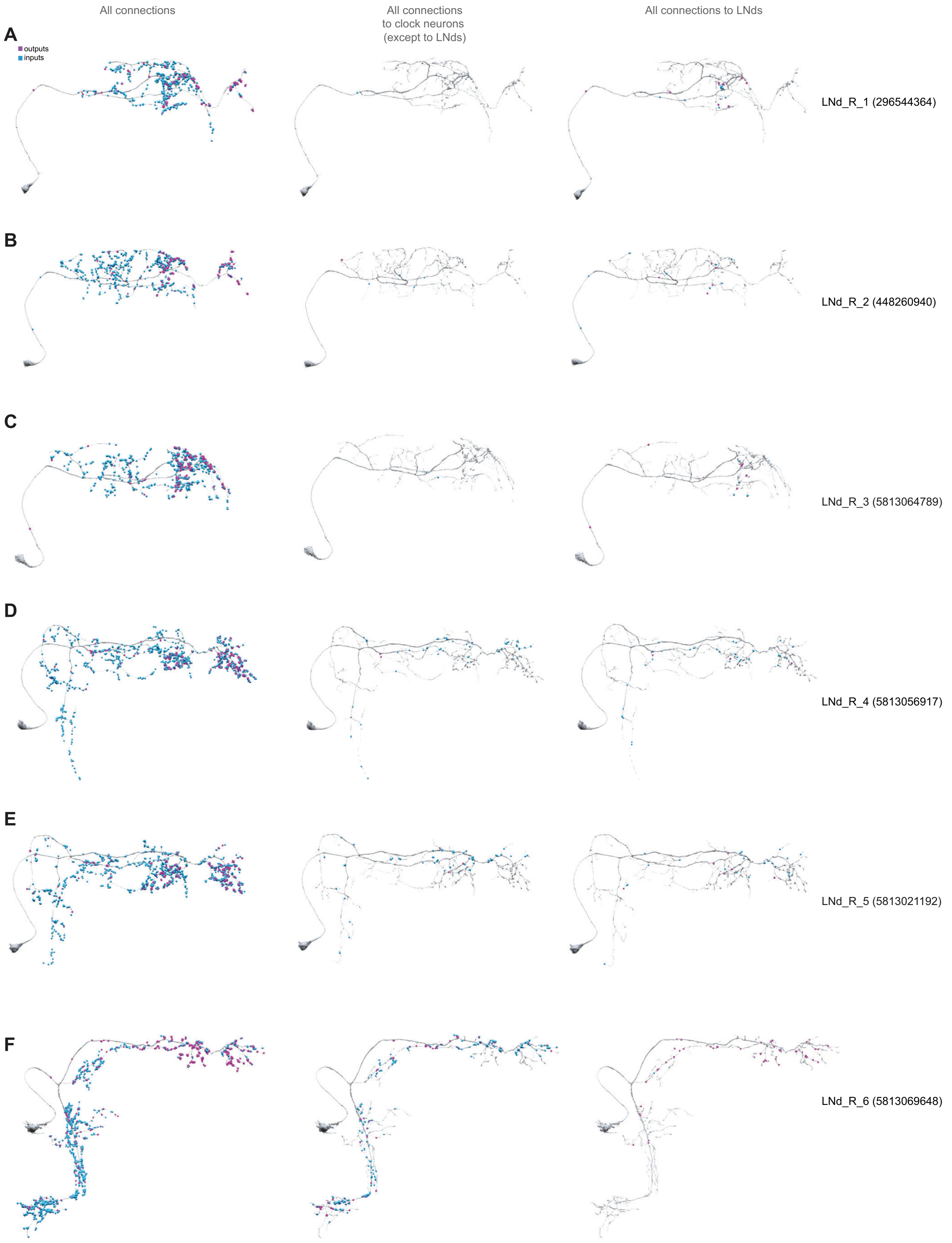
All connections
All connections to clock neurons (except to LNds)
All connections to LNds
A
outputs
inputs
LNd_R_1 (296544364)
B
LNd_R_2 (448260940)
C
LNd_R_3 (5813064789)
D
LNd_R_4 (5813056917)
E
LNd_R_5 (5813021192)
F
LNd_R_6 (5813069648)

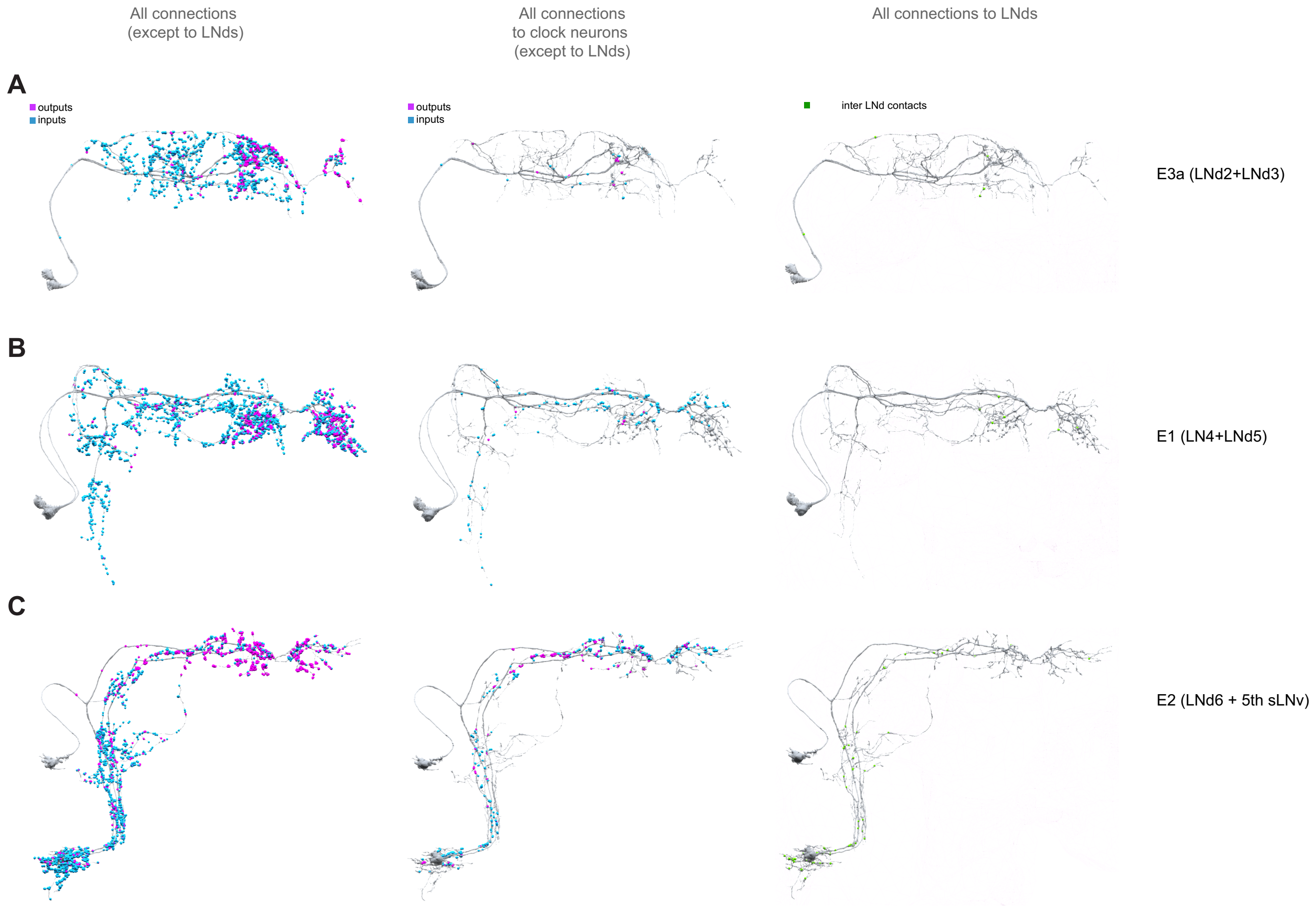
All connections
(except to LNds)
All connections
to clock neurons
(except to LNds)
All connections to LNds
A
outputs
inputs
outputs
inputs
inter LNd contacts
E3a (LNd2+LNd3)
B
E1 (LN4+LNd5)
C
E2 (LNd6 + 5th sLNv)

A

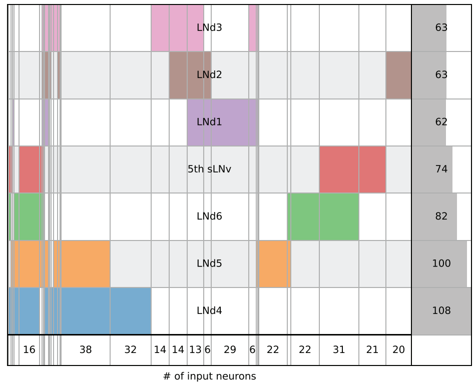
LNd3
LNd2
LNd1
5th sLNv
LNd6
LNd5
LNd4
63
63
62
74
82
100
108
16
38
32
14
14
13
6
29
6
22
22
31
21
20
### of input neurons

B

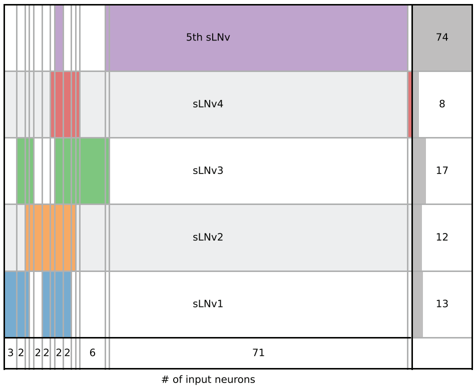
5th sLNv
sLNv4
sLNv3
sLNv2
sLNv1
74
8
17
12
13
3
2
2
2
2
2
6
71
### of input neurons

Figure S5.

**A**

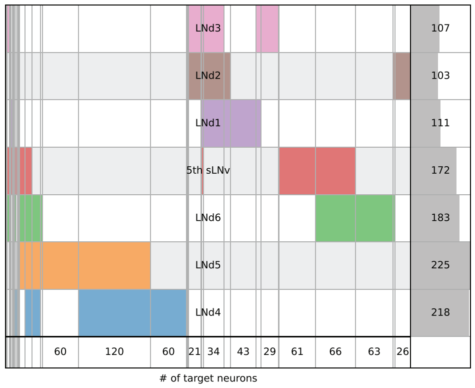

LNd3
LNd2
LNd1
5th sLNv
LNd6
LNd5
LNd4
107
103
111
172
183
225
218
60
120
60
21
34
43
29
61
66
63
26
### of target neurons


**B**

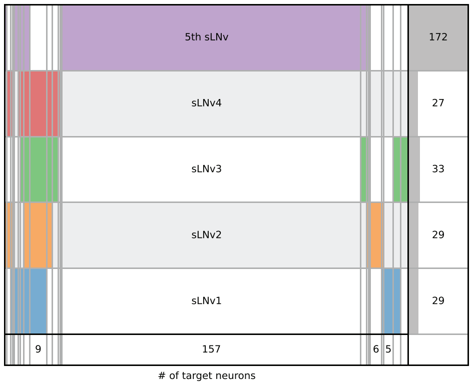

5th sLNv
sLNv4
sLNv3
sLNv2
sLNv1
172
27
33
29
29
9
157
6
5
### of target neurons

**A**

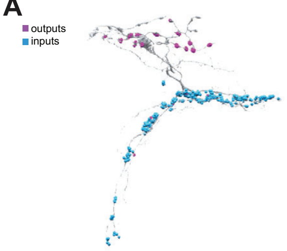


355453590 inputs from all clock cells
and outputs to all clock cells
SLP316_R

**D**

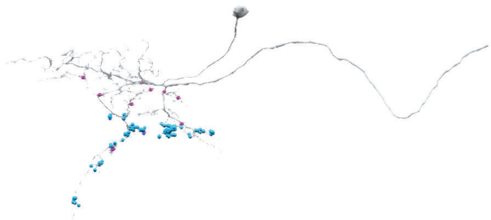

325455002 inputs from all clock cells
and outputs to all clock cells
SLP403_R

**B**

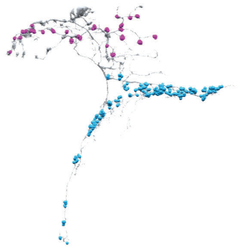

325122525 inputs from all clock cells
and outputs to all clock cells
SLP316_R

**E**

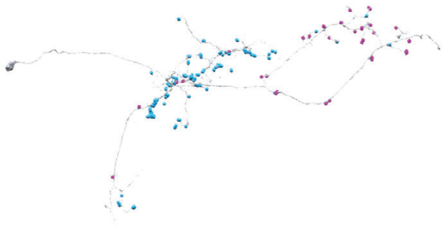

540998882 inputs from all clock cells
and outputs to all clock cells
SLP403_R

**C**

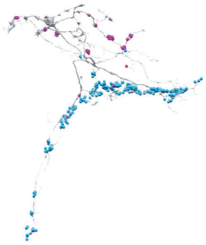

5813047586 inputs from all clock cells
and outputs to all clock cells
SLP316_R

**F**

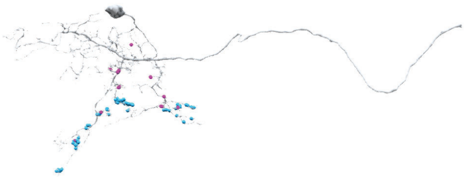

294783216 inputs from all clock cells
and outputs to all clock cells
SMP223_R

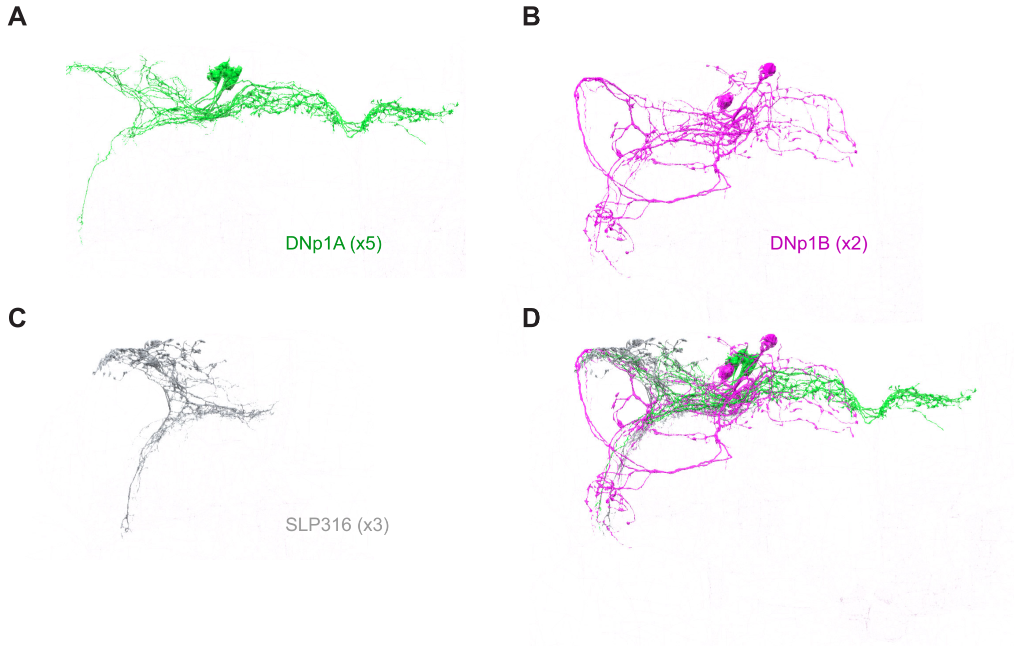
A
DNp1A (x5)
B
DNp1B (x2)
C
SLP316 (x3)
D
