## Supplementary Table 1 for "Connectomic Analysis of the *Drosophila* Lateral Neuron Clock Cells Reveals the Synaptic Basis of Functional Pacemaker Classes"

**Table S1. Inputs to clock neurons**

| | sLNv-1 | sLNv-2 | sLNv3 | sLNv-4 | l-LNv-1 | l-LNV-2 | l-LNv-3 | l-LNv-4 | LNd-1 | LNd-2 | LNd-3 | LNd-4 | LNd-5 | LNd-6 | 5th sLNV | DN1a-1 | DN1a-2 | DN1pA-1 | DN1pA-2 | DN1pA-3 | DN1pA-4 | DN1pA-5 | DN1pB-1 | DN1pB-2 | LPN-1 | LPN-2 | LPN-3 | LPN-4 |
|---|---|---|---|---|---|---|---|---|---|---|---|---|---|---|---|---|---|---|---|---|---|---|---|---|---|---|---|---|
| bodyId_pre | 2.069E+09 | 1.665E+09 | 2.007E+09 | 1.975E+09 | 1.885E+09 | 2.066E+09 | 5.813E+09 | 5.813E+09 | 296544364 | 448260940 | 5.813E+09 | 5.813E+09 | 5.813E+09 | 5.813E+09 | 511051477 | 264083994 | 5.813E+09 | 5.813E+09 | 324846570 | 325529237 | 387944118 | 387166379 | 386834269 | 5.813E+09 | 356818551 | 450034902 | 480029788 | 546977514 |
| 200326126 | | | | | | | | | | | | | | | | 3 | 4 | | | | | | | | | | | |
| 233105330 | | | | | | | | | | | 1 | | 2 | | | | | | | | | | | | | | | |
| 263475451 | | | | | | | | | 2 | | 1 | | | | | | | | | | | | | 1 | 2 | | | |
| 264083994 | | | | | | | | | 1 | | | | | 81 | 55 | | 10 | | | | | | | 1 | 3 | | 1 | |
| 264822904 | | | | | | | | | | | | | | | | | | | | | | | 2 | | | | | |
| 265120589 | | | | | | | | | | | | | | | | | | | | | | | | | 1 | | | |
| 265124786 | | | | | | | | | | 7 | 3 | | | | | | | | | | | | | 1 | | | | |
| 266187477 | | | | | | | | | | | | | | | | | | | | | | | | 1 | | | | |
| 266187480 | | | | | | | | | 1 | | | | | | | | | | | | | | | | | | | |
| 266187559 | | | | | | | | | | | | | | | | | | | | | | | | | 1 | | | |
| 266873389 | | | | | | | | | | 4 | | | | | | | | | | | | | | | | | | |
| 266873401 | | | | | | | | | 1 | 1 | | | | | | | | | | | | | | | | | | |
| 266882091 | | | | | | | | | 2 | | | | | | | | | | | | | | | | | | | |
| 266882171 | | | | | | | | | 5 | 5 | | | | | | | | | | | | | | | | | | |
| 267210124 | | | | | | | | | 4 | | | | | | | | | | | | | | | | | | | |
| 267210191 | | | | | | | | | 5 | | 2 | | | | | | | | | | | | | | | | | |
| 267214250 | | | | | | | | | | 5 | 11 | | | | | | | | | | | | | | | | | |
| 267218804 | | | | | | | | | 1 | | | | | | | | | | | | | | | | | | | |
| 267551622 | | | | | | | | | | 1 | | | | | | | | | | | | | | | | | | |
| 294424196 | | | | | | | | | | | 1 | | | | | 4 | | | 3 | | | | | | | | | |
| 294428266 | | | | | | | | | | | | | | | | | | | | | 1 | | | | 1 | | | |
| 294432626 | | | | | | | | | | | | | | | | 7 | | 3 | | | 1 | | | | 4 | | | |
| 294432760 | | | | | | | | | | 1 | | | | | | | | | | | | | | | | | | |
| 294432807 | | | | | | | | | 1 | | | | | | | | | | | | | | | | | | | |
| 294436967 | | | | | | | | | 1 | 3 | 2 | | 2 | | | 3 | 3 | | | 1 | | | 2 | | 4 | 2 | 1 | |
| 294437328 | | | | | | | | | | 1 | | | | | | | | | | | | | | | | | | |
| 294437347 | | | | | | | | | | | | | | | | | | | | | | | 6 | 1 | 1 | | | |
| 294441510 | | | | | | | | | | 2 | | | | | | | | | | | | | | | | | | |
| 294445804 | | | | | | | | | | 6 | 10 | | | | | 1 | | | | | | | | | | | | |
| 294450578 | | | | | | | | | | 1 | | | | | | | | | | | | | | | | | | |
| 294756576 | | | | | | | | | | | | | | | | | | | | 1 | | | | | | | | |
| 294760699 | | | 1 | | | | | | | 1 | 5 | 7 | 6 | | | | | 6 | 10 | 9 | 6 | 12 | | 1 | 2 | 2 | 1 | |
| 294760837 | | | | | | | | | | 1 | | | | | | | | | | | | | | | 1 | | | |
| 294774792 | | | | | | | | | | | | | | | | | | | 1 | | | | | | | | | |
| 294783216 | | | | 1 | | | | | | | | | | | | | | 2 | 3 | 3 | 2 | 6 | | | | | | |
| 294791509 | | | | | | | | | | | 1 | | | | | | | | | | | | | | | | | |
| 294795963 | | | | | | | | | | | | | | | | | | | | | | | 1 | | | 1 | | |
| 295103172 | | | | | | | | | | 1 | | | | | | | | | | | | | | | | | | |
| 295111734 | | | | | | | | | | 1 | | | | | | | | | | | | | | | | | | |
| 295127956 | | | | | | | | | | | 1 | | | | | | | | | | | | | | | | | |
| 295132845 | | | | | | | | | | | | | | | | | | | | | | | | 1 | | | | |
| 295133778 | | | | | | | | | | | | | | | | | | | | | | 1 | | | | | | |
| 295133902 | | | | | | | | | | | | | | | | | | | | | | | | | 1 | | | |
| 295137019 | | | | | | | | | | | | | | | | | | | | | | | | | 1 | | | |
| 295137392 | | | | | | | | | | | | | | | | | | 1 | | | | | | | | | | |
| 295141618 | | | | | | | | | | | | | | | | | | 1 | | | | | | | | | | |
| 295434457 | | | | | | | | | | | | 2 | | | | | | | | | | | | | | | | |
| 295443136 | | | | | | | | | | 1 | 4 | 10 | 6 | | | | | | | | | | 1 | | | | | |
| 295443724 | | | | | | | | | | 1 | 1 | 2 | 1 | | | 2 | | | | | | | | | 4 | | | |
| 295473947 | | | | | | | | | | | 1 | | | | | | | | | | | | | | | | | |
| 295474250 | | | | | | | | | | 1 | 1 | | | | | | | | | | | | | | | | | |
| 295478082 | | | | | | | | | | 1 | 2 | | | | | | | | | | | | | | | | | |
| 295482340 | | | | | | | | | | | | | | | | | | | | | | | | 1 | | | | |
| 295482551 | | | | | | | | | | 1 | | | | | | | | | | | | | | | | | | |
| 295490863 | | | | | | | | | | | | | | | | | | | | | | | | 1 | | | | |
| 295780154 | | | | | | | | | | 2 | 2 | | | | | | | | | | | | | | | | | |
| 295793991 | | | 1 | | | | | | | | | | | | | | | | | | | | | | | | | |
| 295797312 | | | | | | | | | | 1 | 6 | | | | | | | | | | | | | | | | | |
| 295802733 | | | | | | | | | | 7 | 3 | 7 | 2 | | | | | | | | | | | | | | | |
| 295807429 | | | | | | | | | | 1 | | | | | | | | | | | | | | | | | | |
| 295814932 | | | | | | | | | | | | | | | | | | | | | | | | | 1 | | | |
| 295819998 | | | | | | | | | | | | | | | | | | 1 | | | | | | | | | | |
| 295827899 | | | | | | | | | | | | | | | | | | | | | | | 2 | | | | | |
| 295828029 | | | | | | | | | | | | | | | | | | | | | | | | | | | 1 | |
| 295849217 | | | | | | | | | | | | | | | | | | | | | | | 1 | | | | | |
| 296120593 | | | | | | | | | | 3 | 1 | 1 | | | | | | | 1 | 2 | 1 | | 2 | 4 | 4 | | | |
| 296139882 | | | | | | | | | 2 | | | 1 | | | | | | | | | | | | | | 4 | 5 | |
| 296164879 | | | | | | | | | | 2 | | 1 | | | | 1 | | | | | | | | | 1 | | | |
| 296168382 | | | | | | | | | | | 2 | | | | | | | | | | | | | | | | | |
| 296199026 | | | | | | | | | | | 2 | | | | | | | | | | | | | | | | | |
| 296199149 | | | | | | | | | | | | | | | | | | | | | | | | | | 1 | | |
| 296203440 | | | | | | | | | | | | | 1 | | | | | | | | | | | | | | | |

| | | | | | | | | | | | | | | | | | | | | | | | |
|---|---|---|---|---|---|---|---|---|---|---|---|---|---|---|---|---|---|---|---|---|---|---|---|
| 296203608 | | | | | | | 1 | | | | | | | | | | | | | | | | |
| 296207652 | | | | | | | | | | | | | | | | | | 1 | | | | | |
| 296475214 | | | | | | | | | | | | 3 | | | | | | | | | | | |
| 296489073 | | | | | | 1 | | | | | | | | | | | | | | | | | |
| 296496999 | | | | | | 1 | | | | | | 1 | | | | | | | | | | | |
| 296502002 | | | | | | 1 | | | | | | | | | | | | | | | | | |
| 296509673 | | | | | | | | | | | | | | | | | | 4 | 3 | | | | |
| 296509709 | | | 1 | | | | | | | | | | | | | | | | | | | | |
| 296543992 | | | | | 3 | 2 | 3 | 1 | | | | | | | | | | 6 | 9 | | | | |
| 296544262 | | | | 1 | 5 | | | 1 | 1 | 1 | | | 1 | 5 | 1 | 1 | 4 | 1 | 2 | | | | |
| 296544364 | | | | | | 6 | 3 | | | | | | | | | | | | | | | | |
| 296556885 | | | | | 3 | 2 | | | | | | | | | | | | | | | | | |
| 296832977 | | | | | | | 1 | | | | | | | | | | | | | | | | |
| 296837846 | | | | | 1 | 3 | 1 | | | | | | | | | | | | | | | | |
| 296838079 | | | | | | | | | | | | | | | | | | | | 2 | | | |
| 296838734 | | | | | | 1 | 2 | | | | | | | | | | | | | | | | |
| 296843037 | | | | | | 1 | | | 1 | | | | | 1 | 1 | | | | | | 1 | | |
| 296846109 | | | | | 1 | 1 | | | | | | | | | | | | | | | | | |
| 296859438 | | | | | 1 | | | | 2 | | | | | 1 | | | | 14 | 4 | | | 1 | |
| 296885154 | | | | | 3 | | | | | | | | | | | | | | | | | | |
| 296885362 | | | | | 2 | | | | | | | | | | | | | | | | | | |
| 296889667 | | | | | 4 | 6 | | | | | | | | | | | | | | | | | |
| 296889776 | | | | | 1 | | | | | | | | | | | | | | | | | | |
| 297192193 | | | | | | 1 | | | | | | | | | | | | | | | | | |
| 297192204 | | | | | | | | | | | | | | | | | | | | | | 1 | |
| 297196636 | | | | | | | | 1 | | | | | | | | | | | | 1 | | | |
| 297221971 | | | | | | | 2 | | | | | | | | | | | 2 | | | | | |
| 297225829 | | | | | 1 | | | | | | | | | | | | | | | | | | |
| 297226270 | | | | | | 1 | | | | | | | | | | | | | | | | | |
| 297229974 | | | | | 1 | | | | | | | | | | | | | | | | | | |
| 297230313 | | | | | 1 | | | | | | | | | | | | | | | | | | |
| 297230760 | | | | | 3 | 2 | 2 | | | | | | | | | | | | 2 | 1 | | 1 | |
| 297234448 | | | | | 1 | | | | | | | | | | | | | | | | | | |
| 297239186 | | | | | | | 1 | | | | | | | | | | | | | | | | |
| 297239209 | | | | | 1 | | | | | | | | | | | | | | | | | | |
| 297239231 | | | | | | 1 | | | | | | | | | | | | | | | | | |
| 297243542 | | | | | 53 | 17 | 28 | 3 | | | | | | | | | | | | 1 | | | |
| 297247801 | | | | | | | | | 1 | | 3 | | | 1 | | | | | | | 1 | | |
| 297247805 | | | | | 2 | | | | | | | | | | | | | | | | | | |
| 297251714 | | | | | | | 1 | | | | | | | | | | | | | | 8 | 7 | |
| 297502392 | | | | | | | | | | | | | | | | | | | | 2 | | | |
| 297520036 | | | | | 1 | | | 1 | | | | | | | | | | | | | | | |
| 297524212 | | | | | | 1 | 1 | | | | | | | | | | | | | | | | |
| 297528204 | | | | | 1 | | | | | | | | | | | | | | | | | | |
| 297533212 | | | | | | | 1 | | | | | | | | | | | | | | | | |
| 297541369 | 2 | 2 | 2 | 1 | | | | | | | | | | | | | | 1 | 1 | | | | |
| 297558674 | | | | | | 1 | | | | | | | | | | | | | | | | | |
| 297558862 | | | | | | | 3 | | | | | | | | | | | | | | | | |
| 297562705 | | | | | 1 | | | | | | | | | | | | | | | | | | |
| 297563139 | | | | | | | | 6 | 7 | | | | | | | | | | | | | 4 | |
| 297563270 | | | | | 3 | 2 | | | | | | | | | | | | | | | | | |
| 297571222 | | | | | 3 | | | | | | | | | | | | | | | | | | |
| 297572235 | | | | | 1 | | | | | | | | | | | | | | | | | | |
| 297579726 | | | | | 1 | | | | | | | | | | | | | | | | | | |
| 297584752 | | | | | | | | | | | | | | | | | | 3 | | | | | |
| 297584869 | | | | | 4 | | | | | | | | | | | | | | | | | | |
| 297588882 | | | | | | 1 | | | | | | | | | | | | | | | | | |
| 297588995 | | | | | 2 | 1 | | | | | | | 1 | | | | | | 2 | | | | |
| 297851977 | | | | | 8 | 6 | 2 | | | | | 1 | | | | | | | | | | | |
| 297860964 | | | | | 3 | | | | | | | | | | | | | | | | | | |
| 297899921 | | | | | 6 | 3 | | | | | | | | | | | | | | | | | |
| 297908801 | | | | | 3 | 2 | | | | | | | | | | | | | | | | | |
| 297912117 | | | | | 1 | | | 4 | 3 | | | | | | | | | | | 1 | | | |
| 297913231 | | | | | 2 | | | | | | | | | | | | | | 2 | | | | |
| 297917158 | | | | | | 1 | 2 | | | | | | | | | | | | | | | | |
| 297917363 | | | | | | | | | | | | | | | | | | | | 3 | 1 | 1 | |
| 297917475 | | | | | 5 | 1 | | | | | | | | | | | | | | | | | |
| 297921570 | | | | | | 2 | 1 | | | | | | | | 2 | | | | | | 1 | | |
| 297921608 | | | | | | | | | | | | | | | | | | 1 | | | | | |
| 297921642 | | | | | 1 | | | | | | | | | | | | | | | | | | |
| 297921808 | | | | | 2 | | | | | | | | | | | | | | | | | | |
| 297925478 | | | | | | 1 | 2 | 4 | 4 | 3 | | | | | | | | | | 1 | | | |
| 297925596 | | | | | 2 | 1 | | | | | | | 2 | | | | | | | | | | |
| 297925608 | | | | | | | | | | | | | | | | | | | | 4 | 1 | | 1 |
| 297925628 | | | | | | | | 3 | | | | | | | | | | | | | | 2 | |
| 297925742 | | | | | 1 | | | | | | | | | | | | | | | | | | |
| 297925765 | | | | | 1 | | | | | | | | | | | | | | | | | | |
| 297925887 | | | | | | 1 | | | | | | | | | | | | | | | | | |
| 297929807 | | | | | | 1 | | | | | | | | | | | | | | | | | |
| 297934028 | | | | | 1 | | | | | | | | | | | | | | | | | | |

| | | | | | | | | | | | | | | | | | | | | | | | | |
|---|---|---|---|---|---|---|---|---|---|---|---|---|---|---|---|---|---|---|---|---|---|---|---|---|
| 297934049 | | | | | 2 | | | | | | | | | | | | | | | | | | | |
| 297934346 | | | | | | | 1 | | | | | | | | | | | | | | | | | |
| 297938108 | | | | | | | | | | | | | | | | | | | 4 | | | | | |
| 297938585 | | | | | | | | | | 1 | | | | 15 | 12 | 12 | 14 | 10 | 8 | 9 | 1 | 17 | 8 | |
| 297947227 | | | | | | 1 | | 1 | | | | | | | 1 | | | | | | 1 | | 3 | |
| 298197305 | | | | | 1 | | 2 | | | | | | | | | | | | | | | | | |
| 298201943 | | | | | | 3 | 4 | | | | | | | | | | | | | | | | | |
| 298210543 | | | | | 2 | 2 | 4 | 5 | 6 | | | | | | 2 | | 2 | 1 | | | 1 | | | |
| 298245448 | | | | | 5 | | | | | | | | | | | | | | | | | | | |
| 298249566 | | | | | | 1 | | | | | | | | | | | | | | | | | | |
| 298249624 | | | | | 2 | | | | | | | | | | | | | | | | | | | |
| 298249959 | | | | | 4 | | | | | | | | | | | | | | | | | | | |
| 298254384 | | | | | | 1 | | | | | | | | | | | | | | | | | | |
| 298254517 | | | | | | | | 15 | 13 | | | | | | | | | | | | 1 | 10 | 10 | 1 |
| 298258134 | | | | | 1 | | | | | | | | | | | | | | | | | | | |
| 298258513 | | | | | | | | | 2 | | | | | | | | | | | | | | | |
| 298258530 | | | | | | | | | | | | | | | | | | | | | 1 | | | |
| 298262357 | | | | | | | 1 | | | | | | | | | | | | | | | | | |
| 298262408 | | | | | 2 | | | | | | | | | 1 | | 1 | 1 | 3 | 5 | 4 | | 1 | 3 | 2 |
| 298262442 | | | | | | | | | 2 | | | | | | | | | | | | | 1 | | |
| 298262498 | | | | | 1 | | | | | | | | | | | | | | | | | | | |
| 298262599 | | | | | | | | | | | 1 | | | | | | | | | | | | | |
| 298262644 | | | | | 1 | 6 | 3 | | | | | | | | | | | | | | 2 | 1 | 1 | |
| 298262663 | | | | | 1 | | 1 | | | | | | | | | | | | | | | | | |
| 298266660 | | | | | 1 | | | | | | | | | | | | | | | | | | | |
| 298266716 | | | | | | 2 | | | | | | | | | | | | | | | | | | |
| 298266805 | | | | | | 1 | | | | | | | | | | | | | | | | | | |
| 298266950 | | | | | | | 1 | | | | | | | | | | | | | | | | | |
| 298267120 | | | | | | 2 | | | | | | | | | | | | | | | | | | |
| 298270808 | | | | | 5 | 1 | 3 | | | | | | | | | | 1 | | 2 | 1 | | | | |
| 298270909 | | | | | 1 | | | | | | | | | | | | | | | | | | | |
| 298275024 | | | | | | 1 | | | | | | | | | | | | | | | | | | |
| 298275090 | | | | | | 1 | | | | | | | | | | | | | | | | | | |
| 298275356 | | | | | | | | 1 | | | | | | | | | | | | | | | | |
| 298279629 | | | | | 1 | 2 | 1 | | | | | | | | | | | | | | | | | |
| 298547436 | | | | | | 2 | 1 | | | | | | | | | | | | | | | | | |
| 298586567 | | | | | 3 | | 2 | | | | | | | | | | | | | | | | | |
| 298590924 | | | | | | 1 | | | | | | | | | | | | | | | | | | |
| 298591130 | | | | | 2 | | 1 | 1 | | | | | | | | | | | | | | | | |
| 298599269 | | | | | 1 | 1 | | | | | | | | | | | | | | | | | | |
| 298599310 | | | | | 2 | | | | | | | | | | | | | | | | | | | |
| 298603633 | | | | | | 1 | | 1 | | | | | | | | | | | | | | 1 | | |
| 298883439 | | | | | | 1 | 1 | | | | | | | | | | | | | | | | | |
| 298892294 | | | | | 1 | | | | | | | | | | | | | | | | | | | |
| 298931489 | | | | | 1 | | | | | | | | | | | | | | | | | | | |
| 298935769 | | | | | | | 1 | | | | | | | | | | | | | | | | | |
| 298936099 | | | | | | 2 | | | | | | | | | | | | | | | | | | |
| 298952944 | | | | | | 1 | | | | | | | | | | | | | | | | | | |
| 298957166 | | | | | | | | | | | | | | | | | | | | | | 1 | | |
| 298957344 | | | | | 4 | | | | | | | | | | | | | | | | | | | |
| 298962041 | | | | | | | | 1 | | | | | | | | | | | | | | | | |
| 299083134 | | | | | | 1 | | | | | | | | | | | | | | | | | | |
| 299281053 | | | | | 5 | | | | | | | | | | | | | | | | | | | |
| 299289475 | | | | | 1 | | | | | | | | | | | | | | | | | | | |
| 299289811 | | | | | | | | | | | | | | | | | | | | | | | 1 | |
| 299290105 | | | | | | | | 2 | | | | | | | | 1 | | | | | 4 | 1 | 2 | |
| 299613271 | | | | | 3 | 1 | | | | | | | | | | | | | | | | | | |
| 299617529 | | | | | | | | | 1 | | | | | | | | | | | | 3 | 9 | 11 | |
| 299617731 | | | | | 2 | | | | | | | | | | | | | | | | | | | |
| 299954299 | | | | | 1 | | | | | | | | | | | | | | | | | | | |
| 299954300 | | | | | 2 | | | | | | | | | | | | | | | | | | | |
| 300972942 | | | | | | | | | | | | | | | | | | | | 1 | | | | |
| 324846570 | | | | | | | | 6 | 13 | 37 | 25 | | | 1 | | | 1 | 2 | 3 | 1 | | | | |
| 325113795 | | | | | | | | | | | | | | 1 | | | | | | | | | | |
| 325118117 | | | 1 | | | 1 | | | | | | | | 1 | 1 | | | | | | | | | |
| 325122109 | | | | | | 1 | | | | | | | | | | | 1 | | 56 | 43 | | | | |
| 325122525 | | | | | | 2 | | | | | | | | 19 | 32 | 21 | 10 | 20 | | | | | | |
| 325122802 | | | | | | 1 | | | | | | | | | | | | | | | | | | |
| 325446181 | | | | | | | | | | | | | | | | | | | | | 1 | | | |
| 325455002 | | 1 | | | | | | | | | | | | 1 | 4 | 1 | 2 | 5 | | 1 | | | | |
| 325455473 | | | | | | | | | | | | | | | | 2 | | | | | | | | |
| 325460206 | | | | | | 1 | 1 | | | | | | | | | | | | | | | | | |
| 325472654 | | | | | | | 1 | | | | | | | | | | | | | | | | | |
| 325489440 | | | | | | 1 | | | | | | | | | | | | | | | | | | |
| 325490014 | | | | | | | | | | | | | | | | | | | | | 1 | | | |
| 325529237 | | | | | | 1 | | 5 | 8 | 30 | 33 | | | 2 | 2 | | | 1 | 1 | | | | 1 | |
| 325791855 | | | | | | | | | | | | | | | | | | | | | 2 | | | |
| 325814461 | | | | | | | 2 | | | | | 1 | 4 | | | | 1 | | | 1 | | | | |
| 325815440 | | | | | | | | | | | | 1 | | | | | | | | | 1 | | | |
| 325839288 | | | | | | | | 1 | | | | | | | | | | | | | 1 | | | |

| | | | | | | | | | | | | | | | | | | | | | | | | | | | | |
|---|---|---|---|---|---|---|---|---|---|---|---|---|---|---|---|---|---|---|---|---|---|---|---|---|---|---|---|---|
| 326119769 | | | | | | | | | | | | | | 1 | | | 2 | | | | | | 1 | | 2 | | | |
| 326124243 | | | | | | | | | | | | | | | | | | | | | | | | 1 | | | | |
| 326128670 | | | | | | | | | | 1 | | | | | | | | | | | | | | | | | | |
| 326134056 | | | | | | | | | | | | | | | | | | | | 1 | | | | | | | | |
| 326137566 | | | | | | | | | | 6 | 2 | | | | | | | | | | 1 | | | | | | | |
| 326159907 | | | | | | | | | | 1 | | | | | | | | | | | | | | | | | | |
| 326253554 | | | | | | | | | 2 | | | 2 | 2 | | | | | 1 | | | | 1 | 9 | 6 | 12 | | | |
| 326460989 | | | | | | | | | | | | 1 | | | | | | | | | | | | | | | | |
| 326465302 | | | | | | | | | | | 1 | | | | | | | | | 1 | | | | | | | | |
| 326474963 | | | | | | | | | | | | | | | | | | | | | | | | | 1 | | | |
| 326521082 | | | | | | | | | | | | | | | | | | | | | | | 2 | | | | | |
| 326525255 | | | | | | | | | | | | | | | | | | | | | | | 1 | | | | | |
| 326530038 | | | | | | | | | | | | | | | | | | | | | | | 1 | | | | | |
| 326556230 | | | | | | | | | 1 | | | | | | | | | 1 | | 2 | 1 | | 1 | 2 | 5 | | | |
| 326560782 | | | | | | | | | | | | 1 | | | | | | | | | | | | | | | | |
| 326812225 | | | | | | | | | | 1 | | | | | | | | | | | | | | | | | | |
| 326817510 | | | | | | | | | 2 | 12 | 11 | | | | | | | 1 | | | | | 13 | 7 | 1 | | 1 | |
| 326875404 | | | | | | | | | | | | | | | | | | | | | | | | 1 | | | | |
| 326884223 | | | | | | | | | | | | | | | | | | | | | | | | | 2 | | | |
| 326884250 | | | | | | | | | | 1 | 2 | | | | | | | | | | | | | | | | | |
| 326884264 | | | | | | | | | | 1 | | | | | | | | | | | | | | 1 | | | | |
| 326888433 | | | | | | | | | 1 | | 1 | | | | | | | | | | | | | | | | | |
| 326888609 | | | | | | | | | | | | | | | | | | | | | | | | | | 1 | | |
| 326888675 | | | | | | | | | 1 | | | | | | | | | | | | | | | | | | | |
| 327147292 | | | | | | | | | | | 1 | 10 | 5 | | | | | | | | | | | | | | | |
| 327147610 | | | | | | | | | | 1 | | | | | | | | | | | | | | | | | | |
| 327152465 | | | | | | | | | | 3 | | | | | | | | | | | | | | | 2 | | | |
| 327158831 | | | | | | | | | | | | | | | | | 2 | | | | | | | | 9 | | | |
| 327242169 | | | | | | | | | | 2 | 2 | | | | | | | | | | | | | | | | | |
| 327242287 | | | | | | | | | | | | 1 | | | | | | | | | 1 | 1 | | 1 | 1 | 2 | 1 | |
| 327246381 | | | | | | | | | | | 2 | | | | | | | | | | | | | | | | | |
| 327246931 | | | | | | | | | 7 | 3 | 6 | | 1 | | | | | 3 | 2 | | | | 10 | 18 | 1 | 1 | | |
| 327255181 | | | | | | | | | 1 | 1 | 2 | | | | | | | | | | | | | | | | | |
| 327255543 | | | | | | | | | | | 1 | | | | | | | | | | | | | | | | | |
| 327506825 | | | | | | | | | | | | | | | | | | | | | | | | | 1 | | | |
| 327509674 | | | | | | | | | | 1 | | | | | | | | | | | | | | | | | | |
| 327532246 | | | | | | | | | | 1 | | | | | | | | | | | | | | | | | | |
| 327566137 | | | | | | | | | | | | | | | | | | | | | | | 1 | | | | | |
| 327575026 | | | | | | | | | | | | | | | | | | | | | | | 1 | | | | | |
| 327587601 | | | | | | | | | | | 1 | | | | | | | | | | | | | | | | | |
| 327588074 | | | | | | | | | 21 | 9 | 20 | | | | | | | | | | | | 1 | | | | | |
| 327588208 | | | | | | | | | | | 1 | | | | | | | | | | | | | | | | | |
| 327588225 | | | | | | | | | | | | | 1 | | | | | | | | | | | | | 1 | | |
| 327592562 | | | | | | | | | | | | 1 | | | | | | 14 | 6 | 3 | 1 | 8 | 7 | 6 | | 1 | 1 | |
| 327596481 | | | | | | | | | | 1 | | | | | | | | | | | | | | | | | | |
| 327600366 | | | | | | | | | | | 1 | | | | | | | | | | | | | | | | | |
| 327843160 | | | | | | | | | 1 | 1 | 2 | | | | | | 2 | 2 | | | | | 1 | | 1 | | 1 | |
| 327848116 | | | | | | | | | | | | | | | | | | | | | | | | 1 | | | | |
| 327869598 | | | | | | | | | | | 1 | | | | | | | | | | | | | | | | | |
| 327869910 | | | | | | | | | | 1 | | | | | | | | | | | | | | | | | | |
| 327872615 | | | | | | | | | | | 3 | | | | | | | | | | | | | | | | | |
| 327928411 | | | | | | | | | | | | | | | | | | | | 2 | | | 3 | 3 | | | 2 | |
| 327933008 | | | | | | | | | | | | 1 | | | | | | | | | | | | | | | | |
| 327933027 | | | | | | | | | 1 | 1 | | | | 1 | | | | 1 | 2 | | | | | 1 | | | | |
| 327933814 | | | | | | | | | | | 2 | | | | 1 | | | 1 | | | | | | 1 | | | | |
| 327937328 | | | | | | | | | 4 | | | | | | | | | | | | | | | | | | | |
| 327937494 | | | | | | | | | 1 | | | | | | | | | | | | | | | | | | | |
| 327937506 | | | | | | | | | | | 1 | | | | | | | | | | | | 12 | 8 | | 1 | 1 | 3 |
| 328010530 | | | | | 3 | 2 | 2 | 3 | 2 | | 1 | 1 | 6 | 1 | 2 | | | 2 | | 2 | 3 | 1 | 4 | 14 | 6 | | 1 | 1 |
| 328179117 | | | | | | | | | | 1 | | 1 | 1 | | | | | | | | | | | | | | | |
| 328213227 | | | | | | | | | 1 | | | | | | | | | | | | | | | | | | | |
| 328257630 | | | | | | | | | | | | | | | | | | | | | | | | | 4 | | | |
| 328264983 | | | | | | | | | 1 | | | | | | | | | | | | | | | | | | | |
| 328265389 | | | | | | | | | | 1 | | 19 | 18 | | | | | | | | | | | 1 | 4 | 15 | 21 | 2 |
| 328273806 | | | | | | | | | 7 | 17 | 31 | 1 | | | | | | | | | | | | | 1 | | | |
| 328273858 | | | | | | | | | 2 | | | | | | | | | | | | | | | | | | | |
| 328274706 | | | | | | | | | 1 | 2 | | | | | | | | | | | | | | | | | | |
| 328278125 | | | | | | | | | 1 | 1 | | | | | | | | | | | | | | | | | | |
| 328278368 | | | | | | | | | 1 | | | | | | | | | | | | | | | | | | | |
| 328283528 | | | | | | | | | | | 1 | | | | | | | | | | | | | | | | | |
| 328291455 | | | | | | | | | 1 | | | | | | | | | | | | | | | | | | | |
| 328524867 | | | | | | | | | | | | 1 | | | | | | | | | | | | | | | | |
| 328533896 | | | | | | | | | | | | 1 | 1 | | | | | | | | | | | | | | | |
| 328537891 | | | | | | | | | | 1 | | | | | | | | | | | | | | | | | | |
| 328559607 | | | | | | | | | 11 | 1 | 1 | | | | | | | | | | | | | | | | | |
| 328563364 | | | | | | | | | | | | | | | | | | | | | | | 1 | | | | | |
| 328611467 | | | | | | | | | | 1 | | | | | | | | | | | | | | | | | | |
| 328632707 | | | | | | | | | | | | | | | | | | | | | | | | | 5 | 3 | | |
| 328718110 | | | | | | | | | | 1 | | | | | | | | | | | | | | | | | | |
| 328860948 | | | | | | | | | | | 1 | | | | | | | | | | | | | | | | | |

| | | | | | | | | | | | | | | | | | | | | | | | | |
|---|---|---|---|---|---|---|---|---|---|---|---|---|---|---|---|---|---|---|---|---|---|---|---|---|
| 328870472 | | | | | 1 | | | | | | | | | | | | | | | | | | | |
| 328883562 | | | | | | | 1 | | | | | | | | | | | | | | | | | |
| 328896613 | | | | | 3 | 1 | | | | | | | | | | | | | | | | | | |
| 328900127 | | | | | 1 | | | | | | | | | | | | | | | | | | | |
| 328905011 | | | | | | 3 | | | | | | | | | | | | | | | 1 | | 2 | |
| 328909309 | | | | 1 | | | | | | | | | | | | | | | | | | | | |
| 328943204 | | | | | | | | | | | | | | | | | | | | | | | | 1 |
| 328955898 | | | | | | | 1 | | | | | | | | | | | | | | | | | |
| 329227987 | | | | | | 1 | | | | | | | | | | | | | | | | | | |
| 329271369 | | | | | | 1 | | | | | | | | | | | | | | | | | | |
| 329301513 | | | | | | | | | 3 | | | | | | | | | | | | 2 | | 1 | |
| 329310230 | | | | | | | | 11 | 18 | | | | | | | | | | | 1 | 4 | 7 | 9 | 2 |
| 329561513 | | | | | | | | | | | | | | | | | | | | | 1 | | | |
| 329561679 | | | | | | 1 | | | | | | | | | | | | | | | | | | |
| 329566174 | | | | | | | | | | | | | | | | | | | | | 1 | | | |
| 329566197 | | | | | | | | | | | | | | | | | | | | | 1 | | | |
| 329732855 | | | | | | | | 5 | 5 | | | | | | | | | | | | | | 1 | 1 |
| 329928204 | | | | | | | 1 | | | | | | | | | | | | | | | | | |
| 329966674 | | | | | | | | | | | | 1 | | | | 1 | | 1 | | | | | | |
| 330074628 | | | | | | 1 | | | | | | | | | | | | | | | | | | |
| 330237771 | | | | | | | | 1 | | | | | | | | | | | | | | | | |
| 330246657 | | | | | 1 | 6 | 5 | | | | | | | | | | | | | | 2 | | | |
| 330316013 | | | | | | | | | | | | | | | | | | | | | 1 | 1 | | |
| 330415332 | | | | | | | | 21 | 18 | | | | | | | | | | | | 1 | | | |
| 330640044 | | | | | | | | | | | | | | | | | | | | | | | 1 | |
| 330661786 | | | | | | | | | | | | | | | | | | | | | 1 | | | |
| 330678844 | | | | | | | | | 1 | | | | | | | | | | | | | | | |
| 330998071 | | | | | 1 | | | | | | | | | | | | | | | | | | | |
| 331774819 | | | | | | | | | | | | | | | | | | | | | 5 | | | 1 |
| 355453590 | 2 | | 3 | 2 | | | | | | | | | | 7 | 15 | 26 | 15 | 13 | | | | | | |
| 355816896 | 1 | 4 | 3 | 2 | | 7 | 4 | 1 | | 1 | | | | | | 2 | | 1 | 2 | 1 | 7 | 2 | 3 | |
| 356136008 | | | | | 1 | 2 | | 1 | 2 | | | | | | | | 1 | | | | | | 1 | |
| 356140100 | | | | | | | | | | | | | | | | | | | | 2 | | | | |
| 356144721 | | | | | | 1 | | | | | | | | | | | | | | | | | | |
| 356153135 | | | | | | 1 | | | | | | | | | | | | | | | 1 | | 1 | |
| 356472645 | | | 1 | | | | | | 1 | | | 1 | | 1 | | | | | 46 | 54 | | | | |
| 356472660 | | | | | 1 | 1 | | 1 | | | 1 | | 1 | | | | 2 | | 7 | 4 | | | 1 | |
| 356490891 | | | | | | 1 | | | | | | | | | | | | | | | | | | |
| 356494648 | | | | | | 1 | | | | | | | | | | | | | 2 | | | | | |
| 356503889 | | | | | | 1 | 4 | | | | | | | | 3 | 1 | | 3 | | | | | | |
| 356517307 | | | | | | | | | | | | | | 1 | | | | | | | 1 | | | |
| 356813986 | | | | | | | | | | | | | | | 2 | | 1 | | | | | | | |
| 356818551 | | | | | | | | | 1 | | | 1 | | | | | | | 1 | | | 4 | 2 | |
| 356823065 | | | | | | | | 1 | | | | | | | | 2 | 1 | | | 1 | | | | |
| 356832459 | | | | | | 2 | | | | | | | | | | | | | | | | | | |
| 356835277 | | | | | | 2 | 2 | | | | | | | | | | | | | | 1 | | | |
| 357146296 | | | | | | | 1 | | | | | 1 | | | | | | | | | | | | |
| 357150583 | | | 2 | | | | | | | 4 | | 1 | 1 | 1 | | | | 1 | 7 | 3 | | 1 | 1 | |
| 357163689 | | | | | | 1 | | | | | | | | | | | | | | | | | | |
| 357482735 | | | | | | | | 3 | 4 | 21 | 11 | 3 | 3 | 3 | | 1 | 1 | 1 | 21 | 8 | 5 | 2 | 2 | |
| 357490931 | | | | | | | | | | | | | | | | | | | | | 3 | | | |
| 357515217 | | | | | | 3 | 1 | | | | | | | | | | | | | | 2 | | | |
| 357522097 | | | | | | | | | | | | | | | | | | | | | 4 | | | |
| 357522570 | | | | | | | | | | | | | | | | | | | | | 3 | | | |
| 357569116 | | | | | | 1 | | | | | | | | | | | | | | | | | | |
| 357569231 | | | | | | | | | | | | | | | | | | | 3 | 3 | | | | |
| 357569240 | | | | | | | | | | | | | | | | | | | 2 | | | | | |
| 357573154 | | | | | | | 2 | | | | | | | | | | | | | | | | | |
| 357581846 | | | | | | | | 1 | | | | | | | | | | | | | | | | |
| 357595428 | | | | | 1 | | | | | | | | | | | | | | 1 | | | | | |
| 357599074 | | | | | | | | | | | | | | | | | | | 8 | 11 | 57 | 1 | 4 | |
| 357607932 | | | | | | | | 3 | 4 | 26 | 17 | | | 2 | | 1 | | 4 | 1 | 1 | | | | |
| 357814777 | | | | | | | | | | | 1 | | | | | | | | | | | | | |
| 357823645 | | | | | | | 1 | | | | | 2 | | | | | | | | | | | | |
| 357832762 | | | | | | 2 | | | | | | | | | | | | | | | | | | |
| 357862532 | | | | | | 3 | 1 | | | | | | | | | | | | 3 | | | | | |
| 357863646 | | | | | | | | | | | | | | | | | | | | | 6 | | | |
| 357867646 | | | | | | 1 | | | | | | | | | | | | | | | | | | |
| 357909813 | | | | | | | | | | | | | | | | | | | | 2 | | | | |
| 357919199 | | | | | | | | | | | | | | | | | | | 1 | | | | | |
| 357923095 | | | 2 | | | | | | | | | | | | | | | | 6 | 3 | 2 | | 4 | |
| 357932060 | | | | | 1 | | | | | | | | | 1 | 2 | | | | 5 | 4 | | 1 | 3 | |
| 357940977 | | | | | | | | | | 1 | 1 | | | 2 | | | | 2 | | | 1 | 1 | 2 | |
| 357944721 | | | | | | | | | | | | | | | | | | | 1 | | | | | |
| 357944930 | | | | | | 1 | | 1 | 2 | | | | | | | | | | 1 | 5 | | | | |
| 357945095 | | | | | 3 | 1 | 4 | | 3 | | 1 | | | 1 | 1 | | | | | | | | | |
| 357945155 | | | | | | | | | | | | | | | | | | | | | 1 | | | |
| 357949102 | | | | | | | | | | | 1 | | | 1 | | | | | | | 3 | 1 | 5 | |
| 357949364 | | | | | 2 | | 2 | | | | | | | | | | | | | | | | | |
| 357953145 | | | | | 7 | 8 | 8 | | 1 | | | | | | 1 | | | | 1 | | | | | |

| | | | | | | | | | | | | | | | | | | | | | | |
|---|---|---|---|---|---|---|---|---|---|---|---|---|---|---|---|---|---|---|---|---|---|---|
| 358197600 | | | | | | | 1 | | | | | | | | | | | | | | | |
| 358264919 | | | | | | | | | | | 1 | 1 | | | | | | 1 | | 1 | | |
| 358281935 | | | | | | | | | | | | | | | | | | | | 2 | | 3 |
| 358281997 | | | | | 2 | 1 | | 1 | | | | | 1 | | | | | | | | | |
| 358285424 | | | | 7 | 11 | 16 | | 2 | | | | | | | | | | | | | | |
| 358286194 | | | | | | | 1 | 3 | | | | | | 1 | | | 1 | | 2 | | | 3 |
| 358286442 | | | 1 | | | | 1 | | | | | | | | | | | | | | | 1 |
| 358290727 | | | | | | | | | | | | | | | | | | | | | | 1 |
| 358290805 | | | | | | | | | | | | | | | | | | | | 2 | | |
| 358295017 | | | | | | 1 | | | | | | | | | | | | | | | | |
| 358501372 | | | | | | | 2 | | | | | | | | | | | | | | | |
| 358515278 | | | | | | | | | 1 | | | | | | | 1 | | 1 | | | 1 | |
| 358538040 | | | | | | | | | | | | | | | | | | | | 1 | | |
| 358593042 | | | | | | | | | | | | | | | | | | | | 3 | 1 | 2 |
| 358613806 | | | | | | | | | | | | | | | | | | | | 1 | | |
| 358631295 | | | | | 1 | 2 | | | | | | | | | | | | | 1 | | | |
| 358631384 | | | | 4 | 4 | 3 | 10 | 8 | | | | | | | | | | | | | | |
| 358631450 | | | | | | | | | | | | | | | | | | 1 | 2 | | | |
| 358631537 | | | | 4 | 5 | 12 | | 1 | | | | | | | | | | | | | | |
| 358639742 | | | | | | | 22 | 18 | | | | | | | | | | | | | | |
| 358639884 | | | | 3 | 11 | 7 | | | | | | | | | | | | | 1 | | 1 | |
| 358847551 | | | | | | | 1 | 1 | | | | | | | | | | | | 14 | 7 | 3 |
| 358865451 | | | | | | | | | | | 1 | | | | | | | | | | | |
| 358878546 | | | | | | | | | | | | | | | | | | | | 3 | | |
| 358899943 | | | | | | 1 | | | | | | 1 | | | | | | | | | | |
| 358976440 | | | | 1 | 1 | | 1 | | 1 | | | | 2 | 1 | | | | | 1 | | | |
| 358976515 | | | | 3 | 6 | 10 | 1 | | | | | | | | | | | | | | | |
| 358985893 | | | | | 3 | | 2 | | | | | | | | | | | | | | | |
| 359210479 | | | | | | | 1 | | | | | | | | | | | | | | | |
| 359219263 | | | | | | | | | | | | | | | | | | | | 1 | | |
| 359279388 | | | | | | | | | | | | | | 1 | 2 | 1 | | | | | | |
| 359326692 | | | | 5 | 10 | 9 | 1 | | | | | | 1 | | | | | | 1 | | | |
| 359326764 | | | | 2 | | | | | | | | | | | | | | | | | | |
| 359447314 | | | | | 2 | | | | | | | | | | 1 | | | 2 | | | | |
| 359555152 | | | | | | | | | | | | | | | | | | | | 1 | | |
| 359744616 | | | | | 2 | | | | | | | | | | | | | | | | | |
| 359748709 | | | | 1 | | | | | | | | | | | | | | | | | | |
| 359897526 | | | | | 1 | | | | | | | | | | | | | | | | | |
| 360086302 | | | | | | | 1 | | | | | | | | | | | | | | | |
| 360245971 | | | | | 2 | | | | | | | | | | | | | | | | | |
| 360254108 | | | | 2 | | | | 1 | | | | | | | | | | | | 1 | | |
| 360284300 | | | | | | | | | | | 1 | 1 | | | | | | | | | 1 | 1 |
| 360427529 | | | | 1 | 1 | | | | | | | | | | | | | | | | | |
| 360432213 | | | | 2 | | | | | | | | | | | | | | | | | | |
| 360596183 | | | | | | | | | | | 1 | | | | | | | | | | | |
| 360677632 | | | | | | | | | | | | | | | | | | | | 1 | | |
| 360690414 | | | | | 1 | | | | | | | | | | | | | | | | | |
| 360698817 | | | | | | 1 | | | | | | | | | | | | | | | | |
| 360763630 | | | | | 1 | | | | | | | | | | | | | | | | | |
| 360764170 | | | | 1 | | | | | | | | | | | | | | | | | | |
| 361005850 | | | | | | | | 1 | | | | | | | | | | | 1 | 4 | 4 | 10 |
| 361104677 | | | | | 1 | | | | | | | | | | | | | | | | | |
| 361312146 | | | | | 2 | 1 | | | | | | | | | | | | | | | | |
| 361363973 | | | | | | | | | | | | | | | | | | | | 1 | | |
| 361445621 | | | | | | | 2 | | | | | | | | | | | | | | | |
| 361636071 | | | | | 1 | 1 | | | | | | | | | | | | | | | | |
| 362127690 | | | | | 1 | | | | | | | | | | | | | | | | | |
| 362394990 | | | | | | | 1 | | | | | | | | | | | | | | | |
| 362473525 | | | | | | | 14 | 17 | | | 2 | 3 | | | | | | | 1 | 4 | 3 | 2 |
| 362719326 | | | | | | | 2 | | | | | | | | | | | | | 1 | | |
| 386506469 | | | | 1 | 1 | | | | | | | | | | | | | | | | | |
| 386821054 | | | | | 1 | | | | | | | | | | | | | | | | | |
| 386829647 | | | | | | | | | | | | | | | | | | | 1 | | | |
| 386833812 | | | | 1 | | | | | | | 6 | 6 | 3 | 1 | | | | 4 | 1 | 2 | | 1 |
| 386833850 | 1 | 1 | | | 1 | | 7 | 2 | | | | | 1 | | | | | 24 | 41 | | 3 | |
| 386834269 | 3 | | | | 1 | 2 | 3 | 4 | 4 | 2 | | | 3 | | 1 | 3 | 1 | | 5 | 8 | 1 | |
| 386838513 | | | | | | | 1 | | | | 6 | 3 | | | | | | | | 2 | | |
| 386838814 | | | | | 1 | | | | | | | | | | | | | | | | | |
| 386842280 | | | | | | | | | | | | 1 | | | | | | | | | | |
| 386847799 | | | | | | | | | | | | | | | | | | | | | 1 | |
| 386872886 | | | | | | | | | | | | | 3 | | | | | | | | | |
| 387166379 | | | | | | | 8 | 5 | 30 | 25 | | | 1 | 1 | 2 | 1 | | | 1 | | 1 | |
| 387170738 | | | | | | | | | | | | 1 | | | | | | | | 1 | 1 | |
| 387180498 | | | | | 1 | | 3 | 2 | | | 1 | | 2 | 4 | 2 | 1 | 2 | | | 1 | 1 | |
| 387215056 | | | | | | | | | | | | | 2 | | | | | | | | | |
| 387503511 | | | | | | | | | | | | | | | | | | | | | 1 | 1 |
| 387870860 | | | | | | | | | | | | 3 | | | | | | | | | | |
| 387943673 | | | | | | | 1 | | | | | | | | | | | | | | | |
| 387944118 | | | | | | | 2 | 2 | 34 | 22 | | | 1 | 1 | | | | | | | | 2 |
| 387952104 | | | | | | | | 1 | | | | 1 | | | | | | 1 | | | | |

| | | | | | | | | | | | | | | | | | | | | | | | | | | | | |
|---|---|---|---|---|---|---|---|---|---|---|---|---|---|---|---|---|---|---|---|---|---|---|---|---|---|---|---|---|
| 388284324 | | | | | | | | | | | | | | | | | | | | | | | | | 2 | | | |
| 388289013 | | | | | | | | | | | | | | | | | | | | | | | | | | | | 1 |
| 388293131 | | | | | | | | | | | | 1 | 1 | | | | | | | | | | | | | | | |
| 388297625 | | | | | | | | | | | | 1 | 1 | | 5 | | | | | | | 1 | 2 | 1 | | | 4 | |
| 388625791 | | | | | | | | | | | | | | | | | | | | | | | 1 | | | | | |
| 388638672 | | | | | | | | | 2 | | | 48 | 22 | | | | | 4 | 2 | | | 3 | | | 1 | 1 | 1 | |
| 388854495 | | | | | | | | | | | 1 | | 1 | | | | | | | | | | | | | | | |
| 388859011 | | | | | | | | | | 1 | | | | | | | | | | | | | | | | | | |
| 388877336 | | | | | | | | | | | | | | | | | | | | | | | 1 | | | | | |
| 388881226 | | | | | | | | | | 1 | | | | | | 19 | 10 | | | | | | | | 1 | | | |
| 388954086 | | | | | | | | | | | | 1 | | | | | | | | | | | | | 2 | | 2 | |
| 388975834 | | | | | | | | | | | 2 | | | | | | | | | | | | | | | 1 | 1 | |
| 388984132 | | | | | | | | | | | | 3 | 5 | | 2 | | | | | | | | 2 | 1 | 19 | 25 | 5 | |
| 389195896 | | | | | | | | | | | | | 1 | 1 | | | | 1 | | | | | 2 | 4 | | | | |
| 389199936 | | | | | | | | | | | | 9 | 3 | | | 13 | 21 | 11 | 3 | | 5 | 10 | | 1 | 1 | | | |
| 389204929 | | | | | | | | | | 2 | | | | | | | | | | | | | | | | | | |
| 389217959 | | | | | | | | | | 1 | | | | | | | | | | | | | | | | | | |
| 389251578 | | | | | | | | | | | 1 | | | | | | | | | | | | | | | | | |
| 389325067 | | | | | | | | | 1 | 2 | | | | | | | | | | | | | | | | | | |
| 389527383 | | | | | | | | | | | | | | | | | | | | | 1 | | 2 | 2 | 1 | | | |
| 389536926 | | | | | | | | | | | | | | | | | 1 | | | | | 6 | | | 15 | 5 | 1 | |
| 389545760 | | | | | | | | | | | | 3 | | | | | | | | | | | | | | | | |
| 389546522 | | | | | | | | | | | | | 1 | | | | | | | | | | | | | | | |
| 389563849 | | | | | | | | | | | 1 | | | | | | | | | | | | 4 | 3 | | | | |
| 389657555 | | | | | | | | | 1 | | | | 3 | | | 1 | | | | | | | | | 3 | 10 | 10 | |
| 389687699 | | | | | | | | | 3 | | 3 | | | | | | | | | | | | | | | | | |
| 390024952 | | | | | | | | | | 1 | | | | | | | | | | | | | | | 1 | | | |
| 390097431 | | | | | | | | | 1 | | | | | | | | | | | | | | | | | | | |
| 390271275 | | | | | | | | | | | | | | | | | | | | | | | | | 3 | | | |
| 390279635 | | | | | | | | | | | | | | | | | 1 | | | | | | | | 1 | | | |
| 390331583 | | | | | | | | | | | | 7 | 7 | | | | | | | | | | | | | | 1 | |
| 390344371 | | | | | | | | | | | | | 1 | | | | | | 1 | | | 1 | | 1 | 5 | | | |
| 390356988 | | | | | | | | | | | 1 | | | | | | | | | | | | | | | | | |
| 390366259 | | | | | | | | | 1 | | | | | | | | | | | | | | | | | | | |
| 390378650 | | | | | | | | | | | | | | | | | | 1 | 1 | 2 | 1 | | 4 | 1 | | | | |
| 390442730 | | | | | | | | | 2 | 1 | | | | | | | | | | | | | | | | | | |
| 390443308 | | | | | | | | | 1 | | | | | | | | | | | | | | | | | | | |
| 390560212 | | | | | | | | | | | | | | | | | | 1 | 1 | | | | | | | | | |
| 390579059 | | | | | | | | | | | | | | | | | | | | | | | | | 1 | | | |
| 390590450 | | | | | | | | | | | | | | | | | | | | | | | | | 1 | 7 | 1 | |
| 390641665 | | | | | | | | | | 1 | | | | | | | | | | | | | | | 1 | | 1 | |
| 390702220 | | | | | | | | | 1 | | | | | | | | | | | | | 1 | 1 | | | | | |
| 390706983 | | | | | | | | | 1 | 1 | 1 | | | | | | | | | | | | | | | | | |
| 390779805 | | | | | | | | | | 2 | | | | | | | | | | | | | | | | | | |
| 390784054 | | | | | | | | | | 1 | | | | | | | | | | | | | | | | | | |
| 390918452 | | | | | | | | | | | | | 1 | | | | | | | | | | | | | | | |
| 391026123 | | | | | | | | | | 1 | | | | | | | | | | | 2 | | | | 1 | 1 | | |
| 391034580 | | | | | | | | | | | | | | | | | | 5 | 2 | | 3 | 3 | | 1 | | | | |
| 391052159 | | | | | | | | | | | | | 2 | | | | | | | | | | | | | | | |
| 391120561 | | | | | | | | | | 5 | | | | | | | | | 1 | | | | | | | | | |
| 391362634 | | | | | | | | | | | | 1 | | | | | | | | | | | | | | | | |
| 391462340 | | | | | | | | | 1 | | | | | | | | | | | | | | | | | | | |
| 391802788 | | | | | | | | | 1 | | | | | | | | | | | | | | | | | | | |
| 391971839 | | | | | | | | | | | | | | | | | | | | | | | | | 1 | | | |
| 392057736 | | | | | | | | | | | | | | | | | | | | | | | | | | 1 | | |
| 392130578 | | | | | | | | | | | | | | | | | | | | | | | | | 3 | | | |
| 392325925 | | | | | | | | | | 1 | 5 | | | | | | | | | | | | | | | | | |
| 392411222 | | | | | | | | | | | | | | | | | | | | | | | | | | | 1 | |
| 392490012 | | | | | | | | | 1 | | | | | | | | | | | | | | | | | | | |
| 392821837 | | | | | | | | | | 8 | 5 | | | | | | | | | | | | | | | | | |
| 392822059 | | | | | | | | | | | | 1 | 1 | | | | | | | | | | | | | | | |
| 393063351 | | | | | | | | | | | | 1 | 2 | | | | | | | | | | | | 10 | 11 | 12 | 6 |
| 393776005 | | | | | | | | | | | | | | | | | | | | | | | | | 36 | 1 | 5 | 5 |
| 417143726 | | | | | | | | | | | | 1 | 2 | | | | | | | | | | | | 14 | 20 | 18 | |
| 417190793 | | 1 | | | | | | | | 1 | | | | | | 1 | | 1 | 1 | 3 | 4 | 5 | | | 1 | 1 | | |
| 417199910 | | | | | | | | | | | | | | | | | | 1 | | | | | 4 | 1 | | | | |
| 417204164 | | | | | | | | | | | | | | | | | | | | | | | 1 | | | | | |
| 417204354 | | | | | | | | | | | | | | | | 5 | 7 | | 1 | | | | 6 | 6 | 1 | 1 | 4 | |
| 417221397 | | | | | | | | | | | | | | | | | | 1 | | | | | | 1 | | | | |
| 417536905 | | | | | | | | | 1 | 2 | 1 | | | | | | | | 1 | | | | | 1 | | | | |
| 417540543 | | | | | | | | | | | | | | | 2 | 5 | 6 | | | 1 | 1 | 3 | | 1 | 1 | | | |
| 417550216 | | | | | | | | | | | | 10 | 9 | 1 | | | 1 | 1 | | | 1 | | 1 | 1 | 4 | | | |
| 417554004 | | | | | | | | | | | | | | | | | | | | | | | | 2 | | | | |
| 417554241 | | | | | | | | | | | | | | | | 1 | | | | | | | | | | | | |
| 417558532 | | | | | | | | | | | | | 1 | | | 14 | 13 | | | | | | | | | | | |
| 417882515 | | | | | | | | | | | | | | | | | | | | | 1 | | | | | | | |
| 417890128 | | | | | | | | | | | | | | | | 8 | 14 | | | | | | | 2 | | | | |
| 417908800 | | | | | | | | | | | | 6 | 8 | | | | | | | | | | | | 1 | 1 | | |
| 418205694 | | | | | | | | | | 4 | | | 1 | | | | | 2 | 3 | | 2 | 2 | | | | | | |
| 418214549 | | | | | | | | | | | | | | | | | | | | | | | | | 1 | | | |

| | | | | | | | | | | | | | | | | | | | | | | | | | | | | |
|---|---|---|---|---|---|---|---|---|---|---|---|---|---|---|---|---|---|---|---|---|---|---|---|---|---|---|---|---|
| 418224293 | | | | | | | | | | | | | | | | | | | | | | 1 | | | | 3 | | |
| 418224351 | | | 2 | | | | | | | | | 1 | | | | | | | | | | | 1 | | 3 | 1 | | |
| 418261858 | | | | | | | | | | | 1 | | | | | | | | | | | | | | | | | |
| 418607328 | | | | | | | | | | | | | | | | | | | | | | | | | 2 | | | |
| 418641700 | | | | | | | | | | | 1 | 1 | | 1 | | | | | | | | | 2 | 3 | 1 | 2 | 3 | |
| 418650351 | | | | | | | | | | | | 14 | 15 | 4 | 4 | | | 7 | 6 | 7 | 7 | 7 | | | | | | |
| 418870503 | | | | | | | | | | | | | | | | | | | | | | | | | 1 | | | |
| 418888180 | | | | | | | | | | 1 | | | | | | | | | | | | | | | | | | |
| 418991065 | | | | | | | | | | | | | | | | | | | | | | | | 1 | | | | |
| 419000001 | | | | | | | | | | | | 11 | 5 | 18 | 19 | | | 1 | | | 2 | 4 | | | | 4 | 1 | |
| 419345606 | | | | | | | | | | | | 4 | 1 | | 2 | | | 3 | | | | | | | 15 | 55 | 56 | |
| 419466429 | | | | | | | | | 1 | | | 5 | 6 | 1 | 1 | | | | | | | 1 | | | 19 | 1 | 2 | 34 |
| 419669180 | | | | | | | | | | | | | | | | | | 1 | 4 | | 1 | 2 | 7 | | | | | |
| 419908058 | | | | | | | | | | | | 2 | | | | | | | | | | | | | | | | |
| 420010560 | | | | | | | | | | | | | | | | | | | | | 1 | | | | | 1 | | |
| 420027427 | 1 | | 1 | | | | | | | 2 | 3 | | 1 | | | | 1 | 2 | 2 | | | 1 | 1 | | | 1 | | |
| 420036032 | | | | | | | | | | | 1 | | | | | | | 1 | | | | | | | | | | |
| 420036071 | | | | | | | | | | | | | | | | | | 1 | | | | | | | | | | |
| 420221276 | | | | | | | | | | 2 | | 2 | 3 | | | 1 | 1 | 3 | 4 | 3 | 5 | 2 | 1 | | 1 | | | |
| 420239937 | | | | | | | | | | | | 3 | | | | | | | | | | | | | | | | |
| 420347328 | | | | | | | | | 1 | | | 1 | | | | | | | | | | | 2 | 14 | 4 | 2 | 3 | |
| 420347360 | | | | | | | | | | 2 | | | | | | | | | | | | | | | | | | |
| 420351087 | | | | | | | | | | | | | | | | | | | | | | | | | | | 1 | |
| 420351828 | | | | | | | | | 1 | | 1 | 3 | 2 | | | | | | | | | | | | | 1 | 1 | |
| 420382647 | | | | | | | | | | | | 2 | | | | | | 2 | 2 | 4 | 2 | 1 | 11 | 5 | | | | |
| 420386244 | | | | | | | | | | | | | | | | | | | | | | | | | | 1 | | |
| 420455015 | | | | | | | | | | | | | | | | | | | 1 | | | 1 | | | | | | |
| 420567894 | | | | | | | | | | | | | | | | | | | | | | | | | 2 | | | |
| 420581268 | | | | | | | | | | | | 1 | | | | | | | | | | | | | | | | |
| 420586101 | | | | | | | | | | | | 1 | | | | | | | | | | | | | | | 1 | |
| 420666689 | | | | | | | | | | 1 | | | | | | | | | | | | | | | | | | |
| 420799910 | | | | | | | | | | | 1 | | | | | | | 3 | | 2 | 4 | 2 | 3 | | | | | |
| 420908078 | | | | | | | | | | | | | | | | | 1 | | | | | | | | | 1 | | |
| 421054850 | 1 | | | | | | | | 1 | | | | | | | | | | | | | | | | | | | |
| 421055181 | | | | | | | | | | | 1 | | | | | | | | | | | | | | | | | |
| 421405157 | | | | | | | | | | | | | | | | | | | | | | | | | | 1 | | |
| 421477693 | | | | | | | | | | 1 | | | | | | | | | | | | | | | | | | |
| 421477870 | | | | | | | | | | | | | | 1 | | | | | 1 | | | | | | | | | |
| 421605182 | | | | | | | | | 1 | | 2 | | | | | | | | | | | | | | | | | |
| 421814868 | | | | | | | | | | 1 | | | | | | | | | | | | | | | | | | |
| 421819939 | | | | | | | | | | 1 | | | | | | | | | | | | | | | | | | |
| 421946059 | | | | | | | | | | 1 | | | | | | | | | | | | | | | 1 | | | |
| 422151816 | | | | | | | | | | 1 | | | | | | | | | | | | | | | | | | |
| 422156333 | | | | | | | | | | 1 | | | | | | | | | | | | | | | | | | |
| 422286752 | | | | | | | | | | 1 | | | | | | | | | | | | | | | | | | |
| 422419397 | | | | | | | | | | | | | | | 1 | | | | | | | | | | | 1 | | |
| 422622063 | | | | | | | | | | | | | 1 | | | | | | | | | | | | | 1 | | |
| 422674339 | | | | | | | | | | 1 | | | | | | | | | | | | | | | | | | |
| 422751191 | | | | | | | | | | | | | | | | | | | | | | | | | 1 | | | |
| 423101189 | | | | | | | | | | | | 3 | 2 | | | | | | | | | | | | 18 | 32 | 35 | |
| 423852469 | | | | | | | | | | | | | | | | | | | | | | | | | 1 | | | |
| 424793122 | | | | | | | | | | | 1 | | | | | | | | | | | | | | | | | |
| 425143700 | | | | | | | | | | | 1 | | | | | | | | | | | | | | | | | |
| 425148091 | | | | | | | | | | | | | | | | | | | | | | | | | 33 | 5 | 3 | 7 |
| 425484071 | | | | | | | | | 2 | | | | | | | | | | | | | | | | | | | |
| 425790257 | | | | | | | | | | | | | | | | 1 | | | | | | | | | 1 | | | |
| 448182402 | | | 2 | | | | | | | | | | 1 | | 1 | 1 | | | | | | | 1 | | | | | |
| 448234999 | | | | | | | | | 1 | | 1 | | | | | | | 1 | | 1 | | 2 | | | | | | |
| 448238871 | | | | | | | | | 1 | | | | 1 | 1 | 1 | | | | 1 | | 1 | | | | | | | |
| 448260940 | | | | | | | | | 3 | | 3 | | | | | | | | | | 1 | | | | | | | |
| 448515267 | | | | | | | | | 2 | | | | | | | | | | 1 | | | | | | 6 | 2 | | |
| 448908723 | | | | | | | | | | | | | 1 | | | | | | | | | | | | | | | |
| 448912981 | | | | | | | | | | | | 2 | 1 | | | | | | | | | | | | 1 | | | |
| 448942604 | | | | | | | | | | 1 | | | | | | | | | | | | | | | | | | |
| 448943882 | | | | | | | | | | 1 | | | | | | 3 | 2 | | | | | | | | | 1 | | |
| 448994377 | | | | | | | | | | | | | | | | | 1 | | | | | | | | | | | |
| 449250041 | | | | | | | | | | | | | | | | | | | | | | | | | | 1 | | |
| 449288517 | | | | | | | | | | | | | | | | | | | | | | | | | 4 | | | |
| 449289591 | | | | | | | | | | 1 | | | | | | | | | | | | | | | | 1 | | |
| 449331241 | 2 | | | 1 | | | | | | | | | | | | | | 5 | 6 | 7 | | 7 | 28 | 8 | | | 1 | |
| 449335798 | | | | | | | | | | | | | | | | | | 4 | 2 | 1 | | 5 | 2 | 2 | 1 | 3 | 7 | |
| 449339532 | | | | | | | | | | | | | | | | | | | 1 | | 1 | | | | | | | |
| 449339838 | | | | | | | | | | | 1 | | | | | | | 2 | | | 1 | | | | | 1 | | |
| 449490682 | | | | | | | | | | | | 1 | | | | | | | | | | | | | | | | |
| 449490694 | | | | | | | | | | | | | | | 2 | | | | | 1 | | | | | | | | |
| 449555633 | | | | | | | | | | | | | | | | | | 2 | | | | | | | | | | |
| 449568880 | | | | | | | | | | 1 | | | | | | | | | | | | | | | | | | |
| 449634200 | | | | | | | | | | 1 | | | | | | | | | 1 | 1 | 4 | 4 | | | | | 1 | |
| 449634539 | | | | | | | | | | | | | | | | 1 | 1 | | | | | | | | | 1 | | |
| 449668199 | | | | | | | | | | | 1 | | | | | | | | | | | | 1 | | | | | |

| | | | | | | | | | | | | | | | | | | | | | | | | | | | | |
|---|---|---|---|---|---|---|---|---|---|---|---|---|---|---|---|---|---|---|---|---|---|---|---|---|---|---|---|---|
| 449681157 | | | 1 | 1 | | | | | | | | | | 1 | | 3 | 1 | 3 | | | | | 1 | | 13 | 9 | 12 | |
| 449685246 | | | | | | | | | | | | | | | | | | 1 | | | | | | | | | | |
| 449685778 | | | | | | | | | | | | | 2 | | | | | | | | | | | | | | | |
| 449901277 | | | | | | | | | | 1 | 1 | | | | | | | | | | | | | | | | | |
| 449905648 | | | | | | | | | | | | | | | | | | | | | | | | | | 1 | | |
| 449910481 | | | | | | | | | 2 | | | 6 | 10 | | | | | 1 | 2 | 1 | 2 | 4 | 6 | 2 | | | 1 | |
| 449910491 | | | | | | | | | | | 1 | | | | | | | | | | | | | | | | | |
| 449975227 | | | | | | | | | | 1 | | | | | | | | | | | | | | | 1 | | 2 | |
| 450008671 | | | | | | | | | | | | | | | | | 1 | | | | 1 | | 1 | | | | | |
| 450026507 | 1 | | | 1 | | | | | | | | | | 1 | | | | 6 | 4 | 4 | 2 | 3 | 7 | 15 | | | | |
| 450034902 | | | | | | | | | | | | 2 | 2 | | 1 | | | 1 | | 1 | 1 | 1 | 1 | 1 | 2 | | 30 | |
| 450047573 | | | | | | | | | | | | 3 | | | 2 | | | | | | | 1 | | | | | | |
| 450155802 | | | | | | | | | | | | | | | | | | | | 1 | | | | | | | | |
| 450160102 | | | | | | | | | | | | | | | | | | | | 1 | | | | | | | | |
| 450285679 | | | | | | | | | | 1 | | | | | | | | | | | | | | | 1 | | | |
| 450354788 | | | | | | | | | | | | 1 | | | 1 | | 1 | | | | | | 1 | | 1 | 1 | 1 | |
| 450371612 | | | | | | | | | | | | | | | | | | | | | | | | | 1 | | | |
| 450497356 | | | | | | | | | | | | | | | | | | | | | | | | | | | 1 | |
| 450583886 | | | | | | | | | | | | 20 | 22 | 20 | 12 | 18 | 11 | | 1 | | | | | | 25 | 7 | 5 | |
| 450597298 | | | | | | | | | | | | | | | | | | | | | | | | | | | 2 | |
| 450700075 | | | | | | | | | | | 2 | 4 | 2 | | 1 | | | 1 | 1 | | | | | 1 | 5 | | | |
| 450807268 | | | | | | | | | | | | | | | | | | | | | | | | | | | | 1 |
| 451040974 | | | | | | | | | | | | | | | | | | | | | | | | | 7 | 4 | 4 | |
| 451054039 | | | | | | | | | | | | | | | | | | | | | | | 6 | 3 | 27 | 1 | | 28 |
| 451071417 | | | | | | | | | | | | | | | | | | 2 | | | | | | | | 1 | 1 | |
| 451080648 | | | | | | | | | | | | | | | | | | | 1 | | | | | | | 2 | 4 | |
| 451084400 | | | | | | | | | | | | | | | | | | | | | 1 | | | | | | | |
| 451148412 | | | | | | | | | | | | | | | | | | | 1 | | 2 | | | | | | | |
| 451279196 | | | | | | | | | | | | 1 | | | | | | 1 | | | | 1 | 1 | 1 | | 1 | | |
| 451365099 | | | | | | | | | | | | | | | | | | | | | | | | | | 1 | | |
| 451382570 | | | | | | | | | | | | | | | | | | | | | | | | | | 1 | | |
| 451390676 | | | | | | | | | | | | | 1 | | | | | | | | | | | | 5 | | | |
| 451412500 | | | | | | | | | | | 1 | | | | | | | | | | | | | | | | | |
| 451424866 | | | | | | | | | | | | 5 | 2 | 8 | 4 | 1 | | 2 | | | | 1 | | 1 | 2 | | 1 | |
| 451485030 | | | | | | | | | | | | | | | | | | | | | | | | | | 1 | | |
| 451485461 | | | | | | | | | | 1 | | | | | | | | | | | | | | | | | | |
| 451489835 | | | | | | | | | | | | | | | | | | 1 | | 1 | | 1 | | | | | | |
| 451602687 | | | | | | | | | | | | | | | | 1 | 2 | | | | | | | | | | | |
| 451662040 | | | | | | | | | | | | 4 | | | | | | | | | | | | | 1 | | | |
| 451689001 | | | | | | | | | | | | | | | | | | 1 | | | 2 | 3 | | | | | | |
| 451722668 | | | | | | | | | | | | | 1 | | | | | | | | | | | 1 | 1 | | | |
| 451762072 | | | | | | | | | | | | | | | 2 | | | | 1 | | 1 | 3 | 1 | | | 15 | 8 | |
| 452167805 | | | | | | | | | | 1 | | | | | | | | | | | | | | | | | | |
| 452171673 | | | | | | | | | | 4 | | | | | | | | | | | | | | | | | | |
| 452401052 | | | | | | | | | | 3 | | | | | | | | | | | | | | | | | | |
| 452504314 | | | | | | | | | | | | | | | | | | | | | 1 | | | | | | | |
| 452771533 | | | | | | | | | | | 1 | 1 | | | | | | 1 | | 1 | 2 | | 2 | | | 4 | | |
| 452771913 | | | | | | | | | | | | 1 | 6 | | | | | 1 | 5 | 1 | | 1 | | | | | | |
| 452841019 | | | | | | | | | | | | 1 | 1 | | | | | | | | | | | | | | | |
| 452953198 | | | | | | | | | | | | 4 | 1 | 1 | | | | | | | | | | | | | | |
| 452971138 | | | | | | | | | | | | | | | | | | | | | | | | 1 | | | | |
| 453009665 | | | | | | | | | | 1 | | | | | | | | | | | | | | | | | | |
| 453108601 | | | | | | | | | | | | | | | | | | | | | | | | | 1 | | | |
| 453130054 | | | | | | | | | | | | 18 | 19 | | | | | | | | | | | | | | | 1 |
| 453185918 | | | | | | | | | | 6 | | | | | | | | | | | | | | | | | | |
| 453320065 | | | | | | | | | | | | 3 | 1 | | | | | | | | | | | | | | | |
| 453436589 | | | | | | | | | | | | | | | | | | 1 | | | | 2 | | | | | 2 | |
| 453457927 | | | | | | | | | | | 2 | | | | | | | | | | | | | | | | | |
| 453467404 | | | | | | | | | | | 1 | | | | | | | | | | | | | | | | | |
| 453527730 | | | | | | | | | | | | 21 | 16 | 1 | | | | | | | | | | | | 2 | 1 | 1 |
| 454386099 | | | | | | | | | | | | 2 | 1 | | | | | | | | | | | | | | | |
| 454481372 | | | | | | | | | | | | | | | | | | | | | | | | | 8 | | | |
| 454887480 | | | | | | | | | 1 | | | | | | | | | | | | | | | | | | | |
| 454935401 | | | | | | | | | | | | | | | | | | | | | | | 1 | | | | | |
| 455172836 | | | | | | | | | | 1 | | | 2 | | | | | | | | | | | | | | | |
| 455513316 | | | | | | | | | | | | | | | | | | | | | | | | | 1 | | | 1 |
| 455910660 | | | | | | | | | 1 | | | | | | | | | | | | | | | | | | | |
| 477918443 | | | | 1 | | | | | | | | | | | 1 | | | | | | | | | | | | | |
| 478618011 | | | | | | | | | | | | | | | | | | | | | | | 1 | 1 | | | | |
| 478928628 | | 1 | | | | | | | | | | | | | | 84 | 76 | | | | | | | | | | 4 | |
| 478937243 | | | | | | | | | 4 | 3 | 17 | | | | | | | | | | | | | | | | | |
| 479260921 | | | | | | | | | 4 | 1 | 1 | 1 | | | | | | | | | | | 1 | 2 | | | | |
| 479269268 | | | | | | | | | | | | | | | | | | | 1 | | | | | | | | | |
| 479290982 | | | | | | | | | | | | | | 2 | | 9 | 10 | 1 | | | | 1 | 1 | 1 | 1 | 1 | | |
| 479637825 | | | | | | | | | | | | | 1 | | | | | | | | | | | | | | | |
| 479645393 | | | | | | | | | | | | | 3 | | | | | | | | | | | | | | 1 | |
| 479908437 | | | | | | | | | | | | 2 | | | | | | | | | | | | | | | | |
| 479912666 | | | | | | | | | | | | 1 | | | | | | | | | | | | | | | | |
| 479921537 | | | | | | | | | | | | | | | | | 1 | | | | | | 1 | | | 1 | | |
| 479987085 | | | | | | | | | | | 3 | 1 | 4 | | | 1 | 1 | | | | 1 | | 7 | 4 | 1 | 2 | 3 | |

| | | | | | | | | | | | | | | | | | | | | | |
|---|---|---|---|---|---|---|---|---|---|---|---|---|---|---|---|---|---|---|---|---|---|
| 480029788 | | | | | | 2 | | | | | 1 | | | 1 | | 1 | | 3 | 45 | | |
| 480276374 | | | | | | | | | | | | | | | | | | 2 | | | |
| 480328501 | | | | | | | | | | | | | | | | | | 1 | | | |
| 480331310 | | | | | | | | | 2 | 1 | | | | | | | | | | | |
| 480371354 | | | | | | | | | | | | | | 1 | | | | 1 | | | |
| 480387478 | | | | | | | | | | | | | | | | | | | 1 | 1 | |
| 480391822 | | 1 | 2 | | 9 | 2 | | | | | 3 | | | 1 | 1 | 1 | | 22 | 14 | 5 | |
| 480521586 | | | | | | | | | | | 1 | | 2 | | 1 | 4 | 9 | | 1 | | |
| 480525449 | | | | | | | | 1 | | | | | | | | | | | | | |
| 480590337 | | | | | | | | | | | | | | | | | | | | 2 | |
| 480590566 | | | | | | | | | 6 | 11 | | | | | | | | | 2 | | |
| 480590606 | | | | | | | | | | | 1 | | | | | | | | | | |
| 480595244 | | | | 1 | | | | | | | | | | | | | | | | | |
| 480681197 | | | | | 1 | 1 | | | | | | | | | | | 1 | | | | |
| 480698453 | | 2 | | 6 | | | | | | | | | | | | | | | | | |
| 480737286 | | | | | | | 1 | 3 | | | | 3 | 4 | | | | | | 11 | 10 | |
| 480854134 | | | | | | | | 1 | | | | | | | 1 | | | | | | 3 |
| 480863095 | | | | | | | | 1 | | | | | | | | | | | | | |
| 480866929 | | | | | | | | | | | | | | | 1 | | | | | | |
| 480923210 | | | 2 | | | | | | | | | | | | | | | 1 | | | |
| 480945354 | | | | | | | | | | | | | | | | | | | | 1 | |
| 480978811 | | | | | | | | | | | | | | | | | | 1 | | | |
| 480987689 | | | | | | 1 | | | | | | | | | | | | | | | |
| 481034861 | | | | | 1 | | | | | | | | | | | | | | | | |
| 481053136 | | | | | | | | | | | 3 | 2 | 1 | | | | | 2 | 8 | 9 | |
| 481056857 | 2 | | | | | | | | | | 6 | 5 | 10 | | 1 | 9 | 8 | | | | |
| 481070027 | | | | 1 | 12 | 24 | | | | | | | | | | | | 2 | | | |
| 481190355 | | | | | | | | | | | | | | 1 | | | | | | | |
| 481199594 | | | | | 1 | | | | | | | | | 1 | | | | | | | |
| 481286239 | | | | | 1 | 1 | | | | | | | | | | | | | | | |
| 481406868 | | | | | 2 | | | | | | | | | | | | | 1 | 1 | | |
| 481410841 | | | | | | | | 1 | | | | | | | | | | | | | |
| 481427934 | | | | 1 | | | | | | | | 1 | | 1 | 4 | | | | | | |
| 481501144 | | | | | | | | 1 | | | 1 | 3 | 3 | | 1 | | | | 6 | 2 | |
| 481657750 | | | | | | | | | | | | | | | | | | | | 2 | |
| 481747302 | | | | | 7 | 9 | | | | | | | | | | | | | | | |
| 481770433 | | | | | | | | | | | | | | | | | | | | 2 | |
| 481838035 | | | | | | | | | | | | | 1 | 1 | | | | | | | |
| 482050067 | | | | | | | | | | | 1 | 2 | 1 | | | | | | | | |
| 482071463 | | 1 | | | | | | | | | | | | | | | | | | | |
| 482178905 | | | | | | | | | | | | | | | | | | 1 | | | |
| 482178967 | | | | | | | | | | | | | | 2 | | | | | | | |
| 482179179 | | | | | | | 1 | | | | | | | | | | | | | | |
| 482183354 | | | 2 | | | | | 1 | | | 4 | 1 | | 4 | | | | | | | |
| 482183384 | | | | | 11 | 8 | 18 | 36 | | | 3 | | 1 | | 1 | | 3 | | | | |
| 482511406 | | | | | 8 | 4 | 2 | 2 | | | 2 | | | | | | | | 1 | 1 | |
| 482520165 | | | | 1 | | | | | | | | 1 | | | | | | | | | |
| 482637370 | | | | | | 3 | | | | | | | | | | | | | | | |
| 482754144 | | | | | | | | | | | | | | | | | | 8 | | | |
| 482757982 | | | | | | | | | | | | | | | | | | | 1 | 1 | |
| 482801075 | | | | | | | | | | | | 1 | | | | | | | | | |
| 482805831 | | 2 | | | | | | | | | | | | | | | 1 | | | | |
| 482852299 | | | | | | | | | | | 6 | 3 | | | 7 | | | | | | |
| 482852401 | | | | | | | | | | | | | | | | | | | | 2 | |
| 482852469 | | | | | | | | | | | | | | 1 | | | | | | | |
| 482866008 | | | | | | | | | | | | | | | | | | | | | 3 |
| 483189268 | | | | | | | | | | | 1 | 9 | 8 | 6 | 13 | 3 | 8 | | | | |
| 483324087 | | | | | 1 | | | | | | | | | | | | | | | | |
| 483337285 | | | 2 | | | | | | | | | | | | | | 1 | | | | |
| 483646991 | | | | | 1 | | | | | | | | | | | | | | | | |
| 483720758 | | | | | | | | | | | | | | | | | | 1 | | | |
| 483750501 | | | | | | | | | | | | | | | | | 1 | | | | |
| 483880436 | | | | | | | | | | | 1 | | | 1 | | | | 1 | 1 | | |
| 484005994 | | | | | | | | | | | | | | | | | | 1 | | | |
| 484134987 | | | | | | | | | | | | | | | | | 1 | | | | |
| 484355328 | | | | | | 1 | | | | | | | | | | | | | | | |
| 484501839 | | | | | 1 | | | | | | | | | | | | | 1 | | | |
| 484562428 | | | | | 6 | 1 | | | | | | | | | | | | | | | |
| 484829654 | | | | | 2 | | | | | | | | | | | | | 20 | 4 | 6 | 22 |
| 484838760 | | | 1 | | | | 1 | | | | | | | | | | | 1 | | 1 | |
| 484911881 | | | | | | | | | | | | | | | | | | | | | 1 |
| 484911954 | | | | | | 1 | | | | | | | | | | | | | | | |
| 485171425 | | | | | 2 | | | | | | | | | | | | | 11 | 5 | 3 | 12 |
| 485252796 | | | | | | 1 | | | | | | | | | | | | | | | |
| 485511772 | | 4 | 6 | 13 | 1 | 1 | | | | | | | 1 | | | 1 | | 3 | 3 | 2 | 1 |
| 485572658 | | | 1 | | | | | | | | | | | | | | | | | | |
| 485930943 | | | | | 1 | | | | | | | | | | | | | | | | 2 |
| 485934965 | | | | | 6 | 2 | | | | | | | | | | | | 11 | 13 | 16 | 10 |
| 487100556 | | | | | 11 | 3 | | | | | | | | | | | | | | | |
| 487286529 | | | | | | 1 | | | | | | | | | | | | | | | 1 |

| | | | | | | | | | | | | | | | | | | | | | | | | | | | |
|---|---|---|---|---|---|---|---|---|---|---|---|---|---|---|---|---|---|---|---|---|---|---|---|---|---|---|---|---|
| 487541595 | | | | | | | | | | | | | | | | | | | | | | | | | 1 | | | |
| 487882168 | | | | | | | | | | | | | | | | | | | | | | | | | 1 | | | |
| 487899426 | | | | | | | | | | | | | | | | | | | | | | | | | 1 | | | |
| 488240010 | | | | | | | | | | | | | | | | | | | | | | | | | 2 | | | |
| 488572496 | | | | | | | | | | | | | 1 | | | | | | | | | | | | | | | |
| 509626405 | | | 1 | | | | | | | | | | | | | 1 | 1 | | | | | | 5 | 6 | | | | |
| 509631101 | | | | | | | | | | 1 | | | 1 | | 1 | | | 2 | 1 | | 1 | | | 1 | | | | |
| 509643775 | | | | | | | | | 1 | 1 | 2 | | | | 1 | | | | | 1 | | | 4 | 7 | | 2 | | |
| 509993739 | | | | | | | | | | 2 | | | 1 | | | | 1 | | | | 2 | | | | | | | |
| 510602264 | | | | | | | | | | | | | | | | | | | | | | | | | | | 2 | |
| 510680365 | | | | | | | | | | | | | | | | | | | | | | | | | | 1 | | |
| 511051477 | 1 | 1 | 3 | 1 | | | | | | | 1 | 18 | 25 | 40 | | 3 | 3 | 9 | 8 | 12 | 9 | 14 | 3 | 1 | 4 | 1 | 2 | |
| 511349908 | | | | | | | | | | | | | | | | | | | | | | | 1 | | | | | |
| 511358390 | | | | | | | | | | | | | | | | | | | | | | | | | 1 | | 1 | |
| 511392837 | | | | | | | | | | | | | | | | | | | | | | | 1 | | | | | |
| 511397372 | | | | | | | | | | | | | | | | | | | | | | | | 1 | | | 1 | |
| 511547921 | | | | | | | | | | | | | | | | | | | | | | 1 | | | | | | |
| 511556617 | | | | | | | | | | | | | | | | | | | | 1 | | 3 | | | | | | |
| 511561131 | | | | | | | | | | | 1 | 3 | 1 | | | | | 1 | 4 | 2 | 5 | 1 | 2 | 5 | | | | |
| 511561200 | | | | | | | | | | | | 1 | 1 | | | | | | | | | | | | | | | |
| 511617404 | | | | | | | | | | 1 | | | | | | | | | | | | | | | | | | |
| 511634742 | | | | | | | | | | | 1 | | | | | | | | | | | | | | | | | |
| 511685969 | | | | | | | | | | | | | | | | | | | | | | | | | | | 1 | |
| 511699812 | 3 | 1 | 1 | 2 | | | | | | | | | | | | | | | | | | | | | | | | |
| 511733723 | | | | | | | | | | | | | | | 1 | | | | | | | | | | | | | |
| 511772101 | | | | | | | | | | 1 | 1 | 1 | 3 | | | | | | 1 | 1 | | | | | | 2 | 2 | |
| 511862444 | | | | | | | | | | | | | | | | | | | | 1 | | | | | | | | |
| 511875377 | | | | | | | | | | | | | | 1 | | | | 1 | | | 1 | 2 | | | | | | |
| 511875414 | | | | | | | | | | | | | | | | | | 1 | | | | | | | | | | |
| 511875446 | | | | | | | | | | | | | | | | | | 1 | | | | | | | | | | |
| 511879849 | | | | | | | | | | | | | | | | | | 1 | | | | | | | | | | |
| 511966686 | | | | | | | | | | | | | | | | 3 | | | | | | | | | | | | |
| 512014262 | | | | | | | | | | 1 | | 1 | 2 | | | | | | | | | | | | | | 1 | |
| 512048416 | | | | | | | | | 1 | | 4 | | | | | | | | | | | | 7 | 1 | | | | |
| 512079523 | | | | | | | | | 1 | 4 | 4 | | | | | | | 1 | | | 1 | | 1 | | | | | |
| 512203512 | | | | | | | | | | | | | | 4 | 1 | | | | | | | | | | | | | |
| 512207845 | | | | | | | | | | | | | 1 | | | | | | | | | | | | | | | |
| 512212113 | | | | | | | | | | | | 1 | | 3 | 1 | | | | | | | | | | | | | |
| 512212142 | | | | | | | | | | | | | | | | | | | | | | 2 | | | | | | |
| 512216491 | | | | | | | | | | | | | | | | | | 1 | | | | 3 | | | | | | |
| 512225867 | | | | | | | | | | | | | | | | | | 1 | 4 | | | | | | | | | |
| 512416525 | | | | | | | | | | | | | | | | | | | | | | | | | | 2 | | |
| 512441111 | | | | | | | | | | | | | | | | | | | | | | | | | | 1 | | |
| 512535936 | | | | | | | | | | | | | | | | | | | | 1 | | 1 | | | | | | |
| 512536036 | | | | | | | | | | | | | | | | | | | | | 4 | | | | | | | |
| 512557531 | | | | | | | | | | | | | | | | | | | | | | 2 | | | | | | |
| 512774096 | | | | | | | | | | | | 1 | | | | | | | | 2 | | | | | | | | |
| 512787296 | | | | | | | | | | | | 1 | | | | | | | | | | | | | | | | |
| 513209724 | | | | | | | | | | | | | | | | | | | | | 2 | | | | | | | |
| 513209805 | | | | | | | | | | | | | | | | | | 1 | 1 | | 1 | 1 | | | | | | |
| 513218419 | | | | | | | | | 1 | | | | | | | | | | | | | | | | | | | |
| 513339129 | | | | | | | | | | | | | 1 | | | | | | | | | | | | | | | |
| 513344531 | | | | | | | | | | | | 1 | | | | | | | | | | | | | | | | |
| 513464860 | | | | | | | | | | | | | | | | | | | | | | 1 | | | | | | |
| 513546661 | | | | | | | | | | | | | | | | | | 1 | | | 1 | 1 | | | | | | |
| 513551146 | | | | | | | | | | | | | | | | | | 3 | 2 | 1 | 3 | 6 | | | | | | |
| 513555436 | 3 | | | | | | | | | | | | | | | | | 15 | 5 | 8 | 4 | 9 | 16 | 11 | | | | |
| 513668146 | | | | | | | | | | | | | 1 | | | | | | | | | | | | | | | |
| 513681077 | | | | | | | | | | | | | 1 | | | | | | | | | | | | 2 | | | |
| 513788774 | | | | | | | | | | | | | | | | | | | | | | | | | | | 1 | |
| 513840103 | | | | | | | | | | | | 1 | | | 1 | | | | | | | | | | | 1 | | |
| 513848395 | | | | | | | | | | | | | | | | | | | | | | | | | 1 | 1 | 2 | |
| 513896195 | | | | | | | | | | | | | | 1 | | | | | | | | | | | | | | |
| 514017455 | | | | | | | | | | | | 2 | 1 | | | | | | | | | | | | | | | |
| 514068564 | | | | | | | | | | | | | | | | | 1 | | | | | | | | | | 1 | |
| 514219793 | | | | | | | | | | | | | | | | | | | | | | | | | 2 | | | |
| 514224681 | | | | | | | | | | | | | 1 | 1 | | | | 1 | | | | 1 | | | | | | |
| 514272175 | | | | | | | | | | | | | | | | | | | | | | | | | 1 | 2 | | |
| 514527219 | | | | | | | | | 1 | | | | | 1 | | | | | | | 1 | | | | | | | |
| 514859045 | | | | | | | | | | | | | | 1 | 1 | | | | | 1 | 1 | | | | | | | |
| 515247907 | | | | | | | | | | | | | 1 | | | | | | | | | | | | | | | |
| 515385637 | | | | | | | | | 1 | 2 | | 3 | 5 | | 1 | | 5 | 5 | 3 | 4 | 4 | 2 | 21 | 24 | 3 | | | |
| 515580115 | | | | | | | | | | | 1 | | | | | | | | | | | | | | | | | |
| 515588291 | | | | | | | | | | | | 2 | | | | | | | | | | | | | | | | |
| 515718803 | | | | | | | | | | | | 1 | | | | | | | | | | | | | | | | |
| 515925122 | | | | | | | | | | | | 1 | | 1 | | | | 2 | | | | | 1 | | | | | |
| 515938625 | | | | | | | | | | | | | 1 | | | | | | | | | | | | | | | |
| 515947324 | | | | | | | | | | | | 2 | | | | | | | | | | | | | | | | 9 |
| 516055524 | | | | | | | | | | 1 | 1 | 1 | 3 | 3 | 1 | | | 1 | | 3 | 7 | 3 | 4 | 6 | | | | |
| 516288120 | | | | | | | | | | | | | 1 | | | | | | | | | | | | | | | |

| | | | | | | | | | | | | | | | | | | | | | | | | | | | | |
|---|---|---|---|---|---|---|---|---|---|---|---|---|---|---|---|---|---|---|---|---|---|---|---|---|---|---|---|---|
| 516288489 | | | | | | | | | | | | | | | | | | | | | | | | | | | | 6 |
| 516409549 | | | | | | | | | | | | 2 | 3 | | | | | | | | | | | | | | | |
| 516875221 | | | | | | | | | | | | | | | | | | | | | | | | | | 1 | | |
| 516943720 | | | | | | | | | 1 | 1 | | | | | | | | | | | | | | | | | | |
| 517086196 | | | | | | | | | | | | | 2 | | | | | | | | | | | | | | | |
| 517242832 | | | | | | | | | | | | 1 | | | | | | | | | | | | | | 2 | | |
| 517506265 | | | | | | | | | | | | | | | | | 1 | | | | | | | | | | | |
| 518217691 | | | | | | | | | | | | | | | | | | | | | | | | | 1 | | | |
| 518580489 | | | | | | | | | | | | | | | | | | | | | | | | | 3 | | | |
| 518899771 | | | | | | | | | | | | | | | | | | | | | | | | | 1 | | | |
| 518925962 | | | | | | | | | | | | | 1 | | | | | | | | | | | | | | | |
| 518947623 | | | | | | | | | | | | | | | | | | | | | | | | | | | | 2 |
| 520414997 | | | | | | | | | | | | | | | | | | | | | | | | | | | | 1 |
| 520613226 | | | | | | | | | | 1 | | | | | | | | | | | | | | | | | | |
| 540325242 | | | | | | | | | | | | | | | | | | | | 1 | | 1 | 5 | 3 | | 3 | 3 | |
| 540329317 | | | | | | | | | | | 1 | | | | | | | | 1 | | | | | | | | | |
| 540657374 | | | | | | | | | | | | | | | | 18 | 28 | | | | | | | 1 | | | | |
| 540662210 | | | | | | | | | | | 1 | | | | | | | | | 2 | 2 | 1 | | | | | | |
| 540674783 | | | | | | | | | | | | 32 | 26 | 85 | 41 | | | | | 1 | | 2 | 1 | | 2 | 8 | | |
| 540679089 | | | | | | | | | | | | | | | | | | | | | | | | | 1 | 2 | | |
| 540998783 | | | | | | | | | | | | | | | | 10 | 30 | | | | | | | | | | | |
| 540998882 | 1 | | 1 | | | | | | | | | | | | | 3 | 1 | 2 | | | | | | | 15 | 19 | 12 | |
| 541007502 | | | | | | | | | | | | | | | | | | 1 | 1 | | | 2 | | | | | | |
| 541019599 | | | | | | | | | | | | | | | | | | | | | 2 | | 2 | | | | | |
| 541373868 | | | | | | | | | | | 1 | | | | | 1 | | | | | | | | 1 | 1 | | | |
| 541378802 | | 1 | | 1 | | | | | | | | | 1 | | | | 1 | | | | | | | | 23 | 10 | 7 | |
| 541391609 | | | | | | | | | | | | | | 3 | | | | | | | | | | | | | | |
| 541681271 | | | | | | | | | | | | | | | | | 1 | | | | | | | | | | | |
| 541697718 | | | | | | | | | | | | | | | 2 | 1 | | | | | | | 1 | | | 1 | | |
| 541723731 | | | | | | | | | | | | | 1 | | | | | | | | | | | | | | | |
| 541913280 | | | | | | | | | | | | 1 | | | | | | | | | | | | | | | | |
| 542056766 | | | | | | | | | | | | | | 2 | | 2 | 1 | | | | | 1 | | | | | | |
| 542068823 | | | | | | | | | | | | | 1 | | | | | | | | | | | | | | | |
| 542457618 | | | | | | | | | 1 | | | | | | | | | | | | | | | | | | | |
| 542691005 | | | | | | | | | | | | | 1 | | | | | | | | | | | | | | | |
| 542798778 | | | | | | | | | | | | | | | | | | | | | | | 1 | | | | | |
| 542802879 | | | | | | | | | | | | | | | | | | | 1 | | | | | | | | | |
| 542807704 | | | | | | | | | | | | | | | | | | | | | | | 1 | | | | | |
| 542897839 | | | | | | | | | | | | | | 2 | | | | | | | | | | | | | | |
| 542902050 | | | | | | | | | | | | | | | | | | | 1 | | | | | | | | | |
| 542902307 | | | | | | | | | | | | | | | 1 | | | | | | | | | | | | | |
| 543010474 | | | | | | | | | | 3 | | 2 | | 1 | | 7 | 10 | | | | 1 | | | | 1 | | | |
| 543023799 | | | | | | | | | | | | | | | | | | | | | | | | | 1 | | | |
| 543049093 | | | | | | | | | | | | 1 | 1 | | | | | | | | | | | | | | | |
| 543062097 | | | | | | | | | | | | | | | 2 | | | | | | | | | | | | | |
| 543135607 | | | | | | | | | | | | 1 | | | | | | | | | | | | | 11 | | 1 | |
| 543234243 | | | | | | | | | | | | 1 | | | | | | | | | | | | | | | | |
| 543238468 | | | | | | | | | | | | 2 | | | 2 | | | | | | | | | | | | | 2 |
| 543247668 | | | | | | | | | | | | | 1 | | | | | | | | | | | | | | | |
| 543247684 | | | | | | | | | | | | | | | | | | | | 1 | | | | | | | | |
| 543251883 | | | | | | | | | | | | | | | | | | | 2 | | | | | | | | | |
| 543252056 | | | | | | | | | | | | | | | | | | | | | 2 | 2 | | | | | | |
| 543256347 | | | | | | | | | | | | | | | | | | | | | | | | | | | | 3 |
| 543260688 | | | | | | | | | | | | 1 | 1 | | | | | | | | | | | | | | | |
| 543365167 | | | | | | | | | | | | 2 | 2 | | | 9 | 7 | 3 | | 4 | | 2 | | | 2 | 2 | 7 | |
| 543369332 | | | | | | | | | | | | 1 | | | | | | | | | | | | | | | | |
| 543566744 | | | | | | | | | | | | 4 | | | | | | 1 | | | | | | | 6 | 1 | | |
| 543571255 | | | | | | | | | | | | | | | | | | 2 | 4 | 7 | 1 | 5 | | | | | | |
| 543580198 | | | | | | | | | | | | 1 | | | | | | | | | | | | | | | | |
| 543583770 | | | | | | | | | | | | | | | | | | | 1 | | | | | | | | | |
| 543584523 | | | | | | | | | | | | | | | | | | | 1 | | | 1 | | | 1 | | 1 | |
| 543692985 | | | | | | | | | | | | | | 5 | 1 | | | | | | | | | | | 1 | | |
| 543702186 | | | | | | | | | | | | | | 3 | 4 | | | | | | | | | | | | | |
| 543718301 | | | | | | | | | | | | 1 | | | | | | | | | | | | | | | | |
| 543804534 | | | | | | | | | | | | | | | | | | | | | | | | | | 2 | | |
| 543808842 | | | | | | | | | | | | | | | | | | | | | | | | | | | | 1 |
| 543821838 | | | | | | | | | | | | | | | | | | | | | | | | | | | 1 | |
| 543916401 | | | | | | | | | | | | | | | | | | 1 | | | 1 | | | | | | | |
| 543925435 | | | | | | | | | | | | | 1 | | | | | | | | | | | | | | | |
| 543929121 | | | | | | | | | | | | | | | | | | | 1 | | | | | | | | | |
| 543934429 | | | | | | | | | | | | | | | | | | | | | | | | | | | | 1 |
| 544038013 | | | | | | | | | | | | 2 | | | | | | | | | | | | | | | | |
| 544158940 | | | | | | | | | | | | | | | | | | | | | | | | | | | 1 | |
| 544184932 | | | | | | | | | | | 1 | 1 | | | | | | | | | | | | 4 | | | | |
| 544249161 | | | | | | | | | | | | 1 | | | | | | | | | | | | | | | | |
| 544262349 | | | | | | | | | | | | | | | | | | | | | | | | | | | | 1 |
| 544265926 | | | | | | | | | | | | | | | | | | | | | | | | | | | | 2 |
| 544379269 | | | | | | | | | | | | 2 | 1 | | | | | | | | | | | | | | | |
| 544599014 | | | | | | | | | | | | | | | | | | | | | | | | | | | | 1 |
| 544606883 | | | | | | | | | | | | | | | | | | | | | | | | | | | | 1 |

| | | | | | | | | | | | | | | | | | | | | | | | |
|---|---|---|---|---|---|---|---|---|---|---|---|---|---|---|---|---|---|---|---|---|---|---|---|
| 544606915 | | | | | | | | | | | | | | | | | 3 | | | | | | |
| 544620289 | | | | | | | | | 1 | | | | | | | | | | | | | | |
| 544699051 | | | | | | | | | | | | | | | | | | | | 1 | | | |
| 544939871 | | | | | | | 1 | 4 | | | | | | | | | | | | | | | |
| 544943733 | | | | 1 | | | | | | | | | | | | | | | | | | | |
| 545224254 | | | | | | | | | | | | | 1 | 2 | 2 | 1 | 1 | | | | | | |
| 545284807 | | | | | 1 | | | | | | | | | | | | | | | | | | |
| 545510060 | | | | | | | | | | | | | | | | | | | 1 | | | | |
| 545535468 | | | | | | | 4 | 4 | | | | | | | | | | | | | | | |
| 545625824 | | | | | | | | 3 | | | | | | | | | | | | | | | |
| 545748103 | | | | | | | 2 | 2 | | | | 1 | 1 | | 1 | | 1 | | | | | | |
| 545906392 | | | | 1 | 1 | | | | | | | | | | | | | | | | | | |
| 545907195 | | | | | | | | 1 | | 1 | | | 1 | | | 1 | | | | | 7 | 4 | |
| 545966796 | | | | | | | 3 | | | | | | | | | | | | | | | | |
| 545971090 | | | | | | | | 2 | | | | | | | | | | | | | | | |
| 546217818 | | | | | | | | 1 | | | | | | | | | | | | 15 | 2 | 3 | |
| 546312597 | | | | | | | | | | | | | | | | | | | | | | | 2 |
| 546917498 | | | | | | 2 | | | | | | | | | | | | | | | | | |
| 546925626 | | | | 3 | 1 | 5 | | | | | | | | | | | | | 10 | | | | |
| 546981769 | | | | | | | | | | | | | | | | | | | | | | | 1 |
| 547439565 | | | | | | | 1 | 1 | | | | | | | | | | | | | | | |
| 547612423 | | | | | 1 | | | | | | | | | | | | | | | | | | |
| 547918570 | | | | | | | | | | | | | 2 | 2 | 2 | | 1 | | | | 2 | 4 | |
| 547927681 | | | | | 1 | | | | | | | | | | | | | | | | | | |
| 547978595 | | | | 1 | 3 | 1 | | | | | | | | | | | | | 1 | | | | |
| 548367533 | | | | | | | | | | | | | | | | | | | | | | | 1 |
| 548367551 | | | | | | | | | | | | | | | | | | | | | | | 1 |
| 548377264 | | | | | | | | | | | | | | | | | | | | | | | 1 |
| 548704181 | | | | | | | | | | | | | | | | | | | | | | | 1 |
| 548712377 | | | | | | | | | | | | | | | | | | | | 2 | | | 1 |
| 548955060 | | | | | | | 2 | 7 | | | | | | | | | | | | | | | |
| 549282467 | | | | | | | 1 | | | | | | | | | | | | | | | | |
| 549377313 | | | | | | | | | | | | | | | | | | | | | 1 | | |
| 549381505 | | | | | | | | | | | | | | | | | | | | | | | 1 |
| 549597974 | | | | | | | | | | | | | | | | | | | 1 | | | | |
| 549611256 | | | | | | | 2 | | | | | | | | | | | | | | | | |
| 549714149 | | | | | | | | | | | | | | | | | | | | | | | 3 |
| 550033958 | | | | | | 1 | | | | | | | | | | | | | | | | | |
| 550063586 | | | | | | | | | | | | | | | | | | | | | | | 2 |
| 550297695 | | | | | | | | | 1 | | | | | | | | | | | | | 2 | |
| 550326974 | | | | | | | | | | | | | | | | | | | | 1 | | 1 | |
| 550408962 | | | | | | | | | | | | | | | | | | | | | | | 1 |
| 550617370 | | | | | | | | | | | | | | | | | | | | 1 | | | |
| 550758596 | | | | | | | | | | | | | | | | | | | | | | | 1 |
| 550953198 | | | | 3 | | | | | | | | | | | | | | | | | | | |
| 551086679 | | | | | | | | | | | | | | | | | | | | | | | 1 |
| 551095302 | | | | | | | | | | | | | | | | | | | | | | | 1 |
| 571346887 | | | | | | | | 1 | | | | | | | | | | | | | | | |
| 571355851 | | | | | | | | | 2 | | | | | | | | | | | | | | |
| 571364144 | | | | | | 7 | | | | | | | | | | | | | | | | | |
| 571364778 | | 3 | | | | | | | 1 | | 1 | | | | | | | 2 | 1 | | | | |
| 571368849 | | | | | 2 | | 4 | 1 | 1 | 2 | | | 6 | 5 | 7 | 4 | 10 | 1 | | 1 | | 1 | |
| 571372889 | | | | | | | 3 | 2 | | | | 1 | | | | | | | | 19 | 20 | 11 | |
| 571692115 | | | | | 2 | | | | | | | | | | | | | | 1 | | | | |
| 571692839 | | | 1 | | | | | | | | | | | | | | | | | | | | |
| 571709547 | | | | | 2 | 1 | | | | | | | | | | | | 1 | 3 | | | | |
| 571713905 | | | | | 1 | | | | | | | | | | | | 1 | 3 | 5 | 1 | 1 | | |
| 571717923 | | | | | | | | | | | | | | | | | | | 2 | | | | |
| 571726583 | 1 | | | | | | | | | | | | | | | | | | | | | | |
| 572046627 | | | | | | | | | | | | 1 | | | | | | | | | | | |
| 572050535 | | | | | | | | | | | | 1 | | | | | | | | 1 | 2 | | |
| 572408934 | | | | | | | | 1 | | | | | | | | | | 2 | | 1 | 20 | 27 | |
| 572754751 | | | | | | | | | | | | | | | | | | | | | | 2 | |
| 573090850 | | | | | | | | | | | 2 | 3 | | | | | | | | 1 | | | |
| 573142557 | | 1 | | | | | | | | | | | | 1 | | | | | | | | | |
| 573143028 | | | | | | | | | | | | | | | | | | | 1 | | | | |
| 573289509 | | | | | | | | | | 1 | | | | | | | | | | 2 | 1 | | |
| 573381453 | | | | | | | 1 | | | | | | | | | | | | | | | | |
| 573415212 | | | | | | | 1 | | | | | | | | | | | | | | | | |
| 573774122 | | | | | | | | | 5 | | 7 | 6 | | | | | | | | | | | |
| 574062996 | | | | | | | 3 | 3 | 6 | 2 | | | 1 | 1 | 1 | 2 | | | 2 | 3 | | 1 | |
| 574260596 | | | | | | | | | | | | | | | | | 1 | | | | | | |
| 574395611 | | | | | | | | 1 | | | | | | | | | | 2 | | 2 | | 2 | |
| 574511458 | | | | | | | 19 | 24 | | | | | | | | | | | | 2 | | | |
| 574688051 | | | | | | | | | | | | 2 | | | | | | | | | | | |
| 574740549 | | | | | | | | | | 1 | | | | | | | | | | | | | |
| 574938221 | | | | | | | | | | | | | | | | 1 | | | | | | | |
| 574973454 | | | | | | | | 1 | | | | | | | | | | | | | | | |
| 575063620 | | | | | | | | | | | | | | | | | | | | 1 | | | |
| 575073147 | | | | | | | 1 | | | | | | | | | | | | | | | | |

| | | | | | | | | | | | | | | | | | | | | | | | | | | | | |
|---|---|---|---|---|---|---|---|---|---|---|---|---|---|---|---|---|---|---|---|---|---|---|---|---|---|---|---|---|
| 575206067 | | | | | | | | | | | 1 | | | | | | | | | | | | | | | | | |
| 575301459 | | | | | | | | | | | | 1 | 5 | | | | | | | | | | | | | | | |
| 575323308 | | | | | | | | | | | | | 1 | | | | | | | | | | | | | | | |
| 575327207 | | | | | | | | | | | | | | | | | | | | | | | | | | 3 | 2 | |
| 575651309 | | | | | | | | | | | | 1 | 4 | | | | | | | | | | | | | | | |
| 576082257 | | | | | | | | | | | | | | | | | | | | | | | | | 1 | | | |
| 576260070 | | | | | | | | | 1 | 1 | 5 | | | | | | | 3 | | | | | | | | | | |
| 576574889 | | | | | | | | | | | | | | | | | | | | | | | | | | | 1 | |
| 576597029 | | | | | | | | | | | | | | | | | | | | | | | | 1 | | | | |
| 577244511 | | | | | | | | | | | | | 2 | | | | | | | | | | | | 1 | | | |
| 577304838 | | | | | | | | | | | | | | | | | | | | | 1 | | | | | | | |
| 577330676 | | | | | | | | | | | | | | | | | | | | | | | | | | 1 | | |
| 577554935 | | | | | | | | | | | | | 1 | | | | | | | | | | | | | | | |
| 577645883 | | | | | | | | | | | | | | | | | | | 1 | | | | | | | | | |
| 578284233 | | | | | | | | | | | | | | | | | | | | | | | | | 6 | 1 | | |
| 578284872 | | | | | | | | | | | | | | | | | | | | | | | | | 1 | | | |
| 578686468 | | | | | | | | | | | | | 1 | | | | | | | | | | | | | | | |
| 578811223 | | | | | | | | | | | | 1 | | | | | | | | | | | | | | | | |
| 578919867 | | | | | | | | | | | | | | 1 | | | | | | | | | | | | | | |
| 579014562 | | | | | | | | | | | 2 | | | | | | | | | | | | | | | | | |
| 579666216 | | | | | | | | | 2 | | 1 | | | | | | | | | | | | | | | | | |
| 579695775 | | | | | | | | | | 1 | | | | | | | | | | | | | | | | | | |
| 580010810 | | | | | | | | | | | 2 | | | | | | | | | | | | | | | | | |
| 580714587 | | | | | | | | | | | | | | | | | | | | | | | | | 1 | | | |
| 580745387 | | | | 1 | | | | | | 1 | 1 | 3 | | | | | | | | | | | | | | | | |
| 581073524 | | | | | | | | | | | | | | | | | | | | | | | | | 1 | | | 3 |
| 601712799 | | | | 3 | | | | | | | | | | | 1 | 1 | | | 1 | | | | | | 1 | | | |
| 601712813 | | | 3 | | | | | | | | | 2 | | 1 | | | | | | | 1 | | 3 | | | | | |
| 601716965 | | | 1 | | | | | | | | | | 1 | 1 | 1 | | 1 | | | | | | 1 | | | | | |
| 602044790 | | | | | | | | | | | | | | | | 1 | | | | | | | | | | | | |
| 602071049 | 1 | | | | | | | | | | | | | | | | | | | | | | | | | | | |
| 602382052 | | | | | | | | | | | | | | 1 | 1 | | | | | | 1 | | | | | | | |
| 602399280 | | | | | | | | | | | | | | | | | | | 4 | 1 | | | | | | | | |
| 602675369 | | | | | | | | | | 1 | | | | | | | 3 | | | | | | 4 | 2 | | | | |
| 603089654 | | | | | | | | | | | | | | | | | | | | 2 | | | | | | | | |
| 603095061 | | | | | | | | | | | | | | | | | | | | | | | | 1 | | | | |
| 603465268 | | | | | | | | | | | | | | | | | | 2 | | | | | | | | | | |
| 603466235 | | | | | | | | | 1 | | 1 | | | | | | | | | | | | 1 | | 1 | | | |
| 603482385 | 1 | | | | | | | | | | | | | | | | | | | | | | | | 1 | 6 | 5 | |
| 603755188 | | | | | | | | | | | | | | | | | | | | | | | | | 1 | | | |
| 603785283 | | | | | | | | | | | | 2 | 5 | 5 | 1 | | | | | | | | 1 | 4 | | 2 | 3 | |
| 603810512 | | | | | | | | | | 2 | | | | | | | | | | | | | | | | | | |
| 603810673 | | | | | | | | | | | 1 | | | | | | | | | | | | | | | | | |
| 604161217 | | | | | | | | | | 1 | | | | | | | | | | | | | | | | | | |
| 604194925 | | 1 | | | | | | | | | | | | | | | | | | | | | | | | 3 | | |
| 604460198 | | | | | | | | | | | | | | | | | 1 | | | | | | | | | | | |
| 604476368 | | | | | | | | | | | | | | | | | | | | | | | | | 1 | | | |
| 604510234 | | | | | | | | | | | | | 1 | | | | | | | | | | | | | | | |
| 604825724 | | | | | | | | | | | | | | | | | | | | | | | 1 | | | | | |
| 604847018 | | | | | | | | | | | | | | | | | | | | | | | | | 1 | | | |
| 604864392 | | | | | | | | | | | | | | | | | | | | | | | | | 1 | | | 1 |
| 604889732 | | | | | | | | | | 1 | | | | | | | | | | | | | 1 | | | | | |
| 604976663 | | | | | | | | | | | | | | | | | | | | | | | | | | | | 1 |
| 604989814 | | | | | | | | | | | | | | | | | | | | | | | | | | | | 1 |
| 605554875 | | | | | | | | | | | | | 4 | | | | | | | | | | | | | | | |
| 605775210 | | | | | | | | | | | | | | 18 | 8 | | | | | | | | | | | | | |
| 605999323 | | | | | | | | | | | | 1 | | | | | | | | | | | | | | | | |
| 606944503 | | | | | | | | | | | | | | | | | | | 1 | | | | | | | | | |
| 607264357 | | | | | | | | | | | | | | | | | | | | | | | | | 2 | | | |
| 607273128 | | | | | | | | | | | | | 1 | | | | | | | | | | | | | 2 | | |
| 607273339 | | | | | | | | | | | | | | | | | | | | | | | | | | 1 | | |
| 607294784 | | | | | | | | | | | | | | | | | | 1 | | | 1 | | | | | | | |
| 607597135 | | | | | | | | | | | | 1 | | | | | | | | | | | | | | | | |
| 607929192 | | | | | | | | | | | | 23 | 18 | | | | | | | | | 1 | | | | | 1 | |
| 607955261 | | | | | | | | | | | | 1 | | | | | | | | | | | | | | | | |
| 607998424 | | | | | | | | | | | | | | | | | | | | | 2 | 1 | | | | | | |
| 608377992 | | | | | | | | | | | | | | | | | | | | | | | | | | | | 1 |
| 608715651 | | | | | | | | | | | | | | | | | | | | | | | | | | | | 11 |
| 609384016 | | | | | | | | | | | | | | | | | | | | | | | | | 1 | | | |
| 609384093 | | | | | | | | | | | | | 1 | | | | | | | | | | | | | | | 1 |
| 609587511 | | | | | | | | | | | | | | | | | | | | | | | | | 1 | | | |
| 609655670 | | | | | | | | | | | | 1 | | | | | | | | | | | | | | | | |
| 609656444 | | | | | | | | | | | | | 1 | | | | | | | | | | | | | | | |
| 609660480 | | | | | | | | | | | | 1 | | | | | | | | | | | | | 2 | | | |
| 609997539 | | | | | | | | | | | | | | | | | | | | | | | | | | | | 1 |
| 610074665 | | | | | | | | | | | | | | | | | | | | | | | | | | | | 1 |
| 610558436 | | | | | | | | | 2 | | | 2 | 6 | | | | | | | | | | | | | | | |
| 610666582 | | | | | | | | | | | | | | | | | | | | | | | | | 1 | | | |
| 610670860 | | | | | | | | | | | | | 1 | | | | | | | | | | | | | | | |
| 610704870 | | | | | | | | | 2 | | 1 | | | | | | | | | | | | | | | | | |

| | | | | | | | | | | | | | | | | | | | | | | | | | | | | |
|---|---|---|---|---|---|---|---|---|---|---|---|---|---|---|---|---|---|---|---|---|---|---|---|---|---|---|---|---|
| 611016455 | | | | | | | | | | | | | | | | | | | | | | | | | | | | 1 |
| 611067721 | | | | | | | | | 3 | | | | | | | | | | | | | | | | | | | |
| 611089612 | | | | | | | | | | | | | | | | | | | | | | | | | | | | 16 |
| 611340150 | | | | | | | | | | | | | | | | | | | | | | | | | | | 1 | |
| 611357256 | | | | | | | | | | | | | | | | | | | | | | | | | 1 | | | 1 |
| 611706830 | | | | | | | | | | | | | | | | | | | | | | | | | 1 | | | |
| 611974080 | | | | | | | | | | | | | | | | | 1 | | | | | | | | | | | |
| 633097767 | | | | | | | | | | 1 | | 1 | | | | | 1 | | | | | | | | | | | |
| 633097828 | | | | | | | | | | | | | | | | 10 | 13 | | | | | | | | 1 | | | |
| 633101926 | | | | | | | | | | | | | | | | | | | | | | | | | 1 | | | |
| 633412404 | | | | | | | | | | | | | | | | | | | | | | | 1 | | | | | |
| 633434319 | | | 1 | | | | | | | | | | | | | | | | | | | | | | | | | |
| 633438933 | | | | | | | | | | | | | | | | 30 | 24 | | | | | | | | | 1 | | |
| 633447561 | | | | | | | | | | | 1 | | | | 1 | | | | | | | | | | | 2 | | |
| 633793553 | | | | | | | | | | 2 | | 1 | | | | | 1 | | | | | | | | | 2 | 1 | |
| 634090169 | | | | | | | | | | | | | | | | | | | | | | | | | 1 | | | |
| 634138422 | | | | | | | | | | | | | 1 | | | | | | | | | | | | | | | |
| 634759240 | | | | | | | | | | | | | | | | | 2 | | | | | | | | | | | |
| 634879889 | | | | | | | | | | | | | | | | | | | | | | | | 1 | | | | |
| 635109166 | | | | | | | | | | | | | | | | | 1 | | | | | | | | | | | |
| 635424442 | | | | | | | | | | 1 | | | | | | | | | | | | | | | | | | |
| 635523991 | | | | | | | | | | | | | | | | | | | | | | | | 2 | 1 | 12 | 14 | |
| 635541640 | | | | | | | | | | | | | 2 | | 3 | | | | | | | | | | 1 | | | |
| 635890732 | | | | | | | | | | | | | | | | | | | | | | | 1 | | | 1 | | |
| 635895000 | | | | | | | | | | | | 1 | 1 | | | | | | | | | | | | 8 | 16 | 17 | |
| 636344136 | | | | | | | | | | | | | | 1 | | | | | | | | | | | | | | |
| 636701737 | | | | | | | | | | | | | | | | | | | | | | | | | | | | 1 |
| 636944318 | | | | | | | | | | | | 1 | | | | | | | | | | | | | | | | |
| 637008792 | | | | | | | | | | | | 8 | 6 | | | | | | | | | | | | | | 1 | |
| 637306291 | 1 | | | | | | | | | 2 | 1 | | 4 | | 1 | | | 9 | 3 | 6 | 7 | 5 | 5 | 4 | | 4 | | |
| 637673302 | | | | | | | | | | | | | | | | | | 3 | | 1 | 4 | 1 | | | | | | |
| 637914938 | | 3 | | | | | | | | | | | | | | 4 | 1 | | | | | | | | | | | |
| 638040562 | | | | | | | | | | | | | | | | | | | | | | | | | | 1 | | |
| 638355885 | | | | | | | | | | | | | | | | | | | | 1 | | | | | | | | |
| 638481581 | | | | | | | | | | | | | | | | | 1 | | | | | | | | | | | |
| 638985608 | | | | | | | | | | | | | | | | | | | | | | | | | 1 | | | |
| 639327084 | | | | | | | | | | | | | | | | | | | | | | | | | 6 | | | |
| 639650941 | | | | | | | | | | | | 1 | | | | | | | | | | | | | | | | |
| 639694375 | | | | | | | | | | | | | | | | | | 1 | | | | | | 1 | | | | |
| 640082371 | | | | | | | | | | | | | | | | | | | | | | | | | | | | 4 |
| 640337151 | | | | | | | | | | | | | | | | | | | | | | | | | 1 | | | |
| 640539567 | | | | | | | | | | | | 1 | | | | | | | | | | | | | | | | |
| 640755994 | | | | | | | | | | | | 1 | | | | | | | | | | | | | | | | |
| 640760406 | | | | | | | | | | | | | 1 | | | | | | | | | | | | | | | 2 |
| 640877406 | | | | | | | | | | | | | | | | | | | | 1 | | | | | | | | |
| 641481132 | | | | | | | | | | | | | | | | | | | | | | | | | | | | 1 |
| 642046803 | | | | | | | | | | | | | | | | | | | | | | | | | 1 | | | |
| 642422617 | | | | | | | | | 1 | | | | | | | | | | | | | | | | | | | |
| 642651049 | | | | | | | | | | | | | | | | | | | | | | | | | 2 | | | |
| 662197764 | | | | | | | | | | | | | 1 | | 1 | | | | | | | | | | | | | |
| 663432544 | | | | | | | | | | | | | | | | 161 | 139 | | | | | | | | | | | |
| 664447502 | | | | | | | | | | | | | | | | | | | | | | 1 | | | | 1 | | |
| 664456036 | | | | | | | | | | | | | | | | | | | | | | | | | 1 | 4 | | |
| 664473327 | | | | | | | | | | | | | | | | | 1 | | | | | | | | | | | |
| 664824047 | | | | | | | | | | | | | | | | 1 | 1 | | | | | | | | | | | |
| 665164021 | | | | | | | | | | | | | | | | | | | | | | | | | 1 | | | |
| 665181585 | | | | | | | | | | | | | | | | | | | | | | | | 1 | | | | |
| 665190290 | | | | | | | | | | | | | | | | | | | | | | | | 1 | | | | |
| 665191303 | | | | | | | | | | | | | | | | | | | | | | | | 1 | | | | |
| 665194717 | | | | | | | | | | | | | | | | | | | | | | | | 1 | | | | |
| 665552356 | | | | | | | | | | 2 | 2 | | | | | | | | | | | | | | | | | |
| 666212865 | | | | | | | | | | | | | | | | | | | | | | | | | | 1 | | |
| 666230992 | | | | | | | | | | | | | | | | | | | | | | | | | 1 | 1 | | |
| 666252297 | | | | | | | | | | | | | 1 | 3 | 5 | 7 | 3 | | | | | | | | | | | |
| 666503184 | | | | | | | | | | | | | | | | | | | | | | | | | | | 1 | |
| 666567261 | | | | | | | | | | | | | | | | | | | | | | | | | | | 1 | |
| 666904390 | | | | | | | | | | | | | | | | | | | | | | | | | | 2 | | |
| 666916874 | | | | | | | | | | | | | | | | | | | | | | | | | 1 | | | |
| 668340913 | | | | | | | | | 4 | 26 | 14 | 4 | 7 | 1 | 1 | 4 | 1 | | | 1 | | | | | | | | |
| 668397992 | | | | | | | | | | | | | | | | | | | | | 1 | | | | | | | |
| 669171261 | | | | | | | | | | 2 | | | | | | 4 | | | | | | | | | | | | |
| 669675454 | | | | | | | | | | | | | | | | | | | | | | | | | 6 | | | |
| 669726929 | | | | | | | | | | | | | | | | | | | 1 | | | | | | | | | |
| 669744300 | | | | | | | | | | | | 1 | | | | | | | | | | | | | | | | |
| 670020922 | | | | | | | | | | | | | 1 | | | | | | | | | | | | | | | |
| 670094261 | | | | | | | | | | | | | | | | | | | | | | | | | | | | 7 |
| 670348810 | | | | | | | | | | | | | | | | | | | | | | | | | 1 | | | |
| 670370149 | | | | | | | | | | | | | 1 | | | | | | | | | | | | 2 | | | |
| 670370239 | | | | | | | | | | | | | | | | | | | | | | | | | | | 1 | |
| 670370808 | | | | | | | | | | | | | | | | | | | | | | | | | 7 | | | |

| | | | | | | | | | | | | | | | | | | | | | | | | | | | | |
|---|---|---|---|---|---|---|---|---|---|---|---|---|---|---|---|---|---|---|---|---|---|---|---|---|---|---|---|---|
| 670431030 | | | | | | | | | | | | 1 | | | | | | | | | | | | | | | | 1 |
| 670772147 | | | | | | | | | | | | 1 | | | | | | | | | | | | | | | | 1 |
| 670776253 | | | | | | | | | | | | | | | | | | | | | | | | | | | | 6 |
| 670776480 | | | | | | | | | | | | | | | | | | | | | | | | | | | | 7 |
| 671234167 | | | | | | | | | | | | 1 | | | | | | | | | | | | | 1 | | | |
| 671574812 | | | | | | | | | | | | 1 | | | | | | | | | | | | | | | | |
| 671751354 | | | | | | | | | | | | | | | | | | | | | | | | | | | | 1 |
| 672041363 | | | | | | | | | | | | | | | | | | | | | | | | | 1 | | | |
| 673639143 | | | | | | | | | | | | | 4 | | | | | | | | | | | | | | | |
| 673867858 | | | | | | | | | | | | 1 | | | | | | | | | | | | | | | | |
| 674108632 | | | | | | | | | | | | 12 | 15 | | | | | | | | | | | | | | | |
| 675166212 | | | | | | | | | | | | | 1 | | | | | | | | | | | | | | | |
| 694471913 | | | | | | | | | | | | | | | | 5 | 2 | | | | | | | | | | | |
| 694472338 | | | | | | | | | | | | | | | | 33 | 20 | | | | | | | | | | | |
| 694472667 | | | | | | | | | | | | | | | | 1 | 1 | | | | | | | | | | | |
| 694813676 | | | | | | | | | | | | | | | | 23 | 33 | | | | | | | | 1 | | | |
| 694834412 | | | | | | | | | | | | | | | | | | | | | | | | | | | 1 | |
| 695124651 | | | | | | | 1 | | | | | 13 | 15 | 6 | 4 | | 1 | | | | 2 | 1 | | 1 | | | 1 | |
| 695128320 | | | | | | | | | | | | | 2 | | | | 1 | | | | | | | | | | | |
| 695132361 | | | | | | | | | | | | 2 | 3 | 8 | 5 | | | | | | 1 | | 2 | | | | | |
| 695163759 | | | | | | | | | | | | | | | | | | | | | | | | | | | 1 | |
| 695167675 | | | | | | | | | | | | | | | | 1 | | | | | | | | | | | | |
| 695508439 | | | | | | | | | | | | | | | | | 1 | | | | | | | | | | | |
| 695875649 | | | | | | | | | | | | | | | | | | | | | | | | | 1 | | | |
| 696489026 | | | | | | | | | | | | | | | | | | | | 1 | | | | | 1 | | | |
| 696829800 | | | | | | | | | | | | | | | | | | | | | | 1 | | | 2 | 4 | 1 | |
| 696830498 | | | | | | | | | 1 | | 1 | 8 | 10 | | | | | | 2 | | 6 | | 5 | 2 | | | | |
| 696846594 | | | | | | | | | | | | | | | | | | | | | | | 1 | | | | | |
| 696850811 | | | | | | | | | | | | | 1 | | | 1 | | | | | | | 1 | | | | | |
| 696863848 | | | | | | | | | | | | 2 | | 1 | | | | | | | | | | | | | | |
| 696881909 | | | | | | | | | | | | | | 5 | 2 | | | | | | | | | | | | | |
| 697252464 | | | | | | | | | | 1 | 2 | | 1 | | | | | 1 | | 1 | 4 | 3 | 1 | | | | 1 | |
| 697265869 | | | | | | | | | | | | | | 1 | | | | | | | | | | | | | | |
| 697537598 | | | | | | | | | | 1 | | | | | | | | | | | | | | | | | | |
| 697645622 | | | | 1 | | | | | | | | | | | 1 | | | | | | | | | | | | 2 | |
| 697891313 | | | | | | | | | | | | | 1 | | | | | | | | | | | | | | | |
| 697891769 | | | | | | | | | | | | | | | | | | | | | | | | | 1 | | | |
| 698418398 | | | | | | | | | | | | | | | | | | | | | | | | | | | | 1 |
| 698418640 | | | | | | | | | | | | 2 | | | | | | | | | | | | | | | | |
| 698508565 | | | | | | | | | | | | | | | | | 2 | | | | | | | | | | | |
| 698970725 | | | | | | | | | | | | | | | | 28 | 11 | | | | | | | | 1 | | | |
| 698979607 | | | | | | | | | | | | 2 | 4 | | | | | | | | | | | | | | | |
| 699031185 | | | | 1 | 4 | 3 | 1 | 3 | | | | | | | | | | | | | | | 1 | | | | | |
| 699035090 | | | | | | | | | | | | | | | | | | | | | | | 1 | | | | | |
| 699035753 | | | | | | | | | | | | | | | | | | | | | | | 1 | | | | 2 | |
| 699363923 | | | | | | | | | | | | | 1 | | | | | | | | | | | | | | | |
| 699652692 | | | | | | | | | | | | | | | | 1 | | | | | | | | | | | | |
| 699687305 | | | | | | | | | | | | 16 | 19 | | | | | | | | | | | | | | | |
| 700002191 | | | | | | | | | | | | | | | | | | | | | | | | 1 | | | | |
| 700878295 | | | | | | | | | | | | | | | | 1 | | | | | | | | | | | | |
| 701077130 | | | | | | | | | | | | | | | 1 | | | | | | | | | | | | | |
| 701469455 | | | | | | | | | | | | | | | | | | | | | | | | | | | | 1 |
| 701815778 | | | | | | | | | | | | | 1 | | | | | | | | | | | | | | | |
| 702152113 | | | | | | | | | | | | 11 | 10 | | | | | | | | | | | | 1 | | | |
| 702497335 | | | | | | | | | | | | | 2 | | | | | | | | | | | | | | | |
| 703145276 | | | | | | | | | | 1 | | | | | | | | | | | | | | | | | | |
| 703296640 | | | | | | | | | | | | | 2 | | | | | | | | | | | | | | | |
| 703623655 | | | | | | | | | | | | | | | | | | | | | | | | 1 | | | | |
| 703857647 | | | | | | | | | | | | | | | | | | | | | | | | | | | | 1 |
| 703866010 | | | | | | | | | | | | | 2 | | | | | | | | | | | | | | | |
| 724820565 | | | | | | | | | | | | | | | | 1 | | | | | | | | | | | | |
| 725498630 | | | | | | | | | | | | | | | | 4 | 3 | | | | | | | | | | | |
| 725839388 | | | | | | | | | | | | | | | | 1 | 6 | | | | | | | | 3 | | 1 | |
| 726207450 | | | | | | | | | | | | | | | | 20 | 30 | | | | | | | | | | | |
| 726858507 | | | | | | | | | | | | | | | | 2 | | | | | | | | | | | | |
| 726918599 | | | | | | | | | | | | | | | | | | | | | | | | | 2 | | | |
| 726918990 | | | | | | | | | | | | | | | | | 1 | | | | | | | | | | | |
| 727182822 | | | | | | | | | | 1 | | | | | | | | | | | | | | | | | | |
| 727225324 | | | | | | | | | | | | | | | | | | | | | | | | | 2 | | | |
| 727260771 | | | | | | | | | | 1 | 1 | | | | | | | | 1 | | | | | | | | | |
| 727280915 | | | | | | | | | 2 | | | | | | | | | | | | | | | 1 | | | | |
| 727285326 | | | | | | | | | | | | | | | | | | 1 | | | | | | | 1 | | 1 | |
| 728230740 | | | | | | | | | | | | 4 | | | | | | | | | | | | | | | | |
| 728633073 | | | | | | | | | | | | | | | | 7 | 2 | | | | | | | | | | 1 | |
| 728926409 | | | | | | | | | | | | | | | | | | | | | | | | | | | 1 | |
| 729698667 | | | | | | | | | | | | | 1 | | | | | | | | | | | | | | | |
| 729798521 | | | | | | | | | | | | | | | | | | | | | | | | | | | | 1 |
| 729957687 | | | | | | | | | | | | | | | 1 | | | | | | | | | | | | | |
| 730118077 | | | | | | | | | | | | | | | | | | | | | | | | | | | | 2 |
| 730131007 | | | | | | | | | | | | 2 | | | | | | | | | | | | | | | | |

| | | | | | | | | | | | | | | | | | | | | | | | |
|---|---|---|---|---|---|---|---|---|---|---|---|---|---|---|---|---|---|---|---|---|---|---|---|
| 731062953 | | | | | | | | | | | | | | | | | | | | | 1 | | |
| 731145160 | | | | | | | | 1 | 1 | | | | | | | | | | | | | | |
| 731494823 | | | | | | | | 2 | | | | | | | | | | | | | | | |
| 732034061 | | | | | | | | | | | | 1 | | | | | | | | | | | |
| 732478552 | | | | | | | | | | | | | | | | | | 1 | | | | | |
| 733292029 | | | | | | | | 4 | 3 | | | | | | | | | | | | | | |
| 733458828 | | | | | | | | | | | | | | | | | | | 1 | | | | |
| 734491001 | | | | | | | | 1 | | | | | | | | | | | | | | | |
| 734677385 | | | | | | | | 1 | 1 | | | | | | | | | | | | | | |
| 735008444 | | | | | | | | | 1 | | | | | | | | | | | | | | |
| 735699469 | | | | | | | | | 1 | | | | | | | | | | | | | | |
| 735798583 | | | | | | | | | | | | | | | | | | | | | 1 | | |
| 736860990 | | | | | | | | 1 | | | | | | | | | | | | | | | |
| 737491062 | | | | | | | | | | | | 1 | 1 | | | | | | | | | | |
| 739917109 | | | | | | | | | | | | | | | | | | | | 1 | | | |
| 756502912 | 1 | | | | 1 | | | | | 1 | | | | | | | | | | | | | |
| 756870190 | | | | | | | | | | | | 4 | | | | | | | | | | | |
| 756874510 | | | | | | | | | | | | 1 | 3 | | | | | | | | | | |
| 757215581 | | | | | | | | | | | | 4 | 2 | | | | | | | | | | |
| 757374944 | | | | | | | | | | | | | 1 | | | | | | | | | | |
| 757530901 | | | | | | | | 2 | | 2 | | | | | | | | | | | | | |
| 757556799 | | | | | | | | | | | | 1 | 1 | | | | | | | | | | |
| 758320421 | | | | | | | | | | | | | | | | | | | | | 1 | | |
| 758903321 | | | | | | | | | | | | | 2 | | | | | | | | | | |
| 758955666 | | | | | | | | | 1 | | | | | | | | | | | | | | |
| 758989937 | | | | | | | | | | | | | | | | | | | | | | | 1 |
| 759582005 | | | | | | | | | | | | 4 | 2 | | | | | | | | | | |
| 759632919 | | | | | | | | | | | 3 | 38 | 25 | | | | | | | | | | |
| 759810119 | 1 | | | | | | | | | | | 1 | 2 | | | | | | | | | 2 | 1 |
| 759814218 | | | | | | | | | | | | | | | | | | | | | | | 1 |
| 760522417 | | | | | | | | | | | | | 1 | | | | | | | | | | |
| 760638547 | | | | | | | | | | | | | | | | | | | | | 2 | | |
| 760733959 | | | | | | | | 1 | | | | | | | | | | | | | | | |
| 760828908 | | | | | | | | 1 | | | | | | | | | | | | | | | |
| 761182838 | | | | | | | | 2 | 1 | | | | | | | | | | | | | | |
| 761489398 | | | | | | | | | 2 | | | | | | | | | | | | | | |
| 761494051 | | | | | | | | 2 | | | | | | | | | | | | | | | |
| 761838951 | | | | | | | | | 3 | | | | | | | | | | | | | | |
| 761843188 | | | | | | | | 1 | | | | | | | | | | | | | | | |
| 762176004 | | | | | | | | 1 | | | | | | | | | | | | | | | |
| 763435729 | | | | | | | | | | | | | | | | | | | 1 | | | | |
| 764817903 | | | | | | | | | | | | | | | | | | | | | 1 | | |
| 766385396 | | | | | | | | 1 | | | | | | | | | | | | | | | |
| 766389431 | | | | | | | | 1 | | | | | | | | | | | | | | | |
| 766734991 | | | | | | | | 1 | | | | | | | | | | | | | | | |
| 767473384 | | | | | | | | | 1 | | | | | | | | | | | | | | |
| 769544445 | | | | | | | | | | | | | | | | | | | | 1 | | | |
| 787227302 | | | | | | | | | | | | 2 | 2 | | | | | | | | | | |
| 787563782 | | | | | | | | | | | | 1 | | | | | | | | | | | |
| 787564302 | | | | | | | | | | | | 8 | 3 | | | | | | | | | | |
| 787882939 | 1 | 3 | | 2 | | | | | | 1 | | | | 3 | | 1 | 7 | 2 | 6 | 1 | | | |
| 787883174 | | | | | | | | 1 | 2 | 3 | | | | | | | 4 | 1 | | 1 | 8 | 13 | 13 |
| 787883482 | | | | | | | | | 1 | 1 | | | | 1 | | | 4 | 2 | 1 | | 4 | 9 | 18 |
| 787904798 | | | | | | | | | | | | 1 | | | | | | | | | | | |
| 787908929 | | | | | | | | | | | | 9 | 4 | | | | | | | | | | |
| 787909611 | | | | | | | | | | | | | 1 | | | | | | | | | | |
| 789330613 | | | | | | 1 | 1 | | | | | | | | 1 | | 1 | 1 | 1 | | 1 | 26 | 19 |
| 789851418 | | | | | | | | | | | | | | | | 1 | | | | | | | |
| 789934899 | | | | | | | | | | | | | 1 | | | | | | | | | | |
| 790305346 | | | | | | | | | | | | | 1 | | | | | | | | | | |
| 791738711 | | | | | | | | | | | | | | 1 | 3 | | 2 | 1 | 1 | 1 | | | 1 |
| 792692885 | | | | | | 1 | | | | | | 1 | 3 | | | | 1 | | 1 | | 2 | | 1 |
| 794104770 | | | | | | | | | | | | | 1 | | | | | | | | | | |
| 794574805 | | | | | | | | | 1 | | | | | | | | | | | | | | |
| 795546494 | 1 | | 1 | 1 | | | | | | | | | | | | | | | | | | | |
| 796733296 | | | | | | | | 4 | 1 | | | | | | | | | | | | | | |
| 801257348 | | | | | | | | | | | | | | | | | | | | | 1 | | |
| 818577473 | | | 1 | | | | | 1 | 1 | 1 | 1 | | | | | | 1 | | 3 | | | | |
| 818931040 | | | | | | | | | | | | 1 | | | | | | | | 1 | | | |
| 819992660 | | | | | | | | | 1 | | | | | | | | | | | | | | |
| 820015203 | | | | | | | | | | | | | | | | | | | | | | | 1 |
| 820334823 | | | | | | | | | | | | | | | | | | | | | 1 | | |
| 821007911 | | | | | | | | | | 2 | | | | | | | | | | | | | |
| 821702605 | | | | | | | | | 1 | | | | | | | | | | | | | | |
| 821780859 | | | | | | | | | | | 1 | | | | | | | | | | | | 1 |
| 822833681 | | | | | | | | 1 | | | | | | | | | | | | | | | |
| 823063281 | | | | | | | | | | | 2 | | | | | | | | | | | | |
| 824115747 | | | | | | | | 1 | 1 | | | | | | | | | | | | | | |
| 827262627 | | | | | | | | | | | | | | | | | | | | | | 1 | |
| 828282250 | | | | | | | 3 | 3 | 2 | | | | | | | | | | | | | | |

| | | | | | | | | | | | | | | | | | | | | | | | | | | | |
|---|---|---|---|---|---|---|---|---|---|---|---|---|---|---|---|---|---|---|---|---|---|---|---|---|---|---|---|
| 829141094 | | | | | | | | | | | | | | | 1 | | | | | | | | | | | | |
| 849270788 | | | | | | | | | 1 | 4 | 2 | 4 | 5 | 1 | 1 | 1 | 2 | 5 | 2 | 6 | 14 | 11 | 33 | 34 | | | |
| 849612438 | 2 | 1 | 1 | | | | | | | | | | 2 | 1 | | | | | | | | | 3 | | | | |
| 850294342 | | | | | | | | | | | | | | | | 2 | | | | | | | | | | | |
| 850703925 | | | | | | | | | | | | | | | | | 1 | | | | | | | | | | |
| 850717220 | | | | | | | | | | | | 1 | | 3 | 1 | | 1 | | | | | | 4 | 1 | 1 | | |
| 851459972 | | | | | | | | | | | | | | 1 | | 2 | 1 | | 1 | | | | 1 | | 3 | | 2 |
| 851727478 | | | | | | | | | | | | | | | | | | | | | | | | | 1 | | |
| 852046834 | | | | | | | | | | | | | | 4 | | | | | | | | | | | | | |
| 853376855 | | | | | | | | | | | | | | | | 1 | | | | | | | | | | | |
| 853717974 | | | | | | | | | | | | | | | | | 1 | | | | | | | | | | |
| 853726809 | | | | | | | | | | | | | | | | | 2 | | | | | | | | | | |
| 853791166 | | | | | | | | | | | | 1 | 2 | | | | | | | | | | | | | | |
| 855043038 | | | | | | | | | | 1 | | | | | | | | | | | | | | | | | |
| 859006021 | | | | | | | | | | | | 2 | | | | | | | | | | | | | | | |
| 859152863 | | | | | | | | | | | | 1 | 1 | | | | | | | | | | | | | | |
| 859265651 | | | | | | | | | | | | | | | | | | | | | | | | | 2 | | |
| 880305613 | | | 2 | | | | | | | | | | | | | | | | | | | | | | | | |
| 880987657 | | | | | | | | | | | | 4 | | 3 | | 10 | 12 | | | | | | | | 1 | | |
| 880987844 | | | | | | | | | | | | | | 1 | | 3 | 1 | | | | | | | | | | |
| 884140963 | | | | | | | | | | | | 1 | 1 | | | | | | | | | | | | | | |
| 886130319 | | | | | | | | | | | | | | | | | 2 | | | | | | | | | | |
| 889497360 | | | | | | | | | | | | 1 | | | | | | | | | | | | | | | |
| 890883360 | | | | | | | | | | | | 1 | | | | | | | | | | | | | | | |
| 911332304 | | | | | 3 | 5 | | 1 | | | | | | 37 | 55 | | 1 | | | | | | | | | | |
| 911681895 | | | | | | | | | | | | | | | | 1 | | | | | | | | | | | |
| 912045178 | | | | | | | | | | | | | | | | 4 | 2 | | | | | | | | | | |
| 912398383 | | | | | | | | | | | | | | | | | 1 | | | | | | | | | | |
| 912601209 | | | | | | | | | | | | | | 1 | | | | | | | | | | | | | |
| 914492772 | | | | | | | | | | | | | 1 | | | | | | | | | | | | | | |
| 914492942 | | | | | | | | | | | | 1 | 1 | | 1 | | | | | | | | | | | | |
| 915511932 | | | | | | | | | | | | | 1 | 1 | 2 | | | | | | | | 1 | | | | |
| 915852626 | | | | | | | | | | | | | | 1 | | | | | | | | | | | | | |
| 915866098 | | | | | | | | | | | | | 1 | | | | | | | | | | | | | | |
| 917656984 | | | | | | | | | | | | | | | | | | | | | | | | 2 | | | |
| 921589471 | | | | | | | | | | | | | 1 | | | | | | | | | | | | | | |
| 921935093 | | | | | | | | | | | | 4 | | | | | | | | | | | | | | | |
| 921939924 | | | | | | | | | | | | 1 | | | | | | | | | | | | | | | |
| 922595498 | | | | | | | | | | | | | 1 | | | | | | | | | | | | | | |
| 943071287 | | | | | | | | | | | | | | | | 3 | 2 | | | | | | | | | | |
| 943822621 | | | | | 1 | | | 1 | | | | | | | 1 | | | | | | | | | | | | |
| 944146138 | | | | | | | | | | | | | | | 1 | | | | | | | | | | | | 1 |
| 945161760 | | | | | | | | | | | | | | | | | | | | | | | | | | 1 | |
| 946204816 | | | | | | | | | | | | 2 | 2 | | | | | | | | | | | | | | |
| 946214177 | | | | | | | | | | | | | | | 1 | | | | | | | | | | | | |
| 946546270 | | | | | | | | | | | | 1 | 4 | | | | | | | | | | | | 1 | | |
| 946550009 | | | | | | | | | | | | | | 1 | | | | | | | | | | | | | |
| 947193705 | | | | | | | | | | | | 1 | | | | | | | | | | | | | | | |
| 948363764 | | | | | | | | | | | | | | | 1 | | | | | | | | | | | | |
| 948812702 | | | | | | | | | | | | | | | | | | | | | | | | | 3 | | |
| 949045811 | | | | | | | | | | | | | 1 | | 1 | | | | | | | | | | | | |
| 950229431 | | | | | | | | | | | | | | | | | | | | | | | 1 | | | | |
| 950591319 | | | | | | | | | | | | | | | | | | | | | | | | | 3 | | |
| 951738965 | | | | | | | | | | | | | | | | | | | | | | | | | 1 | | |
| 951934694 | | | | | | | | | 1 | | | | | | | | | | | | | | | | | | |
| 952792408 | | | | | | | | | | | 1 | 1 | | | | | | | | | | | | | | | |
| 973423536 | | | | | | | | | | | | | | | | 1 | | | | | | | | | | | |
| 974498400 | | | | | | | | | | | | | | 2 | 1 | | | | | | | | | | | | |
| 974498428 | | | | | | | | | | | | | | 2 | 1 | | | | | | | | | | | | |
| 974498485 | | | | | | | | | | | | | | 9 | 3 | | | | | | | | | | | | |
| 974498491 | | | | | | | | | | | | | | 10 | 2 | | | | | | | | | | | | |
| 974498492 | | | | | | | | | | | | | | 1 | 1 | | | | | | | | | | | | |
| 974839580 | | | | | | | | | | | | | | 2 | 1 | | | 1 | | 1 | 3 | | | 1 | 14 | 24 | 20 |
| 974839683 | | | | | | | | | | | | | | | 3 | | | | | | | | | | | | |
| 974843957 | | | | | | | | | | | | | | 14 | 14 | | 5 | | | | | | | | | | |
| 975872458 | | | | | | | | | | | | | | | | | | | | | | | | | 1 | | |
| 976730002 | | | | | | | | | | | | | | | | | | | | | | | | | | 1 | |
| 977710985 | | | | | | | | | | | | | | | | | | | | | | | | 1 | | | |
| 977934593 | | | | | | | | | | | | | | | | | | | | | | | | | | | 3 |
| 978336888 | | | | | | | | | | | | | | | | | | | | | | | 1 | | | | |
| 982110108 | | | | | | | | | | | | | | | | | | | | | | | 1 | | | | |
| 982994156 | | | | | | | | | | | | | | | | | | | | | | | | 1 | | | |
| 983146199 | | | | | | | | | | | | | | | | | | 1 | | 2 | 1 | | | | | | |
| 984018803 | | | | | | | | | | | | | | | | | | | | | | | | 1 | | | |
| 987759904 | | | | | | | | | | | | | | | | | | | | | | | | 1 | | | |
| 1005102825 | | | | | | | | | | | | | | | | | | | | | | | | | 1 | | |
| 1005119041 | | | | | | | | | | | | 1 | | | | | | | | | | | | | | | |
| 1005174948 | | | | | | | | | | | | | | 1 | | | | 1 | | | 2 | | | | 5 | 8 | 10 |
| 1005520576 | | | | | | | | | | | | | | 1 | | | | | | | | | | | | | |
| 1005542502 | | | | | | | | | | | | | | 1 | 1 | | | | | | | | | | | | |

| | | | | | | | | | | | | | | | | | | | | | | | | | | | | |
|---|---|---|---|---|---|---|---|---|---|---|---|---|---|---|---|---|---|---|---|---|---|---|---|---|---|---|---|---|
| 1005857873 | | | | | | | | | | | | | | | | | 1 | | | | | | | | | | | |
| 1005857944 | | | | | | | | | | | | | | | | | | | | | | | | | 1 | | 1 | |
| 1006199303 | | | | | | | | | | | | | | | | | | | | | | | | | | 1 | | |
| 1006539760 | | | | | | | | | | | | | | 2 | 2 | | | | | | | | | | | | | |
| 1007402796 | | | | | | | | | | | | | 1 | | | | | | | | | | | | | 1 | | |
| 1010731508 | | | | | | | | | | | | | | | | | | | | | | | 1 | 1 | | | | |
| 1014327691 | | | | | | | | | | | | 1 | | | | | | | | | | | | | | | | |
| 1036546847 | | | | | | | | | | | | | | 4 | | | | | | | | | | | | | | |
| 1036550962 | | | | | | | | | | 3 | 1 | | | | | | | | | | | | | 1 | | | | |
| 1036637638 | | | | | | | | | | | | | | | | | | | | | | | | 1 | | | | |
| 1037915305 | | | | | | | | | | | | | | 1 | | | | | | | | | | | | | | |
| 1041541944 | | | | | | | | | | | | | | | | | | | | | | | | | | | | 1 |
| 1042495376 | | | | | | | | | | | | | | | | | | | | | | | 2 | 1 | | | | |
| 1046039987 | | | | | | | | | | | | | | | | | | | | | | | 1 | | | | | |
| 1046385221 | | | | | | | | | | | | | | | | | | | | | | | 2 | | | | | |
| 1046726043 | | | | | | | | | | | | | | | | | | | | | | | 1 | | | | | |
| 1046726280 | | | | | | | | | | | | | | | | | | | | | | | 2 | | | | | |
| 1047075623 | | | | | | | | | | | | | | | | | | | | | | | | 1 | | | | |
| 1047412496 | | | | | | | | | | | | | | | | | | | | | | | 1 | | | | | |
| 1047425857 | | | | | | | | | | | | | | | | | | | | | | | 1 | | | | | |
| 1066511487 | | | | | 1 | | 1 | | | | | 28 | 28 | 135 | 63 | 7 | 10 | 1 | | 4 | 7 | | 1 | 2 | 3 | 3 | 1 | |
| 1072063538 | | | | | | | | | | | | 1 | | 5 | 3 | | 1 | | | | | | | | 1 | | | |
| 1072805036 | | | | | | | | | | | | | | | | | | | | | | | | 1 | | | | |
| 1076724912 | | | | | | | | | | | | | | | | | | | | | | | 4 | 3 | | | | |
| 1077122311 | | | | | | | | | | | | | | | | | | | | | | | | 1 | | | | |
| 1077415831 | | | | | | | | | | | | | | | | | | | | | | | 1 | | | | | |
| 1077415919 | | | | | | | | | | | | | | | | | | | | | | | 2 | 1 | | | | |
| 1077420174 | | | | | | | | | | | | | | | | | | | | | | | 2 | 2 | | | | |
| 1077756731 | | | | | | | | | | | | | | | | | | | | | | | 1 | | | | | |
| 1077756950 | | | | | | | | | | | | | | | | | | | | | | | | 1 | | | | |
| 1077761277 | | | | | | | | | | | | | | | | | | | | | | | | 1 | | | | |
| 1077761302 | | | | | | | | | | | | | | | | | | | | | | | 5 | 3 | | | | |
| 1078102419 | | | | | | | | | | | | | | | | | | | | | | | 1 | | | | | |
| 1078102490 | | | | | | | | | | | | | | | | | | | | | | | | 1 | | | | |
| 1078443080 | | | | | | | | | | | | | | | | | | | | | | | 2 | 1 | | | | |
| 1078447707 | | | | | | | | | | | | | | | | | | | | | | | 4 | 5 | | | | |
| 1078788519 | | | | | | | | | | | | | | | | | | | | | | | 1 | | | | | |
| 1096872469 | | | | | | | | | | | | | | | | | | | | | | | | 1 | | | | |
| 1096959028 | | | | | | | | | | | | | | 2 | | | | | | | | | | | | | | |
| 1104004164 | | | | | | | | | | | | | | | | | | | | | | | 1 | 1 | | | | |
| 1107081827 | | | | | | | | | | | | | | | | | | | | | | | | | 1 | | | |
| 1107423142 | | | | | | | | | | | | | | | | | | | | | | | 1 | | | | | |
| 1107423158 | | | | | | | | | | | | | | | | | | | | | | | | 1 | | | | |
| 1107764206 | | | | | | | | | | | | | | | | | | | | | | | 1 | 1 | | | | |
| 1108109641 | | | | | | | | | | | | | | | | | | | | | | | 1 | | | | | |
| 1108109684 | | | | | | | | | | | | | | | | | | | | | | | 1 | | | | | |
| 1108113869 | | | | | | | | | | | | | | | | | | | | | | | 3 | 1 | | | | |
| 1108446169 | | | | | | | | | | | | | | | | | | | | | | | | 1 | | | | |
| 1108809626 | | | | | | | | | | | | | | | | | | | | | | | 1 | 1 | | | | |
| 1109478369 | | | | | | | | | | | | | | | | | | | | | | | 1 | | | | | |
| 1109832699 | | | | | | | | | | | | | | | | | | | | | | | 1 | 1 | | | | |
| 1110173530 | | | | | | | | | | | | | | | | | | | | | | | 1 | | | | | |
| 1110173958 | | | | | | | | | | | | | | | | | | | | | | | 1 | 1 | | | | |
| 1127635306 | | | | | | | | | | | | | | | 3 | | | | | | | | | | | | | |
| 1127640198 | | | | | | | | | | | | | | 2 | | | | | | | | | | | | | | |
| 1127886209 | | | | | | | | | | | | 1 | 2 | 14 | 6 | | 2 | | | | | | | | | | | |
| 1127894310 | | | | | | | | | | | | 1 | 1 | 12 | 5 | 1 | | | | | | | | | | | | |
| 1127894824 | | | | | | | | | | | | 3 | 1 | 14 | 8 | 1 | 2 | | | | | | | | 1 | | | |
| 1128227087 | | | | | | | | | | | | | | | 1 | | | | | | | | | | | | | |
| 1128274593 | | | | | | | | | | | | 1 | | 6 | 17 | | | | | | | | 1 | | | | | |
| 1128572709 | | | | | | | | | | | | | | | | | | | | | | | 1 | | | | | |
| 1128573330 | | | | | | | | | | | | | | 1 | 1 | | | | | | | | | | | | | |
| 1129033939 | | | | | | | | | | | | | | 1 | | 2 | | | | | | | | | | | | |
| 1129306180 | | | | | | | | | | | | | | 1 | | | | | | | | | | | | | | |
| 1137081105 | | | | | | | | | | | | | | | | | | | | | | | 1 | 1 | | | | |
| 1137767646 | | | | | | | | | | | | | | | | | | | | | | | 3 | 3 | | | | |
| 1137767694 | | | | | | | | | | | | | | | | | | | | | | | 2 | 4 | | | | |
| 1137767724 | | | | | | | | | | | | | | | | | | | | | | | | 2 | | | | |
| 1137780791 | | | | | | | | | | | | | | | | | | | | | | | 1 | | | | | |
| 1138453934 | | | | | | | | | | | | | | | | | | | | | | | 3 | 2 | | | | |
| 1138462517 | | | | | | | | | | | | | | | | | | | | | | | 2 | | | | | |
| 1138807874 | | | | | | | | | | | | | | | | | | | | | | | 1 | | | | | |
| 1139144718 | | | | | | | | | | | | | | | | | | | | | | | 2 | 1 | | | | |
| 1139494179 | | | | | | | | | | | | | | | | | | | | | | | | 1 | | | | |
| 1139826933 | | | | | | | | | | | | | | | | | | | | | | | | 2 | | | | |
| 1140012085 | | | | | | | | | | | | | | | | | | | | | | | | 1 | | | | |
| 1140168071 | | | | | | | | | | | | | | | | | | | | | | | 1 | | | | | |
| 1140176410 | | | | | | | | | | | | | | | | | | | | | | | 3 | | | | | |
| 1140180513 | | | | | | | | | | | | | | | | | | | | | | | 1 | | | | | |
| 1140188920 | | | | | | | | | | | | | | | | | | | | | | | 1 | | | | | |

| | | | | | | | | | | | | | | | | | | | | | | | | | | | |
|---|---|---|---|---|---|---|---|---|---|---|---|---|---|---|---|---|---|---|---|---|---|---|---|---|---|---|---|
| 1140853994 | | | | | | | | | | | | | | | | | | | | | | | 1 | | | | |
| 1140863129 | | | | | | | | | | | | | | | | | | | | | | | 2 | 1 | | | |
| 1141035086 | | | | | | | | | 3 | | | | | | | | | | | | | | | | | | |
| 1141545043 | | | | | | | | | | | | | | | | | | | | | | | 1 | | | | |
| 1158631810 | | | | | 1 | | | | | | | | | | | | | | | | | | | | | | |
| 1158635682 | | | | | | | | | | | | | | | | | | | | | | | 2 | | | | |
| 1158636183 | | | | | | | | | | | | | | 1 | | | | | | | | | | | | | |
| 1158644769 | | | | | 1 | | | | | | | | | | | | | | | | | | | | | | |
| 1158657609 | | | | | | | | | | | | | | | 1 | | | | | | | | | | | | |
| 1159262075 | | | | | | | | 2 | | | | | | | | | | | | | | | | | | | |
| 1159607159 | | | | | | | | | | | | | | 1 | | | | | | | | | | | | | |
| 1159607813 | | | | | | | | | | | | | | 4 | | | | | | | | | | | | | |
| 1159607872 | | | | | | | | | | | | | | 2 | | | | | | | | | | | | | |
| 1159611751 | | | | | | | | | | | | | | 1 | | | | | | | | | | | | | |
| 1159611765 | | | | | | | | | | | | | | 3 | 2 | | | | | | | | | | | | |
| 1159611828 | | | | | | | | | | | | | | 1 | | | | | | | | | | | | | |
| 1159948125 | | | | | | | | | | | | | | 1 | 1 | | | | | | | | | | | | |
| 1159952969 | | | | | | | | | | | | | | | 1 | | | | | | | | | | | | |
| 1159953225 | | | | | | | | | | | | | | | 2 | | | | | | | | | | | | |
| 1167779265 | | | | | | | | | | | | | | | | | | | | | | | 1 | | | | |
| 1169169208 | | | | | | | | | | | | | | | | | | | | | | | | | | 1 | |
| 1170179364 | | | | | | | | | | | | | | | | | | | | | | | 1 | | | | |
| 1170866180 | | | | | | | | | | | | | | | | | | | | | | | 1 | | | | |
| 1172929315 | | | | | | | | | | | | | | | | | | | | | | | 3 | | | | |
| 1190339995 | | | | | | | | | | | | | | 5 | 3 | | 2 | | | | | | | | | | |
| 1190340083 | | | | | | | | | | | | | | 3 | 2 | | | | | | | | | | | | |
| 1190344036 | 1 | | | | | | | | | | | | | | | 1 | | | | | | | | | | | |
| 1190638276 | | | | | | | | | | | | | | | 1 | | | | | | | | | | | | |
| 1190668746 | | | | | | | | | | | | | | | | | | | | | | | | | 7 | | |
| 1198132041 | | | | | | | | | | | | | | | 1 | | | | | | | | | | | | |
| 1199491728 | | | | | | | | | | | | | | | | | | | | | | | | 1 | | | |
| 1200178237 | | | | | | | | | | | | | | | | | | | | | | | | 1 | | | |
| 1200527805 | | | | | | | | | | | | | | | | | | | | | | | 1 | | | | |
| 1202250779 | | | | | | | | | | | | | | | | | | | | | | | 1 | | | | |
| 1203774508 | | | | | | | | | | | | | | | | | | | | | | | | | | 1 | |
| 1220049442 | | | | | | | | | | | | | | | | | | | | | | | 3 | | | | |
| 1225605932 | | | | | | | | | | | | | | | | | | | | | | | 2 | | | | |
| 1226029612 | | | | | | 2 | | | | | | | | | 1 | | | | | | | | | | | | |
| 1231213221 | | | | | | | | | | | | | | | | | | | | | | | 3 | 2 | | | |
| 1234662524 | | | | | | | | | | | | | | | | | | | | | | | 1 | | | | |
| 1252409535 | | | | | | | 2 | | | | | | | | | | | | | | | | | | | | |
| 1283405893 | 1 | | | | | | | | | | | | | | | 2 | 5 | | | | | | | | | | |
| 1283406432 | | | | 2 | 25 | 19 | 24 | 29 | | | | | | 14 | 33 | | | | | | | | | | | | |
| 1284117939 | | | | | | | | | | | | | | | | 9 | 2 | | | | | | | | | | |
| 1284373356 | | | | | | | | | | | | | | | | | 2 | | | | | | | | | | |
| 1313115480 | | | | | | | | 1 | | | | | | | 1 | | 1 | | | | | | | | | | |
| 1313248733 | | | | | | | | | | | | | | | | | | | | | | | | | 1 | | |
| 1313538112 | | | | | | 2 | | 1 | | | | | | | | | | | | | | | | | | | |
| 1313733623 | | | | | | | | | | | | | | 1 | | | | | | | | | | | | | |
| 1314130144 | | | | | | | | | | | | | | | 2 | | | | | | | | | | | | |
| 1315469002 | | | | | | | | | | | | | | 4 | | | | | | | | | | | | | |
| 1317130193 | | | | | | | | | | | | | | 2 | 2 | | | | | | | | | | | | |
| 1317471331 | | | | | | | | | | | | | | 1 | 1 | | | | | | | | | | | | |
| 1320177531 | | | | | | | | | | | | | | 1 | | | 1 | | | | | | | | | | |
| 1344828428 | | | | | | | | | | | | | | | 1 | | | | | | | | | | | | |
| 1345164753 | | | | | | | 2 | 1 | | | | | | | | | | | | | | | | | | | |
| 1345493520 | | | | | | | | | | | | | | 1 | | 5 | 2 | | | | | | | | | | |
| 1346477190 | | | | | | | | | | | | | | | | | | | | | | | | 2 | | | |
| 1346814892 | | | | | 6 | | 2 | 4 | | | | 2 | | 2 | 3 | 1 | | | | | | | | | | | |
| 1347638401 | | | | | | | | | | | | | | | 1 | 4 | 3 | | | | | | | | | | |
| 1348510469 | | | | | | | | | | | | | | 6 | 4 | | | | | | | | | | | | |
| 1348510522 | | | | | | | | | | | | | 1 | 5 | | | | | | | | | | | | | |
| 1375206889 | | | | | | | | | | | | | | | | | | | | | | | | | | 1 | |
| 1375841046 | | | | | | | | | | | | | | | 1 | | | | | | | | | | | | |
| 1376203864 | | | | | | | | | | | | | | | 1 | | | | | | | | | | | | |
| 1378182447 | | | | | | | | | | | | | | 1 | | | | | | | | | | | | | |
| 1378212488 | | | | | 5 | 1 | 3 | 12 | | | | | | 27 | 26 | | | | | | | | | | | | |
| 1383292291 | | | | | | | | | | | | | | | | | | | | | | | 1 | | | | |
| 1407558431 | | | | | | | | | | | | | | | | 1 | | | | | | | | | | | |
| 1414327340 | | | | | | | | | | | | | | | | | | | | | | | 1 | | | | |
| 1418618235 | | | 2 | 2 | | | 2 | | | | | | | | 1 | | | | | | | | | | | | |
| 1436335458 | | | | | | | | | | | | | | 1 | | | | | | | | | | | | | |
| 1439621300 | | 1 | | | | | | 3 | | | | | | | | | | | | | | | | | | | |
| 1448118399 | | | | | | | | | | | | | 1 | | | | | | | | | | | | | | |
| 1467374847 | | | | | | | | | | | | | | 1 | | | | | | | | | | | | | |
| 1481736987 | | | | | | | | | | | | | | 1 | 4 | | | 1 | 2 | | | 1 | | | | 2 | 1 |
| 1498410212 | | | | | | | | | | | | | | 1 | | | | | | | | | | | | | |
| 1501332510 | | | | | 23 | 8 | 15 | 21 | | | | | | 19 | 24 | | | | | | | | | | | | |
| 1501392572 | | | | | | | | | | | | | | | | | | | | | | | 1 | 2 | | | |
| 1505209641 | | | | | | | 1 | | | | | | | | | | 1 | | | | | | | | | | |

| | | | | | | | | | | | | | | | | | | | | | | |
|---|---|---|---|---|---|---|---|---|---|---|---|---|---|---|---|---|---|---|---|---|---|---|
| 1512246869 | | | | | | | | | | 1 | | | | | | | | | | | | |
| 1535709211 | | | | | 1 | | 2 | 2 | | | | | | | | | | | | | | |
| 1563419167 | | | | | 1 | | 1 | | | | | | 2 | | | | | | | | | |
| 1563929167 | | 1 | 1 | | 3 | 1 | 2 | 2 | | | | 1 | 1 | | | | | | | | | |
| 1570684665 | | | | | | | | | | | | | | 1 | 5 | | | | | | | |
| 1594117455 | | | | | 23 | 14 | 22 | 31 | | | | 8 | 9 | | | | | | | | | |
| 1622540715 | | | | | | 1 | | | | | | | | | | | | | | | | |
| 1625139569 | | | | | | | | | | | | | | 1 | | | | | | | | |
| 1653234875 | | | | | | | 1 | 1 | | | | | | | | | | | | | | |
| 1656523777 | | | | | | | 1 | | | | | | | | | | | | | | | |
| 1664980698 | 6 | | 3 | 1 | | | | | | | | | | | | | | | | | | |
| 1670597454 | | | | | | | | | | | | | | 8 | 6 | 2 | 1 | 2 | 3 | | | |
| 1671608112 | | | | | | | | | | | | | | 4 | 1 | | | | | | | |
| 1684256618 | | | | | | | | | | | | | | | 1 | | | | | | | |
| 1692278031 | | | | | | | | | | | | | | | | | | 3 | | | | |
| 1693486219 | | | | | | | | | | | | | | | 1 | | | | | | | |
| 1702306047 | | | | | | | | | | | | | | 1 | 2 | | | | | | | |
| 1727979406 | | | | | | | | | | | | | | | 2 | | | | | | | |
| 1749628974 | | | | 2 | | | | 10 | | | | | 1 | | | | | | | | | |
| 1781687187 | | | | | | | | | | | | 1 | | | | | | | | | | |
| 1810654510 | | | | | | | | | | | | | | | | | | 1 | | | | |
| 1812380613 | | | | | | | | | | | | | 1 | | | | | | | | | |
| 1838364577 | | | | | | | | | | | | | | 1 | | | | | | | | |
| 1849683877 | | | | | | | | | | | | | | 1 | | | | | | | | |
| 1850024931 | | | | | | | | | | | | | | 1 | | | | | | | | |
| 1858901026 | | | | | | | | | | 1 | | | | 117 | 88 | | | | | 58 | 29 | 34 |
| 1870241189 | | | | | | | 1 | | | | | | | | | | | | | | | |
| 1874450213 | | | | | | | | | | | | 1 | | | | | | | | | | |
| 1881059780 | | | | | | | | | | | | | | 2 | | | | | | | | 1 |
| 1881193508 | | | | | 5 | | 2 | | | | | | | | | | | | | | | |
| 1881401277 | | | | | | | | | | 1 | 1 | | | | 1 | | | | | | 1 | |
| 1881751117 | | | | | | | | | | | | | | | 1 | | | | | | | |
| 1884625521 | | | | | | 1 | 2 | 1 | | | | | 1 | | | | | | | | | |
| 1887163927 | | | | | | | | | | | | | | | 1 | | | | | | | |
| 1910886083 | | | | | | | | | | | | 1 | 4 | | | | | | | | | |
| 1912436017 | | | | | | | | | | | | | | | | | | | | | 7 | 8 |
| 1912789965 | | | | | | | | | | | | | | | 1 | | | | | | | |
| 1917866877 | | | | | | | | | | | | | | | 1 | | | | | | | |
| 1937164446 | | | | | | | | | | | | | | | | | | | 1 | | | |
| 1941243752 | | | | | | | | | | | | | | | | | | 2 | 4 | | | |
| 1941925302 | | | | | | | | | | | | | 1 | | | | | | | | | |
| 1943811736 | | | | | | | | | | | | | | 6 | 4 | | | | | | | |
| 1943812176 | | | | | | | | | | | | | | 2 | 2 | | | | | 1 | | 1 |
| 1943950268 | | | | | | | 3 | | | | | | | | | | | | | | | |
| 1944502935 | | | | | | | | | | | | | | 4 | 2 | | | | | | | |
| 1948107641 | | | | | | | | | | | | | | 1 | 2 | | | | | | | |
| 1962171889 | | | | | | | 1 | | | | | | | | | | | | | | | |
| 1968215498 | | | | | | | | | | 1 | | | | | | | | | | | | |
| 1974846717 | | | | | | | | | | | | | | | | | | 3 | | | 3 | 1 |
| 1975187675 | | | | | | | | | | | | | | 11 | 11 | | | | | | | |
| 1975347348 | 3 | | 5 | | | | | | | | | | | | | | | | | | | |
| 1975878958 | | | | | | | | | | | | | | 69 | 90 | | | | | | | |
| 2000606012 | | | | | | | | | | | | | | | | | | 3 | | | | |
| 2004665019 | | | | | | | 1 | | | | | | | | | | | | | | | |
| 2005005494 | | | | | 1 | | | | | | | | | | | | | | | | | |
| 2007068523 | 3 | 5 | | 5 | | | | | | | | | | | | | | | 1 | | | |
| 2007725158 | | | | | | | 1 | | | | | 1 | | | | | | | | | | |
| 2008061833 | 4 | 4 | 4 | | 19 | 7 | 19 | 17 | | | | 1 | | | | | | | | | | |
| 2024906500 | | | | | 2 | | | | | | | | | | | | | | | | | |
| 2031343357 | | | | | | | | | | | | | | | 1 | | | | | | | |
| 2035708242 | | | | | 2 | | | 2 | | | | | | | | | | | | | | |
| 2036048508 | | | | | 1 | | | | | | | | | | | | | | | | | |
| 2036730835 | | | | 1 | 4 | 2 | 2 | 5 | | | | | | | | | | | | | | |
| 2037067448 | | | | | | | | 3 | | | | 1 | | | | | | | | | | |
| 2038082105 | | | | | 3 | 1 | 2 | | | | | | | | | | | | | | | |
| 2039096834 | | | | | 5 | 1 | 2 | 3 | | | | | 2 | | | | | | | | | |
| 2039100722 | | | 1 | | 1 | 3 | 1 | 1 | | | | | | | | | | | | | | |
| 2064165421 | | | | | | | | | | | | | | | 1 | | | | | | | |
| 2064713562 | | | | | | | | | | | | | | | | | | 2 | 1 | | | |
| 2065745704 | | | | | 3 | | 1 | 1 | | | | | | | | | | | | | | |
| 2067088743 | | | | | 1 | | | 1 | | | | 1 | | | | | | | | | | |
| 2067748663 | | | | | | | | | | | | | 2 | | | | | | | | | |
| 2068443690 | | | | | 1 | | | | | | | | | | | | | | | | | |
| 2068780445 | | | | | | 2 | | 2 | | | | | | | | | | | | | | |
| 2068801704 | | 4 | 5 | 9 | | | | | 1 | | | | | | | | | 1 | 1 | | | |
| 2069117095 | 1 | 1 | 3 | | 16 | 11 | 17 | 14 | | | | | 3 | | | | | | | | | |
| 2069648663 | | | | | | | | | | | | | | 1 | 3 | | | | | | | |
| 2094397733 | | | | | | | | | | | | | | | | | | 4 | 2 | | | |
| 2097462638 | | | | | 1 | | 1 | 2 | | | | 9 | 16 | | | | | | | | | |
| 2098791953 | | | | | | | 1 | | | | | | | | | | | | | | | |

| | | | | | | | | | | | | | | | | | | | | | | | | | | | |
|---|---|---|---|---|---|---|---|---|---|---|---|---|---|---|---|---|---|---|---|---|---|---|---|---|---|---|---|
| 2098796739 | | | | | 1 | | | 3 | | | | | | | | | | | | | | | | | | | |
| 2099141470 | | | | | 1 | | | | | | | | | | | | | | | | | | | | | | |
| 2099146512 | 3 | 3 | 1 | 1 | 17 | 5 | 15 | 8 | | | | | | | | | | | | | | | | | | | |
| 2128838418 | | | | | | | 1 | | | | | | | | | | | | | | | | | | | | |
| 2129848864 | 3 | 3 | 1 | | 5 | 3 | 9 | 6 | | | | | | | | | | | | | | | | | | | |
| 2130509045 | | | | | | | | 3 | | | | | | | | | | | | | | | | | | | |
| 2155478189 | | | | | 1 | | | | | | | | | | | | | | | | | | | | | | |
| 2157831742 | | | | | | | | | | | | | | | | 1 | | | | | | | | | | | |
| 2161190280 | | | | | 1 | | | | | | | | | | | | | | | | | | | | | | |
| 2221273723 | | | | | | | | 1 | | | | | | | | | | | | | | | | | | | |
| 2222612442 | | | | | | | | | | | | | | | | 1 | | | | | | | | | | | |
| 2254986534 | | | | | 1 | | | | | | | | | | | | | | | | | | | | | | |
| 2289223313 | | | | | | | | | | | | | | | | | | | | | | | 1 | 1 | | | |
| 2315060417 | | | | | 3 | | | 2 | | | | | | | | | | | | | | | | | | | |
| 5812979048 | | | | | 1 | | | 1 | | | | | | 1 | | 1 | | | | | | | | | | | |
| 5812979935 | | | | | | | | | | | | 1 | | | | | | | | | | 1 | | | | | |
| 5812979938 | | | | | | | | | | 1 | 1 | | | | | | | | | | | | | | | | |
| 5812979995 | | | | | | | | | | 1 | | | | | | | | | | | | | | | | | |
| 5812980006 | | | | | | | | | | 17 | 13 | 1 | | | | | | 1 | | | 1 | | | 1 | | | 1 |
| 5812980010 | | | | | | | | | | | | 1 | | | | | | | | | | | | | 14 | 6 | 2 |
| 5812980016 | | | | | | | | | | | 1 | | | | | | | | | | | | | | | | |
| 5812980024 | | | | | | | | | | | | | | | | | | | | | | | | | 1 | | |
| 5812980047 | | | 2 | | | | | | | | | | | | 1 | | | | | | | | | | | | |
| 5812980079 | | | | | | | | | 1 | | | | | | | | | | | | | | | | | | |
| 5812980142 | | | | | | | | | 1 | 6 | 8 | 8 | 4 | | | | | | | | | | | | | | |
| 5812980250 | 2 | 1 | 5 | 3 | | | | | 1 | 1 | 4 | 4 | 2 | | | | | | 3 | 2 | 8 | 3 | | 1 | 2 | 3 | |
| 5812980253 | | | | | | | | | | | | | | | | | | | | | | | | 1 | 1 | | |
| 5812980270 | | | | | | | | | | 2 | 1 | | | | | | | | | | | | | | | | |
| 5812980289 | | | | | | | | | 1 | | | | | | 1 | 12 | 3 | | | | | | | | | 1 | |
| 5812980316 | | | 2 | | | | | | | | | | | | | | | | | | | | | | | | |
| 5812980473 | | | | | | | | | | | | 2 | | | | | | | | | | | | | | | |
| 5812980494 | | | | | | | | | | | | 2 | | | | | | | | | | | | | | | |
| 5812980501 | | | | | | | | | | | | | | | | | | | | | | | 1 | | | | |
| 5812980507 | | | | | | | | | | | | | 4 | 4 | 2 | | | | | | | | | | 1 | | 1 |
| 5812980513 | | | | | | | | | | | 3 | | | | | | 3 | | | | | | | | 1 | | |
| 5812980662 | | | | | | | | 1 | | | | | | 14 | 19 | | 2 | | | | | | | | | | |
| 5812980673 | | | | | | | | | | | | 1 | | | | | | | | | | | | | | | |
| 5812980710 | | | | | | | | | | | | | | | 1 | | | | | | | | | | | | 1 |
| 5812980742 | | | | | | | | | 1 | | | | | | | | | | | | | | | | | | |
| 5812980884 | | | | | | | | | | | | | 1 | | | | | | | | 1 | | | | | | |
| 5812980982 | | | 2 | | | | | | | | | | | | | | | | | | | | | | | | |
| 5812980984 | | | | | | | | | | | 1 | | | | | | | | | | | | | | | | |
| 5812981000 | | | | | | | | | | | | | 1 | | | | | | | | | | | | | | |
| 5812981016 | | | | | | | | | | | | 1 | 1 | | | 10 | 3 | | | | | | | 1 | | | |
| 5812981228 | | | | | | | | | | | | | 1 | | | | | | | | | | | | | | |
| 5812981237 | | | | | | | | | | | | | | | | | | | | | | | | | | 4 | 2 |
| 5812981460 | | | | | | | | | | | | 30 | 32 | 135 | 83 | 6 | 8 | 1 | | | 3 | 2 | 1 | 3 | 3 | 1 | |
| 5812981506 | | | | | | | | | | | | | | | | | | | | | | | | | | 1 | |
| 5812981517 | | | | | | | | | | | | | | | | | | | | | | | | | | 1 | |
| 5812981643 | | | | | | | | | | | | | | | | | | | | | | 1 | | | | | |
| 5812981715 | | | 2 | 2 | | | | | | | | | | | 4 | | | | | | | | | | | | |
| 5812981763 | | | | | | | | | | | | | | 2 | 1 | | | | | | | | | | | | |
| 5812982241 | | | | | | | | | | | | | | 1 | | | | | | | | | | | | | |
| 5812982278 | | | | | | | | | | | | | | 1 | 2 | | | | | | | | | | 1 | | |
| 5812982324 | | | | | | | | | | | | 4 | 2 | | | | | | | | | | | | 1 | 10 | 7 |
| 5812982566 | | | | | | | | | | | | | | 1 | 3 | | | | | | | | | | | | |
| 5812982573 | | | | 1 | | | | | | | | | | | | | | | | | | | | | | | |
| 5812982574 | | | | | | | | | | | | | | 1 | | | | | | | | | | | | | |
| 5812982632 | | | | | | | | | | | | | | | | | | | 2 | 3 | | | | | | | |
| 5812983533 | | | | | | | | | | | | | | | | | | | | | | | 1 | | | | |
| 5812983721 | | | | | | | | | | | | | | 1 | | | | | | | | | | | | | |
| 5812983722 | | | | | | | | | | | | | | 1 | | | | | | | | | | | | | |
| 5812983942 | | | | | | | | | | | | | | 2 | 2 | | | | | | | | | | | | |
| 5812983943 | | | | | | | | | | | | | | | 1 | | | | | | | | | | | | |
| 5812986257 | | | | | | | | | | | | | | | | | | | | | | | | 1 | | | |
| 5812986871 | | | | | | | | | | | | | | | | | | | | | | | | 1 | | | |
| 5812987418 | | | | | | | | | | | | | | 2 | | | | | | | | | | | | | |
| 5812987907 | | | | | | | | | | | | | | 1 | | | | | | | | | | | | | |
| 5812988181 | | | | | | | | | | | | | | | | | | | | | | | 1 | | | | |
| 5812990740 | | | | | | | | | | | | | | | | | | | | | | | 1 | | | | |
| 5812994948 | | | | | | | | | | | | 1 | | | | | | | | | | | | | | | |
| 5812995141 | | | | | | | | | | | | | 1 | | | | | | | | | | | | | | |
| 5812995298 | | | | | | | | | | | | | | | | 6 | 1 | 1 | 5 | 1 | 1 | 1 | 5 | 8 | 2 | | |
| 5812996678 | | | | | 5 | | | 1 | | | | | | | | 1 | 1 | | | | | | | | | | |
| 5812996681 | | | 1 | | 1 | 7 | 3 | | | | | | | | | | | | | | | | | | | | |
| 5812996748 | | | | | | | | | | | | | | | | | | 1 | | | | | | 1 | 1 | | |
| 5812997016 | | | | | | | 2 | | | | | | | | | | | | | | | | | | | | |
| 5813000129 | | | | | 1 | | 1 | 2 | | | | | | | | | | | | | | | | | | | |
| 5813000891 | | | | | | 1 | | 1 | | | | | | | | | | | | | | | | | | | |
| 5813001741 | | | | | 3 | 1 | | | | | | | | | 2 | | | | | | | | | | | | |

| | | | | | | | | | | | | | | | | | | | | | | | | | | | | |
|---|---|---|---|---|---|---|---|---|---|---|---|---|---|---|---|---|---|---|---|---|---|---|---|---|---|---|---|---|
| 5813002641 | | | | | | | | | | | | 1 | | | | | | | | | | | | | | | | |
| 5813002721 | | | | | 2 | 2 | 5 | 1 | | | | | | | | | | | | | | | | | | | | |
| 5813002722 | | | 1 | | 5 | 4 | 9 | 2 | | | | | | | | | | | | | | | | | | | | |
| 5813002810 | | | | | | | | | | | | | | | | | | 1 | | | | 4 | | | | | | |
| 5813002827 | | | | | | | | | | | | | | | | | | | | | | 1 | | | | | | |
| 5813002832 | | | | | | | | | | | | | | | | | | 1 | | | | | | | | | | |
| 5813005406 | | | | | 1 | | | | | | | | | | | | | | | | | | | | | | | |
| 5813007108 | | | | | | | | | | | | 3 | 1 | | | | | | | | | | | 1 | 2 | 5 | 3 | |
| 5813008796 | | | | | | | | | | | | | | | | | | | | | | | 1 | | | | | |
| 5813008892 | | | | | | | | | | | | | | | | | | | | | | 1 | | | 4 | | | |
| 5813008911 | | | | | | | | | | 1 | | | | | | | | | | | | | | | | | | |
| 5813008919 | | | | | | | | | | 2 | | | | | | | | | | | | | | | | | | |
| 5813008924 | | | | | | | | | 4 | 2 | | | | | | | | | | | | | | | | | | |
| 5813008983 | | | | | | | | | 1 | 2 | 1 | | | | | | | | | | | | | | | | | |
| 5813009260 | | | | | | | | | | | | | | | | | | | | | | | | 1 | | | | |
| 5813009352 | | | | | | | | | | 2 | 1 | | | | | | | | | | | | | | | | | |
| 5813009378 | | | | | | | | | | | | 1 | 2 | 1 | | | | | | | | | 1 | | | | | |
| 5813009391 | | | | | | | | | | 5 | 2 | | | | | | | | | | | | | | | | | |
| 5813009449 | | | | | | | | | | | | 2 | 1 | | | 1 | | | | | | | | | | | | |
| 5813009486 | | | | | | | | | | 1 | | 1 | | | | | | 2 | | | | | | | | | | |
| 5813009496 | | | | | | | | | 2 | | 2 | | | | | | | | | | | | | | | | | |
| 5813009497 | | | | | | | | | | 1 | | | | | | | | | | | | | | | | | | |
| 5813009500 | | | | | | | | | 4 | 1 | 4 | | | | | | | | | | | | | | | | | |
| 5813009589 | | | | | | | | | | | | | | | | | | | | | | | | | | | 1 | |
| 5813009609 | | | | | | | | | 2 | | | | | | | | | | | | | | | | | | | |
| 5813009670 | | | | | | | | | 1 | | | | | | | | | | | | | | | | | | | |
| 5813009676 | | | | | | | | | 2 | | | | | | | | | | | | | | | | | | | |
| 5813009705 | | | | | | | | | 5 | | | | | | | | | | | | | | | | | | | |
| 5813009725 | | | | | | | | | | | | | 1 | 1 | | | | | | | | | | | | | | |
| 5813009749 | | | | | | | | | | | | | | 3 | | | | | | | | | | | | | | |
| 5813009781 | | | | | | | | | | 1 | | | | | | | | | | | | | | | | | | |
| 5813009791 | | | | | | | | | | 1 | | | | | | | | | | | | | | | | | | |
| 5813009845 | | | | | | | | | | | | 1 | | | | | | | | | | 1 | | | | 1 | 1 | |
| 5813009881 | | | | | | | | | 1 | | | | | | | | | | | | | | | | | | | |
| 5813009882 | | | | | | | | | 2 | | | | | | | | | | | | | | | | | | | |
| 5813009905 | | | | | | | | | | | 1 | | | | | | | | | | | | | | | | | |
| 5813009946 | | | | | | | | | | 1 | | | | | | | | | | | | | | | | | | |
| 5813010120 | | | | | | | | | | | | | 1 | | | | | | | | | | 1 | 1 | | | | |
| 5813010121 | | | | | | | | | | | | | | | | | | | | | | | 1 | | | | | |
| 5813010135 | | | | | | | | | | | | | | | | | | | | | | | 1 | | | | | |
| 5813010139 | | | | 2 | | | | | | | | | | 1 | 1 | | | | | | | | | 1 | | | | |
| 5813010153 | | | | | | | | | | | | 9 | 6 | 15 | 25 | | | | 1 | | 1 | 2 | | 2 | 1 | 4 | 1 | |
| 5813010376 | | | | | | | | | | | | | 1 | | | | 2 | 1 | | | | | 1 | 2 | | 1 | | |
| 5813010377 | | | | | | | | | | | | | | | | | | | | | | | 1 | | | | | |
| 5813010385 | | | | | | | | | | | | | | | | | | | | | | | 3 | 1 | | | | |
| 5813010435 | | | | | | | | | | | | | | | | | | 1 | | | | | | 1 | 2 | | | |
| 5813010448 | | | | | | | | | 1 | | | | | | | | | | | | | | | | | | | |
| 5813010489 | | | | | | | | | | | | | 2 | | | | | | | | | | | | | | | |
| 5813010682 | | | | | | | | | | | | | | | | | | | | | | 1 | 2 | 3 | | | | |
| 5813010683 | 1 | 1 | | 1 | | | | | | | 1 | | | | | | | 8 | 9 | 6 | 4 | 2 | 3 | 4 | 1 | | | |
| 5813010690 | | | | | | | | | | | | 1 | | 1 | | | | 2 | | 3 | | 3 | 1 | 1 | 2 | | 1 | |
| 5813010721 | | | | | | | | | | | | | 1 | | | | | | | | | | | | | | | |
| 5813010738 | | | | | | | | | | | | | | | | | | 1 | | | | | | | | | | |
| 5813010766 | | | | | | | | | 1 | | 1 | | | | | | | | | | | | | | | | | |
| 5813010770 | | | | | | | | | | | | | | | | 1 | 1 | | | | | | | | | | | |
| 5813010948 | | | | | | | | | | | | | | | | | | | | | | | | | 1 | | | |
| 5813010996 | | | | | | | | | | | | | | | | | | | | 2 | | | | | | | | |
| 5813011015 | | | | | | | | | | 2 | 1 | | | | | | | | | | | | 1 | 3 | | | 2 | |
| 5813011061 | | | | | | | | | | | | | | | | | | | | | | | | | | | 1 | |
| 5813011066 | | | | | | | | | 1 | | | | | | | | | | | | | | | | | | | |
| 5813011067 | | | | | | | | | | | | | | | | | | | | | | | | | | | | 1 |
| 5813011133 | | | | | | | | | | | | | | | | | | | | | | | | | 1 | | | |
| 5813011327 | | | | | | | | | | | | | | | | | | | | 1 | | | | | | | | |
| 5813011384 | | | | | | | | | | | | | | | | | | | | | 1 | | | | | | | |
| 5813011423 | | | | | | | | | | | | 2 | 2 | | | | | | | | | | | | | | | |
| 5813011483 | | | | | | | | | | | | 2 | | | | | | | | | | | | | | | | |
| 5813011660 | | | | | 1 | | | | | 1 | 3 | | | | | | | | | | | | | | 1 | | | |
| 5813011683 | | | | | 42 | 14 | 26 | 46 | | | | | | 15 | 48 | 1 | | | | | | | | | | | | |
| 5813011709 | | | | | | | | | | | | | | | | | | | | 1 | | 1 | | | | | | |
| 5813011717 | | | | | | | | | | | | 2 | | | | | | | | | | | | | | | | |
| 5813012028 | | | | | | | | | | | | | | | | | | | | | | | | | 1 | | | |
| 5813012048 | | | | | | | | | | | | | | | | | | | | | | | | | 1 | | | |
| 5813012053 | | | | | | | | | | | | | 4 | 11 | 6 | | | | | | | | | 3 | | | | |
| 5813012126 | | | | | | | | | | | | | | | | | | | | | | | | | | | | 1 |
| 5813012366 | | | | | | | | | | | | | | | | | | | | | | | | | 6 | 3 | | |
| 5813012781 | | | | | | | | | | | | | | | | 2 | 1 | | | | | | | | | | | |
| 5813013091 | | | | | | | | | | | | | | | | | | | 1 | | | | | | | 1 | | |
| 5813013185 | | | | | | | | | | | | | | | | | | | | | | | | | | | | 3 |
| 5813013237 | | | | | | | | | | | | | 1 | | | | | | | | | | | | | | | |
| 5813013436 | | | | | | | | | | | | | | | | | | | | | | | 1 | | | | | |

| | | | | | | | | | | | | | | | | | | | | | | | | | | | | |
|---|---|---|---|---|---|---|---|---|---|---|---|---|---|---|---|---|---|---|---|---|---|---|---|---|---|---|---|---|
| 5813013965 | | | | | | | | | | | | 1 | | | | | | | | | | | | | | | | |
| 5813014166 | | | | | | | | | | | | | | | | 2 | | | | | | | | | | | | |
| 5813014218 | | | | | | | | | | | | | | | | 3 | 9 | | | | | | | | | | | |
| 5813014572 | | | | | 1 | | | 1 | | | | | | 2 | 1 | 1 | | | | | | | | 1 | | | 2 | |
| 5813014615 | | | | | | | | | | | | 1 | 2 | | | | | | | | | | | | | | | |
| 5813014765 | | | | | | | | | | | | 1 | | | | | | | | | | | | | | | | |
| 5813014910 | | | 1 | | | | | | | | | | | | | | | | | | | | | | | | | |
| 5813014911 | | | | | | | | | | | | | 1 | | | | | | | | | | | | | | | |
| 5813016450 | | | | | | | | | | | | | | | | | 1 | | | | | | | | | | | |
| 5813016743 | | | | | | | | | | | | | | | 2 | | | | | | | | | | | | | |
| 5813017614 | | | | | | | | | | | | | | 6 | 4 | | | | | | | | | | | | | |
| 5813019487 | | | | | | | | | 2 | | | | | | | | | | | | | | | | 1 | | | |
| 5813019517 | | | | | | | | | | 1 | | | | | | | | | | | | | | | | | | |
| 5813019526 | | | | | | | | | 2 | 1 | 2 | 2 | | | | | | | | | | | | | | | | |
| 5813019528 | | | | | | | | | | | | | | | | | | | | | | | | | 1 | 1 | 1 | |
| 5813019543 | | | | | | | | | | | | | | | | | | | | | | | 1 | | | | | |
| 5813019567 | | | | | | | | | 1 | | | | | | | | | | | | | | | | | | | |
| 5813019724 | | | | | | | | | | | | | | 16 | 5 | | | | | | | | 1 | 1 | | | | |
| 5813019868 | 2 | 2 | 1 | | | | | | 1 | | | | 2 | | | 2 | | 8 | 10 | 7 | 18 | 8 | 86 | 79 | 12 | | | |
| 5813020118 | | | | | | | | | | | | | | 18 | 9 | | 1 | | | | | | | 1 | | | | |
| 5813020136 | | | | | | | | | | | | | | | | 5 | 6 | | | | | | | | | | | |
| 5813020347 | | | | | | | | | | | | 1 | | | | | | | | | | | | | | | | |
| 5813020673 | | | | | | | | | | | | | | | | | | | | | | | | | 8 | 6 | 4 | |
| 5813020696 | | | | | | | | | | | | 1 | 1 | | | | | | | | | | | | | 1 | 1 | |
| 5813020785 | | | | | | | | | | 1 | | | | | | | | | | | | | | | | | | |
| 5813020840 | | | | | | | | | | | | | | | | | | | | | | | 1 | | | | | |
| 5813020969 | | | | | | | | | | | | | | | | | | | | | | | | | | 1 | | |
| 5813020993 | | | | | | | | | 1 | | | | | | | | | | | | | | | | | | | |
| 5813020996 | | | | | | | | | | | | | | | | | | | | | | | | | | | | 1 |
| 5813021094 | | | | | | | | | | 1 | 1 | | | | 1 | 3 | 5 | | | | | | 1 | 2 | 2 | 1 | | |
| 5813021192 | | | | | | | | | | | | 5 | | 1 | | | | | | | | | | | 2 | | 1 | |
| 5813021196 | | | | | | | | | | | | | | 1 | | | | | | | | | | | | | | |
| 5813021202 | | | | | 18 | 16 | 11 | 15 | | | | | | | | | | | | | | | | | | | | |
| 5813021217 | | | | | 13 | 8 | 10 | 26 | | | | | | 1 | 10 | | | | | | | | | | | | | |
| 5813021221 | | | | | | | | | | | | | | 2 | | | | | | | | | | | | | 1 | |
| 5813021225 | | | | | | | | | | | | 2 | 1 | | | | | | | | | | | | | | | |
| 5813021291 | | | | | | | | | | | | | | | | 3 | 6 | | | | | | | | | | | |
| 5813021336 | | | | | | | | | | | | | | | | | 1 | | | | | | | | | | | |
| 5813021480 | | | | | | | | | | | | | | 2 | | | | | | | | | | | | | | |
| 5813021485 | | | | | | | | | | | | 1 | | | | | | | | | | | | | | | | |
| 5813021535 | | | | | | | | | | | | 1 | | | | | | | | | | | 1 | | | | | |
| 5813021564 | | | | | | | | | | | | | | | | | | | | | | | | | 2 | | | |
| 5813021572 | | | | | | | | | | | | | | | | | | | | | | | | | 1 | | | |
| 5813021602 | | | | | | | | | | | | | | | | | | | | | | | | | 2 | | | |
| 5813021615 | | | | | | | 1 | 2 | | | | 1 | | 26 | 38 | 13 | 8 | | | | | | 1 | | | | | |
| 5813021632 | | | | | | | | | | | | | | | | | 1 | | | | | | | | | | | |
| 5813021666 | 1 | | | | | | | | | | | | | 81 | 71 | 2 | 1 | | | | | | | 1 | 1 | | | |
| 5813021671 | | | | | | | | | | | | | | 2 | 3 | 2 | | | 1 | | | | 1 | | | | | |
| 5813021882 | | | | | | | | | | | | | | 1 | | | | | | | | | | | | | | |
| 5813021894 | | | | | | | | | | | | | | | | 1 | | | | | | | | | | | | |
| 5813021899 | | | | | | | | | | | | | | | | | | | | | | | | 1 | | | | |
| 5813021943 | | | | | | | | | | | | | | | | | | | | | | | | | 1 | | | |
| 5813021948 | | | | | 11 | 2 | 2 | 11 | | | | 1 | | 2 | 3 | | | | | | | | | | | | | |
| 5813022150 | | | | | | | | | | | | | | 2 | | 1 | 1 | | | | | | | | | | | |
| 5813022182 | | | | | | | | | | | | | | 2 | | | 1 | | | | | | | 2 | | | 1 | |
| 5813022274 | | | | | | | | | | | | 1 | | 75 | 63 | 12 | | | | | | 1 | | | 3 | | 1 | |
| 5813022429 | | | | | | | | | | | | | | | | | | | | | | | | | | | | 1 |
| 5813022453 | | | | | | | | | | | | | | | | 1 | 5 | | | | | | | | | | | |
| 5813022753 | 18 | 26 | 23 | 22 | 23 | 16 | 25 | 16 | | | | | | 8 | 6 | | | | | | | | | | | | | |
| 5813022839 | 1 | | | | | | | | | | | | | 1 | | | | | | | | | | | | | | |
| 5813023097 | | | | 1 | | 1 | | | | | | | | | | | | | | | | | | | | | | |
| 5813024125 | | | 1 | | 18 | 8 | 13 | 16 | | | | | | | 1 | | | | | | | | | | | | | |
| 5813024789 | | | | | 1 | 1 | 1 | 3 | | | | | | | | | | | | | | | | | | | | |
| 5813024792 | | | 3 | | 4 | 2 | 7 | 2 | | | | | | | 1 | | | | | | | | | | | | | |
| 5813025330 | | | | | 1 | 1 | 1 | | | | | | | | | | | | | | | | | | | | | |
| 5813026385 | | | | | | | | | | | | | | | | | | | | | 1 | | | | | | | |
| 5813026579 | | | | | | | | | | | | 1 | | | | | | | | | | | | | | | | |
| 5813026589 | | | | | | | | | | | | 23 | 23 | | | | | | | | | | | | 1 | 14 | 12 | 1 |
| 5813026613 | | | | | | | | | | | | | 1 | | | | | | | | | | | | 2 | 8 | 10 | |
| 5813026615 | | | | | | | | | 1 | | | | | | | | | | | | | | | | | | | |
| 5813026661 | | | | | | | | | | | | | | | | | | | | | | | | | 1 | | | |
| 5813026718 | | | | | | | | | | | | | | | | | | | | | | | | | 1 | | | |
| 5813026773 | | | | | | 1 | 2 | | | | | | | | | | | | | | | | | | | | | |
| 5813026780 | | | | | | | | | | | | | | | | | | | | | | | 1 | | | | | |
| 5813027028 | | | | | | | | | | | | | | 1 | | | | | | | | | | | | | | |
| 5813027244 | | | | | | | | | | | | | | | | 12 | 16 | | | | | | | | 1 | | | |
| 5813027248 | | | | | 1 | | | | | | | | | | | | | | | | | | | | | | | |
| 5813032640 | | | | | | | | | 2 | 5 | 4 | | | | | | | | | | | | | | | | | |
| 5813032715 | | | | | 1 | | | | | | | | | | | | | | | | | | | | | | | |
| 5813032849 | | | | | | | | | | | | | 1 | | | | | | | | | | | | | | | |

| | | | | | | | | | | | | | | | | | | | | | | | | | | | | |
|---|---|---|---|---|---|---|---|---|---|---|---|---|---|---|---|---|---|---|---|---|---|---|---|---|---|---|---|---|
| 5813032924 | | | 1 | | | | | | | | | | | | | | | | | | | | 1 | | | | | |
| 5813033279 | | | | | 1 | | 2 | 1 | | | | | | | 3 | | | | | | | | | | | | | |
| 5813035064 | | | | | | | | | 3 | | 1 | | | | | | | | | | | | | | | | | |
| 5813035070 | | | | | | | | | | | | 1 | 1 | | | | | | | | | | | | | | | 13 |
| 5813035213 | | | | | | | | | | | | | | | | | | | | | | | | | 3 | 4 | 2 | |
| 5813035242 | | | | | | | | | | 2 | | | | | | | | | | | | | | | | | | |
| 5813035328 | | | | | | | | | 3 | | 2 | 4 | 7 | | | | | | | | | | | | | | | |
| 5813039660 | | | | | | | | | | | | | | 1 | | | | | | 1 | | | | | | | | |
| 5813039860 | | | | | | | | | | 1 | | | | | | | | | | | | | | | | | | |
| 5813039875 | | | | | | | | | | 5 | 1 | | | | | | | | | | | | 3 | | | | | |
| 5813039877 | | | | | | | | | | 1 | 2 | | | | | | | | | | | | | | | | | |
| 5813039902 | | | | | | | | | | | | | | | | | | | | | | | | | | | 1 | |
| 5813039910 | | | | | | | | | 1 | | | 1 | | | | | | | | | | | | | | | | |
| 5813039936 | | | | | | | | | | | | | | | | | | | | | 1 | | | | 1 | | 1 | |
| 5813039952 | | | | | | | | | | | | | | | | | | | | 1 | | | | | | | | |
| 5813040118 | | | | | | | | | | | | | | | | | | | | | | | | | | 1 | | |
| 5813040119 | | | | | | | | | | | 1 | 2 | | | | | | | | | | | | | | | | |
| 5813040216 | | | | | | | | | | | | | | | | | | | | | | | 4 | 2 | | | | |
| 5813040224 | | | | | | | | | | | | 1 | | | | | | | | | | | | | | | | |
| 5813040494 | | | | | 1 | 1 | 4 | 3 | | | | | | | 1 | | | | | | | | | | | | | |
| 5813040707 | | | | | | | | | 1 | | | | | | | | | | | | | | | | | | | |
| 5813040711 | | | | | | | | | | | | | | | | | | | 2 | 1 | 1 | | | | | | | |
| 5813040712 | | | | | | | | | | | | 22 | 23 | | | | | | | | | | | | | | | 1 |
| 5813040731 | | | | | | | | | | | | | | 2 | | | | | | | | | | | | | | |
| 5813040744 | 9 | 12 | 8 | 9 | 29 | 16 | 20 | 21 | | | | | | 11 | 3 | | 1 | | | | | | | | | | | |
| 5813041131 | | | | | | | | | | | | | | | | | | | | | | | 1 | | | | | |
| 5813041244 | | | | | | | | | | | | | | | | | 2 | | | | | | | | | | | |
| 5813041426 | | | | | | | | | | | | | | 2 | | | | | | | | | | | | | | |
| 5813044177 | | | | | | | | | | | | | | | | 1 | | | | | | | | | | | | |
| 5813045601 | | | | | | | | | | | | | | | 1 | | 1 | | | | | | | | | | | |
| 5813046572 | | 1 | | 2 | 10 | 6 | 11 | 10 | | | | | | | 1 | | | | | | | | | | | | | |
| 5813046674 | | | | | | | | | | | | | | | | | | | 1 | | | | | | | | | |
| 5813046951 | | | | | | | | | 1 | | | 1 | | | | | | | | | | | | | | | | |
| 5813046963 | | | | | | | | | | | | | | | | | | | | | | | | 1 | | | | |
| 5813046991 | | | | | | | | | | 1 | | | | | | | | | | | | | | | | | | |
| 5813047012 | | | | | | | | | | | | 1 | | | | | | | | | | | | | | | | |
| 5813047053 | | | | | | | | | | | | | | | | | | | | | | | 1 | | | | | |
| 5813047065 | | | 1 | | | | | | | | | | 2 | | | 2 | 1 | | 2 | | 3 | | 1 | | | | | |
| 5813047101 | | | | | | | | | | | | 1 | | | | | | | | | | | | | | | | |
| 5813047183 | | | | | | | | | 1 | | | | | | | | | | 1 | | | | | | | | | |
| 5813047188 | | | | | | | | | | | | | | | | | | | | | | | | | | 1 | | |
| 5813047189 | | | | | | | | | | | | | | | | | 1 | | | | | | | | | | | |
| 5813047284 | | | | | | | | | | | | 6 | 12 | 10 | 31 | | | 2 | 1 | 1 | | 1 | | | | 1 | 1 | |
| 5813047489 | | | | | | | | | | | | | | | | | | | | | | | | | 1 | | | |
| 5813047586 | 1 | 2 | 2 | | | | | | | | 1 | | | | | | | 11 | 5 | 20 | 4 | 10 | 1 | 4 | | | | |
| 5813047713 | | | | | | | | | 11 | 10 | 5 | | | | | | | | | | | | | | | | | |
| 5813047739 | | | | | | | | | 1 | 3 | 3 | 14 | 12 | 1 | | | | | | | | 1 | | | 50 | 8 | 3 | 14 |
| 5813047774 | | | | | | | | | | | | 6 | 1 | 1 | 1 | 3 | 3 | | | | 1 | 1 | | | | | | |
| 5813049105 | | | | | | | | | | | | | | | | | | | | | | | | | | 1 | | |
| 5813049136 | | | | | | | | | | | | | | 1 | | | | | | | | | | 1 | | | | |
| 5813049438 | | | | | | | | | | | | | | | | | | 7 | 15 | 10 | 16 | 18 | | | | | | |
| 5813049907 | | | | | | | | | | 1 | | | | | | | | | | | | | | | | | | |
| 5813050292 | | | | | | 1 | | | | | | | | | | | | | | | | | | | | | | |
| 5813054835 | 13 | 13 | 19 | 22 | 1 | 1 | 1 | | | | | | | | 2 | | | | | | | | | | | | | |
| 5813055164 | | | | | | 1 | | | | | | | | | | | | | | | | | | | | | | |
| 5813055734 | | | | | | | | | | | | | | | | | | | | 4 | | | | | | | | |
| 5813055748 | | | | | | | | | | | | | | | | | | | | 2 | | | | | | | | |
| 5813055759 | | | | | | | | | | | 2 | 3 | | | 1 | | | | | | | | | | 2 | | | |
| 5813055768 | | | | | | | | | 2 | | | | | | | | | | | | | | | | | | | |
| 5813055790 | | | | | | | | | | 2 | 1 | 2 | | | | | | 3 | | 1 | | | 2 | | 1 | | | |
| 5813055800 | | | | | | | | | | | | 1 | | | | | | | | | | | | | | | | |
| 5813055813 | | | | | | | | | | | | 1 | | | | | | | | | | | | | | | | |
| 5813055817 | | | | | | | | | 8 | 6 | 6 | 2 | 3 | 1 | | | | 1 | | | | | | 1 | 1 | 1 | 2 | |
| 5813055822 | | | | | | | | | | | | | | | | | | | 1 | 1 | 1 | | | | | | | |
| 5813055853 | | | | | | | | | | 1 | | | | | | | | | | | | | | | 1 | | | |
| 5813055862 | | | | | | | | | | | | | | | | | | | | | | | | | 1 | | 1 | |
| 5813055949 | | | | | | | | | | | | 1 | | | | | | | | | | | | | | 1 | | |
| 5813056020 | | | | | | | | | | | | | | | 2 | | | | | | | | | | | | | |
| 5813056026 | | | | | | | | | | 1 | | | | | | | | | | | | | | | | | | |
| 5813056028 | | | | | | | | | | | | | | | 1 | | | | | | | | | | | | | |
| 5813056062 | | | | | | | | | 1 | | 3 | 2 | | | | | | | 1 | 1 | | 3 | | | | | | |
| 5813056072 | | | | | | | | | | | | | | | | 7 | 11 | | | | | | | | | | | |
| 5813056423 | | | | | | | | | | | | | | | 2 | | | | | | | | | | | | | |
| 5813056759 | | | | | 1 | | | | | | | | | | | | | | | | | | | | | | | |
| 5813056887 | | | | | | | | | 55 | 13 | 5 | 1 | 7 | 7 | 3 | | | | | | | | | 5 | 1 | | | |
| 5813056917 | | | | | | | | | | | 2 | | 1 | | | | | | | | | | 2 | | | | | |
| 5813056920 | | | | | | | | | | | | | 1 | | | | | | | | | | | | | | | |
| 5813057148 | | | | | | | | | | | 1 | | | | | | | | | | | | | | | | | |
| 5813057153 | | | | | | | | | | | | | | | 1 | 1 | | | | | | | 1 | 1 | | | | |
| 5813057319 | | | | | 2 | | | | | | | | | | | | | | | | | | | | | | | |

| | | | | | | | | | | | | | | | | | | | | | | | | | | | | |
|---|---|---|---|---|---|---|---|---|---|---|---|---|---|---|---|---|---|---|---|---|---|---|---|---|---|---|---|---|
| 5813057606 | | | | | 1 | 1 | 3 | | | | | | | | | | | | | | | | | | | | | |
| 5813057615 | | | | | | | | | | | | | | 2 | 6 | | | | | | | | | | | | | |
| 5813057933 | | | | | | | | | | | | 1 | | | | | | | | | | | | | | | | |
| 5813057973 | | | | | | | | | | | | | | | | | | | | | | | | | | | 1 | |
| 5813058028 | | | | | | | | | | | | | | | 1 | 4 | 3 | | | | | | | | | | | |
| 5813058097 | | | | | | | | | | | | 2 | | | | | | | | | | | | | | | | |
| 5813058236 | | | | | | | | | | | | | | 1 | | | | | | | | | | | | 1 | | |
| 5813058431 | | | | | | 7 | 1 | 7 | | | | | | 40 | 46 | 4 | 7 | | | | | | | | | | | |
| 5813060768 | | | | | | | | | | | | | | | | | | | | | | | 1 | | 1 | | | |
| 5813060781 | | | | | | | | | | | | | | | 4 | | | | | | | | | | | | | |
| 5813061110 | | | | | | | | | | | | | | | | | 1 | | | | | | | | 1 | | | |
| 5813061112 | | | | | | | | | | | | | | | 1 | | | | | | | | | | | | | |
| 5813061156 | | | | | | | 1 | | | | | | | | | | | | | | | | | | | | | |
| 5813061260 | | | | | | | | | | | | | | | | | | | | | | | 1 | | | | | |
| 5813061262 | | | | | | | | | | | | | | | | | | | | | | | | | | | | 1 |
| 5813061344 | | | | | | 1 | | | | | | | | | | | | | | | | | | | | | | |
| 5813061690 | | | | | | | | | | | | | | | | | | | | | | | | | 4 | | | |
| 5813062772 | | | | | | | | | | | | | | | | | | | | | | | | | 1 | | | |
| 5813063151 | | | | | | | | | | | | | 1 | | | | | | | | | | | | | | | |
| 5813063227 | | | | | 2 | 1 | | 9 | | | | | | | | | | | | | | | | | | | | |
| 5813063239 | | | | | | | | | | | | | | | | | | | | | | | | | 13 | 3 | 6 | |
| 5813063356 | | | | | | 1 | | 4 | | | | | | | | | | | | | | | | | | | | |
| 5813063907 | | | | | | | | | | | | | | | | | | | | | | | | 2 | | | | |
| 5813064332 | | | | | | | | | | | | | | | | | | | | | | | | | 1 | 1 | | |
| 5813064410 | | | | | | | | | | | | | | | | | | | | | | | | | 1 | 1 | | |
| 5813064789 | | | | | | | | | 6 | 3 | | | | | | | | | | | | | | | | | | |
| 5813067337 | | | | | | | | | 1 | 17 | 12 | | | | | | | | | | | | | | | | | |
| 5813067452 | | | | | | | | | 16 | 4 | 8 | 1 | | | | | | | | | | | 1 | 3 | | | | |
| 5813067463 | | | | | | | | | 8 | 3 | | | | | | | | | | | | | | | | | | |
| 5813067666 | | | | | | | | | | | | | | | | | | | 2 | 2 | 4 | 1 | | | | | | |
| 5813067776 | | | | | | | | | | | | | | | 1 | | | | | | | | | | | | | |
| 5813068563 | | | | | | | | | | | | | 1 | | | | | | | | | | | | | | | |
| 5813068707 | | | | | | | | | 1 | | | | | | | | | | | | | | | | | | | |
| 5813068976 | | | | | 1 | 4 | | 1 | | | | | | | | | | | | | | | | | | | | |
| 5813069193 | | | | | 1 | 1 | | 2 | | | | | | | 2 | | | | | | | | | | | | | |
| 5813069255 | | | 1 | | | | 6 | 1 | | | | | | | 1 | | | | | | | | | | | | | |
| 5813069630 | | | | | | | | | 2 | 1 | 1 | | | | | | | | | | | | | | | | 1 | |
| 5813069648 | | | | | 1 | 1 | 2 | 1 | | | | 31 | 21 | | 37 | 1 | 3 | 10 | 9 | 14 | 10 | 13 | 1 | | | | 1 | |
| 5813069844 | 3 | 5 | 3 | 3 | 1 | | 1 | | | | | | | | | | | | | | | | | | | | | |
| 5813069859 | | | | | | | | | 1 | | | 16 | 12 | | | 3 | 1 | | | | | | | 1 | 1 | | 4 | |
| 5813071068 | | | | | | | | | | | | | | | | | | | | | | | | | | | | 1 |
| 5813071078 | | | | | | | | | | | | | 1 | | | | | | | | | | | | | | | |
| 5813071288 | | | | | | | | | | | 1 | | | | | | | | 1 | | | | | | | | | |
| 5813071319 | | | 1 | | | | | | | 1 | | 4 | 5 | 11 | 4 | | | 3 | 1 | 1 | 3 | 2 | 5 | | 2 | 3 | 4 | |
| 5813071348 | | | | | | | | | | | | | | | | 2 | 9 | | | | | | | | | | | |
| 5813075020 | | | | | | | | | | | | | | | | | 2 | | | | | | | | | | | |
| 5813075715 | | | | | 5 | 2 | 7 | 7 | | | | | | 1 | 3 | | | | | | | | | | | | | |
| 5813075931 | | | | | | | | | | | | | | | | | | | | | | | | | | | 1 | |
| 5813076017 | | | | | | | | | | | | | | | | | | | | | | | 1 | | | | | |
| 5813076981 | 1 | 2 | 1 | 1 | 14 | 5 | 12 | 13 | | | | | | | | | | | | | | | | | | | | |
| 5813077488 | | | | | | | | | | | | | | | | | | | | | | | | | | | | 1 |
| 5813077562 | | | | | | | | | 21 | 31 | 33 | 4 | 3 | 1 | | | 1 | | 7 | 2 | 2 | 1 | 1 | 2 | | | | |
| 5813077567 | | | | | | | | | | 1 | | | | | | | | | | | | | | | | | | |
| 5813077788 | | | | | | | | | | | | | | | | | | | | | | | | | | 1 | | |
| 5813078542 | | | | | | | | | | 1 | | | | | | | | | | | | | | | | | | |
| 5813078557 | | | | | | | | | | | | | | 6 | 14 | 2 | 5 | | | | | | | | | | | |
| 5813078563 | | | | | | | | | | | | | | | | 1 | | | | | | | | | | | | |
| 5813078595 | | | | | | | | | | | | | | 1 | 2 | | | | | | | | | | | | | |
| 5813078603 | | | | | | | | | | | | | | 3 | 2 | | | | | | | | | | | | | |
| 5813078743 | | | | | | | | | | | | | | | | | | | | | | | | | 1 | | | |
| 5813082948 | | | | | | | | | | | | | | | | | | | | | | | 2 | | | | | |
| 5813083733 | | | | | | | | | 1 | | | | | | | | | | | | | | | | | | | |
| 5813087531 | | | | | | | | | | | | | | | 1 | | | 4 | 1 | 2 | 1 | 2 | 1 | | 1 | 14 | 16 | |
| 5813087551 | | | | | | | | | | | | | | | | 2 | 1 | | | 1 | | | | | | | | |
| 5813087839 | | | | | | | | | | | | | | | 1 | | | | | | | | | | | | | |
| 5813088391 | | | | | | | | | | 1 | | | | | | | | | | | | | | | | | | |
| 5813088525 | | | | | | | | | | | | | 1 | | | | | | | | | | | | | | | |
| 5813089132 | | | | | | | | | | | | | | | | | | 2 | 4 | 4 | 2 | 3 | | | | | | |
| 5813090530 | | | | | | | | | | | | | | | | 1 | | | | | | | | | | | | |
| 5813091698 | | | | | | | | | | 1 | | | 1 | | | | | | | 1 | | | 1 | | 2 | 1 | | |
| 5813092319 | | | | | | | | | 2 | | 2 | 5 | 4 | | | | | | | | | | | | | 3 | 3 | |
| 5813094916 | | | | | | | | | | | | | | | | | | | 1 | | 1 | | | | | | 3 | |
| 5813095998 | | | | | | | | | | | | | | | | | | | | | 1 | | | | | | | |
| 5813097458 | 1 | 2 | 1 | 2 | 7 | 2 | 5 | 7 | | | | | | | | | | | | | | | | | | | | |
| 5813097632 | | | | | | | | | | 1 | | 2 | | | | | | | | | | | | | | | | |
| 5813097957 | | | | | | | | | | | | | | 2 | 1 | | | | | | 1 | | | | | | | |
| 5813098374 | | | | | | | | | | | | | | | | | | | | | | | 1 | | | | | |
| 5813105726 | | | | | | | | | | | | 3 | | | | | | | | | | | | | | | | |
| 5813108084 | | | | | | | | | | | | | | | | 2 | 4 | | | | | | | | | | | |
| 5813109120 | | | | | | | | | | | | | | 2 | | | | | | | | | | | | | | |

| | | | | | | | | | | | | | | | | | | | | | | | | | | | | |
|---|---|---|---|---|---|---|---|---|---|---|---|---|---|---|---|---|---|---|---|---|---|---|---|---|---|---|---|---|
| 5813111989 | | | | | | | | | | | | | | | | | | | | | | | | | | | | 3 |
| 5813128431 | | | | | | | | | | | | | | | | | | | | | | | 2 | | | | | |
| 5813129572 | | | 2 | | | | | | | 2 | | 1 | 1 | | | | | 1 | | | | | | | | 1 | | |
| 5813129573 | | | | | | | | | 1 | | | | | | | | | | | | | | | 1 | | | | |
| 5813134255 | | | | | | | | | | | | | | | 2 | | | | | | | | | | | | | |
| 5813134294 | | | | | | | | | | | | | | | | | | | | | | | | | | | | 1 |
| 5901118375 | | | | | | | | | | | | | | | | | | | | | | | | | | | 1 | |
| 5901118414 | | | | | | | | 5 | | | | | | | | | | | | | | | | | | | | |
| 5901193006 | | | | | | | | | | | | | | | | | | | | | 1 | | | | | | | |
| 5901193416 | | | | | | | | | | | | | | | | | | | | | | | | | | 1 | | |
| 5901193482 | | | | | | | | | | | | | | | | | | | | | | | 1 | | | | | |
| 5901194250 | | | | 1 | | | | | | | | | | | | | | | 1 | 1 | 3 | | | | | | | |
| 5901194556 | | | | | | | | | | | | | | | | 1 | | | | | | | | 1 | | | | |
| 5901195361 | | | | | | | | | | 1 | | | | | | | | | | | | | | | | | | |
| 5901195498 | | | | | | | | | | 1 | | | | | | | | | | | | | | | | | | |
| 5901195772 | | | | | | | | | | | | | | | | | | | | | | | 1 | | | | | |
| 5901197274 | | | | | | | | | | | | 1 | | | | | | | | | | | | | | | | |
| 5901197793 | | | | | | | | | | | | 2 | | 3 | | | | | | | | | | | | | | |
| 5901201383 | | | | | | | | | | 1 | | | 1 | | | | | | | | | | | | | | | |
| 5901203588 | | | 1 | | | | | | | | | | | | | | | | | | | | | | | | | |
| 5901203987 | | | | | | | | | 1 | | 3 | | | | | | | | | | | | | | | | | |
| 5901204788 | | | | | | | | | | | | | | | | | | | | | | | 2 | 5 | | | | |
| 5901207157 | | | | | | | | | 2 | 1 | 1 | | | | | | | | | | | | | | | | | |
| 5901207822 | | | | | | | | | | | | | | | | | | | | | | | 1 | | | | | |
| 5901211896 | | | 4 | | | | | | | | | | | | | 4 | 5 | | | | | | | | | | | |
| 5901213790 | | | | | | | | | 1 | | | | | | | | | | | | | | | | | | | |
| 5901221083 | | | | | | | 2 | 1 | | | | | | | | | | | | | | | | | | | | |
| 5901221199 | | | | | | | | | | | | | | | | 1 | | | | | | | | | | | | |
| 5901221363 | | | | | | | | | | | 1 | | | | | | | | | | | | 1 | | | | | |
| 5901221890 | | | | | | | | | 3 | | | | | | | | | | | | | | | | | | | |
| 5901222080 | | | | | | | | 1 | | | | | | | | | | | | | | | | | | | | |
| 5901222742 | 2 | 1 | | | 15 | 2 | 14 | 12 | | | | | | 1 | 18 | | | | | | | | | | | | | |
| 5901227104 | | | | | | | | | | | | | | 2 | 1 | | | | | | | | | | | | | |
| 5901232053 | | | | | | | | | | | | 5 | 2 | | | | | | | | | | | | 14 | 12 | 16 | 8 |
| 6400000773 | | | | | | | | | | | | | | | | | | | | | | | | | 3 | | | |
| 7112612987 | | | | | | | | | | | | | | | | 1 | | | | | | | | | | | | |
| 7112613012 | | | | | | | | | | | | | | | | | | | | 1 | | | | | | | | |
| 7112616299 | | | | | 1 | 1 | | | | | | | | 1 | | | | | | | | | | | | | | |
| 7112616416 | | | | | | | | | | 1 | | | | | | | | | 1 | | 1 | | | | | | 2 | |
| 7112616960 | | | | | | | | | | | | 1 | | 4 | | | 1 | | | | | | 1 | | | | | |
| 7112617046 | | | | | | | | | | | | | | 3 | | | | | | | | | | | | | | |
| 7112622017 | | | | | | | | | | | | | | | | | | | 1 | | | | | | | | | |
| 7112626023 | | | | | | | | | | | | | | | | | | | | | | | | | 1 | | | |
