## Supplementary Table 2 for "Connectomic Analysis of the *Drosophila* Lateral Neuron Clock Cells Reveals the Synaptic Basis of Functional Pacemaker Classes"

**Table S2. Outputs of clock neurons**

| | sLNv-1 | sLNv-2 | sLNv3 | sLNv-4 | l-LNv-1 | l-LNV-2 | l-LNv-3 | l-LNv-4 | LNd-1 | LNd-2 | LNd-3 | LNd-4 | LNd-5 | LNd-6 | 5th sLNV | DN1a-1 | DN1a-2 | DN1pA-1 | DN1pA-2 | DN1pA-3 | DN1pA-4 | DN1pA-5 | DN1pB-1 | DN1pB-2 | LPN-1 | LPN-2 | LPN-3 | LPN-4 |
|---|---|---|---|---|---|---|---|---|---|---|---|---|---|---|---|---|---|---|---|---|---|---|---|---|---|---|---|---|
| bodyId_pre | 2.069E+09 | 1.665E+09 | 2.007E+09 | 1.975E+09 | 1.885E+09 | 2.066E+09 | 5.813E+09 | 5.813E+09 | 296544364 | 448260940 | 5.813E+09 | 5.813E+09 | 5.813E+09 | 5.813E+09 | 511051477 | 264083994 | 5.813E+09 | 5.813E+09 | 324846570 | 325529237 | 387944118 | 387166379 | 386834269 | 5.813E+09 | 356818551 | 450034902 | 480029788 | 546977514 |
| 203253253 | | | | | | | | | | | | | 1 | | | | | | | | | | | | | | | |
| 203598647 | | | | | | | | | | | | | | 1 | | | | | | | | | | | | | | |
| 233105330 | | | | | | | | | 1 | | | | | | | | 1 | | | | | | 2 | | | | | |
| 234630133 | | | | | | | | | | | | 10 | 3 | | | | | | | | | | | | | | | |
| 234965803 | | | | | | | | | | | | | | | | | | | | 1 | | 2 | | | | | | |
| 264083994 | | | | | | | | | | | | | | 1 | 3 | | 12 | | | | | | | | 1 | | | |
| 264438143 | | | | | | | | | | | | | | | | | | | | | | | 2 | | | | | |
| 264822904 | | | | | | | | | | 1 | | | | | | | | | | | | | 1 | | | | | |
| 265120424 | | | | | | | | | | | | | | | | | | | | | | | 6 | 2 | | | | |
| 265120467 | | | | | | | | | | | | | | | | | | | | | | | | 2 | | | | |
| 265124786 | | | | | | | | | | | | | | | | | | | | | | | | 1 | | | | |
| 265155147 | | | | | | | | | | | | | | | | | | | | | | | 1 | | | | | |
| 266187342 | | | | | | | | | 1 | | | | | | | | | | | | | | | | 1 | | | |
| 266187383 | | | | | | | | | 5 | 3 | 6 | | 1 | | | | | | | | | | | | 1 | | | |
| 266187477 | | | | | | | | | 2 | 3 | 2 | | | | | | | | | | | | | | | | | |
| 266187480 | | | | | | | | | 6 | 1 | 1 | | | | | | | | | | | | | | | | | |
| 266187532 | | | | | | | | | 1 | | | | | | | | | | | | | | | | | | | |
| 266187559 | | | | | | | | | | | | | | | | | | | | | | | | | 3 | | | |
| 266191395 | | | | | | | | | | | | | | | | | | | | | | | | 2 | | | | |
| 266191407 | | | | | | | | | 2 | | 2 | | | | | | | | | | | | | | | | | |
| 266195641 | | | | | | | | | 1 | | | | | | | | | | | | | | | | | | | |
| 266200011 | | | | | | | | | | | | | | | | | | | | | | | | 1 | | | | |
| 266527996 | | | | | | | | | | 1 | | | | | | | | | | | | | | | | | | |
| 266528044 | | | | | | | | | | 1 | | | | | | | | | | | | | | | | | | |
| 266528078 | | | | | | | | | 2 | 2 | 1 | | | | | | | | | | | | | | | | | |
| 266528086 | | | | | | | | | 1 | 1 | | | | | | | | | | | | | | | | | | |
| 266847236 | | | | | | | | | | | | | | | | | | | | | | | | 1 | | | | |
| 266873389 | | | | | | | | | | | | | | 3 | | | | | | | | | | | | | | |
| 266873591 | | | | | | | | | 9 | 6 | 14 | | | | | | | | | | 1 | | | | 1 | | | |
| 267214250 | | | | | | | | | 15 | 8 | 12 | | | | | | | | | | | | | | | | | |
| 267551614 | | | | | | | | | 1 | | | | | | | | | | | | | | | | | | | |
| 267551639 | | | | | | | | | | 1 | | | | | | | | | | | | | | | | | | |
| 267896360 | | | | | | | | | 2 | | 1 | | | | 1 | | | | | | | | | | 3 | 1 | 1 | |
| 268731005 | | | | | | | | | | | | | | | | | | | | | | | | | | | | 10 |
| 294424196 | | | | | | | | | | 1 | | | | | | | | | | | | | | | | | | |
| 294428266 | | 2 | | | | | | | | | | | 1 | | | | | | | | | | | 3 | | | | |
| 294432626 | | | | | | | | | | | | | | | | | | | | | | | | | 1 | 1 | 1 | |
| 294436967 | | | | | | | | | | | | | | | | 1 | 1 | | | | | | | | | | | |
| 294437347 | | | | | | | | | | | | | | 1 | | | | | | 1 | | | | | | | | |
| 294445804 | | | | | | | | | | | | | | | | | 1 | | | | | | | | | | | |
| 294760699 | | | | | | | | | | | | | 2 | | | | | | 1 | | | | 1 | | | | | |
| 294783216 | 19 | 24 | 12 | 17 | | | | | | | | | | | | | 2 | | | | | | | | | | | |
| 294791642 | | | | | | | | | | | | | | | | | | | | | | | | 1 | | | | |
| 294795822 | | | | | | | | | | | | | | | | | | | | | | | 6 | 6 | | | | |
| 295063181 | 1 | 1 | | 5 | | | | | | | | | 2 | 9 | 25 | | | | | | | 1 | | | | | | |
| 295103172 | | | | | | | | | | | | | 1 | | | | | | | | | | | | | | | |
| 295132077 | | | | | | | | | | | | | | | | | | | | | | | 1 | 4 | | | | |
| 295132551 | | | | | | | | | | | | | | | | | | | | | | | | 3 | | | | |
| 295132845 | | | | | | | | | | | | | | | | | | | | | | | | 1 | | | | |
| 295133015 | | | | | | | | | | | | | | | | | | | | | | | | 3 | | | | |
| 295133743 | | | | | | | | | | | | | | | | | | | | | | | | 1 | | | | |
| 295133778 | | | | | | | | | | | | | | | 1 | | | 2 | 3 | 2 | 1 | | 16 | 12 | | | | |
| 295133927 | | | | | | | | | | | | | | | | | | | | | | | | 1 | | | | |
| 295136450 | | | | | | | | | | | | | | | | | | | | | | | | 1 | | | | |
| 295136668 | | | | | | | | | | 1 | | | | | | | | | | | | | | | | | | |
| 295137392 | | | | | | | | | | | | | | | | | | | | | | | 4 | 3 | | | | |
| 295137515 | | | | | | | | | | | | | | | | | | | | | | | | 1 | | | | |
| 295443724 | | | | | | | | | | | | 1 | | | | | | | | | | | | | 1 | | | |
| 295477467 | | | | | | | | | | | | | | | | | | | | | | | 4 | 3 | | | | |
| 295477903 | | | | | | | | | | | | | | | | | | | | | | | | | | 2 | | |
| 295478389 | | | | | | | | | | | | | | | | | | | | | | | 2 | | | | | |
| 295793991 | 1 | | 3 | | | | | | | 1 | | | | | | | | | | | | | 5 | 3 | | 1 | | |
| 295802733 | | | | | | | | | | 1 | | | | | | | | | | 1 | | | 2 | | 1 | | | |
| 295814411 | | | | | | | | | | | | | | | | | 1 | | | | | | | | | | | |
| 295814630 | | | | | | | | | | 1 | | | | | | | | | | | | | | | | | | |
| 295814932 | | | | | | | | | | | 1 | | | | | | | | | | | | | | | | | |
| 295815100 | | | | | | | | | | | | | | | | | | | | | | | | | 1 | | | |
| 295815955 | | | | | | | | | | | | | | | | | | | | | | | | | 1 | | | |
| 295819998 | | | | | | | | | | | | | | | | | | | | | | | 8 | 1 | | | | |
| 295820069 | | | | | | | | | | | | | | | 2 | | | | | | | | | | | | | |
| 295849217 | | | | | | | | | | 1 | | | 1 | 1 | 5 | | | 4 | | 2 | | 2 | 33 | 12 | | | | |
| 295849410 | | | | | | | | | | | | | | | | | | | | | | | | 1 | | | 1 | |

| | | | | | | | | | | | | | | | | | | | | | | | | | | | | |
|---|---|---|---|---|---|---|---|---|---|---|---|---|---|---|---|---|---|---|---|---|---|---|---|---|---|---|---|---|
| 296120593 | | | | | | | | | | | | | | | | | | | | | | | | 1 | | | | |
| 296139882 | | | | | | | | | | 1 | | | | | | | | | | | | | 1 | | 3 | 5 | 8 | |
| 296153150 | | | | | | | | | | | | | | | | | | | | | | | | | 1 | | | |
| 296168382 | | | | | | | | | | | | | | | | | | | | | | | 2 | | | | | |
| 296194535 | | | | | | | | | 2 | | | | | | | | | | | | | | | | | | | |
| 296199026 | | | | | | | | | 7 | 6 | 23 | | | | | | | 1 | | 1 | | | | | | 1 | | |
| 296203440 | | | | | | | | | | | | 1 | | | | | | | | | | | | 1 | | | | |
| 296358430 | | | | | | | | | | | | 2 | 4 | | | | | | | | | | | | | | | 1 |
| 296502002 | | | | | | | | | | | 1 | | | | | | | | | | | | | | | | | |
| 296505463 | | | | | | | | | | | | | | | | | | | | | | | | | 2 | | | |
| 296509709 | 4 | 1 | 1 | | | | | | | | 1 | | | | 3 | | | | | | | | 6 | 4 | | | | |
| 296535619 | | | | | | | | | | | | | | | | | | | | | | | 1 | | | | | |
| 296535868 | | | | | | | | | | 1 | | | | | | | | | | | | | | | | | | |
| 296543992 | | | | | | | | | | 1 | | | | | | | | | | | | | 3 | 2 | | | | |
| 296544262 | | | 1 | | | | | | | | | | | 1 | 4 | | | | | | | | 2 | | | | | |
| 296544364 | | | | | | | | | | 3 | 6 | | | | | 1 | | | | | | | | | | | | |
| 296829531 | | | | | | | | | 1 | | | | | | | | | | | | | | | | | | | |
| 296838079 | | | | | | | | | | | | | | | | | | | | | | | 1 | | | | | |
| 296838229 | | | | | | | | | | | | | | | | | | | | | | | | 1 | | | | |
| 296841676 | | | | | | | | | | | | | | | | | | | | | | | | | 1 | | | |
| 296846314 | | | | | | | | | | 1 | | | | | | | | | | | | | | | | | | |
| 296850634 | | | | | | | | | | | | | | | | | | | | | | | 1 | | | | | |
| 296851132 | | | | | | | | | | | | | | | | | | | | | | | 3 | 6 | | | | |
| 296851235 | | | | | | | | | | | | | | | | | | | | | | | 1 | | | | | |
| 296851261 | | | | | | | | | | | | | | | | | | | | | | | 1 | | | | | |
| 296859399 | | | | | | | | | | | | | | | | | | | | | | | 1 | | | | | |
| 296859438 | | | | | | | | | 1 | | | 1 | | | | | | | 1 | | | | 3 | 2 | 1 | | | |
| 296863817 | | | | | | | | | | | | | | | | | | | | | | | | 1 | | | | |
| 296868077 | | | | | | | | | 2 | | 1 | | | | | | | | | | | | 1 | | | | | |
| 296885362 | | | | | | | | | 13 | 2 | 2 | | | | | | | | | | | | 2 | 2 | 2 | 1 | 1 | |
| 297166197 | | | | | | | | | | | | | | | | | | | | | | | | | 1 | | | |
| 297221971 | | | | | | | | | 1 | 2 | | | | 3 | 3 | | | | | | | | 1 | 1 | | 2 | | |
| 297230359 | | | | | | | | | 1 | | | | | | | | | | | | | | | | | | | |
| 297230760 | | | | | | | | | 10 | 12 | 8 | | | | | | | | | | | | | | 3 | 1 | 1 | |
| 297239396 | | | | | | | | | | 4 | 4 | | | | | | | | | | | | | | | | | |
| 297243542 | | | | | | | | | 49 | 12 | 14 | 1 | | 4 | 5 | | | | | | | | | | | | 2 | |
| 297247801 | | | | | | | | | | | | | | 1 | | | | 1 | | | | | | | | | | |
| 297251714 | | | | | | | | | | | | 2 | | 1 | | | | | | | | | | | | 9 | 14 | |
| 297251765 | | | | | | | | | 6 | 7 | 9 | | | | | | | | | | | | | | | | | |
| 297502392 | | | | | | | | | | | | | | | | | | | | | | | | | 2 | | | |
| 297519807 | | | | | | | | | | 2 | | | | | | | | | | | | | | | | | | |
| 297528741 | | | | | | | | | 1 | 2 | | | | | | | | | | | | | | | | | | |
| 297533212 | | | | | | | | | | | | | | | | | | | | | | | 1 | | | | | |
| 297541369 | 1 | 9 | 7 | 3 | | | | | | | | | | | | 1 | | | | | | | 1 | 5 | | 1 | | |
| 297558862 | | | | | | | | | | | | | | | | | | | | | | | | 2 | | | | |
| 297563139 | | | | | | | | | 3 | 3 | 3 | 5 | 4 | | | | | | | | | | | | | 1 | 2 | |
| 297580512 | | | | | | | | | 10 | 2 | 2 | | | | | | | | | | | | | | | | | |
| 297580589 | | | | | | | | | | | | | | | | | | | | | | | | | 1 | | | |
| 297584752 | | | | | | | | | | 1 | | | | 2 | 4 | | | | 2 | 2 | | 2 | 9 | 8 | | | | |
| 297584869 | | | | | | | | | 4 | 7 | 4 | | | | | | | | | | | | | | | | | |
| 297588775 | | | | | | | | | 4 | 3 | 5 | | | | | | | | | | | | | | | | | |
| 297588821 | | | | | | | | | 2 | 2 | | | | | | | | | | | | | | | | | | |
| 297588838 | | | | | | | | | 4 | 2 | 5 | | | | | | | | | | | | | | | | | |
| 297588882 | | | | | | | | | 4 | 17 | 12 | | | | | | | | | | | | | | | | | |
| 297588995 | | | | | | | | | 3 | 2 | | | | | | | | | | | | | 2 | | | | | |
| 297593055 | | | | | | | | | 1 | 9 | 1 | | | | 1 | | | | | | | | | | | | | |
| 297899921 | | | | | | | | | 1 | | | | | | | | | | | | | | | | | | | |
| 297908595 | | | | | | | | | | | | | | | | | | | | | | | | 1 | | | | |
| 297908801 | | | | | | | | | | 2 | | | | | | | | | | | | | | | | | | |
| 297912117 | | | | | | | | | | | 1 | | | | | | | | | | | | | | | | | |
| 297917158 | | | | | | | | | | 3 | 5 | | | 1 | 1 | | | | | | | | | | | | | |
| 297917363 | | | | | | | | | | | | | 2 | | 1 | | | | | | | | | | 68 | 3 | 9 | |
| 297917474 | | | | | | | | | | | | | | | | | | | | | | | | | 1 | | | |
| 297917475 | | | | | | | | | | 1 | 2 | | | 1 | | | | | | | | | | 1 | | | | |
| 297921570 | | | | | | | | | | | | | | | | | | | 1 | | | | 3 | | | | 1 | |
| 297921608 | | | | | | | | | | | | | | | | | | | | | | | 1 | 1 | | | | |
| 297921642 | | | | | | | | | | | | | 2 | | | | | | | | | | | | | | | |
| 297921802 | | | | | | | | | | | 2 | | | | | | | | | | | | | | | | | |
| 297925478 | | | 1 | | | | | | 2 | 5 | 14 | | | 2 | | 1 | 1 | | | | | | | | | | | |
| 297925596 | | | | | | | | | | 2 | 1 | | | | | | | | | | | | | | | | | |
| 297925608 | | | | | | | | | | 1 | | | 1 | | | | | | | | | | | | 83 | 11 | 5 | |
| 297925628 | | | | | | | | | | | | | 1 | 2 | | | | | | | | | | | | 1 | | |
| 297925656 | | | | | | | | | 1 | 1 | | | | | | | | | | | | | | | | | | |
| 297925719 | | | | | | | | | 6 | 1 | 2 | | | | | | | | | | | | | | | | | |
| 297925742 | | | | | | | | | | 1 | | | | | | | | | | | | | | | | | | |
| 297925887 | | | | | | | | | | 7 | 12 | | | | | | | | | | | | | | | | | |
| 297929768 | | | | | | | | | 1 | | | | | | 4 | | | | | | | | 2 | | | | 1 | |
| 297929807 | | | | | | | | | 3 | | 2 | | | | | | | | | | | | | | | | | |
| 297929902 | | | | | | | | | 7 | | 1 | | | | | | | | | | | | | | | | | |
| 297930040 | | | | | | | | | 2 | 2 | 10 | | | | | | | | | | | | | | | | | |

| | | | | | | | | | | | | | | | | | | | | | | | | | | | | |
|---|---|---|---|---|---|---|---|---|---|---|---|---|---|---|---|---|---|---|---|---|---|---|---|---|---|---|---|---|
| 297930119 | | | | | | | | | | 1 | 1 | | | | | | | | | | | | | | | | | |
| 297934028 | | | | | | | | | 1 | 1 | | | | | | | | | | | | | | | | | | |
| 297934049 | | | | | | | | | 3 | 3 | 1 | | | | | | | | | | | | | | | | | |
| 297934349 | | | | | | | | | | 2 | 3 | | | | | | | | | | | | | | | | | |
| 297938303 | | | | | | | | | 3 | | | | | | | | | | | | | | | | | | | |
| 297938585 | | | | | | | | | 2 | | 1 | | | | 1 | | | | 1 | | 1 | 2 | | 2 | | 13 | 5 | |
| 297947227 | | | | | | | | | 2 | 1 | 2 | 1 | | | | | | 1 | | | | 1 | | | | 13 | 13 | |
| 298055794 | | | | | | | | | | | | | 1 | | | | | | | | | 1 | | | | | | 1 |
| 298197305 | | | | | | | | | 2 | | | | | | | | | | | | | | | | | | | |
| 298201943 | | | | | | | | | 2 | | | | | | | | | | | | | | | | | | | |
| 298214577 | | | | | | | | | | | | | | | | | | | | | | | | | 1 | | | |
| 298250090 | | | | | | | | | | | | | | | | | | | | | | | | | 1 | | | |
| 298254517 | | | | | | | | | | | | 11 | 15 | | 1 | | | | | | | | | | 1 | 2 | 3 | 9 |
| 298258354 | | | | | | | | | | 3 | 10 | | | | | | | | | | | | | | | 1 | | |
| 298258399 | | | | | | | | | | 1 | 1 | | | | | | | | | | | | | | | | | |
| 298258513 | | | | | | | | | | | | | | | | | | | | | | | | | 1 | | | |
| 298258714 | | | | | | | | | | 1 | 11 | | | | | | | | | | | | | | 2 | | | |
| 298262408 | | | | | | | | | | | | | | | | | | | | | 1 | | | 1 | 2 | 1 | | |
| 298262498 | | | | | | | | | 1 | | | | | | | | | | | | | | | | | | | |
| 298262644 | | | | | | | | | | | | | | | 2 | 1 | | | | 1 | | | 1 | 1 | 22 | 73 | 61 | |
| 298262663 | | | | | | | | | | | 2 | | | | | | | | | | | | | | 5 | | | |
| 298266660 | | | | | | | | | | | | | | 1 | | | | | | | | | | | 2 | 1 | | |
| 298266716 | | | | | | | | | 1 | | 2 | | | | | | | | | | | | | | | | | |
| 298266783 | | | | | | | | | 1 | | | | | | | | | | | | | | | | | | | |
| 298266797 | | | | | | | | | | | 1 | | | | | | | | | | | | | | | | | |
| 298266903 | | | | | | | | | | | 4 | | | | | | | | | | | | | 1 | | | | |
| 298266950 | | | | | | | | | 1 | 19 | 11 | | | | | | | | | | | | | | | | | |
| 298270808 | | | | | | | | | 1 | 2 | 4 | | | | | | | | | | | | | | 1 | | | |
| 298275024 | | | | | | | | | | 1 | | | | | | | | | | | | | | | | | | |
| 298275090 | | | | | | | | | 1 | | | | | | | | | | | | | | | | | | | |
| 298275356 | | | | | | | | | | | | | | | 1 | | | | | | | | | | | 20 | 16 | |
| 298279428 | | | | | | | | | 1 | | | | | | | | | | | | | | | | | | | |
| 298279629 | | | | | | | | | 17 | 18 | 6 | | | | | | | | | | | | | 1 | | | | |
| 298288301 | | | | | | | | | 4 | 2 | 2 | | | | | | | | | | | | | | 1 | | | |
| 298542439 | | | | | | | | | 1 | | | | | | | | | | | | | | | | | | | |
| 298586553 | | | | | | | | | | 1 | | | | | | | | | | | | | | | 9 | 1 | | |
| 298590924 | | | | | | | | | | 3 | | | | | | | | | | | | | | | 2 | | | |
| 298591130 | | | | | | | | | 9 | 11 | 17 | 1 | | | | | | | | | | | 1 | | | | | |
| 298599269 | | | | | | | | | 18 | 1 | 1 | | | | | | | | | | | | | | | | | |
| 298599310 | | | | | | | | | 1 | | | | | | | | | | | | | | | | | | | |
| 298599672 | | | | | | | | | | | 1 | | | | | | | | | | | | | | | | 2 | |
| 298603633 | | | | | | | | | | 1 | 2 | 1 | | | | | | | | | | | | | | 1 | | |
| 298607679 | | | | | | | | | | | | | | | | | | | | | | | | | 1 | | | |
| 298612190 | | | | | | | | | 4 | 2 | 1 | | | | | | | | | | | | | | | | | |
| 298616588 | | | | | | | | | | 1 | | | | | | | | | | | | | | | 22 | | | |
| 298625276 | | | | | | | | | 10 | 5 | 5 | 1 | | | | | | | | | | | | | | | | |
| 298935769 | | | | | | | | | 4 | | 2 | | | | | | | | | | | | | | | | | |
| 298944620 | | | | | | | | | 2 | 1 | 3 | | 2 | | 1 | | | | | | | | | | | | | |
| 298948785 | | | | | | | | | | | | | | | | | | | | | | | 2 | 3 | | | 1 | |
| 298952944 | | | | | | | | | | | | 3 | 4 | | | | | | | | | | | | | | | |
| 298957166 | | | | | | | | | | | | | | | | | | | | | | | | | | | 1 | |
| 298961895 | | | | | | | | | 5 | 15 | 4 | | | | | | | | | | | | | | | | | |
| 298962041 | | | | | | | | | | | | | | | | | | | | | | | | | | 5 | 1 | |
| 299043307 | | | | | | | | | 1 | 3 | 3 | | | | | | | | | | | | | | | | | |
| 299043443 | | | | | | | | | 1 | 1 | | | 3 | | | | | | | | | | | | | | | 2 |
| 299272434 | | | | | | | | | | | | | | | | 4 | 4 | | | | | | | | | | | |
| 299289811 | | | | | | | | | | | | | | | | | | | | | | | | | 5 | | | |
| 299290105 | | | | | | | | | | | 2 | 2 | 1 | | | | | 5 | 2 | 4 | | 3 | 1 | 2 | 21 | 47 | 52 | |
| 299298593 | | | | | | | | | 1 | 1 | | | | | | | | | | | | | | | | | | |
| 299617529 | | | | | | | | | | | | | | | | | | | | | | | | 1 | 2 | 2 | 1 | |
| 299621755 | | | | | | | | | | | | | | | | | | | | | | | | | | | | 5 |
| 324846570 | | | | | | | | | | | | | | 9 | 8 | | | 1 | | 2 | 1 | 1 | | 1 | | | | |
| 325113795 | 1 | | | | | | | | | | | | | | | | | | | | | | | | | | | |
| 325122109 | | | | | | | | | 1 | | | | | | | 1 | 1 | | | | | | 2 | 5 | | | | |
| 325122525 | 64 | 60 | 71 | 56 | | | | | | | | | | | | | | | | | | | 1 | 2 | | | | |
| 325446181 | | | | | | | | | | 1 | | | | | | | | | | | | | | | | | | |
| 325455002 | 20 | 31 | 23 | 22 | | | | | | | | | | | | | | | | | | | 1 | | | | | |
| 325529237 | | | | | | | | | | | | | | 14 | 12 | | | | | | | 2 | 1 | 1 | | 1 | | |
| 325814461 | | | | | | | | | | | | | | | | 10 | 2 | | | | | | 1 | | | | | |
| 325815440 | | | | | | | | | | | | | | | | | | | | | | | | | 1 | | | |
| 326119769 | | | | | | | | | | | | | | | | 4 | 1 | | | | | | | | | | 1 | |
| 326137566 | | | | | | | | | | | | | | | | | | | | | | | | | 2 | | | |
| 326159897 | | | | | | | | | | | | | | | | | | | | | | | | | 2 | | | |
| 326164634 | | | | | | | | | | | 1 | | | | | | | | | | | | | | | | | |
| 326253554 | | | | | | | | | | | | | | | 1 | | | 1 | 1 | 1 | | 1 | 2 | 1 | 2 | | | |
| 326460989 | | | | | | | | | | | | 2 | 2 | | | | | | | | | | | | | | | |
| 326465302 | | | | | | | | | | | | 1 | | | | | | | | | | | 1 | 1 | | | | |
| 326508583 | | | | | | | | | | | | | | | | | | | | | | | 1 | | | | | |
| 326509224 | | | | | | | | | | | | | | | | | | | | | | | | | 2 | | | |
| 326530038 | | | | | | | | | | | | 1 | | | | | | | | | | | 2 | 1 | | 7 | 8 | |

| | | | | | | | | | | | | | | | | | | | | | |
|---|---|---|---|---|---|---|---|---|---|---|---|---|---|---|---|---|---|---|---|---|---|
| 326542980 | | | | | | | | | | | | | | | | 4 | | | | | |
| 326556230 | | | | | | | | | | | | | | | | | | | | 1 | |
| 326560782 | | | | | | | | 1 | 2 | | | | | | | | | | | | |
| 326817565 | | | | | | | | | | | | | | | | | | 1 | | | |
| 326875404 | | | | | | | | | | | | | | | | 1 | 1 | | | | |
| 326888433 | | | 10 | 6 | 13 | | | | | | 1 | | | 1 | | | | | 1 | | |
| 326888609 | | | | | | | | | | | | | | | | | | 1 | | | |
| 326888675 | | | 2 | | 2 | | | | | | | | | | | | | | | | |
| 326905710 | | | | | | | | | | | | | | | | | | 1 | | | |
| 326905980 | | | 1 | 1 | 4 | | | | | | | | | | | | | | | | |
| 327225145 | | | | | | | | | | | | | | | | | 1 | | | | |
| 327229641 | | | | | | | | | | | | | | | | | 2 | | | | |
| 327242169 | | | 3 | 8 | 3 | | | | | | | | | | | | | | | | |
| 327242287 | | | | | | | | | | | 1 | 1 | 1 | 2 | 1 | 36 | 53 | | 2 | 2 | |
| 327246381 | | | | | | | 1 | | 1 | | | | | | | | | 1 | | | |
| 327246931 | | | 3 | 1 | 1 | 1 | | 1 | 4 | | | | | | | | | 1 | | | |
| 327251085 | | | | | 1 | | | | | | | | | | | | | | | | |
| 327255181 | | | 3 | 3 | 5 | | | | | | | | | | | | | | | | |
| 327499164 | | | | | | | | | | | | | | | | | | 1 | | | |
| 327532246 | | | 1 | | | | | | | | | | | | | | | | | | |
| 327584307 | | | | 3 | 1 | | | | | | | | | | | | | | | | |
| 327587580 | | | 4 | 8 | 9 | | | | | | | | | | | | | | | | |
| 327587644 | | | 9 | 16 | 27 | | | 2 | 4 | | | | | | | | | | 2 | | |
| 327588225 | | | | | | | 2 | | | | | | | | | | | 31 | 9 | 19 | 4 |
| 327588446 | | | 2 | | | 3 | 6 | 38 | 44 | | 2 | | | 2 | 1 | | 1 | | | 1 | |
| 327592562 | | | | | | | | 2 | 3 | | | | | 1 | | 7 | 11 | | | | |
| 327596481 | | | 1 | 2 | 11 | | | | | | | | | | | | | | | | |
| 327600366 | | | 3 | 2 | | | | | | | | | | | | | | | | | |
| 327843160 | 1 | | | | | 1 | | | 1 | | | | | | | 3 | 1 | | | 1 | |
| 327872592 | | | 1 | 1 | 1 | | | | | | | | | | | | | | | | |
| 327928411 | | | | | | | | | | | | | | | | 1 | 1 | | 3 | | |
| 327932807 | | | 4 | 8 | 11 | | | 1 | | | | | | | | | | | | | |
| 327932921 | | | | 2 | | | | | | | | | | | | | | | | | |
| 327933008 | | | | | 1 | | | | | | | | | | | | | | | 1 | |
| 327933027 | | | | 1 | | | | 2 | 13 | | | | | | | 2 | 3 | | 5 | 6 | |
| 327933061 | | | | | | | | | | | | | | | | | | 1 | | | |
| 327933208 | | | 12 | 8 | 9 | | | | | | | | | | | | | | | | |
| 327933679 | | | 11 | 11 | 13 | | | | | | | | | | | | | | | 1 | |
| 327933814 | | | 3 | 1 | 2 | | 1 | 6 | 10 | | 4 | 1 | 3 | 9 | 1 | 10 | 19 | 1 | | 5 | |
| 327937328 | | | | | 2 | | | | | | | | | | | | | | | | |
| 327937494 | | | 4 | 16 | 18 | | | | | | | 1 | | | | | | | | 1 | |
| 327937506 | | | | | | | | | | | | | | | | | 1 | 1 | | 1 | |
| 327937628 | | | 9 | 1 | 9 | | | | | | | | | | | | | | | | |
| 327941614 | | | 1 | | | | | | | | | | | | | | | | | | |
| 327941844 | | | | 1 | 2 | | | | | | | | | | | | | | | | |
| 327945864 | | | 2 | 2 | 1 | | | | | | | | | | | | | | | | |
| 328010530 | | | | | | | | | 1 | | 4 | 1 | 5 | | 1 | 2 | 2 | | | 1 | 1 |
| 328213824 | | | | | | | | | | | | | | | | 1 | | | | | |
| 328244591 | | | | | | | | | | | | | | | | | | 1 | | | |
| 328257630 | | | | | | | | | | | | | | | | | | 1 | | | |
| 328264983 | | | | | | | | | | | | | 1 | | | 1 | 1 | | | | |
| 328265182 | | | | 2 | 2 | | | | | | | | | | | | | | | | |
| 328265389 | | | | | | 12 | 6 | 2 | 1 | | | | | | | | | 20 | 21 | 11 | 2 |
| 328269766 | | | 7 | 6 | 2 | | | | | | | | | | | | | 1 | | | |
| 328273806 | | | 2 | 10 | 20 | | | | 1 | | | | | | | | | | | | |
| 328274706 | | | 5 | 7 | 6 | | | | | | | | | | | | | | | | |
| 328278125 | | | 2 | 1 | | | | | | | | | | | | | | | | | |
| 328278368 | | | 9 | 4 | 4 | | | | | | | | | | | | 2 | | | | |
| 328278449 | | | | | | 1 | | | | | | | | | | | | 2 | | | |
| 328283406 | | | | 1 | 3 | | | | | | | | | | | | 2 | | | | |
| 328283521 | | | | | | | | | | | | | | | | | 2 | | | | |
| 328283528 | | | | | 3 | | | | | | | | | | | | | | | | |
| 328287175 | | | | 1 | | | | | | | | | | | | | | | | | |
| 328291455 | | | | 1 | | | | | | | | | | | | | | | | | |
| 328373131 | | | | | | 5 | 1 | 1 | | | | | | | | | | | | | |
| 328551180 | | | 1 | | | | | | | | | | | | | | | | | | |
| 328559607 | | | 2 | | | | | | | | | | | | | | | | | | |
| 328611004 | | | 2 | | | | | | | | | | | | | | | | | | |
| 328628560 | | | | | 2 | | | | | | | | | | | | | | | | |
| 328632707 | | | | 1 | | | | 1 | 2 | | | | | | | | 1 | 43 | 23 | 15 | 9 |
| 328709642 | | | 1 | | | | | | | | | | | | | | | | | | |
| 328714074 | | | 2 | 6 | | | | | | | | | | | | | | | | | |
| 328714144 | | | 4 | 1 | | | | | | | | | | | | | | | | | |
| 328718126 | | | | | | 1 | | | | | | | | | | | | | | | |
| 328883562 | | | 2 | | | | | | | | | | | | | | | | | | |
| 328909309 | | 1 | | | | | | | | | | | | | | | | | | | |
| 328935472 | | | | | | | | | | | | | | | | | | 2 | | | |
| 328943204 | | | | | | | | | | | | | | | | | | | | | 5 |
| 328952123 | | | | | | 1 | | | | 1 | | | | | | | | 1 | 3 | 2 | |
| 328955898 | | | | | 1 | | | | | | | | | | | 3 | | 1 | | | |

| | | | | | | | | | | | | | | | | | | | | | | | | | | | | |
|---|---|---|---|---|---|---|---|---|---|---|---|---|---|---|---|---|---|---|---|---|---|---|---|---|---|---|---|---|
| 329050894 | | | | | | | | | | | | | | | 1 | | | | | | | | | | | | | |
| 329206392 | | | | | | | | | | | | | | | | | | | | | | | | 2 | | | | |
| 329254539 | | | | | | | | | 1 | | | | | | | | | | | | | | | | | | | |
| 329301513 | | | | | | | | | | | | 1 | 12 | | 1 | | | | | | | | 5 | | 5 | | 2 | |
| 329301762 | | | | | | | | | | | | | | | | | | | | | | | | | 1 | | | |
| 329310230 | | | | | | | | | | | | 6 | 11 | 1 | | | | | | | | | | | 1 | 5 | 3 | 8 |
| 329395900 | | | | | | | | | 1 | | | | | | | | | | | | | | | | | | | |
| 329396828 | | | | | | | | | 4 | | | 2 | | | | | | | | | | | | | | | | |
| 329566174 | | | | | | | | | | | | | | | | | | | | | | | | 1 | | | | |
| 329566197 | | | | | | | | | | 4 | 9 | | | | | | | | | | | | | | | | | |
| 329732855 | | | | | | | | | | | | 50 | 27 | 62 | 27 | | | 1 | 3 | | 1 | | | | 1 | 2 | | |
| 329738037 | | | | | | | | | 2 | | | | 2 | | | | | | | | | | | | | | | |
| 329897255 | | | | | | | | | | | | | | | | | | | | | | | | | 2 | | | |
| 329966674 | | | | | | | | | | | | | | 2 | | 1 | | | | | | | | | 2 | | | |
| 330074340 | | | | | | | | | | | | | | | | | | | | | | 2 | | | | | | |
| 330074405 | | | | | | | | | 3 | 1 | | | | | | | | | | | | | | | | | | |
| 330074628 | | | | | | | | | 6 | 4 | | | | | | | | | | | | | | | | | | |
| 330079329 | | | | | | | | | | 1 | | | | | | | | | | | | | | | | | | |
| 330079381 | | | | | | | | | 1 | | | 1 | | | | | | | | | | | | | | | | |
| 330259272 | | | | | | | | | | | | | | | | | | | | | | | | | 1 | | | |
| 330260421 | | | | | | | | | | | | 1 | | | | | | | | | | | | | | | | |
| 330316013 | | | | | | | | | | | 10 | 1 | | | | | | | | | | | | | 1 | | 1 | |
| 330319977 | | | | | | | | | | | | | | | | | | | | | | | 1 | 2 | | | | |
| 330415160 | | | | | | | | | 2 | | | | | | | | | | | | | | | | | | | |
| 330415332 | | | | | | | | | | | | 17 | 16 | | | | | | | | | | | | | | | 2 |
| 330415373 | | | | | | | | | 13 | 3 | | 1 | | | | | | 1 | | 1 | | | | | | | | |
| 330640044 | | | | | | | | | | | | | | | | | | | | | | | | | 8 | 1 | 1 | |
| 330661786 | | | | | | | | | | | | | | | | | | | | | | | | | | | | 2 |
| 331006655 | | | | | | | | | | | | 2 | 2 | | | | | | | | | | | | 6 | 1 | 2 | |
| 331097097 | | | | | | | | | 2 | | | | | | | | | | | | | | | | | | | |
| 331338959 | | | | | | | | | 4 | | | 2 | 2 | | | | | | | | | | | | | | | |
| 331343089 | | | | | | | | | | | | | | | | | | | | | | | | | 1 | | | |
| 331774819 | | | | | | | | | | | | 1 | | | | | | | | | | | | | 4 | | | 20 |
| 331774931 | | | | | | | | | | 1 | 3 | | | | | | | | | | | | | | | | | |
| 355453590 | 75 | 61 | 97 | 72 | | | | | | | | | | | | | | | | | | | | | | | | |
| 355816896 | | 1 | | | | | | | | | | 1 | 1 | | | 1 | 1 | | | | | | | | 3 | 2 | 2 | |
| 356136008 | | | | | | | | | | | | 3 | 2 | | 1 | | | | | 1 | | | | | 1 | 1 | 1 | |
| 356140100 | | 1 | 3 | | | | | | | | | 1 | 1 | 25 | 34 | 1 | 3 | | | 1 | | | 1 | 2 | | 1 | | |
| 356153135 | | | | | | | | | | | | | | | | | 4 | | | | | | | | | | | |
| 356429209 | | | | | | | | | | | | | | | | | | | | | | | 1 | | | | | |
| 356472628 | | | | | | | | | | | | | | 1 | | | | | | | | | | 1 | 2 | | | |
| 356472645 | 2 | 1 | | | | | | | 1 | 1 | | 1 | | | | 3 | 2 | | | | | | | 4 | 1 | | | |
| 356472660 | | | | 1 | | | | | | | | | 2 | | | | | | | | | | | 3 | 1 | | | |
| 356499529 | | | | | | | | | | | | | | | | | | | | 1 | | | | | | | | |
| 356503889 | | | | | | | | | | | | | | | | 7 | 5 | | | | | | | | | | | |
| 356809556 | 2 | | | 1 | | | | | | 2 | | | | | 1 | | | | | | | | | | | | | |
| 356818551 | | | | | | | | | | | | | 2 | | 4 | 3 | 3 | 1 | | | | | 8 | 2 | | 2 | 3 | |
| 356823065 | | | | | | | | | | | | | | | | | 1 | | | | | | | | | | | |
| 356835277 | | 1 | | | | | | | | 1 | 1 | | | | 2 | | | | | | | | 1 | | | | | |
| 356913605 | | | | | | | | | | | | | | | | | | | | | | | | | | | 1 | |
| 357150583 | | 1 | | | | | | | | | | 1 | | | | | | | | | | | 1 | 2 | | | | |
| 357159104 | | | | | | | | | | | | | | | | 1 | | | | | | | | | | | | |
| 357224041 | | | | | | | | | | | | | | | | 2 | | | | | | | | | | | | |
| 357482735 | | 1 | | | | | | | | | | 2 | 1 | 1 | 1 | 3 | 2 | | | | | | 13 | 11 | 1 | 3 | 1 | |
| 357490931 | | | | | | | | | | | | | | | | | | | | | | | | | 1 | | | |
| 357515217 | | | | | | | | | | | | | | | | | | | | | | | 2 | 2 | | | | |
| 357522097 | | | | | | | | | | | | | | | | | | | | | | | 6 | 10 | 2 | | | |
| 357569231 | | | | | | | | | | | | | | | | | | | | | | | | 1 | | | | |
| 357569362 | | | | | | | | | | | | | | | | | | | | | | | | 1 | | | | |
| 357573154 | | | | | | | | | | | | | | | | | | | | | | | 2 | | | | | |
| 357581846 | | | | | | | | | | | | 6 | | | | | | | | | | | | | | | | |
| 357581860 | | | | | | | | | | | | | | | | | | | | | | | | | | | 1 | |
| 357599074 | | | | | | | | | | | | | | | | | | | | | | | | | 7 | 2 | 1 | |
| 357607932 | | | | | | | | | | | | | | 5 | 1 | | | 1 | 1 | 1 | | 2 | 1 | 1 | | | | |
| 357814777 | 4 | 2 | | | | | | | | | | 2 | 1 | 2 | 1 | | | | | | | | | | | | | |
| 357862532 | | | | | | | | | | | | | | | | | | | | | | | 1 | | | | | |
| 357863646 | | | | | | | | | | | | | | | | | | | | | | | 3 | | | | | |
| 357909813 | | | | | | | | | | | | | | | | | | | | | | | 1 | 1 | | | | |
| 357923095 | | | | | | | | | | | | | | 1 | 1 | | | | | | 2 | | 1 | 3 | | 5 | 2 | |
| 357932060 | 1 | | | | | | | | | 1 | | | | 4 | 1 | | | | | | 1 | | 3 | 2 | | | | |
| 357940977 | | | | | | | | | | | 3 | 1 | | 15 | 9 | | | 7 | 3 | 5 | 4 | 1 | | 4 | | 1 | 1 | |
| 357944721 | | | | | | | | | | 2 | | 1 | 2 | 14 | 21 | | | | | | | | | | | | | |
| 357944930 | 1 | | | | | | | | | | | | | | | | | 3 | 1 | 2 | | 1 | 5 | | | 1 | 6 | |
| 357945095 | 1 | | | | | | | | | | | | | 2 | 3 | | | | 2 | 1 | | | 3 | 1 | | | | |
| 357945155 | | | | | | | | | | | | | 2 | 1 | | | | | | | | | | 2 | 56 | 19 | 20 | |
| 357949102 | | | | | | | | | | | 1 | 1 | 1 | 21 | 21 | | 1 | 5 | 6 | 4 | 1 | 8 | 10 | 28 | | | | |
| 357949364 | | | | | | | | | 3 | 2 | | | | | 3 | | | 2 | 1 | | 2 | | 16 | 23 | 1 | | | |
| 357953145 | | | | | | | | | 1 | 1 | | | | 1 | | | | | | | | | | | | | | |
| 358082499 | | | | | | | | | | 2 | | | | 1 | | | | | | | | | | | | | | |
| 358251878 | | | | | | | | | | | | | | | | | | | | | | | | 1 | | | | |

| | | | | | | | | | | | | | | | | | | | | | | | | |
|---|---|---|---|---|---|---|---|---|---|---|---|---|---|---|---|---|---|---|---|---|---|---|---|---|
| 358264919 | | | | | | | | | | | | | | | | | | | | 1 | | | | |
| 358277352 | | | | | | | | | | | | | | | | 1 | | | | | | | | |
| 358281935 | | | | 1 | | | | | 1 | 1 | 1 | | | 11 | 7 | 5 | 7 | 1 | | | | 3 | 1 | |
| 358281997 | | | | | | | | | | | | | | | | | | | | 1 | | | | |
| 358285424 | | | | | | 5 | 9 | | | | 1 | | | | | | | | | | | | | |
| 358286194 | | | | | | | 1 | | | | | | | 1 | | 4 | 1 | | | | | 25 | 29 | |
| 358286442 | 4 | 2 | 11 | | | | | 1 | | | | | | | | | | | 4 | 7 | | | | |
| 358290805 | | | | | | | | 1 | 1 | | 1 | | | | | | | | | | 6 | | | 21 |
| 358295017 | | | | | | | 4 | | | | | | | | | | | | | | | | | |
| 358501372 | | | | | | | | 1 | | | | | | | | | | | | | | | | |
| 358515278 | | | | | | | | | | 1 | | 1 | | | | | | | | | | 1 | | |
| 358593042 | | | | | | | | | | | 6 | | | | | | | | 3 | 4 | | | | |
| 358613806 | | | | | | | | | | | | | | | | | | | | 1 | | | | |
| 358627224 | | | | | | | | | | | | | | 1 | | | 2 | | | | | | | |
| 358631295 | | | | | | 1 | | | | | | | | | | | | | | | | | | |
| 358631384 | | | | | | | | | 1 | | | | | 1 | | | | | | | | | | |
| 358631450 | | | | | | 1 | | | | 2 | 9 | | | | 1 | | | 2 | 3 | 3 | | | 1 | |
| 358631537 | | | | | | 2 | 2 | 2 | 2 | 3 | 3 | | | | | | | | 2 | 2 | | 1 | | |
| 358635284 | | | | | 1 | 2 | 7 | | | | | | | | | | | | | | 1 | | | |
| 358639742 | | | | | | | | 16 | 30 | | | | | | | | | | | | | 1 | | |
| 358639884 | | | | | 1 | 2 | 7 | | | 1 | 10 | | 1 | | | | | | | | | | | |
| 358725902 | | | | | | | | | | | | | | | | | | | | 1 | | | | |
| 358847551 | | | | | | | | 1 | 4 | | | 1 | | | | | | | | | 3 | 1 | | |
| 358879127 | | | | | | | | 1 | | | | | 1 | | | | | | | | | | | |
| 358890923 | | | | | | | | | | | | | | | | | | | | | 6 | | | |
| 358976440 | | | | | | | 2 | | | 15 | 27 | | | 13 | 10 | 8 | 6 | 9 | 12 | 19 | 2 | 1 | | |
| 358976515 | | | | | 2 | 5 | 10 | | | 2 | | | | | | | | | 1 | | | | | |
| 358985893 | | | | | | | | | | | | | | 1 | | | | | | | | | | |
| 359027729 | | | | | 2 | | | | | | | | | | | | | | | | | | | |
| 359317340 | | | | | | | | | | | | | | | | | | | | | 1 | | | |
| 359326692 | | | | | | | 1 | | | | | | | | | | | | 1 | 1 | | | | |
| 359326764 | | | | | | | 1 | | | | 2 | | | 1 | 1 | 1 | 2 | 1 | 15 | 12 | | 1 | | |
| 359330004 | | | | | | 1 | | | | | | | | | | | | | | | | | | |
| 359407882 | | | | | | 2 | | | | | | | | | | | | | | | | | | |
| 359447314 | | | | | | | | | | | | | | | | | | | 2 | | | | | |
| 359560701 | | | | | | | | | | | | | | | | | | | | 2 | | | | |
| 359744616 | | | | | | 1 | | | | | | | | | | | | | | | | | | |
| 359744768 | | | | | | 2 | | | | | | | | | | | | | | | | | | |
| 359749513 | | | | | 3 | 7 | | | | | | | | | | | | | | | | | | |
| 360081309 | | | | | | 3 | | | | | | | | | | | | | | | | | | |
| 360081407 | | | | | 1 | | | | | | | | | | | | | | | | | | | |
| 360091049 | | | | | 1 | | | | | | | | | | | | | | | | | | | |
| 360091238 | | | | | 2 | | | | | | | | | | | | | | | | | | | |
| 360242173 | | | | | | | 1 | | | | | | | | | | | | | | | | | |
| 360254108 | | | | | 12 | 15 | 15 | | | | | | | | | | | | | | 1 | | | |
| 360263797 | | | | | | | | | 3 | | | | | | | | | | | | | | | |
| 360426854 | | | | | 3 | | | | | | | | | | | | | | | | | | | |
| 360427510 | | | | | | 2 | | | | | | | | | | | | | | | | | | |
| 360431375 | | | | | 3 | | | | | | | | | | | | | | | | | | | |
| 360432213 | | | | | 4 | 2 | | | | | | | | | | | | | | | | | | |
| 360677632 | | | | | | | | 8 | 11 | 11 | 2 | | | | 2 | | | | | | 1 | 1 | | |
| 360690414 | | | | | 2 | 11 | 13 | | 1 | 2 | 3 | | | | | | | | | | | | | |
| 360698817 | | | | | 14 | 4 | 3 | | | | | | | | | | | | | | | | | |
| 360763630 | | | | | 1 | 1 | | | | | | | | | | | | | | | | | | |
| 360764170 | | | | | 5 | 3 | | | | | | | | | | | | | | | | | | |
| 360768754 | | | | | | 1 | | | | | | | | | | | | | | | | | | |
| 361005850 | | | | | | | | | | | | | | | | | | | | | 1 | | 1 | |
| 361040812 | | | | | | | | | 2 | | | | | | | | | | | | | | | |
| 361104490 | | | | | | | | | | | | | | | | | | | | | | | | 1 |
| 361298637 | | | | | | 3 | | | | | | | | | | | | | | | | | | |
| 361312146 | | | | | 1 | 9 | 3 | | | | | | | | | | | | | | | | | |
| 361312808 | | | | | | | | | | | | | | | | | | | | | 2 | | | |
| 361355033 | | | | | | | 2 | | | | | | | | | | | | | | | | | |
| 361376643 | | | | | | | | | | | | | | | | | | | | | 4 | | | |
| 361455153 | | | | | 2 | | | | | | | | | | | | | | | | | | | |
| 361636071 | | | | | 2 | 4 | 2 | | | | | | | | | | | | | | | | | |
| 361704572 | | | | | | | | | | | | | | | | | | | | | 3 | | | |
| 361722354 | | | | | | | | | 1 | | | | | | | | | | | | | | | |
| 361787030 | | | | | | 1 | | 3 | 1 | | | | | | | | | | | | | | | 1 |
| 361791920 | | | | | 2 | | | | | | | | | | | | | | | | | | | |
| 362019898 | | | | | | | | | | | | | | | | | | | | | 5 | | | |
| 362123158 | | | | | | | | | | | 1 | | | | | | | | | | | | | |
| 362394990 | | | | | | | | | | | | | | | | | | | 4 | | | | | |
| 362473525 | | | | | | | | 3 | 1 | | | | 1 | 1 | | 1 | 1 | | | | | | 1 | 21 |
| 362719326 | | | | | | | | | | | | | | | | | | | | | 1 | | | |
| 362809689 | | | | | 1 | 1 | | | | | | | | | | | | | | | | | | |
| 362814519 | | | | | 1 | | | 1 | | | | | | | | | | | | | | | | |
| 362862047 | | | | | | | | | | | | | | | | | | | | | | | | 1 |
| 386506469 | | | | | | | | | 1 | | | | | | | | | | | | 4 | | | |
| 386829647 | | | | | | | | | | 1 | | | | | | | | | | | | | | |

| | | | | | | | | | | | | | | | | | | | | | | | |
|---|---|---|---|---|---|---|---|---|---|---|---|---|---|---|---|---|---|---|---|---|---|---|---|
| 386833812 | | | | | | | | | | 1 | | 4 | | | | | | | 1 | | 1 | 1 | 1 |
| 386833850 | | | | | | | | | 1 | 1 | 1 | 17 | 21 | | | | | | 3 | 3 | | | |
| 386834269 | 1 | | | | | | | 2 | | 1 | 3 | | | | 3 | 1 | | | | 5 | 1 | 1 | 1 |
| 386838814 | | | | | | | | | | | | 1 | 4 | | | | | | | | | | |
| 386847799 | | | | | 1 | | 3 | | | 1 | | | | | | | | | 1 | 3 | | | |
| 386877197 | | | | | | | | | | | | | | | | | | | 3 | | | | |
| 387166379 | | | | | | | | | | 13 | 14 | | 1 | 2 | 2 | 1 | | | 1 | 2 | | 1 | |
| 387170738 | | | | | | | | | | | | | | | | | | | 1 | | | | |
| 387180498 | | | | | | | | 1 | | | | | 1 | | | | | | 1 | | | | |
| 387503511 | | 1 | | 1 | | | | | | 2 | 1 | | | | | | | | 9 | 1 | 1 | 1 | |
| 387507495 | | | | | | | | | | | | | | | | | | | 1 | 4 | | | 1 |
| 387556524 | | | | | | | | | | | | | | | | | | | | | | 2 | |
| 387943673 | | | | | | | | | | | | | | | | | | | | 1 | | | |
| 387944118 | | | | | | 1 | | | | 10 | 9 | | | 1 | 1 | | | 1 | 3 | 3 | | 1 | 1 |
| 387952104 | | | | | | | | | 1 | | | | | | | | | | 21 | 3 | 2 | | 2 |
| 388280297 | | | | | | | | | | | 1 | | | | | | | | | | 3 | 2 | 1 |
| 388280529 | | | | | | | | | 2 | 4 | 1 | | | | | | | | | | | | |
| 388289013 | | | | | | | | | | | | | | 1 | | | 1 | | | | | | |
| 388289350 | | | | | | | | | | | | | | | | | | | | 1 | | | |
| 388293040 | | | | | | | | | | 1 | | | | | | | | 1 | | | | | |
| 388293048 | | | | | | | | | | | | | | | | | | | | 3 | | | |
| 388293131 | | | | | | | | | | | | | | | | | | | 1 | 2 | | | 1 |
| 388297625 | | | | | | | | | | 1 | | 4 | | | | | | | | | | | 1 |
| 388517539 | | | | | | | | | | | | | | | | | | | 1 | 5 | | | |
| 388605118 | | | | | | | | | | | | | | | | | | | 1 | | | | |
| 388612388 | | | | | | | | | | | | | | | | | | | 2 | 2 | | | |
| 388612768 | | | | | | | | | | | | | | | | | | | | 1 | | | |
| 388638672 | | | | | | | | 3 | 3 | 1 | | | | | 1 | | | 1 | | | 3 | 3 | 1 |
| 388859011 | | | | | | | | | | | | | | | | | | | | 3 | | | |
| 388877336 | | | | | | | | | | | | | 1 | | | | | | | | | | |
| 388881226 | 1 | | | | | | | 13 | 12 | 9 | 3 | 23 | 36 | | | | | | 1 | | | | 1 |
| 388924424 | | | | | | | | | | | | 1 | | | | | | | | | | | |
| 388954086 | | | | | | 1 | | 8 | 2 | | 2 | | | | | | | | | | 2 | | 1 |
| 388975834 | | | | | | | | | | 1 | | | | | | | | | | | 31 | 19 | 13 |
| 388984132 | | | | | | | | 1 | | | 1 | | | | | | | | 1 | 1 | 3 | 3 | 9 |
| 389195896 | | | | | | | | | | 1 | | | | | | | | | 4 | 3 | 2 | | |
| 389199936 | | | | | | | | 1 | 2 | 4 | 8 | 4 | 5 | | | 1 | | | | 2 | | 1 | |
| 389210230 | | | | | | | | | | | | | | | | | | | | | 1 | | |
| 389247395 | | | | | | | 1 | | | | | | | | | | | | | | | | |
| 389286412 | | | | | | | | | | | | | | | | | | | | 3 | | | |
| 389311717 | | | | | | | | | | | | | | | | | | | 2 | 3 | | 2 | |
| 389325067 | | | | | | | 2 | | | | | | | | | | | | | | | | |
| 389419768 | | | | | | | | | | 4 | 1 | | | | 1 | | 1 | | | | | | |
| 389419777 | | | | | | | | 1 | | 1 | 1 | | | | | 1 | | | | | | | |
| 389419787 | | | | | | | | | | 6 | 3 | | | | | | | | | | | | |
| 389527383 | | | | | | | | | | | | | | | | | | | 8 | 43 | 4 | | |
| 389563849 | | | | | | | | 1 | | | | | | | | | | | | | | | |
| 389568268 | | | | | | 1 | 1 | | | | | | | | | | | | | | | | |
| 389589423 | | | | | | | | | | | | | | | | | | | 1 | 2 | | | |
| 389626867 | | | | | | | | | | | | | | | | | | | | | 1 | | |
| 389657555 | | | | | | | 1 | | 1 | 1 | | | | | | | | 1 | 2 | 1 | 5 | 43 | 59 |
| 389687699 | | | | | | | | | 1 | | | | | | | | | | | 1 | 3 | 1 | |
| 389760846 | | | | | | | | 2 | | | | | | | | | | | | | | | |
| 389976940 | | | | | | | | | | | | | | | | | | | | | 1 | | |
| 390003153 | | | | | | 2 | 1 | | | | | | | | | | | | | | | | |
| 390024875 | | | | | 2 | 2 | 2 | | | | | | | | | | | | | | | | |
| 390024952 | | | | | 1 | | | | | | | | | | | | | | | | | | |
| 390097182 | | | | | 1 | | | | | | | | | | | | | | | | | | |
| 390101630 | | | | | | | | | | | 1 | | | | | | 1 | | | | | | |
| 390101921 | | | | | | 1 | | | | | | | | | | | | | | | | | |
| 390280026 | | | | | | | | | | | | | | | | | | | | | 16 | | |
| 390331583 | | | | | 2 | | | 33 | 30 | 72 | 65 | | | | 1 | | | | 1 | 1 | 1 | 5 | 3 |
| 390344371 | | 2 | 3 | 4 | | 1 | 6 | | | 1 | 1 | 3 | 2 | | | | | | 1 | 1 | | | |
| 390352384 | | | | | | | 1 | | | | | | | | | | | | | | 1 | | |
| 390356988 | | | | | 2 | 1 | 4 | | | | | 3 | 2 | | | | | | | | | | |
| 390361224 | | | | | 8 | 5 | 13 | | | | | | | 2 | | | | | | | | 1 | |
| 390366259 | | | | | | | 1 | | | | | | | | | | | | | | | | |
| 390378650 | | | | | 1 | | 1 | | | | | | | | 1 | | | | | | | | |
| 390438574 | | | | | | 1 | | | | 15 | 9 | | | 4 | 1 | | | 2 | | | | | |
| 390442730 | | | | | | | 2 | | | | | | | | | | | | | | | | |
| 390443308 | | | | | 1 | | | | | | | | | | | | | | | | | | |
| 390447010 | | | | | 2 | 6 | | | | | | | | | | | | | | | | | |
| 390473497 | | | | | | | | | | | | | | | | 1 | | 3 | | | | | |
| 390579059 | | | | | | | | | | | | | | | | | | | | | 2 | | |
| 390590450 | | | | | | | | 1 | 2 | 1 | | | | | | 1 | | | 30 | 27 | 3 | 8 | 4 |
| 390697959 | | | | | | 1 | 1 | | | | | | | | | | | | | | | | |
| 390706983 | | | | | | | | 1 | 1 | | | | | 2 | 1 | 2 | 2 | 2 | 6 | 3 | | | |
| 390779758 | | | | | | | | 1 | | | | | | | | | | | | | | | |
| 390780035 | | | | | | | | | | 1 | 1 | | | | | | | | | | | | |
| 390918452 | | | | | | | | | | | | | | | | | | | 1 | | | | |

| | | | | | | | | | | | | | | | | | | | | | | | | |
|---|---|---|---|---|---|---|---|---|---|---|---|---|---|---|---|---|---|---|---|---|---|---|---|---|
| 391016873 | | | | | 2 | | | | | 14 | 12 | | | | | | | | | | | | | |
| 391026123 | | | | | | | | 1 | | | 2 | 1 | 2 | 2 | 1 | | 1 | 2 | 5 | 8 | 3 | | | |
| 391034580 | | | | | | | | | | | | | | 6 | | | 1 | | | | 1 | 3 | 4 | |
| 391052159 | | | | | 5 | 2 | 2 | 2 | 1 | 6 | | | | | | | | | | | | | | |
| 391056169 | | | | | 1 | | 2 | | | | | | | | | | | | | | | | | |
| 391120561 | | | | | 1 | | | | | 1 | | | | | | | | | | | | | | |
| 391121449 | | | | | | 2 | | | | | | | | | | | | | | | | | | |
| 391124880 | | | | | 3 | | | | | | | | | | | | | | | | | | | |
| 391362634 | | | | | | | | | | | | 3 | 4 | | | | | | | | | | | |
| 391457659 | | | | | 5 | | | | | | | | | | | | | | | | | | | |
| 391458054 | | | | | 2 | 3 | | | | | | | | | | | | | | | | | | |
| 391462340 | | | | | 1 | | | | | | | | | | | | | | | | | | | |
| 391496898 | | | | | | | | | | | | | | | | | | | | | | | | 2 |
| 391618398 | | | | | | | | | | | | | | | | | | | | | 1 | | | |
| 391725625 | | | | | | | 1 | | | | | | | | | | | | | | | | | |
| 391799282 | | | | | | 1 | | 5 | 4 | | | | | | | | | | | | | | | |
| 391802788 | | | | | 1 | 1 | 1 | | | | | | | | | | | | | | | | | |
| 392002352 | | | | | | | | | | | | | | | | | | | 4 | 2 | | | | |
| 392062145 | | | | | | | 1 | | | | | | | | | | | | | | | | | |
| 392130578 | | | | | | | | | | | | | | | | | | | | | | | | 23 |
| 392325925 | | | | | 5 | 5 | 2 | | | | | | | | | | | | | | | | | |
| 392368893 | | | | | | | | | | | | 1 | | | | | | | | | | | | |
| 392411222 | | | | | | | 1 | | 4 | 1 | | | | | | | | | 1 | | 46 | 14 | 6 | |
| 392636270 | | | | | | | | | | | | | | | | | | | | | 2 | | | |
| 392748193 | | | | | | 1 | 1 | | | | | | | | | | | | | | | | | |
| 392821837 | | | | | 4 | 2 | 1 | | | | | | | | | | | | | | | | | |
| 392822059 | | | | | | | | 5 | 6 | | | | | | | | | | | | | | | 1 |
| 393063351 | | | | | | | | 8 | 6 | 5 | 4 | | | | | | | | | | 5 | 5 | 2 | 3 |
| 393158731 | | | | | 1 | | | | | | | | | | | | | | | | | | | |
| 393162877 | | | | | | 2 | | | | | | | | | | | | | | | | | | |
| 393215059 | | | | | | | | | | | | | | | | | | | | | | | | 1 |
| 393503853 | | | | | 2 | 1 | | | | | | | | | | | | | | | | | | |
| 393766777 | | | | | | | | | | | | | | | | | | | | | 3 | | | |
| 393776005 | | | | | | | | | | | | | | | | | | | | | 104 | 12 | 14 | 9 |
| 394095053 | | | | | | | | | | | | | | | | | | | | | 2 | | | |
| 417143726 | | | | | | | | | | 21 | 37 | | 2 | 5 | | 13 | 1 | 3 | 15 | 13 | 1 | 1 | 2 | |
| 417190793 | | 4 | | | | | | 1 | 3 | 3 | 10 | | 4 | 1 | | 1 | | | 2 | | 1 | | | |
| 417199910 | | | | | | | 1 | | | | | | | | | | | | 1 | 1 | | | | |
| 417204164 | | | | | | | | | | | 2 | | | | | | | | 4 | | | | | |
| 417204354 | | | | 1 | | | 1 | | | 1 | 3 | 2 | | | | | | | | | | | 3 | |
| 417221397 | | | | | | | | | | 1 | | | | | | | | | | 1 | | | | |
| 417536905 | | | | | | 1 | 5 | | | 4 | 9 | | | | | | | | 2 | | | 1 | | |
| 417540543 | | | | | | | | | | 2 | 1 | 1 | | | | | | | | | | | | |
| 417554241 | | | | | | | | | | | | | 1 | | | | | | | | 1 | | | |
| 417558532 | | | | | | | | | | | | 1 | 2 | | | | | | | | | | 2 | |
| 417869138 | | | | | | | | | | | | | | | | | | | | 1 | | | | |
| 417882515 | | | | | | | | | | 2 | 1 | 1 | | | | | | | | 2 | 1 | 2 | 1 | |
| 417890128 | | | | | | | | | | 1 | | 21 | 21 | | | | | | | | | | | |
| 417908800 | | | | | | 1 | | 6 | 9 | 1 | | | | | | | | | | | | | | |
| 418224106 | 2 | 1 | 2 | | | | | | | | | 1 | | | | | | | 1 | | | | | |
| 418224351 | | | | | | | | | | | | | | | | | 1 | | | 1 | 1 | | 1 | |
| 418261858 | | | | | | | | 1 | | 7 | 2 | | | | | | | | | | | | | |
| 418607328 | | | | | | | | | | | | | | | | | | | 1 | | | | | |
| 418641700 | | | | | | | | 1 | | 2 | | | | | 2 | 1 | | | 2 | 5 | | 18 | 18 | |
| 418645869 | | | | | | | | 1 | 1 | 22 | 19 | | | 2 | | | | | | | | | | |
| 418650337 | | | | | | | | | | | | | | 1 | | | | | | | | | | |
| 418650351 | | | | | | | | | 2 | 7 | 2 | | | 26 | 32 | 21 | 36 | 30 | | | | 2 | 1 | |
| 418888180 | | | | | | | | 1 | | | | | 2 | | | | | | 9 | 33 | | | | |
| 418969956 | | | | | | | | | | | | | | | | | | | | 1 | 3 | | 1 | |
| 418987009 | | | | | | | | | | | | | | | | | | | | 5 | 1 | | 2 | |
| 419000001 | | | | | | | | | | 2 | 2 | | | | | | | 1 | | | | | | |
| 419337157 | | | | | | 3 | 4 | | | | | | | | | | | | | | | 1 | | |
| 419341292 | | | | | | | | 1 | | | | | | | | | | | | | 2 | | | |
| 419345466 | | | | | 3 | 1 | 2 | | | | | | | | | | | | | | | | | |
| 419345606 | | | | | | | | 3 | 2 | 1 | | | | | | | | | 2 | 1 | | 20 | 19 | |
| 419466429 | | | | | | | | 4 | 4 | 2 | 1 | | | | | | | | | 1 | 3 | 11 | 10 | 12 |
| 419669180 | | | | | | | | | | | | | | | | | | | | 2 | | 4 | 1 | |
| 419669547 | | | | | | | | | | | | | | 2 | | | | | | | | | | |
| 419807055 | | | | | | | | | | | 3 | | | | | | | | | | | | | |
| 420010739 | | | | | | | | | | | | | | | | | | | | | 3 | | | |
| 420027427 | 1 | | | | | 1 | | | | 17 | 20 | | | 10 | 3 | 7 | 16 | 7 | 7 | 32 | 1 | | | |
| 420027557 | | | | | 2 | | | | | | | | | | | | | | | | | | | |
| 420036032 | | | | | | | | | | | | | | | | | | | 1 | | | | 1 | |
| 420113535 | | | | | | | | | | | 1 | | | | 1 | | | 2 | | | | | | |
| 420221276 | | | | | | | | 1 | 1 | 1 | 3 | | 1 | 1 | 1 | 3 | | | | | | | | |
| 420325753 | | | | | | | | | | 1 | | | | | | | | | | | 2 | | | |
| 420347328 | | | | | | | | | | 1 | | | | | | | | | 1 | | | | | |
| 420347360 | | | | | | | | | | | | | | | | | | | 1 | 3 | | | | |
| 420351087 | | | | | | | | | | 1 | 1 | | | | 1 | | | | 1 | 2 | 27 | 46 | 50 | |
| 420351828 | | | | | 4 | 1 | | | 2 | 1 | | | | | | | | | | | | 3 | 6 | 1 |

| | | | | | | | | | | | | | | | | | | | | | | | | | | | | |
|---|---|---|---|---|---|---|---|---|---|---|---|---|---|---|---|---|---|---|---|---|---|---|---|---|---|---|---|---|
| 420359820 | | | | | | | | | | 1 | | | | | | | | | | | | | | | | | | |
| 420382647 | | | | | | | | | | | | | | 1 | | | | | | 1 | 1 | | | 2 | | | 1 | |
| 420386244 | | | | | | | | | | | | | | 1 | 2 | | | 1 | | | | 2 | | | 2 | | | |
| 420454837 | | | | | | | | | | | | | | | | | | | | | | 1 | | | | | | |
| 420455015 | | | | | | | | | | | | | | | | | | | | | 1 | | | | | | | |
| 420571563 | | | | | | | | | | | | | | 1 | | | | | | | | | | 1 | | | | |
| 420586101 | | | | | | | | | | | | 1 | 3 | 9 | | | | | | | | | | 1 | | 2 | | |
| 420688241 | | | | | | | | | | | | | | | | | | | | | | | | | | 1 | | |
| 420688294 | | | | | | | | | | | | | | 1 | | | | | | | | | | | 2 | | | |
| 420723066 | | | | | | | | | | | | | 2 | | | | | | | | | | | | | 1 | | |
| 420795489 | | | | | | | | | 1 | 1 | | | | | | | | | | | | | | | | | | |
| 420796184 | | | | | | | | | | 1 | | | | | | | | | | | | | | | | | | |
| 420799910 | | | | | | | | | | | | 1 | | | | | | | | | 1 | | 2 | | | | | 1 |
| 420834927 | | | | | | | | | | | | | 1 | | | | | | | | | | | | | | | |
| 420904348 | | | | | | | | | | | 1 | | | | | | | | | | | | | 6 | | | | |
| 421033734 | | | | | | | | | | | 3 | 1 | | 16 | 9 | | | | | | | | | | | 1 | | |
| 421041565 | | | | | | | | | | | 1 | | | | | | | | | | | | | | | | 1 | |
| 421054850 | | | | | | | | | 1 | | | | | 2 | 3 | | | | | | | | 8 | 28 | | | | |
| 421055181 | | | | | | | | | | | | | | 2 | | | | 2 | | | | 1 | 2 | 25 | | 1 | | |
| 421068062 | | | | | | | | | | | | 6 | 6 | 32 | 39 | | | 3 | 2 | | | | | | | 1 | 4 | |
| 421128142 | | | | | | | | | | 2 | | | | | | | | | | | | | | | | | | |
| 421132639 | | | | | | | | | | 1 | | | | | | | | | | | | | | | | | | |
| 421136849 | | | | | | | | | 1 | | | | | | | | | | | | | | | | | | | |
| 421137065 | | | | | | | | | 1 | | | | | | | | | | | | | | | | | | | |
| 421137107 | | | | | | | | | | 1 | | | | | | | | | | | 2 | | | | | | | |
| 421271305 | | | | | | | | | | 1 | | | | | | | | | | | | | | | | | | |
| 421357275 | | | | | | | | | | | | | | | | | | | | | | | | | 2 | | | |
| 421405157 | | | | | | | | | 1 | | | | | 5 | 2 | | | 5 | 2 | 3 | 2 | 1 | 7 | 8 | 12 | 38 | 35 | |
| 421469164 | | | | | | | | | 1 | 2 | | | | | | | | | | | | | | | | | | |
| 421469304 | | | | | | | | | 1 | 1 | | | | | | | | | | | | | | | | | | |
| 421477870 | | | | | | | | | | | | | | 9 | 4 | | | 2 | 4 | | 2 | 1 | | | | | | |
| 421605182 | | | | | | | | | | | 1 | | | | | | | | | | | | | | | | | |
| 421746220 | | | | | | | | | | 1 | | | | | | | | | | | | | | | | | | |
| 421810160 | | | | | | | | | | 1 | | | | | | | | | | | | | | | | | | |
| 421810214 | | | | | | | | | 3 | | | | | | | | | | | | | | | | | | | |
| 421814868 | | | | | | | | | | | | 2 | | | | | | | | | | | | | | | | |
| 421815053 | | | | | | | | | 2 | 6 | | | | 4 | | | | | | | | | | | | | | |
| 421946059 | | | | | | | | | | | | | | | | | | | | | | | | | 5 | | | |
| 422052346 | | | | | | | | | | | | | 7 | 3 | 3 | | | | 2 | | | | | | 6 | 5 | 4 | |
| 422056521 | | | | | | | | | | | | | 1 | | | | | | | | | | | | | | | |
| 422151417 | | | | | | | | | 1 | | | | | | | | | | | | | | | | | | | |
| 422155460 | | | | | | | | | | 3 | | 1 | | | | | | | | | | | | | | | | |
| 422155824 | | | | | | | | | | 1 | | | | | | | | | | | | | | | | | | |
| 422156254 | | | | | | | | | | 2 | | | | | | | | | | | | | | | | | | |
| 422156333 | | | | | | | | | | 3 | | | | | | | | | | | | | | | | | | |
| 422160219 | | | | | | | | | 1 | | | | | | | | | | | | | | | | | | | |
| 422419397 | | | | | | | | | | | | | 3 | 12 | 5 | | | | | | | | | | | | 1 | |
| 422492727 | | | | | | | | | | | | 1 | | | | | | | | | | | | | | | | |
| 422496798 | | | | | | | | | | 1 | | | | | | | | | | | | | | | | | | |
| 422496879 | | | | | | | | | | 1 | | | | | | | | | | | | | | | | | | |
| 422497047 | | | | | | | | | | 1 | | | | | | | | | | | | | | | | | | |
| 422622063 | | | | | | | | | | | | | | | | | | | | | | | | | 1 | | | |
| 422777746 | | | | | | | | | 2 | | | | | | | | | | | | | | | | | | | |
| 422833659 | | | | | | | | | | 1 | | | | | | | | | | | | | | | | | | |
| 422833681 | | | | | | | | | | 2 | | | | | | | | | | | | | | | | | | |
| 423101189 | | | | | | | | | | | | 1 | 5 | | | | | | | | | | | | | 10 | 6 | |
| 423174813 | | | | | | | | | 3 | 1 | | | | | | | | | | | | | | | | | | |
| 423183023 | | | | | | | | | | 2 | | | | | | | | | | | | | | | | | | |
| 423442357 | | | | | | | | | 3 | | 5 | | | | | | | | | | | | | | | | | |
| 423516229 | | | | | | | | | | | 1 | | | | | | | | | | | | | | 1 | | | |
| 423792440 | | | | | | | | | | | | | | | | | | | | | | | | | 2 | | | |
| 423852469 | | | | | | | | | | | | | | | | | | | | | | | | | 1 | | | 3 |
| 424037884 | | | | | | | | | | 2 | | | | | | | | | | | | | | | | | | |
| 424193190 | | | | | | | | | | | | | | | | | | | | | | | | | | | | 5 |
| 424193645 | | | | | | | | | | | | | | | | | | | | | | | | | | | | 1 |
| 424443911 | | | | | | | | | | | | | | 3 | | | | | | | | | 1 | | | | | |
| 424452682 | | | | | | | | | | | | | | | | | | | | | | | | | 2 | | | |
| 424792976 | | | | | | | | | | | | | | | | | | | | | | | | | | 1 | | |
| 424793122 | | | | | | | | | | | | | | | | | | | | | | | | | 2 | | | |
| 424923616 | | | | | | | | | | | | 1 | | | | | | | | | | | | | | | | |
| 425143700 | | | | | | | | | 2 | | | | | | | | | | | | | | | | | | | |
| 425148091 | | | | | | | | | | | | | | | | | | | | | | | | | 80 | 5 | 8 | 13 |
| 447599745 | | | | | | | | | | | | | | | | 1 | 3 | | | | | | | 2 | | | | |
| 448182402 | 1 | 2 | | | | | | | | | | | | | | | | | | | | | | | | | | |
| 448234999 | | 1 | | | | | | | | 1 | 4 | 1 | | | | | | | | | | | | 1 | | 1 | | |
| 448238871 | | | | 1 | | | | | | | | | | 7 | 11 | | | 4 | 2 | 6 | 3 | 2 | 3 | 2 | | 1 | | |
| 448260940 | 1 | | | | | | | | 6 | | 3 | | | | | | | | | 1 | | | 1 | 1 | | | | |
| 448515267 | | | | | | | | | | | 1 | | | | | | | 6 | 4 | 8 | 5 | | | 7 | | | | |
| 448942604 | | | | | | | | | | | | | 2 | | | | | | | | | | | | | | 1 | |
| 448994377 | | | | | | | | | | | | | | | | 1 | | | | | | | | | | | | |

| | | | | | | | | | | | | | | | | | | | | | | | | |
|---|---|---|---|---|---|---|---|---|---|---|---|---|---|---|---|---|---|---|---|---|---|---|---|---|
| 449236636 | | | | | | | | | | | | | | | | | | | 6 | 11 | | | | |
| 449244969 | | | | | | | | | | | | | | | | | | | | | | | 2 | |
| 449285328 | | | | | | | | | | | | | | | | | | | 1 | | 1 | | | |
| 449288517 | | | | | | | | 9 | 34 | | | | | 1 | 1 | 1 | 1 | | 1 | | | 1 | 1 | |
| 449296930 | | | | | | | | 1 | | | | | | | | | | | | | 6 | | | |
| 449331241 | | | | | | | | | | 1 | 1 | | | 1 | 1 | 2 | 1 | 1 | 1 | 2 | | | | |
| 449335798 | 1 | | 1 | | | | | | | 14 | 24 | | | 10 | 5 | 4 | 6 | 10 | 27 | 21 | | 2 | | |
| 449339510 | | | | | | | | | | 4 | | | | | | | | | | | | | | |
| 449339532 | | | | | | | | | | | | | | | | | 1 | | | | | | | |
| 449339838 | | | | | | | | | | | | | | | | 3 | 1 | | | | | | | |
| 449344351 | | | | | | | | | | | | | | | 1 | 1 | | | | | | 2 | | |
| 449490682 | | | | | | | | | | | | | | 1 | 3 | 2 | | | | | | | | |
| 449634539 | | | | | | | | | | | | | | | | | | | | | 1 | 1 | | |
| 449681157 | 5 | 4 | 6 | 3 | | | | | | | 1 | 11 | 13 | 1 | 1 | | | | 2 | 1 | | 1 | | |
| 449685778 | | | | | | | | | | 2 | 7 | | | 2 | | | | 3 | | 2 | 4 | 13 | 18 | |
| 449910741 | | | | | | | | | | 1 | | | | | | | | | | | | | | |
| 449975227 | | | | | | | | 2 | | 2 | 5 | | | | | | | | | 1 | 1 | 2 | 4 | |
| 450008671 | | | | | | | | | | | | | | | | | | | 12 | 2 | 1 | | | |
| 450026507 | | | | | | 1 | | | | | 2 | | | | | | | | 1 | 1 | | 6 | 3 | |
| 450034796 | | | | | | | | | | | | | | | | 1 | | | | | | | | |
| 450034902 | | | | | | | | | | | 1 | | | 4 | | | | 1 | 1 | 3 | 4 | | 45 | |
| 450035275 | | | | | 1 | | | | | | | | | | | 1 | | | | | | | | |
| 450164398 | | | | | | | | | | | | | | | | 2 | | | | | | | | |
| 450302778 | | | | | 1 | | | | | | | | | | | | | | | | | | | |
| 450354788 | | | | | | | | | | | 4 | | 1 | | | | | | 2 | 1 | | | | |
| 450371612 | | | | | 2 | | | 1 | 1 | 3 | | | | | | | | 2 | | 2 | | | 1 | |
| 450376017 | | | | | 2 | 1 | | | | | | | | | | | | | | | | | | |
| 450380493 | | | | | 2 | | | | | | | | | | | | | | | | | | | |
| 450496881 | | | | | | | | | | 1 | | | | 1 | 1 | 3 | | | | | | | | |
| 450497246 | | | | | | | | | | | | | | 1 | | | | 1 | | | | | | |
| 450497356 | | | | | | | | | | | | | | 2 | | | | 3 | | 1 | | | | |
| 450497386 | | | | | | | | | | | 1 | | | | | | | | | | | | | |
| 450583886 | | | | 1 | | | | | | 2 | 2 | 4 | 1 | | | | | | 3 | 6 | 4 | 1 | | |
| 450617762 | | | | | | | | | | | | | | | | | | | 2 | 1 | | | | |
| 450656955 | | | | | 1 | | | | | | | | | | | | | | | | | | | |
| 450700075 | 1 | | | 2 | | 1 | 1 | | 2 | | 2 | 8 | 6 | | | | | | | 2 | | | | |
| 450807268 | | | | | | | | 2 | | | | | | | | | | | | | | | | |
| 450838353 | | | | | | | | | | | | | | 6 | 15 | 5 | 4 | 6 | | | | | | |
| 451040974 | | | | | | | | | | | | | | | | | | | | | 1 | | | |
| 451049385 | | | | | | | | | | | | | | | | | | | 4 | 5 | | | 1 | |
| 451054039 | | | | | | | | | | | 2 | | | | | | | | 1 | | | | | 5 |
| 451057982 | | | | | 2 | | | | | | | | | | | | | | | | | | | |
| 451062462 | | | | | | | | | | | | | | | | | | | | 3 | 5 | | | |
| 451071081 | | | | | | | | | | | 1 | | | | | | | | | | | | 2 | |
| 451071417 | | | | | 3 | | 1 | | | | 1 | | | | | | | | | | | 1 | | |
| 451080648 | | | | | | | | | | | | | | 1 | | | | | | | | | | |
| 451118413 | | | | | | | | | | | | | | 1 | | | | | | | | | | |
| 451148412 | | | | | | | | | | | | | | | | | | 1 | | | | | | |
| 451148508 | | | | | 1 | | | | | | | | | | | | | | | | | | | |
| 451148559 | | | | | | | | | | | | | | | 3 | 1 | 2 | 2 | | | | | | |
| 451148659 | | | | | | | | | 1 | 1 | | | | | | | | | | | | | | |
| 451148838 | | | | | | | | 2 | 7 | 34 | 21 | | | 6 | 4 | 1 | 1 | 3 | | | | | | |
| 451279196 | | | | | | | | 2 | | | | | | | | | | | | | 1 | | | |
| 451356082 | | | | | | 1 | | | | | | | | | | | | | | | | | | |
| 451365099 | | | | | | | | | | | | | | | | | | | | | 1 | 1 | 1 | |
| 451390676 | | | | | | | | 3 | 2 | | 1 | | | | | | | | 1 | | | | 1 | |
| 451399140 | | | | | 1 | | 1 | | | | | | | | | | | | | | | | | |
| 451424866 | | | | | | | | | | 2 | | | 1 | | 1 | 2 | | 3 | 2 | 3 | | | | |
| 451485030 | | | | | | 1 | | | | | | | | | | | | | | | | | | |
| 451485434 | | | | | | | | 1 | | | | | | | | | | | | | | | | |
| 451485461 | | | | | | 1 | | | | | | | | | | | | | | | | | | |
| 451489835 | | | | | | | | 3 | | 1 | 1 | | | | 2 | | | | | | | | | |
| 451490214 | | | | | | | | 1 | | | | | | | | | | | | | | | | |
| 451662040 | | | | | | | | | | | | | | | | | | | | | 2 | 1 | | |
| 451689001 | | 1 | | | | | | | 1 | 1 | | | | | 1 | | 1 | | 1 | 1 | 1 | 5 | 6 | |
| 451722668 | | | | | | | | 1 | 1 | | | | | | | | | | 2 | 8 | | | 1 | |
| 451735916 | | | | | 1 | | | | | | | | | | | | | | | | | | | |
| 451762072 | | | | | | | | | | | 2 | | | | | | | 2 | | | | | 1 | |
| 451765891 | | | | | | | | | | | 1 | | | | | | | | | | | | | |
| 451822015 | | | | | | 2 | | 3 | | | | | | | | | | | | | | | | |
| 452158327 | | | | | | | | | | | 1 | | | | | | | | | | | | | |
| 452158730 | | | | | | | | | | | 1 | | | | | | | | | | | | | |
| 452162916 | | | | | | | | 2 | 1 | 2 | | | | | | | | | | | | | | 21 |
| 452167518 | | | | | | | | | 1 | 10 | 7 | | | 6 | 2 | 1 | 4 | 3 | | | | | | |
| 452167550 | | | | | | 3 | | | | | | | | | | | | | | | | | | |
| 452167805 | | | | | | 3 | | | | | | | | | | | | | | | | | | |
| 452171360 | | | | | | | | | | 2 | 1 | | | | 2 | | 2 | 1 | | | | | | |
| 452171507 | | | | | | | | 2 | | 1 | 2 | | | 2 | | | 2 | | | | | | | |
| 452171631 | | | | | | 1 | | | | | | | | | | 2 | | | | | | | | |
| 452171673 | | | | | | 1 | | | | | | | | | | | | | | | | | | |

| | | | | | | | | | | | | | | | | | | | | | | | | | | | | |
|---|---|---|---|---|---|---|---|---|---|---|---|---|---|---|---|---|---|---|---|---|---|---|---|---|---|---|---|---|
| 452504314 | | | | | | | | | | | | | | 1 | | | | | | | | | | | | | | |
| 452509129 | | | | | | | | | | | | 1 | | | | | | | | | | | | | | | | |
| 452771533 | | | | | | | | | | | | | | 1 | | | | | 1 | | | | 1 | 2 | 5 | 1 | | |
| 452771913 | | | | | | | | | | | 1 | | | | | | | | | 1 | | | | | | | | |
| 452841019 | | | | | | | | | | | | 1 | 2 | 2 | 4 | | | | | | | | | | | | | 1 |
| 452841244 | | | | | | | | | | | | | | | 4 | | | | | | | | | | | | | |
| 452841273 | | | | | | | | | | | | 4 | | | | | | | | | | | | | | | | |
| 452841283 | | | | | | | | | | | | 2 | | 1 | | | | | | | | | | | | | | |
| 452845606 | | | | | | | | | | 2 | | 1 | | | | | | | | | | | | | | | | |
| 452849628 | | | | | | | | | 1 | | | | | | | | | | | | | | | | | | | |
| 452849699 | | | | | | | | | | 1 | | | | | | | | | | | | | | | | | | |
| 452953198 | 1 | 1 | 2 | 2 | | | | | | | | | 1 | 1 | | | | | | | | | | | | | | |
| 453130054 | | | | | | | | | | | | 28 | 15 | | | | | | | | | | | | 6 | 1 | | |
| 453185918 | | | | | | | | | | 6 | | 2 | | | | | | | | | | | | | | | | |
| 453195281 | | | | | | | | | | 1 | | | | | | | | | | | | | | | | | | |
| 453423834 | | | | | | | | | | | | | | | | | | | | | | | 2 | | | | 1 | |
| 453436589 | | | | | | | | | 3 | | | | | | | | | 1 | | 1 | | 2 | | | | | 1 | |
| 453457927 | | | | | | | | | | | | 2 | 1 | | | | | | | | | | | 9 | | | | |
| 453467404 | | | | | | | | | | 1 | | 1 | 1 | 3 | | | | | 1 | | | | | | | | | |
| 453522986 | | | | | | | | | | | | 3 | 4 | 14 | 6 | | | | | | | | | | | | | |
| 453527730 | | | | | | | | | | | | 1 | 1 | | | | | | | | | | | | | | | |
| 453863806 | | | | | | | | | | | | | | | | | | | | | 1 | | | | | | | |
| 453868633 | | | | | | | | | | 4 | | | | | | | | | | | | | | | | | | |
| 453873395 | | | | | | | | | | | 1 | | | | | | | | | | | | | | | | | |
| 454003707 | | | | | | | | | | | | | | | | | | | | | | | | | 1 | | | |
| 454089465 | | | | | | | | | | | | | | | | | | | | | | | | | | 3 | 1 | |
| 454217919 | | | | | | | | | | 1 | 2 | | | | | | | | | | | | | | | | | |
| 454477603 | | | | | | | | | | | | 1 | | | | | | | | | | | | | 1 | | | |
| 454481372 | | | | | | | | | | | | | | | | | | | | | | | | | 12 | 5 | | |
| 454546908 | | | | | | | | | | 2 | | | | | | | | | | | | | | | | | | |
| 454554631 | | | | | | | | | | 1 | | | | | | | | | | | | | | | | | | |
| 454822542 | | | | | | | | | | | 1 | | | | | | | | | | | | | | | | | |
| 455229034 | | | | | | | | | | 1 | | | | | | | | | | | | | | | | | | |
| 455232361 | | | | | | | | | 2 | | | | | | | | | | | | | | | | | | | |
| 455815977 | | | | | | | | | | | 1 | 7 | 2 | | 4 | | | | | | | | 1 | | 24 | 2 | 3 | |
| 455910660 | | | | | | | | | 3 | 1 | | | | | | | | | | | | | | | | | | |
| 455911138 | | | | | | | | | 2 | | | | | | | | | | | | | | | | | | | |
| 456942698 | | | | | | | | | 1 | 2 | | | | | | | | | | | | | | | | | | |
| 477918443 | 1 | 7 | | 3 | | | | | | | | | | | | | | | | 1 | 2 | | 28 | 41 | | | | |
| 478618011 | | | | | | | | | | | | | | 1 | | | | | | | | | | | | | | |
| 478928628 | | | | | | | | | | | | | | 1 | | 1 | 5 | | | | | | | | 1 | | | |
| 478937243 | | | | | | | | | 1 | 9 | 10 | | | 1 | 2 | | | | | 1 | | | 1 | | | | | |
| 479260891 | | | | | | | | | | | | 1 | | | | | | | | | | | | | | 4 | 3 | |
| 479260921 | | | | | | | | | | 3 | 2 | | | 1 | | 1 | 1 | | | 1 | | 1 | 8 | 7 | | 1 | | |
| 479290982 | 2 | 1 | | 1 | | | | | | 1 | | | | 4 | | 3 | | | | | | | | | 1 | | | |
| 479636867 | | | | | | | | | | | | | | 1 | | | | | | | | | | | | | | |
| 479637825 | | | | | | | | | | | | 1 | 9 | | | | | | | | | | | | | | | |
| 479645393 | | | | | | | | | | | | | | 1 | | | | | | 1 | | | 1 | 1 | | | | |
| 479921537 | | | | | | | | | | | | | | | | | 2 | | | | | | 30 | 67 | 1 | | | |
| 479935033 | | | | | | | | | | | | | | | | | | | | | | | 1 | | | | | |
| 479952665 | | | | | | | | | | | | | | | | 1 | | | | | | | | | | | | |
| 479987085 | | | | 2 | | | | | | | | | | | 11 | 22 | 15 | | | | | 1 | 2 | 4 | 2 | 4 | 6 | |
| 480029788 | | | | | | | | | | | | | 1 | 1 | 2 | 1 | 1 | 1 | | 1 | 2 | | | 4 | 2 | 30 | | |
| 480328501 | | | | | | | | | | | | 1 | | 3 | 2 | 1 | | | | | | | | | | | | |
| 480331310 | | | | | | | | | | | | | | | | | | | | | | | 1 | | | | | |
| 480378793 | | | | | | | | | | | | | | | | | | 1 | 1 | | 1 | 2 | | | | 4 | | |
| 480387478 | | | | | | | | | | | | | 1 | 7 | 7 | | | | 1 | 3 | | | | 1 | 3 | 9 | 19 | |
| 480391822 | | | | | | | | | | | | 1 | 1 | | 6 | | | | | | | | | | | 10 | 6 | |
| 480521256 | | | | | | | | | | | | | | | | | | | | 2 | | | | | | | | |
| 480521493 | | | | | | | | | | | | | | | 1 | | | | | | | | | | | | | |
| 480521586 | | | | | | | | | | | | | | | 1 | | | | | | | | | | | | | |
| 480525903 | | | | | | | | | | | | | | | | | | | 1 | | | 1 | | | | | | |
| 480581727 | 1 | 1 | 3 | | | | | | | | | | | 5 | 3 | | | | | | | | | | | | | |
| 480590566 | | | | | | | | | | | | | | | | 1 | | | | | | | | | | | | |
| 480681197 | | | | | | | | | | | | 1 | 2 | | | | | | | | | | | | 4 | | | |
| 480698453 | | | | | | | | | | 3 | 2 | 1 | 2 | | | | | | | | | | | | | | 2 | |
| 480737286 | | | | | | | | | | | | | | 2 | | | | | 1 | 1 | | 3 | | | | 1 | 1 | |
| 480849477 | | | | | | | | | | | | | | | | | | | | | | 1 | | | | | | |
| 480854105 | | | | | | | | | | | | | | | | | | | | 1 | | | | | | | | |
| 480854134 | | | | | | | | | | | | | | | 1 | | | | | | | | | | | | | 4 |
| 480854154 | | | | | | | | | | | | | | | | | | | | 1 | | | | | | | | |
| 480857926 | | | | | | | | | | | | | | | | | | | 1 | | | | | | | | | |
| 480866929 | | | | | | | | | | | | | | 1 | 3 | | | 1 | 1 | 1 | | | | | | | | |
| 480987689 | | | | | | | | | | | | | 4 | 3 | | | | | | | | | | | | | | |
| 481053136 | | | | | | | | | 1 | 3 | 2 | 1 | | 1 | 1 | | | 4 | 13 | 3 | 6 | 2 | | | | 7 | 2 | |
| 481056857 | | | | | | | | | | | | | | | | | | | | | | | 1 | | | 6 | 5 | |
| 481070027 | | | | | | | | | | | | 21 | 16 | 2 | | | | 2 | | 1 | | | | 2 | 11 | 8 | 18 | |
| 481190355 | | | | | | | | | | | | 3 | | 3 | | | | | 1 | 2 | | | | | | | | |
| 481190745 | | | | | | | | | | | | | | | | | | | | 1 | | | | | | | | |
| 481191030 | | | | | | | | | | | | | | 1 | | | | | | 2 | | 4 | | | | | | |

| | | | | | | | | | | | | | | | | | | | | |
|---|---|---|---|---|---|---|---|---|---|---|---|---|---|---|---|---|---|---|---|---|
| 481199594 | | | | | | | | | | | 1 | | | 1 | | | | | | |
| 481199836 | | | | | 9 | 17 | 1 | 1 | | | 1 | | | | | | | | | |
| 481204297 | | | | | | | | | | | | 1 | | | | | | | | |
| 481337148 | | | | | | 1 | | | | | | | | | | | | | | |
| 481359062 | 3 | | | | 1 | 1 | | | | | | | | | 22 | 21 | | | | |
| 481390070 | | | | | | | | | | | | | | | | | | 1 | | |
| 481394217 | | 4 | 1 | 3 | | 1 | 8 | 6 | | | | | | | | | | 1 | | |
| 481406868 | | | | | 1 | 5 | 2 | 2 | | | | | | | | | | 4 | 3 | |
| 481410841 | | | | | | | | | | | | | | | | | 4 | 1 | 1 | |
| 481427934 | | | | | | | | 2 | | 4 | 9 | 4 | 4 | 5 | | | | 1 | 1 | |
| 481501073 | | | | | | | | | | | | 1 | 2 | 3 | | | | | | |
| 481501144 | | 1 | | | | | | | | | 2 | 1 | | | | | | | 1 | |
| 481532281 | | | | | | | 6 | 4 | | 2 | | 2 | | | | | | | | |
| 481747302 | | | | | 8 | 25 | | | | | | | | | | | 13 | 7 | 11 | |
| 481770433 | | | | | | | | | | 1 | | 1 | | 1 | | | | | 1 | |
| 481837842 | | | | | | 4 | 2 | 1 | | | 1 | | | | | | | | | |
| 481837848 | | | | | | | | | | | | 2 | | | | | | | | |
| 481838017 | | | | | | | | | | 1 | | | | | | | | | | |
| 481842257 | | | | | | | | | | 1 | | | 2 | | | | | | | |
| 481842486 | | | | | | 1 | 1 | | | | | | | | | | | | | |
| 481846454 | | | | | | | 1 | 2 | | 2 | 7 | 2 | 3 | 6 | | | | | | |
| 482071463 | | 1 | | | 1 | | | | | | | | | | | | | | | |
| 482174774 | | | | | | | 2 | | | | | | | | | | | | | |
| 482178967 | | | | | | | 1 | | | | | | | 1 | | | | | | |
| 482179179 | | | | | | | | | | 2 | 1 | 1 | 1 | | | | | | | |
| 482183354 | | | | | 1 | | | | | | | 1 | | | | | | | | 1 |
| 482183384 | | | | | | | 1 | 3 | | | | | | | | 1 | | | | |
| 482511406 | | | | | | | 3 | 5 | | 7 | 13 | 24 | 13 | 20 | | 1 | | | | |
| 482520165 | | | | | | | | | | 4 | 1 | 1 | 2 | 3 | | | | | | |
| 482520823 | | | | | | | | 1 | | | | | | | | | | | | |
| 482520889 | | 1 | | | | | | | | | | | | | | | | | | |
| 482524464 | | | | | 4 | 1 | 2 | | | | | | | | | | | | | 18 |
| 482754144 | | | | | | | | 1 | | | | | | | 2 | 2 | 2 | | | |
| 482783862 | | | 1 | 1 | | 3 | | 4 | | | | | | | | | | | | |
| 482805831 | | | | | | | | | | 1 | | | | | | | | | | |
| 482852299 | | | | | | | 1 | 2 | | 1 | | 1 | 1 | | | | | | | |
| 482852401 | | | | | | | | | | | | | | | 1 | 15 | | | | |
| 482856632 | | | | | | | | | | 1 | | 1 | 3 | 3 | | | | | | |
| 482856872 | | | | | | | | | | | | | | 2 | | | | | | |
| 482861459 | | | | | 1 | | 6 | | | | | | | | | | | | | |
| 482869559 | | | 1 | | | | | | | | | | | | | | | | | |
| 483142996 | | | | | | 1 | | 1 | | | | | | | | | | | | |
| 483189268 | | | | | | | | | | | 1 | | | | 1 | | | 1 | | |
| 483193732 | | | | | 20 | 14 | 3 | | | | | | | | | | | | | |
| 483198033 | | | 1 | | | | | | | | | | | | | | | | | |
| 483202128 | | 1 | | | | | | | | | | | | | | | | | | |
| 483206709 | | | | | 1 | 5 | 7 | 5 | | | | | | | | | | | | |
| 483206849 | | | | | | | | 1 | | 1 | 2 | 5 | | 1 | | | | | | |
| 483337285 | | | | | | | | | | | | | | | | 1 | | | | |
| 483457318 | | | 2 | | | | | | | | | | | | | | | | | |
| 483539538 | | | 1 | | | | | | | | | | | | | | | | | |
| 483547827 | | | 1 | | | | | | | | | | | | | | | | | |
| 483646991 | | | | | | | | | | | | | | | | | 1 | | | |
| 483879756 | | 2 | | | | | | | | | | | | | | | | | | |
| 483880436 | | | | | | | | | | 1 | | | | | | | 1 | | | |
| 484126552 | | | | | | | | | | | | | | | | | 11 | | | |
| 484213034 | | | | | 5 | 4 | | | | | | | | | | | | | | |
| 484217280 | | | 2 | | | | 1 | | | | | | | | | | | | | |
| 484350664 | | | | | | | | | 1 | | | | | | | | | | | |
| 484355328 | | | | | 4 | 1 | | | | | | | | | | | | | | |
| 484501839 | | | | | | | 10 | 6 | | | | | | | | | | | | |
| 484553282 | | | | | | | | | | 2 | | | | | | | | | | |
| 484554412 | | | | | 6 | 3 | 1 | | | | | | | | | | | | | 6 |
| 484562428 | | | | | 2 | 2 | 1 | 1 | | | | | | | | | | | | |
| 484566597 | | | | | | | 4 | 2 | | | 1 | | 3 | 5 | | | | | | |
| 484829654 | | | | | | | 1 | | | | | | | | | | 11 | 1 | 1 | 50 |
| 484838760 | | | | | 2 | 1 | 2 | 8 | | | | | | | | | | | | |
| 484890037 | | 1 | 1 | | 1 | 2 | 25 | 10 | | 2 | | | | 1 | | | | | | 1 |
| 484898739 | | | | | | | | | | | 1 | | | | | | | | | |
| 484903376 | | | 1 | | | | | | | | | | | | | | | | | |
| 484911881 | | | | | | 1 | | | | | | | | | | | | | | 2 |
| 484911954 | | | | | | 1 | | | | | | | | | | | | | | |
| 484942773 | | | | | 3 | 1 | | | | | | | | | | | | | | |
| 485171425 | | | | | | | | | | | | | | | | | 17 | 3 | 2 | 44 |
| 485231414 | | | | | | | | | | | | | | | | | | | | 1 |
| 485249150 | | | | | | 1 | | | | | | | | | | | | | | 11 |
| 485252796 | | | | | | 1 | | | | | | | | | | | | | | |
| 485499217 | | | | | | | | | | | | | | | | | 2 | | | |
| 485511772 | | 5 | 2 | 1 | | | 1 | 1 | | | | | | | 1 | 2 | | | 1 | |
| 485568100 | | 2 | 1 | | | | | | | | | | | | | | | | | |

| | | | | | | | | | | | | | | | | | | | | | | | | | | | | |
|---|---|---|---|---|---|---|---|---|---|---|---|---|---|---|---|---|---|---|---|---|---|---|---|---|---|---|---|---|
| 485572658 | | | | | | | | | 2 | 2 | | | | | | | | | | | | | | | | | | |
| 485576944 | | | | | | | | | | | | | | | | | | | | 3 | | | | | | | | |
| 485580811 | | | | | | | | | 2 | | | | | | | | | | | | | | | | | | | |
| 485589846 | | | | | | | | | | | | | | | | | | | | | | | | | | | | 1 |
| 485628489 | | | | | | | | | | | | | | | 1 | | | | | | | | | | | | | 1 |
| 485745069 | | | | | | | | | | | | | 31 | | | | | | | | | | | | | | | |
| 485930943 | | | | | | | | | | | | 3 | 3 | | | | | | | | | | | | | | | |
| 485930996 | | | | | | | | | | | | | | | | | | | | | | | | | | | | 1 |
| 485934965 | | | | | | | | | | | | 12 | 7 | | | | | | | | | | | | | 1 | | 1 |
| 486103577 | | | | | | | | | 5 | 2 | | | | | | | | | | | | | | | | | | |
| 486255488 | | | | | | | | | 9 | 2 | 4 | | | | | | | | | | | | | | | | | |
| 486646991 | | | | | | | | | | | | 8 | 5 | 8 | 7 | | | 1 | | 2 | | 3 | | | | | | |
| 486842208 | | | | | | | | | 1 | | | | | | | | | | | | | | | | | | | |
| 486867877 | | | | | | | | | | | | | | | | | | | | | | | | | 1 | | | |
| 486876782 | | | | | | | | | | | | 1 | | | | | | | | | | | | | 1 | | 1 | |
| 486932201 | | | | | | | | | | | | | | | | | | | | | | | | | 4 | | | 30 |
| 486941224 | | | | | | | | | | 2 | | | | | | | | | | | | | | | | | | |
| 486945479 | | | | | | | | | | | | | | | | | | | | | | | | | | | | 1 |
| 487212916 | | | | | | | | | | | | | | | | | | | | | | | | | 1 | | | |
| 487290306 | | | | | | | | | | | | | | | | | | | | | | | | | 5 | | | |
| 487541595 | | | | | | | | | | | | | | | | | | | | | | | | | 1 | | | |
| 487636204 | | | | | | | | | 4 | 5 | | | | | | | | | | | | | | | | | | |
| 487887067 | | | | | | | | | | | | | | | | | | | | | | | | | 7 | | | |
| 487899426 | | | | | | | | | | | 1 | | | | | | | | | | | | | | | | | |
| 488231939 | | | | | | | | | | | | | | | | | | | | | | | | | | | 2 | |
| 488572496 | | | | | | | | | | | | | 2 | 4 | 1 | | | | | | | | | | 3 | 1 | | |
| 509626405 | | 1 | 1 | | | | | | | | | | | | | | | | | | | | | | | | | |
| 509626609 | | 7 | | | | | | | | | | | | | | | | | | | | | | | | | | |
| 509631101 | | | | | | | | | | | | | | 4 | 10 | | | 7 | 7 | 11 | 4 | 6 | 1 | 4 | | | | |
| 509643775 | | | | | | | | | | | | | | 1 | 17 | 3 | 5 | 1 | | 1 | | | 5 | | 8 | 5 | 1 | |
| 509945954 | | | | | | | | | | | | | 3 | | | | | | | | | | | | | | | |
| 509968249 | | | | | | | | | | | | | | | | | | | | | | | | | 1 | | | |
| 509993739 | | | 2 | | | | | | | | | | | | 1 | | | | | | | | 1 | 2 | | | | |
| 510602264 | | | | | | | | | | | | | | | 1 | | 1 | | | | | | | | | | | |
| 510676580 | | | | | | | | | | | | | | | | | | | | | | | 3 | | | | | |
| 510680365 | | | | | | | | | | | | | | | | 5 | 12 | | | | | | | | 1 | | | |
| 510680957 | | | | | | | | | | 4 | 6 | | | 2 | 1 | | | | | | | | | | | | | |
| 510987433 | | | | | | | | | | | | | | | | | | | | | | | | | 1 | | | |
| 511051477 | | | | | 1 | | 2 | | | | | | | 37 | | 55 | 63 | 25 | 25 | 33 | 22 | 25 | 2 | 4 | | 1 | | |
| 511056466 | | | | | | | | | | | | | | | 2 | | | | | | | | | | | 3 | 2 | |
| 511068685 | | | | | | | | | | | | | | | | | | | | | | | | | | | 1 | |
| 511219394 | | | | | | | | | | | | | | | | | | | 1 | | | | | | | | | |
| 511219601 | | | | | | | | | | | | | | | 1 | | | 3 | 3 | 1 | 1 | 6 | | | | | | |
| 511353266 | | | | | | | | | | | | | 2 | | | | | | | | | | 6 | 4 | | | | |
| 511358390 | | | | | | | | | | | | | | | | | | | | | | | | | | | 1 | |
| 511422245 | | | | | | | | | | | | | | | | | | | | 1 | | | | | | 1 | | |
| 511543135 | | | | | | | | | | | | | | | | | | | | 1 | | | | | | | | |
| 511547921 | | | | | | | | | | | | | | 5 | 1 | | | 1 | 5 | 1 | | 5 | | | | | | |
| 511556310 | | | | | | | | | | | | | | 1 | | | | | | 1 | | | | | | | | |
| 511561200 | | | | | | | | | | | | | | | | | | | | 2 | | | | | | | | |
| 511681569 | 1 | | | | | | | | | | | | | 1 | | | | | | | | | | | | | | |
| 511685969 | | | | | | | | | | | | 1 | | | | | | | | | | | | | | | | |
| 511686666 | | | | | | | | | | | | | | 1 | | | | | | | | | | | | | | |
| 511699812 | 4 | | 5 | 1 | | | | | | | | | | | | | | | | | | | | | | | | |
| 511733723 | | | | | | | | | | | | | | | | | | | | | | | 2 | | | | | |
| 511772101 | | | | | | | | | | | | 4 | | 12 | 7 | | | 4 | 7 | 11 | 8 | 5 | | | | 4 | 1 | |
| 511862444 | | | | | | | | | | | | | | | | | | 3 | 3 | | | 1 | | | | | | |
| 511875377 | | | | | | | | | | | | | | | | | | 1 | | | 1 | | | | | | | |
| 511875414 | | | | | | | | | | | | | | | | | | 1 | | | | | | | | | | |
| 511875446 | | | | | | | | | | | | | | | | | | 1 | | | | | | | | | | |
| 511879849 | | | | | | | | | | | | | | | | | | 1 | | | | | | | | | | |
| 511889371 | | | | | | | | | | | | | | 1 | 1 | | | | | | | | | | | | | |
| 511901988 | | | | | | | | | | | | | | 1 | | | | | | | | | | | | | | |
| 512014262 | | | | | | | | | | | | 8 | 15 | 2 | 2 | | | | | | | | | | | | | |
| 512048416 | | | | | | | | | | | | | | | | | | | | | | | 2 | | 4 | | | |
| 512079523 | | | | | | | | | 2 | 2 | 1 | | | | 2 | | | | | | | | | 1 | | | | |
| 512100489 | | | | | | | | | | | | | | | | | | | | | | | | | | 2 | 1 | |
| 512122154 | | | | | | | | | | | | | | | | | | 2 | 3 | | | | | | | | | |
| 512122177 | | | | | | | | | | | | | | 1 | | | | | 1 | 4 | | 3 | | | | | | |
| 512207845 | | | | | | | | | | | | | | | | | | | | | | 3 | | | | | | |
| 512207847 | | | | | | | | | | | | | | 2 | 3 | | | | 3 | 3 | | 3 | | | | | | |
| 512207870 | | | | | | | | | | | | | | 5 | 2 | | | 1 | | | | | | | | | | |
| 512212113 | | | | | | | | | | | | | | | | | | | | 1 | | | | | | | | |
| 512212128 | | | | | | | | | | | | | | 1 | | | | | | 1 | | | | | | | | |
| 512216447 | | | | | | | | | | | | | 1 | | | | | | | 1 | | | | | | | | |
| 512216460 | | | | | | | | | | | | | 1 | | | | | | | 1 | | | | | | | | |
| 512216491 | | | | | | | | | | | | | | 9 | 4 | | | 4 | 2 | 1 | 1 | 3 | | | | | | |
| 512225306 | | | | | | | | | | | | | | | | | | | | 1 | | | | | | | | |
| 512225656 | | | | | | | | | | | | | | | | | | | | 1 | | | | | | | | |
| 512225867 | | | | | | | | | | | | | | 6 | 2 | | | 1 | 1 | 1 | 6 | 1 | | | | | | |

| | | | | | | | | | | | | | | | | | | | | | |
|---|---|---|---|---|---|---|---|---|---|---|---|---|---|---|---|---|---|---|---|---|---|
| 512243445 | | | | | | | | | | | | | 1 | | | | | | | | |
| 512377806 | | | | | | | 3 | | | | | | | | | | | | 1 | | |
| 512411244 | | | | | | | | | | | | | | | | | | | 2 | | |
| 512411317 | | | | | | | | | | | | | | | | | | 1 | | | |
| 512441111 | | | | | 10 | 22 | 1 | 1 | | | | | | | | | | 18 | 2 | 2 | |
| 512441442 | | | | | 2 | 1 | | | | | | | | | | | | 34 | 16 | 33 | |
| 512535936 | | | | | 1 | | | | | | | | | | | | | | | | |
| 512536036 | | | | | | | | | | | | | 1 | 2 | 2 | | | | | | |
| 512536060 | | | | | | | | | | | | 1 | | | | | | | | | |
| 512536126 | | | | | | | 1 | 4 | | | | 1 | 1 | | | | | | | | |
| 512557531 | | | | | | 1 | | | | | | | | | 1 | | | | | | 2 |
| 512566365 | | | | | | | | | | | 2 | | | | | | | | | | |
| 512566853 | | | | | | | 1 | 1 | | | | | | | | | | | | | |
| 512566925 | | | | | | | 1 | | | | | | | | | | | | | | |
| 512774096 | | | | | 2 | | 1 | | | | | | | | | | | | 1 | | |
| 512782759 | | | | | | | 1 | | | | | | | | | | 1 | 4 | 10 | 7 | |
| 512787296 | | | | | 10 | 10 | | | | | | | | | | | | 8 | 2 | | |
| 512872629 | | | | | 3 | | 1 | 1 | | | | 1 | 2 | | 3 | | | | | | |
| 512872831 | | | | | 1 | | | 1 | | | | | | | | | | | | | |
| 512873031 | | | | | 2 | | | | | | | | 1 | 1 | | | | | | | |
| 513209746 | | | | | | | | | | | | | | 2 | 2 | | | | | | |
| 513209805 | | | | | | | 2 | | | | | | | 1 | | | | | | | |
| 513214039 | | | | | 1 | | | | | | 1 | | | | | | | | | | |
| 513218590 | | | | | 1 | | | | | | | | | | | | | | | | |
| 513344531 | | | 1 | | 3 | 5 | 1 | | | | | | | | | | | | 1 | | |
| 513464860 | | | | 1 | | | | | | | | | | | | | | | | | |
| 513551146 | | | | | | | | 1 | | | | 1 | | 1 | | | | | | | |
| 513551342 | | 1 | 1 | | 1 | | | | | | | 1 | | | | | | | | | |
| 513555079 | | | | | | | | | | | | | | | 1 | | | | | | 1 |
| 513555436 | | | | | | | | | | | | | 1 | | | | | | 1 | 1 | |
| 513668146 | | | | | 12 | 6 | | | | | | | | | | | | | | | |
| 513788774 | | | | | | | | | | | | | | | | | 1 | 8 | 3 | 1 | |
| 513840103 | | | | | | | | 1 | | | | | | | | | | | | | |
| 513848395 | | | | 2 | | | | | | | | | | | | | | | | | |
| 513887511 | | | | | | | 1 | 1 | | | | | | | | | | | | | |
| 513887940 | | | | | 1 | | 4 | 2 | | | 2 | | | | 1 | | | | | | |
| 513896195 | | | | | 1 | 2 | 2 | | | | | | | | | | | | | | |
| 513900305 | | 1 | | | | | | | | | | | | | | | | | | | |
| 513904588 | | | | | | | | | | | | | | 4 | | | | | | | |
| 514068564 | | | | | | | | | 3 | | | | | | | 1 | 1 | | 4 | 3 | |
| 514219793 | | | | | | | | | | | 2 | | 1 | | | | | | | | |
| 514224681 | | | | | | 1 | 7 | 5 | | | 4 | 7 | 5 | 2 | 2 | | | | | | |
| 514225046 | | | | | | | | | | | | | 1 | 1 | | | | | | | |
| 514271457 | | | | | | | | | | | | | | | | | | | | | 2 |
| 514272175 | | | | | | | | | | | | | | | | 1 | | | | | |
| 514475106 | | | | | | | | | | | | | | | | | | | 3 | 3 | |
| 514522242 | | | | | | | | | | | | | | | | | 2 | | | | |
| 514527219 | | | 1 | | | | 1 | | | | | | | | | | | | | | |
| 514530554 | | | | | | | | | | | | 2 | | | | | | | | | |
| 514561054 | | | | | | | 1 | | | | | | | | | | | | | | |
| 514565269 | | | | | | | | | | | | | | | | | | | | | 2 |
| 514565428 | | | | | | 1 | 7 | | | | | | | | | | | | | | |
| 514569836 | | | | | | | 1 | | | | | | | | | | | | | | |
| 514583142 | | | | | | | 1 | | | | | | | | | | | | | | |
| 514807275 | | | | 1 | | | | | | | | | | | | | | | | | |
| 514842241 | | | | 1 | | | | | | | | | | | | | | | | 1 | |
| 514859045 | 1 | | | | | | 5 | 3 | | | 5 | 5 | 7 | 4 | 8 | 3 | 2 | | | | |
| 514958793 | | | | | | | | | | | | | | | | | | | | | 1 |
| 515247194 | | 1 | 1 | | | | 1 | | | | | | | 1 | | | | | | | |
| 515247907 | | | | | | 1 | | | | | | | | | | | | | | | |
| 515251693 | | | | | 13 | 7 | | | | | | | | | | | | | | | 8 |
| 515385637 | | | | | | | | | 5 | 1 | | | | | 1 | | 1 | | 1 | 1 | |
| 515580115 | | 4 | | | | | | | | | | | | | | | | | | | |
| 515588291 | | | | | 7 | 14 | | | | | | | | | | | | | | | 6 |
| 515852217 | | | | | | | | | | | | | | | | | | 5 | | | |
| 515869245 | | | | | | | | | | | | | | | | | | 1 | | 1 | 1 |
| 515925122 | | | | | | | 2 | | | | 3 | | | | | | | | 1 | | 1 |
| 515947324 | | | | | 2 | 5 | | | | | | | | | | | | | | | 2 |
| 516055524 | | | | | | | | 1 | | | | | | | | | | | | | |
| 516163285 | | | | | | | | | | | | | | | | 2 | | 1 | | | |
| 516214576 | | | | | 5 | 3 | 1 | | | | | | | | | | | 7 | 1 | 1 | |
| 516288489 | | | | | 7 | 8 | | | | | | | | | | | | | | | 17 |
| 516857964 | | | | | | 1 | | | | | | | | | | | | | | | |
| 516875221 | | | | | 6 | 11 | | | | | | | | | | | | 30 | 1 | 2 | |
| 516943720 | | 3 | 1 | | | | | | | | | | | | | | | | | | |
| 516943929 | | | 1 | | | | | | | | | | | | | | | | | | |
| 516965856 | | | | | 7 | 4 | | | | | | | | | | | | | | | 12 |
| 517242832 | | | | | | | | | | | | | | | | | | | 1 | | |
| 517280671 | | | | | | | 1 | | | | | | | | | | | | | | |
| 517358921 | | 3 | | | | | | | | | | | | | | | | | | | |

| | | | | | | | | | | | | | | | | | | | | | | | | | | | | |
|---|---|---|---|---|---|---|---|---|---|---|---|---|---|---|---|---|---|---|---|---|---|---|---|---|---|---|---|---|
| 517647609 | | | | | | | | | 3 | 2 | | | | | | | | | | | | | | | | | | |
| 517652222 | | | | | | | | | | | | 3 | 2 | | | | | | | | | | | | | | | 1 |
| 517682957 | | | | | | | | | | | | | | | | | | | | | | | | | 1 | | | | |
| 517872280 | | | | | | | | | | | | | | | | | | | | | | | | | | 1 | | | |
| 517902689 | | | | | | | | | | | | | | 1 | 1 | | | | | | | | | | | 2 | | 1 | |
| 517910959 | | | | | | | | | | | | | | | | | | | | | | | | | | 5 | | | |
| 517916020 | | | | | | | | | | 2 | | | | | | | | | | | | | | | | | | | |
| 517928417 | | | | | | | | | | | 1 | | | | | | | | | | | | | | | | | | |
| 518576422 | | | | | | | | | | | | 2 | 6 | 10 | 1 | | | | | | | | | | | | | | |
| 518580489 | | | | | | | | | | | | | | | | | | 1 | 1 | | | | | | | | | | |
| 518899771 | | | | | | | | | | | | | | | | | | | | | | | | | | 7 | | | |
| 518917592 | | | | | | | | | | | | | | | | | | | | | | | | | | 3 | | | |
| 518930199 | | | | | | | | | | | | 1 | | | | | | | | | | | | | | | | | |
| 518947623 | | | | | | | | | | | 1 | | | | | | | | | | | | | | | | | | |
| 519275692 | | | | | | | | | | | 1 | | | | | | | | | | | | | | | | | | |
| 519279166 | | | | | | | | | | | | | | | | | | | | | | | | | | 2 | | | |
| 519339923 | | | | | | | | | | 1 | 1 | | | | | | | | | | | | | | | | | | |
| 519591210 | | | | | | | | | | | | | | | | | | | | | | | | | | 1 | | | |
| 519940322 | | | | | | | | | | | | | | | | | | | | | | | | | | 3 | | | |
| 540001084 | | | | | | | | | | | | | | | | | | | | | | | | 1 | | | | | |
| 540325242 | | | | | | | | | | | | 1 | 2 | 12 | 28 | | | 1 | 6 | 6 | 13 | 3 | | 15 | 36 | 1 | 1 | 1 | |
| 540329317 | | 2 | 2 | | | | | | | | | 1 | | 2 | | | | | | 1 | | | | 16 | 10 | | | | |
| 540657374 | | | | | | | | | | | | | | | | 9 | 6 | | | | | | | | | | | | |
| 540662210 | | | | | | | | | | | | | | | | | 2 | | | | | | | | | | | | |
| 540674783 | | | | 2 | | | | | | | | | | 2 | 2 | | 1 | | | | | | | 1 | 1 | | 1 | | |
| 540679089 | | | | | | | | | | | | | 3 | 2 | 10 | 16 | 25 | | | | | | | 1 | | | | | |
| 540998783 | | | | | | | | | | | | | | | | 12 | 8 | | | | | | | | | | | 2 | |
| 540998882 | 13 | 18 | 13 | 13 | | | | | | | | 3 | 2 | | 1 | 35 | 29 | | | | | | | 1 | 1 | | 3 | | |
| 541007502 | | | | | | | | | 1 | | | | | | | 1 | 1 | | | | | | | | | | | | |
| 541011631 | | | | | | | | | | | | 1 | | | | | | | | | | | | | | | | | |
| 541373868 | | | | | | | | | | | | | | | 1 | | | | | | | | | | | 1 | | | |
| 541378802 | 5 | 6 | 9 | 7 | | | | | | | | | 1 | 5 | 4 | 17 | 19 | | | | | | | 1 | | | 1 | | |
| 541663112 | | | | | | | | | | | | | | | | | 1 | | | | | | | | | | | | |
| 541681271 | | | | | | | | | | | | | | 1 | | | | | | | | | | | | 1 | | | |
| 541697718 | | | | | | | | | | | | 2 | 1 | 3 | 2 | | | | | | | | | | | | | | |
| 541723731 | | | | | | | | | | | | 1 | | | | | | | | | | | | | | | | | |
| 541732518 | | | | | | | | | | | | | | | 1 | | | | | | | 3 | | | | | | | |
| 541741406 | | | | | | | | | | | | | | | | | | 1 | | 3 | | | | | | | | | |
| 541913280 | | | | | | | | | | | | 1 | | | | | | | | | | | | | | | | | 1 |
| 542056766 | 1 | 3 | | 1 | | | | | | | | | | 4 | 1 | 2 | 5 | | | | | | | | | | | | |
| 542103594 | | | | | | | | | | | | | | | | | | | | | | | | | 4 | | | | |
| 542103645 | | | | | | | | | | | | | | | | | | 3 | | | | | | | | | | | |
| 542104127 | | | | | | | | | | | | | | | | | | | 1 | | | | | | | | 2 | 1 | |
| 542111941 | | | | | | | | | | | | | | | | | | | | | | | | 2 | | | | | |
| 542410481 | | | | | | | | | | | | 6 | 1 | | | | | | | | | | | | | | | | |
| 542423139 | | | | | | | | | | | | | | | | | | | | | | | | 1 | | | | | |
| 542457618 | | | | | | | | | | | | | | | 1 | | | | | | | | | 1 | | 4 | | | |
| 542573741 | | | | | | | | | | | | | | | 1 | | | 1 | 1 | 2 | 1 | | | | | | | | |
| 542578154 | | | | | | | | | | | | | | | | | | 1 | | | | | | | | | | | |
| 542691005 | | | | | | | | | | | | | 2 | | | | | | | | | | | | | 1 | | 2 | |
| 542734736 | | | | | | | | | | | | | | | | | | | | | | | | | | 1 | | | |
| 542747457 | | | | | | | | | | | | | | 1 | | | | | | | | | | 7 | 5 | | | | |
| 542751938 | | | | | | | | | | | | | | | | | | | | | | | | 1 | | | | | |
| 542781165 | | | | | | | | | | | | | | | | | | | 2 | | | | | | | | | | |
| 542798778 | | | | | | | | | | | | | | | | | | | | | 1 | | | | | | | | |
| 542902307 | | | | | | | | | | | | | | | | | | 1 | | 2 | 1 | 1 | | | | | | | 1 |
| 542906401 | | | | | | | | | | | | | | | | | | 1 | | | | | | | | | | | |
| 542906584 | | | | | | | | | | | | | | | | | | | 1 | | | 1 | | | | | | | |
| 542910915 | | | | | | | | | | | | | | 2 | 1 | | | 1 | 2 | | | 2 | | | | | | | |
| 542910924 | | | | | | | | | | | | | | 1 | | | | | 4 | 2 | | | | | | | | | |
| 542923445 | | | | | | | | | | | | | | 1 | | | | | | | | | | | | | | | |
| 543010474 | | | | | | | | | | | | | | | | | | | | | | | | 1 | 2 | | | | |
| 543023799 | | | | | | | | | | | | | 2 | | | | | | | | | | | | | | | | |
| 543049093 | | | | | | | | | | | | 4 | 9 | | | | | | | | | | | | | | | | |
| 543062097 | | | | | | | | | | | | | | 1 | | 7 | 8 | | | | | | | | | 1 | | | |
| 543092817 | | | | | | | | | | | | 13 | 5 | | | | | | | | | | | | | 1 | | | |
| 543135607 | | | | | | | | | | | | | | | | | | | | | | | | | 1 | 6 | 6 | 8 | |
| 543234229 | | | | | | | | | | | | | | | 2 | | | 3 | 3 | 1 | 2 | 5 | | | | | | | |
| 543234427 | | | | | | | | | | | | | | | | | | | | 2 | | | | | | | | | |
| 543238468 | | | | | | | | | | | | | | | | | | | | | | | | | | | | | 2 |
| 543238882 | | | | | | | | | | | | | | | | | | | | | | 1 | | | | | | | |
| 543243444 | | | | | | | | | | | | | | | | | | | | | | 1 | | | | | | | |
| 543247309 | | | | | | | | | | | | | | | 2 | | | | | | | | | | | | | | |
| 543247387 | | | | | | | | | | | | | | 6 | | | | | 2 | | | | | | | | | | |
| 543247616 | | | | | | | | | | | | | | | | | | | 3 | 1 | | | | | | | | | |
| 543247668 | | | | | | | | | | | | | | | | | | | 1 | | | | | | | | | | |
| 543247684 | | | | | | | | | | | | | | | | | | | 1 | | 1 | | | 3 | 3 | | | | |
| 543247857 | | | | | | | | | | | | | 2 | | | | | | | | 1 | | | | | | | | |
| 543252056 | | | | | | | | | | | | | | | | | | | | | 2 | 1 | | | | | | | |
| 543255853 | | | | | | | | | | | | | | | | | | | 2 | | 2 | | | | | | | | |

| | | | | | | | | | | | | | | | | | | | | | | | | | | | | |
|---|---|---|---|---|---|---|---|---|---|---|---|---|---|---|---|---|---|---|---|---|---|---|---|---|---|---|---|---|
| 543256347 | | | | | | | | | | | | 6 | 2 | | 4 | | | 2 | | | 1 | | | | | | | 3 |
| 543365167 | 2 | 1 | | | | | | | | | | 2 | | 2 | 15 | 11 | | 1 | | 2 | 1 | 2 | | | | 1 | |
| 543498242 | | | | | | | | | | | | | | | | | | | | | | | | | 1 | | |
| 543566744 | | | | | | | | | | | | | | | | | | | | | 1 | 6 | 10 | 1 | 2 | 1 | |
| 543566772 | | | | | | | | | | | | 3 | | 6 | 4 | | | 1 | | 2 | | | | | | | | |
| 543575336 | | | | | | | | | | | | | | | | | | | | 1 | | | | | | | | |
| 543579922 | | | | | | | | | | | | | | | | | | | | 1 | | | | | | | | |
| 543580198 | | | | | | | | | | | | 4 | 2 | | | | | | | | | | | | | | | |
| 543580245 | | | | | | | | | | | | | | | | | | | | 1 | | | | | | | | |
| 543583771 | | | | | | | | | | | | | | | | | | | 1 | | | | | | | | | |
| 543584240 | | | | | | | | | | | | | | 1 | | | | | | | | 1 | | | | | | |
| 543584523 | | | | | | | | | | | | | | | | | | 2 | 1 | | 4 | 2 | | | | | | |
| 543584751 | | | | | | | | | | | | | | | | | | | | 2 | | | | | | | | |
| 543588162 | | | | | | | | | | | | | | 1 | | | | | | | | | | | | | | |
| 543588915 | | | | | | | | | | 2 | | | | | | | | | | | | | | | | | | |
| 543589080 | | | | | | | | | | | | | | | 1 | | | 2 | | | 2 | | | | | | | |
| 543692985 | | | | | | | | | | | | | | | | 1 | | | | | | | | | | | | |
| 543702186 | | | | | | | | 3 | | | | | | | | | | | | | | | | | | | | |
| 543718301 | | | | | | | | | | | | | | | | | | | | | | | 1 | 1 | | | | |
| 543808842 | | | | | | | | | | | | 2 | | 6 | 3 | | | | | | | | | | | | | |
| 543821838 | | | | | | | | | | | | | 2 | | | | | | | | | | | 1 | | | | |
| 543907986 | | | | | | | | | | | | | | | | | | | 2 | | 1 | | | | | | | |
| 543912097 | | | | | | | | | | | | | | | | | | 1 | | | 1 | | | | | | | |
| 543916401 | | | | | | | | | | 3 | | | | 4 | | | | 4 | 5 | 4 | | 2 | | | | | | |
| 543916894 | | | | | | | | | | | | | | | | | | | | 1 | | | | | | | | |
| 543925435 | | | | | | | | | | | | 1 | 1 | | | | | | | | | | | | | | | |
| 543929121 | | | | | | | | | | | | 1 | | 5 | 3 | | | 2 | | | 1 | | | | | | | |
| 543934592 | | | | | | | | | | | | | | | | | | | | | | | | | | | | 4 |
| 544038123 | | | | | | | | | | | | | | | | | | | | | | | | | | 1 | |
| 544145565 | | | | | | | | | | | | | | | 1 | | | | | | | | | | | | | |
| 544158132 | | | | | | | | | | | | | | | | | | | | | | 1 | | 1 | | | | |
| 544158940 | | | | | | | | | | | | | | | | | | | | | | | | 28 | 3 | | |
| 544171197 | | | | | | | | | | | 1 | | | | | | | | | | | | | | | | | |
| 544184932 | | | | | | | | | | | | | | | | | | | | | | | | | 2 | | |
| 544240309 | | | | | | | | | | | | | | | 1 | | | 3 | 4 | 5 | | 1 | | | | | | |
| 544249066 | | | | | | | | | 3 | 1 | | 1 | | | | | | | | | | | | | | | | |
| 544262349 | | | | | | | | | | | | 19 | 17 | 1 | | | | | | | | | | | | | | 3 |
| 544265891 | | | | | | | | | | | | 4 | 6 | 2 | | | | 1 | | | | | | | | | | 2 |
| 544265926 | | | | | | | | | | | | | | | | | | | | | | | | | | | | 1 |
| 544265948 | | | | | | | | | | | | | | | | | | | | | | | | | | | | 1 |
| 544266001 | | | | | | | | | | | | | | | | | | | | | | | | | | | | 3 |
| 544266246 | | | | | | | | | | | | 1 | | | | | | | | | | | | | | | | 2 |
| 544270890 | | | | | | | | | | | | | | | | | | | | | | | | | | | | 14 |
| 544270910 | | | | | | | | | | | | 3 | | 1 | | | | | | | | | | | | | | |
| 544274492 | | | | | | | | | | | | | | | | | | 1 | | | | | | | | | | |
| 544512309 | | | | | | | | | | | | | | | | | | | | | | | 2 | | | | | |
| 544534612 | | | | | | | | | | | | | | 3 | | | | | | 2 | | | | | | | | |
| 544594211 | | | | | | | | | | | | | | 1 | | | | | | | | | | | | | | |
| 544599100 | | | | | | | | | | | | | 1 | 2 | | | | | | | | | | | | | | |
| 544602598 | | | | | | | | | 2 | 10 | | | | | | | | | | | | | | | | | | |
| 544602710 | | | | | | | | | | 1 | | | | | | | | | | | | | | | | | | |
| 544606883 | | | | | | | | | | | | | | | | | | | | | | | | | | | | 1 |
| 544607508 | | | | | | | | | | | | | 1 | | | | | | | | | | | | | | | |
| 544611593 | | | | | | | | | | | | 2 | 2 | | 1 | | | | | | | | | | | | | 1 |
| 544611771 | | | | | | | | | | | | 1 | | | | | | | | | | | | | | | | |
| 544611801 | | | | | | | | | | | | | | | | | | | | | | | | | | | | 4 |
| 544615880 | | | | | | | | | | | | | | 5 | | | | | | | | | | | | | | |
| 544620289 | | | | | | | | | | | | 3 | 2 | 2 | 1 | | | | | | | | | | | | | |
| 544831533 | | | | | | | | | | | | | | 1 | | | | | | | | | | | | | | |
| 544840037 | | | | | | | | | | | 3 | | | | | | | | | | | | | | | | | |
| 544857968 | | | | | | | | | | | 1 | | | | | | | | | | | | | | | | | |
| 544918228 | | | | | | | | | | | | | | | | | | | 1 | | | | | | | | | |
| 544918416 | | | | | | | | | | | | | | | | | | 3 | | | | | | | | | | |
| 544931723 | | | | | | | | | | 1 | | | | | | | | | | | | | | | | | | |
| 544939669 | | | | | | | | | | 1 | | | | | | | | 1 | | | | | | | | | | |
| 544939871 | | | | | | | | | | | | 2 | 1 | | | | | | | | | | | | | | | |
| 544943610 | | | | | | | | | | 5 | | | | | | | | | | | | | | | | | | |
| 544948071 | | | | | | | | | | | | | 4 | | | | | | | | | | | | | | | |
| 544952587 | | | | | | | | | | | | | | | | | | | | | | | | | | | | 4 |
| 544953224 | | | | | | | | | | | | | | | | | | | | | | | | | | | | 2 |
| 545254708 | | | | | | | | | | | | | | 1 | 4 | | | | | | | | | | | | | |
| 545259664 | | | | | | | | | | | | | | 1 | | | | | | | | | | | | | | |
| 545263525 | | | | | | | | | | | | 1 | | | | | | | | | | | | | | | | |
| 545263627 | | | | | | | | | | | | 1 | | | | | | | | | | | | | | | | |
| 545280478 | | | | | | | | | 1 | 1 | | | 1 | | | | | | | | | | | | | | | |
| 545289772 | | | | | | | | | | | | | 2 | | | | | | | | | | | | | | | |
| 545293434 | | | | | | | | | | | | | | | | | | | | | | | | | | | | 5 |
| 545444996 | | | | | | | | | | | | | | | | | | | | | | | | 1 | | | | |
| 545501062 | | | | | | | | | | | | | 1 | | | | | | 2 | 2 | | | 5 | 4 | | | 2 | |
| 545608742 | | | | | | | | | 2 | | | | | | | | | | | | | | | | | | | |

| | | | | | | | | | | | | | | | | | | |
|---|---|---|---|---|---|---|---|---|---|---|---|---|---|---|---|---|---|---|
| 545612931 | 2 | | | | | | | | | | | | | | | | | |
| 545617845 | | 1 | | | | | | | | | | | | | | | | |
| 545621581 | | | | | 1 | | | | | | | | | | | | | |
| 545621597 | | 2 | | | | | | 1 | | | 1 | | | | | | | |
| 545621779 | | | | | 1 | | | | | | | | | | | | | |
| 545625956 | | 2 | | | | | | | | | | | | | | | | |
| 545629975 | | 2 | | | | | | | | | | | | | | | | |
| 545630021 | | | | 2 | | | | | | | | | | | | | | |
| 545748103 | | | | 4 | 1 | | | | | | | | | | | | | |
| 545872467 | | | | | | | | | | | | | | | 1 | | | |
| 545903200 | | | | 1 | | | | | | | | | | | | 12 | 16 | |
| 545906392 | | | | | | | | | | | | | 1 | | | | | |
| 545907195 | | | | | 1 | | | | | | | | | | | 4 | 3 | |
| 545933292 | | | | | | | | 1 | | 1 | | | | | | | | |
| 545941764 | | | | | | | | | | | | | | | | | | 1 |
| 545962610 | | | | | 1 | | | | | | | | | | | | | |
| 545966742 | | | | 1 | 2 | | | | | | | | | | | | | 1 |
| 545966857 | | | | | 1 | | | | | | | | | | | | | |
| 545967081 | | | | 4 | 3 | | | | | | | | | | | | | |
| 545971070 | | | | | 1 | | | | | | | | | | | | | |
| 545971266 | | | | | | | | | | | | | | | | | | 1 |
| 545971650 | | | | | 1 | | | | | | | | | | | | | |
| 545975516 | | | | | | | | | | | | | | | | | | 1 |
| 545979714 | | | | | | | | | | | | | | | | | | 1 |
| 545984286 | | | | | 5 | | | | | | | | | | | | | |
| 546217818 | | | | 1 | 2 | | | | | | | | | | 49 | | | |
| 546234932 | | | | | | 4 | 2 | 1 | | | | | | | | | | |
| 546287086 | | | | 6 | 5 | 3 | 1 | | | | | | | | | | | 1 |
| 546312139 | | | | | | | | | | | | 1 | | | | | | 1 |
| 546312597 | | | | | | | | | | | | | | | | | | 4 |
| 546316510 | | | | | | | | | | | | | | | | | | 6 |
| 546316520 | | 1 | | | | | | | | | | | | | | | | |
| 546316765 | 1 | 1 | | | | | | | | | | | | | | | | |
| 546559025 | | | | | | | | | | | | | | | 1 | | | |
| 546640628 | | | | | | | | | | | | | | | | | | 1 |
| 546641024 | | 2 | | | | | | | | | | | | | | | | |
| 546641044 | | | | | | | | | | | | | | | 1 | | | |
| 546645399 | | 1 | | 2 | | | | | | | | | | | | | | |
| 546653091 | | | | | | | | | | | | | | | | | | 2 |
| 546653603 | | | | | 1 | | | | | | | | | | | | | |
| 546657497 | | 2 | | | | | | | | | | | | | | | | |
| 546917498 | | | | 1 | 1 | | | | | | | | | | | 1 | | |
| 546925626 | 1 | | | | | | | | | | | | | 1 | | | | |
| 546956605 | | | | | 1 | 2 | | | | | | | | | | | | |
| 546981769 | | | | | | | | | | | | | | | | | | 1 |
| 546989946 | | | | | | | | | | | | | | | | | | 3 |
| 547002880 | | | | | 1 | | | | | | | | | | | | | |
| 547240545 | | | | 4 | 5 | | | | | | | | | | | | | |
| 547258283 | | | | | 1 | 3 | | | | | | | 1 | | | 1 | | |
| 547310321 | | | | 8 | | | | | | | | | | | | | | 2 |
| 547318372 | | | | | | | | | | | | | | | | | | 1 |
| 547331124 | | | | | | | | | | | | | | | | | | 2 |
| 547612423 | | 1 | | | | | | | | | | | | | | | | |
| 547659598 | | | | | | | | | | | 1 | | | | | | | |
| 547664568 | | | | | | | | | 1 | | | | | | | | | |
| 547676867 | | | | 1 | 5 | | | | | | | | | | | | | |
| 547918570 | | | | | | | | | | | | | | 1 | 1 | 4 | 3 | |
| 547927681 | | | | 1 | 2 | 1 | 1 | | | | | | | | | | | |
| 547936371 | | | | | | | | | | | | | | | 3 | | | |
| 547978595 | 3 | | | | | | | | 1 | | | | | 2 | | | | |
| 548017378 | | | | | | | | | | | | | | | | | | 1 |
| 548225503 | | | | | | | | | | | | | | | 2 | | | |
| 548272677 | | | | | | | | | | | | | | | 1 | | | |
| 548342008 | | | | 7 | 6 | | | | | | | | | | | | | |
| 548367551 | | | | | | | | | | | | | | | | | | 1 |
| 548367581 | | | | | | | | | | | | | | | | | | 1 |
| 548371852 | | | | 1 | 4 | | | | | | | | | | | | | |
| 548571023 | | | | | | | | | | | | | | | 1 | | | |
| 548678665 | | | | | | | | | | | | | | | | | | 1 |
| 548691397 | | | | | 1 | | | | | | | | | | | | | |
| 548699489 | | | | | | | | | | | | | | | | | | 1 |
| 548704181 | | | | | | | | | | | | | | | | | | 2 |
| 548712377 | | | | | | | | | | | | | | | 25 | | | 1 |
| 548718223 | | | | | | | | | | | | | | | | | | 7 |
| 548718249 | | | | 7 | 3 | 4 | 2 | | | | | | | | | | | |
| 548722401 | | | | | | | | | | | | | | | | | | 6 |
| 548949855 | | | | | | | | | | | | | | | 2 | | | |
| 548972278 | | | 2 | | | | | | | | | | | | | | | |
| 549032565 | | | | | | | | | | | | | | | | | | 1 |
| 549049279 | | | | | | | | | | | | | | | | | | 2 |

| | | | | | | | | | | | | | | | | | | | | | | | | | | | | |
|---|---|---|---|---|---|---|---|---|---|---|---|---|---|---|---|---|---|---|---|---|---|---|---|---|---|---|---|---|
| 549175510 | | | | | | | | | | | | 1 | | | | | | | | | | | | | | | | |
| 549347735 | | | | | | | | | | | | | | | 1 | | | | | | | | | | | | | |
| 549377313 | | | | | | | | | | | | | | 8 | | | | | | | | | | | | | | |
| 549377612 | | | | | | | | | | | | 1 | | 1 | | | | | | | | | | | | | | |
| 549381505 | | | | | | | | | | | | | | | | | | | | | | | | | | | | 2 |
| 549597974 | | | | | | | | | | | | | | | | | | | | | | | | | 2 | | | |
| 549611256 | | | | | | | | | 1 | 1 | 1 | | | | | | | | | | | | | | | | | |
| 549714149 | | | | | | | | | | | | | | | | | | | | | | | | | | | | 9 |
| 549718224 | | | | | | | | | | | | | | | | | | | | | | | | | | | | 8 |
| 549718269 | | | | | | | | | | | | | | | | | | | | | | | | | | | | 2 |
| 549726953 | | | | | | | | | | | | | | | | | | | | | | | | | | | | 2 |
| 549735839 | | | | | | | | | | | | | | | | | | | | | | | | | | | | 1 |
| 550033958 | | | | | | | | | | | 1 | | | | | | | | | | | | | | | | | |
| 550063586 | | | | | | | | | | | | | | | | | | | | | | | | | | | | 13 |
| 550063644 | | | | | | | | | | | | | | | | | | | | | | | | | | | | 1 |
| 550275952 | | | | | | | | | | | | | | | | | | | | | | | | | 2 | | | |
| 550280614 | | | | | | | | | | | | | | | | | | | | | | | | | 6 | | | |
| 550297695 | | | | | | | | | | | | | | 3 | | 1 | 1 | | | | | | 10 | 10 | 2 | 16 | 35 | |
| 550322829 | | | | | | | | | | | 1 | | | | | | | | | | | | | | | | | |
| 550326974 | | | | | | | | | | | | | | | | | | | | | | | | | 3 | 1 | 1 | |
| 550400302 | | | | | | | | | | | | 1 | | | | | | | | | | | | | | | | |
| 550404686 | | | | | | | | | | | | 5 | 2 | | | | | | | | | | | | | | | |
| 550408962 | | | | | | | | | | | | | | | | | | | | | | | | | | | | 6 |
| 550617370 | | | | | | | | | | | | | | | | | | | | | | | | | 1 | | | |
| 550629848 | | | | | | | | | | | | | | | | | | | | | | | | | 2 | | | |
| 550750102 | | | | | | | | | | | | | | | | | | | | | | | | | | | | 2 |
| 550988122 | | | | | | | | | | | | | | | | | | | | | | | | | 1 | | | |
| 551090979 | | | | | | | | | | | | | | | | | | | | | | | | | | | | 4 |
| 551410783 | | | | | | | | | | | | | | | | | | | | | | | | | | | | 2 |
| 551436366 | | | | | | | | | | | | | | | | | | | | | | | | | | | | 1 |
| 571018594 | 5 | 8 | 1 | 3 | | | | | | | | | | | | | | | | | | | | | | | | |
| 571018713 | 5 | 1 | 1 | | | | | | | | | | | | | | | | | | | | | | | | | |
| 571346836 | | | | 1 | | | | | | | | | | 1 | | 2 | | | | | | | 2 | | | | | |
| 571346887 | | | | | | | | | | | 1 | | | | | | | | | | | | | | 2 | | | |
| 571355851 | | | | | | | | | | | | | | 1 | | | | | | | | | | | | | | |
| 571364144 | | | | | | | | | 1 | 4 | 17 | | 2 | | | 1 | | | | | | | 1 | | | | | |
| 571364778 | 3 | | | | | | | | | | | | | | | | | | | | | | | | | | | |
| 571368849 | | | | | | | | | | | | | | | | 7 | 3 | 3 | 1 | 1 | | 2 | 3 | 7 | | | | |
| 571372889 | 6 | 8 | 10 | 8 | | | | | | | | | | 4 | 12 | 19 | 23 | | | | | | | 2 | 1 | | | |
| 571376785 | | | | | | | | | | | | | | | | | | | | | | | 2 | | | | | |
| 571376908 | | | | | | | | | | | | | | | | | | | | | | | 2 | | | | | |
| 571381258 | | | | | | | | | | | | | | | | | | | | | | | | 2 | | | | |
| 571692839 | | | | 1 | | | | | | | | | | | 9 | | | | | | | | 8 | 10 | | | | |
| 571709547 | | | | | | | | | 3 | 6 | 3 | | 1 | 1 | 3 | 1 | | | | 5 | 1 | 2 | 15 | 25 | | | | |
| 571713905 | | | | | | | | | | | 1 | | | 5 | 22 | 34 | 23 | | | | | | 6 | | 12 | 1 | | |
| 571726583 | | | | | | | | | | | | | | | | | | | | | | | 1 | | | | | |
| 572051031 | | | | | | | | | | | | 1 | | | | | | | | | | | | | | | | |
| 572076327 | | | | | | | | | | | | | | | | | | | | | | | 2 | | | | | |
| 572754059 | | | | | | | | | | | | | | 2 | | 1 | | | | | | | | | | | | |
| 572758750 | | | | | | | | | | | | | | | | | | 2 | | 1 | | 1 | | | | | | |
| 572780218 | | | | | | | | | | | | | | | | | | | | | | | 1 | | | | | |
| 573090850 | | | | | | | | | | | | | | | | | 3 | | | | | | 4 | | 1 | | | |
| 573142557 | | 1 | 2 | 2 | | | | | | | | | | 1 | 2 | | | 3 | 2 | 1 | | | 4 | 2 | | 2 | | |
| 573142803 | | | | | | | | | | | | | | 3 | 2 | | | | | 1 | | | | 1 | | | | |
| 573143028 | | | | | | | | | | | | | | | | | | | | | | | 1 | | | | | |
| 573151229 | | | | | | | | | | | | | | | | | | | | | | | 4 | 7 | | | | |
| 573289509 | | | | | | | | | | | | | | | 4 | | | | | | | | | | | | | |
| 573289993 | | | | | | | | | | | | | 1 | | | | | | | | | | | | | | | |
| 573449228 | | | | | | | | | | | | | | | | | | | | | | | 3 | | | | | |
| 573618167 | | | | | | | | | | | | | | | 2 | | | | | | | | | | | | | |
| 573627110 | | | | | | | | | | | | | 1 | | | | | | | | | | | | | | | |
| 573774122 | | | | | | | | | | | | | | | | | 1 | | | | | | | | | | | |
| 573825361 | | | | | | | | | | | | | | | | | | | | | | | | | 3 | | | |
| 573833492 | | | | | | | | | | | | | | 4 | | | | | | | | | | | | | | |
| 573833723 | | | | | | | | | | | | | 3 | | | | | | | | | | | | | | | |
| 573837867 | | | | | | | | | | | | | | | | | | | | | | | | | | 6 | 2 | |
| 573937163 | | | | | | | | | | | | 4 | | | | | | | | | | 2 | | | | | | |
| 574062996 | | | | | | | | | | | | | | 2 | | 1 | | 2 | 4 | 4 | 2 | 1 | 4 | 6 | | | | |
| 574161719 | | | | | | | | | | | | | | | | | | | | | | | | | | 2 | 1 | |
| 574377845 | | | | | | | | | | | | | | | | 2 | | | | | | | | | | | | |
| 574395611 | | | | | | | | | | | | 1 | 1 | 4 | 15 | 25 | 38 | | | | | | 1 | 1 | 1 | 2 | | |
| 574511458 | | | | | | | | | | | | 11 | 13 | 1 | 2 | | | | | | 1 | 1 | | | | | | 2 |
| 574515806 | | | | | | | | | | | | 6 | 1 | 1 | | | | | | | | | 2 | | | | | |
| 574601891 | | | | | | | | | | | | | | 1 | | | | | | | | | | | | | | |
| 574736017 | | | | | | | | | | | | | 1 | | | | | | | | | | | | | | | |
| 574740549 | | | | | | | | | | | | | | 2 | | | | | | | | | | | | | | |
| 574882311 | | | | | | | | | | | | | | | | | | 2 | | 3 | | | | 5 | | | | |
| 574955896 | | | | | | | | | | | | | 2 | | | | | | | | | | | | | | | |
| 574969092 | | | | | | | | | | | | | 1 | | | | | | | | | | | | | | | |
| 574973454 | | | | | | | | | | | | 5 | 4 | 2 | | | | | | | | | | | | | | |

| | | | | | | | | | | | | | | | | | | | | | | | | | | | | |
|---|---|---|---|---|---|---|---|---|---|---|---|---|---|---|---|---|---|---|---|---|---|---|---|---|---|---|---|---|
| 574978101 | | | | | | | | | | | | | | 3 | | | | | | | | | | | | | | |
| 575197482 | | | | | | | | | | 1 | | | | | | | | | | | | | | | | | | |
| 575301459 | | | | | | | | | | | | 1 | | | | | | | | | | | | | | | | |
| 575305924 | | | | | | | | | | | | 2 | 3 | 1 | | | | | | | | | | | | | | 3 |
| 575323308 | | | | | | | | | | | | | | | 1 | | | | | | | | | | | | | |
| 575564352 | | | | | | | | | | | | | | | | | | | | 1 | 1 | | 2 | | | | | |
| 575573640 | | | | | | | | | | | | | | | 1 | | | | | | | | | | | | | |
| 575612047 | | | | | | | | | | | | | | | | | | | | 1 | | | | | | | | |
| 575620465 | | | | | | | | | | | | | | 1 | | | | | | | | | | | | | | |
| 575647041 | | | | | | | | | | | | 1 | | | | | | | | | | | | | | | | |
| 575651309 | | | | | | | | | | | | 6 | 7 | 6 | 1 | | | | | | | | | | | | | |
| 575768294 | | | | | | | | | | | | | | | | 1 | | | | | | | | | | | | |
| 575915136 | | | | | | | | | | | | | | | 2 | | | | | | | | | | | | | |
| 576079106 | | | | | | | | | | | | | | | | 4 | 2 | | | | | | | | | | | |
| 576082257 | | | | | | | | | | | | 9 | 17 | 5 | | | | | | | | | | | | | 1 | |
| 576208238 | | | | | | | | | | | | | | | | | | | | | | | | | | 1 | 1 | |
| 576216182 | | | | | | | | | | | 1 | | | | | | | | | | | | | | | | | |
| 576243061 | | | | | | | | | | | | 1 | 3 | 1 | | | | | | | | | | | | | | |
| 576260070 | | | | | | | | | | | 4 | | | 2 | | | | | | | | | | | | | | |
| 576319940 | | | | | | | | | | 1 | | | | | | | | | | | | | | | | | | |
| 576574889 | | | | | | | | | | | | | 1 | | | | | | | | | | | | | 1 | | |
| 576683512 | | | | | | | | | | | | | 2 | | | | | | | | | | | | | | | |
| 576985092 | | | | | | | | | | | | 1 | | | | | | | | | | | | | | | | |
| 576999054 | | | | | | | | | 2 | | | | | | | | | | | | | | | | | | | |
| 577244511 | | | | | | | | | | | | 4 | 7 | 8 | 2 | | | | | | | | | | 1 | | 1 | |
| 577330676 | | | | | | | | | | | | | | | | | | | | | | | | | | | | 3 |
| 577554935 | | | | | | | | | | | | | | | | | | | | | | | | | | | | 2 |
| 577624125 | | | | | | | | | | | | | | | | | | | | 1 | | | | | | | | |
| 577709776 | | | | | | | | | | | | | | | | | | | | | | | | | | | | 1 |
| 578016879 | | | | | | | | | | | | | | | | | | | | | | | | 2 | | | | |
| 578254032 | | | | | | | | | | | | | | | | | | | | | | | | | | 1 | | |
| 578284233 | | | | | | | | | | | | 1 | 2 | | | | | | | | | | | | 22 | 1 | 1 | |
| 578587259 | | | | | | | | | | | | 3 | 3 | 4 | 1 | | | | | | | | | | | | | |
| 578587420 | | | | | | | | | | | | | | | | | | | | | | | | | 1 | | | |
| 578621125 | | | | | | | | | | | | | 1 | | | | | | | | | | | | | | | |
| 578642656 | | | | | | | | | | 2 | | | | | | | | | | | | | | | | | | |
| 578686468 | | | | | | | | | | | | 3 | 1 | 6 | 2 | | | | | | | | | | | | | |
| 578698726 | | | | | | | | | | | | | | 1 | | | | | | | | | | | | | | |
| 578919867 | | | | | | | | | | | | | | | 2 | | | | | | | | | | | | | |
| 578970891 | | | | | | | | | | | | | | | | | | | | | | | | | 3 | | | |
| 579014562 | | | | | | | | | 2 | | 1 | | | | | | | | | | | | | | | | | |
| 579239039 | | | | | | | | | | | | | | | | | | | | | | | | | | | 1 | |
| 579575594 | | | | | | | | | | | | | | | | | | | | | | | | | 1 | | | |
| 579700707 | | | | | | | | | | 2 | | | | | | | | | | | | | | | | | | |
| 580071186 | | | | | | | | | | | | 1 | | | | | | | | | | | | | | | | |
| 580348303 | | | | | | | | | 2 | | 2 | | | | | | | | | | | | | | | | | |
| 580662960 | | | | | | | | | | | | | | | | | | | | | | | | | 1 | | | |
| 580714587 | | | | | | | | | | | | | | | | | | | | | | | | | 1 | | | |
| 580719155 | | | | | | | | | 4 | 7 | 4 | | | | | | | | | | | | | | | | | |
| 580732172 | | | | | | | | | | | | | | | | | | | | | | | | | | | | 4 |
| 580732377 | | | | | | | | | | | | | | | | | | | | | | | | | | | | 1 |
| 580736603 | | | | | | | | | | | | | | | | | | | | | | | | | | | | 1 |
| 580745387 | | | | | | | | | | | | 1 | | | | | | | | | | | | | 1 | | | |
| 580987056 | | | | | | | | | | | | | | | | | | | | | | | | | 3 | | | |
| 581004609 | | | | | | | | | | | | | | | | | | | | | | | | | 2 | | | |
| 581055513 | | | | | | | | | 6 | 3 | 9 | | | | | | | | | | | | | | | | | |
| 581073524 | | | | | | | | | | | | | | | | | | | | | | | | | | | | 1 |
| 581409749 | | | | | | | | | | | | | | | | | | | | | | | | | | | | 1 |
| 581432310 | | | | | | | | | | | | 9 | 9 | 2 | 1 | | | | | | | | | | | | | |
| 581436615 | | | | | | | | | | | | 1 | | | | | | | | | | | | | | | | |
| 581755530 | | | | | | | | | | 1 | 4 | | | | | | | | | | | | | | | | | |
| 582381047 | | | | | | | | | | | | | | | | | | | | | | | | | 1 | | | |
| 583132200 | | | | | | | | | | | | | | | | | | | | | | | | | | | | 1 |
| 601712799 | | | 2 | | | | | | | | | | | | | | | | | | | | | | | | | |
| 601712813 | | | | | | | | | | | | | | | | | | | | | | | | 1 | | | | |
| 602044783 | | 3 | 2 | 1 | | | | | | | | | | | | | | | | | | | | | | | | |
| 602071049 | 6 | 5 | 3 | 6 | | | | | | | | | | | 5 | | 1 | | | 1 | | | | | 2 | | 1 | |
| 602382052 | | | | | | | | | | | | | | | | 1 | 1 | | | | | | | | | | | |
| 602399280 | | | | | | | | | | | | | | | | | | | | 1 | | | | | | | | |
| 602645152 | | | | | | | | | | | 1 | | | | | 2 | 1 | | | | | | | | | | | |
| 602675369 | 4 | 2 | 2 | | | | | | | | | | | 4 | 16 | 43 | 46 | | | | | | 4 | | 7 | 4 | 1 | |
| 602723103 | | | | | | | | | | | | | | | | | | | | | | | 3 | 8 | | | | |
| 602749386 | | | | | | | | | | | | | | | | | | | | | | | | | | | 1 | |
| 603068370 | | | | | | | | | | | | 1 | | | | | | | | | | | | | | | | |
| 603095061 | | | | | | | | | 1 | | | | | | 1 | | | | | | | | | 1 | | | | |
| 603466235 | | | | | | | | | 1 | | | | | | | | | | | | | | | | 5 | | | |
| 603482385 | | | 1 | | | | | | | | | | | | | 1 | | | | | | | | | | 2 | 1 | |
| 603785283 | | | | | | | | | | | | | | | | | | | | | | | | | | | 1 | |
| 603824120 | | | | | | | | | | | | | | | | | | | | | | | | 1 | | | | |
| 603824307 | | | | | | | | | | | | | | | | | | | | | | | 3 | | | | | |

| | | | | | | | | | | | | | | | | | | | | | | | | | | | | |
|---|---|---|---|---|---|---|---|---|---|---|---|---|---|---|---|---|---|---|---|---|---|---|---|---|---|---|---|---|
| 604070461 | | | | | | | | | | | | | | | | 2 | 1 | | | | | | | | | | | |
| 604182144 | | | | | | | | | | | | | | | | | | | | | | | | | 1 | | | |
| 604194925 | 2 | 4 | 6 | 2 | | | | | | 1 | | 1 | | | | | | | | | | | 4 | 1 | | | | |
| 604195063 | | 1 | 5 | 1 | | | | | | | | | | 1 | 6 | 1 | | | | | | | | | 1 | | | |
| 604195247 | 3 | | | | | | | | | | | | | 1 | 1 | 1 | 1 | 1 | 2 | 2 | 1 | | 1 | 2 | 1 | 5 | 2 | |
| 604476368 | | | | | | | | | | | | | | | | | | | | | | | | | | | 1 | |
| 604510234 | | | | | | | | | | | | | | | | | | | | | | | 2 | | | | | |
| 604510524 | | | | | | | | | | | | | | | | | | | | | | | | | 7 | 1 | | |
| 604527772 | | | | | | | | | | | | | | | | | | | | | | | | | | | 1 | |
| 604634863 | | | | | | | | | | | | | | | | | | | 1 | 1 | | | | | | | | |
| 604652566 | | | | | | | | | | | | | | | 1 | | | | | | | | | | | | | |
| 604847018 | | | | | | | | | | | | | | | | | | | | | | | | | | 1 | | |
| 604856102 | | | | | | | | | | | | | | | | | | | | | | | | | 6 | | | |
| 604864322 | | | | | | | | | | | | 4 | | | | | | | | | | | | | 4 | 5 | 15 | |
| 604864392 | | | | | | | | | | | | 6 | 8 | 5 | | | | | | | | | | | | | 2 | |
| 604868757 | | | | | | | | | | | | | 3 | | | | | | | | | | | | | | | |
| 604877772 | | | | | | | | | | | | | | | | | | | | | | | | | | 8 | | |
| 604886083 | | | | | | | | | | | | | | | | | | | | | | | 1 | | | | | |
| 604889732 | | | | | | | | | | | | | | | | | | | | | | | 14 | 6 | | | | |
| 604976663 | | | | | | | | | | | | | | | | | | | | | | | | | | | | 1 |
| 604980831 | | | | | | | | | | | | | | | 1 | | | | | | | | | | | | | |
| 604989814 | | | | | | | | | | | | 9 | 9 | 1 | 1 | | | | | | | | | | | | | 5 |
| 604998092 | | | | | | | | | | | | | | | | | | | | | | | | | | | | 4 |
| 605154073 | | | | | | | | | | | | | 1 | | | | | | | | | | | | | | | |
| 605317863 | | | | | | | | | | 1 | | | 1 | | | | | | | | | | | | | | | |
| 605550395 | | | | | | | | | | | | 29 | 15 | | | | | | | | | | | | 1 | | | |
| 605554875 | | | | | | | | | | | | 2 | 11 | | | | | | | | | | | | | | | |
| 605654689 | | | | | | | | | | | | | | | | | | | | | | | | | | | | 1 |
| 605662705 | | | | | | | | | | | | 14 | 7 | 1 | | | | | | | | | | | | | | 2 |
| 605662755 | | | | | | | | | | | | | | | | | | | | | | | | | | | | 1 |
| 605775210 | | | | | | | | | | | | | | 1 | | 1 | | | | | | | | | | | | |
| 605994959 | | | | | | | | | | | | 8 | 8 | | 1 | | | | | | | | | | | | | |
| 605999323 | | | | | | | | | | | | 7 | 5 | | | | | | | | | | | | | | | |
| 606193500 | | | | | | | | | | | | | | | | 1 | | | | | | | | | | | | |
| 606340818 | | | | | | | | | | | | 1 | | 3 | | | | | | | | | | | | | | |
| 606603565 | | | | | | | | | | | | | | | 1 | | | | | | | | | | | | | |
| 606914301 | | | | | | | | | | | 2 | | | | | | | | | | | | | | | | | |
| 607010448 | | | | | | | | | | 3 | | | | | | | | | | | | | | | | | | |
| 607260192 | | | | | | | | | | | | | | | | | | | | | | | | | 7 | | | |
| 607273128 | | | | | | | | | | | | 7 | 13 | 11 | 7 | | | | | | | | | | 4 | 1 | | |
| 607273339 | | | | | | | | | | | | 1 | 5 | 8 | 4 | 1 | | | | | | | | | 1 | | | |
| 607294784 | | | | | | | | | | | | | | | | | | | | 1 | | | | | | | | |
| 607346347 | | | | | | | | | | | | | 1 | | | | | | | | | | | | | | | |
| 607635892 | | | | | | | | | | | | | | | | | | | | | | 1 | | | | | | |
| 607929192 | | | | | | | | | | | 1 | 6 | 16 | 1 | | | | 2 | | 1 | 1 | | 3 | | | 7 | 3 | |
| 607938091 | | | | | | | | | | | | | | | | 1 | | | | | | | | | | | | |
| 607955261 | | | | | | | | | | | | 2 | 1 | | | | | | | | | | | | | | | |
| 607998424 | | | | | | | | | | | | | | | | | | | | 1 | | 1 | | | | | | |
| 608042044 | | | | | | | | | | | | 2 | 1 | | | | | | | | | | | | | | | |
| 608261812 | | | | | | | | | | | | 1 | | | 1 | | | | | | | | | | | | | |
| 608278754 | | | | | | | | | | | 2 | 9 | | | | | | | | | | | | | 1 | | | |
| 608301148 | | | | | | | | | | | | | | | | | | | | | | | | | 9 | 1 | 1 | |
| 608347901 | | | | | | | | | | | | | | | | | | 1 | | | | | | | | | | |
| 608602824 | | | | | | | | | | | | 7 | 6 | | | | | | | | | | | | | | | |
| 608715651 | | | | | | | | | | | | 5 | 5 | | | | | | | | | | | | | | | 39 |
| 609000216 | | | | | | | | | | | | | | | | | | | | | | | | | 1 | | | |
| 609384093 | | | | | | | | | | | | 6 | 16 | | | | | | | | | | | | | | | 6 |
| 609587511 | | | | | | | | | | | | | | | | | | | | | | | | | 2 | | | |
| 609591758 | | | | | | | | | | | | | | | | | | | | | | | | | 1 | | | |
| 609591880 | | | | | | | | | | | | | | | | | | | | | | | | | 5 | | | |
| 609591989 | | | | | | | | | | | | | | | | | | | | | | | | | 1 | | | |
| 609655670 | | | | | | | | | | | | 16 | 35 | 10 | 2 | | | | | | | | | | 3 | | | |
| 609656444 | | | | | | | | | | | | | 2 | | | | | | | | | | | | | | | |
| 609773847 | | | | | | | | | | | | | | | | | | | | | | | | | | | | 2 |
| 609993550 | | | | | | | | | | | | | 2 | | | | | | | | | | | | | | | |
| 610018976 | | | | | | | | | | | | | | | 2 | | | | | | | | | | | | | |
| 610049447 | | | | | | | | | | 4 | 3 | | | | | | | | | | | | | | | | | |
| 610074665 | | | | | | | | | | | | | | | | | | | | | | | | | | | | 1 |
| 610385481 | | | | | | | | | | 1 | | | | | | | | | | | | | | | | | | |
| 610415691 | | | | | | | | | | | | 18 | 12 | | | | | | | | | | | | | | | 20 |
| 610666582 | | | | | | | | | | | | 1 | 8 | | | | | | | | | | | | 5 | 1 | 4 | |
| 610704870 | | | | | | | | | 1 | 1 | | | | | | | | | | | | | | | | | | |
| 611015907 | | | | | | | | | | | | | 1 | | | | | | | | | | | | 1 | | | |
| 611016455 | | | | | | | | | | | | | | | | | | | | | | | | | 2 | 5 | 7 | 1 |
| 611029037 | | | | | | | | | | | | | | | | | | | | | | | | | 24 | | | |
| 611067721 | | | | | | | | | 4 | | 1 | | | | | | | | | | | | | | | | | |
| 611089612 | | | | | | | | | | | | 4 | 7 | | | | | | | | | | | | | | | 1 |
| 611340150 | | | | | | | | | | | | 11 | 30 | 3 | 1 | | | | | | | | | | 1 | 1 | | |
| 611974080 | | | | | | | | | | | | | | | | | 1 | | | | | | | | | | | |
| 612073194 | | | | | | | | | | 2 | | | | | | | | | | | | | | | | | | |

| | | | | | | | | | | | | | | | | | | | | | | | | | | | | |
|---|---|---|---|---|---|---|---|---|---|---|---|---|---|---|---|---|---|---|---|---|---|---|---|---|---|---|---|---|
| 612121276 | | | | | | | | | | 3 | | | | | | | | | | | | | | | | | | |
| 612134763 | | | | | | | | | | | | | | | | | | | | | | | | | | | | 1 |
| 612358673 | | | | | | | | | | | | | | | | | | | | | | | | | 2 | | | |
| 612695501 | | | | | | | | | | | | | | | | | | | | | | | | | 6 | | | |
| 613040803 | | | | | | | | | | | | | | | | | | | | | | | | | 1 | | | |
| 613161596 | | | | | | | | | | | | | | | | | | | | | | | | | | | | 3 |
| 613179816 | | | | | | | | | | | | | | | | | | | | | | | | | | | | 1 |
| 613822435 | | | | | | | | | 1 | 2 | 6 | | | | | | | | | | | | | | | | | |
| 613873982 | | | | | | | | | | | | | | | | | | | | | | | | | | | | 1 |
| 614189799 | | | | | | | | | | | | | | | | | | | | | | | | | | | | 1 |
| 633093547 | | | | | | | | | | | | | | | | 1 | 6 | | | | | | | | | | | |
| 633097767 | | | | | | | | | | | | | | | 1 | 2 | | | | | | | 1 | 2 | 1 | | | |
| 633097828 | | | | | | | | | | | | | | | | 78 | 70 | | | | | | | | | | | |
| 633101926 | | | | | | | | | | | | | 1 | | | | | | | | | | | | | | | |
| 633412502 | | | | | | | | | | | | | | | | | | | | | | | | 10 | | | | |
| 633412554 | | | | | | | | | | | | | | | | | | | | | | | 2 | 8 | | | | |
| 633438933 | | | | | | | | | | | | | | | | 3 | 1 | | | | | | | | | | | |
| 633447458 | | | | | | | | | | | | | | | | | 1 | | | | | | | | | | | |
| 633793553 | | | | 1 | | | | | | | | | | | | 68 | 71 | | | | | | | | 1 | | | |
| 634501239 | | | | | | | | | | | 1 | | | | | | | | | | | | | | | | | |
| 634867234 | | | | | | | | | | | | | | | | | | | | | | | | 1 | 1 | | 1 | |
| 634879889 | | | 2 | 1 | | | | | | | | 1 | | | 4 | 2 | | | | | | | | 2 | 2 | 2 | | |
| 635523991 | | | | | | | | | | | | | | | | | | | | | | | | | 1 | | | 2 |
| 635540653 | | | | | | | | | | | | | | | | | | | | | | | | 2 | | | | |
| 635541640 | | | | | | | | | | | | | | | 2 | | | | | | | | | 1 | | | | |
| 635545357 | | | | | | | | | | | | | | | | | | | | | | | | | 7 | | | |
| 635860358 | | | | | | | | | | | | | | 1 | | 2 | | | | | | | | | 1 | | | |
| 635878327 | | | | | | | | | | | | | | | | | | | | | | | | | 1 | | | |
| 635895000 | | | | | | | | | | | | 2 | 1 | | | | | | | | | | | | | 1 | 3 | |
| 635903360 | | | | | | | | | | | | | | | | | | | | | | | | | | 1 | 1 | |
| 635903433 | | | | | | | | | | | | | | | | | | | | | | | | | 3 | | 1 | |
| 635903703 | | | | | | | | | | | | | | | | | | | | | | | | | 1 | | 1 | |
| 635907365 | | | | | | | | | | | | | | | | | | | | | | | | | | | 1 | |
| 636244893 | | | | | | | | | | | | | | | | | | | | | | | | | | | 1 | |
| 636248936 | | | | | | | | | | | | | | | | | | | | | | | | | | 5 | 2 | |
| 636344136 | | | | | | | | | | | | 1 | | 12 | 3 | | | 3 | 2 | 2 | 2 | | | | | | | |
| 636356423 | | | | | | | | | | | | | 1 | 1 | | | | | | | | | | | | | | 1 |
| 636589318 | | | | | | | | | | | | 1 | | | | | | | | | | | | | | | | |
| 636594376 | | | | | | | | | | | | | | 1 | | 7 | | | | | | | | | | 2 | 1 | |
| 636602582 | | | | | | | | | | | | 2 | | | | | | | | | | | | | 1 | | | |
| 636667489 | | | | | | | | | | | | | | 3 | | | | | | | | | | | | | | |
| 636926507 | | | | | | | | | | | | | | | | | | | | | | | | | | | 1 | |
| 636944318 | | | | | | | | | | | | | | 2 | | | | | | | | | | | 12 | 2 | | |
| 636947974 | | | | | | | | | | | | | | | | | | | | | | | | | | | 1 | |
| 637034367 | | | | | | | | | | | | | | | | | | | | | | | | | | | | 1 |
| 637047419 | | | | | | | | | | | | | | | | | | | | | | | | | | | | 1 |
| 637280110 | | | | | | | | | | | | 1 | | | | | | | | | | | | | | | | |
| 637289438 | | | | | | | | | | | | | 4 | | | | | | | | | | | | | | | |
| 637306291 | | | | | | | | | | | | | 1 | 1 | 4 | | | 1 | 4 | 1 | | | 8 | 16 | | | | |
| 637371565 | | | | | | | | | | | | | | | | | | | | | | | | | | | | 1 |
| 637371568 | | | | | | | | | | | | | | | | | | | | | | | | | | | | 1 |
| 637621553 | | | | | | | | | | | | 1 | 2 | | 1 | | | | | | | | | | | | | |
| 637673302 | | | | | | | | | | | | | | | | | | 2 | | | | 1 | | | | | | |
| 637699424 | | | | | | | | | | | | 1 | | | | | | | | | | | | | | | | |
| 637914938 | | | 1 | | | | | | | | | | | | | 2 | 6 | | | | | | | | 4 | | | |
| 637949445 | | | | | | | | | | | 3 | | | | | | | | | | | | | | | | | |
| 637957861 | | | | | | | | | | | | | | | 1 | | | | | | | | | | | | | |
| 638040562 | | | | | | | | | | | | | | 1 | | | | | | | | | | | | | | |
| 638057952 | | | | | | | | | | | | 1 | | | | | | | | | | | | | | | | |
| 638316334 | | | | | | | | | | | | | | | | | | | | | | | | | | 4 | 1 | |
| 638360015 | | | | | | | | | | | | | 1 | 1 | | | | | | | | | | | | | | |
| 638740232 | | | | | | | | | | | | | 1 | | | | | | | | | | | | | | | |
| 638985608 | | | | | | | | | | | | | | | | | | | | | | | 1 | | | | 1 | |
| 639007697 | | | | | | | | | | | | | | | | | | | | | | | | | | 2 | | |
| 639008013 | | | | | | | | | | | | | | 6 | 1 | | | | | | | | | | | | | |
| 639050664 | | | | | | | | | | | | | | | | | | | | | | | | | | | | 2 |
| 639063702 | | | | | | | | | | | | 1 | 2 | | | | | | | | | | | | | 3 | 4 | |
| 639327084 | | | | | | | | | | | | 3 | 4 | | | | | | | | | | | | 6 | | | |
| 639405054 | | | | | | | | | | | | | | | | | | | | | | | | | | | | 1 |
| 639650941 | | | | | | | | | | | | 6 | 4 | | | | | | | | | | | | | | | |
| 639655378 | | | | | | | | | | | | 1 | | | | | | | | | | | | | | | | |
| 639659397 | | | | | | | | | | | | 4 | 7 | | | | | | | | | | | | | | | |
| 639667528 | | | | | | | | | | | | | | | | | | | | | | | | | | 2 | | |
| 639995972 | | | | | | | | | | | | | | | | | | | | | | | | | 3 | | | |
| 640082371 | | | | | | | | | | | | | | | | | | | | | | | | | | | | 2 |
| 640104852 | | | | | | | | | | | | | | | | | | | | | | | | | | | | 1 |
| 640427907 | | | | | | | | | | | | | | | | | | | | | | | | | | | | 1 |
| 640539567 | | | | | | | | | | | | | 1 | | | | | | | | | | | | | | | |
| 640596169 | | | | | | | | | | | | | | | | | | | | | | | | 2 | | | | |
| 640691030 | | | | | | | | | | | | 2 | | | | | | | | | | | | | | 1 | 2 | |

| | | | | | | | | | | | | | | | | | | | | | | | | | | | |
|---|---|---|---|---|---|---|---|---|---|---|---|---|---|---|---|---|---|---|---|---|---|---|---|---|---|---|---|
| 640691209 | | | | | | | | | | | | | | | | | | | | | | | | 27 | | | |
| 640755994 | | | | | | | | | | | 14 | 16 | 16 | 1 | | | | | | | | | | | | | 5 |
| 640760406 | | | | | | | | | | | 7 | 6 | | | | | | | | | | | | | | | |
| 640963092 | | | | | | | | | | | | | | | | | | | | | | | | 1 | | | |
| 641057518 | | | | | | | | | | | | | | | | | | | | | | | | | | | 1 |
| 641105431 | | | | | | | | | | | | | | | | | | | | | | | | | | | 2 |
| 641110014 | | | | | | | | | | | | | | | | | | | | | | | | | | | 1 |
| 641403248 | | | | | | | | | | 1 | | | | | | | | | | | | | | | | | |
| 641688553 | | | | | | | | | | | | | | | | | | | | | | | | 2 | | | |
| 641701225 | | | | | | | | | | | | | | | | | | | | | | | | | | 2 | |
| 642033429 | | | | | | | | | | | | | | | | | | | | | | | | 4 | | | |
| 642046803 | | | | | | | | | | | 1 | | | | | | | | | | | | | | | | 1 |
| 642085399 | | | | | | | | | | 3 | | | | | | | | | | | | | | | | | |
| 642146802 | | | | | | | | | | | | | | | | | | | | | | | | | | | 1 |
| 642262407 | | | | | | | | | | | | | | | | | | | | | | 1 | | | | | |
| 642387793 | | | | | | | | | | | 1 | | | | 1 | | | | | | | | | | | | |
| 643156727 | | | | | | | | | | | 1 | | | | | | | | | | | | | | | | |
| 643393020 | | | | | | | | | | | | 3 | | | | | | | | | | | | | | | |
| 643393688 | | | | | | | | | | | 1 | | | | | | | | | | | | | | | | |
| 643401816 | | | | | | | | | | | | | | | | | | | | | | | | 4 | | | |
| 643502099 | | | | | | | | | | | | | | | | | | | | | | | | | | | 1 |
| 643860525 | | | | | | | | | | | | | | | | | | | | | | | | | | | 1 |
| 644433538 | | | | | | | | | | | | 1 | | | | | | | | | | | | | | | |
| 662197764 | | | | | | | | | | | | | | | | | | | | | | 7 | 3 | | | 1 | |
| 663432544 | | | | | | | | | | | | | | | 1 | 2 | | | | | | | | | | | |
| 663437249 | | | | | | | | | | | | | 1 | | | 2 | | | | | | | | | | | |
| 664128120 | | | | | | | | | | | | | | | 1 | | | | | | | | | | | | |
| 664447502 | | | | | | | | | | | | | 1 | | | | | | | | | | | | | | |
| 664456036 | | | | | | | | | | | | | | | | | | | | | | | | 2 | | | |
| 664490596 | | | | | | | | | | | | | | | | 3 | | | | | | | | | | | |
| 664824047 | | | | | | | | | | | | 1 | | | | 1 | | | | | | | | 4 | | | |
| 664827233 | | | | | | | | | | | | | | | | 2 | | | | | | | | | | | |
| 665466951 | | | | | | | | | | | | 2 | 4 | 2 | | | | | | | | | | | | | |
| 665552356 | | | | | | | | | 3 | 1 | 3 | | | | | | | | | | | 2 | 1 | | | | |
| 665556968 | 2 | 7 | 2 | 3 | | | | | | | 1 | | 1 | 2 | | | | | | | | | | 1 | | 3 | |
| 665820084 | | | | | | | | | | | | | 1 | | | | | | | | | | | | | | |
| 665897487 | 3 | 1 | 6 | 3 | | | | | | | | | | 6 | 1 | | | | | | | | | 3 | | | |
| 666162219 | | | | | | | | | | | 2 | | 1 | | | | | | | | | | | | | | |
| 666230992 | | | | | | | | | | | | | | | | | | | | | | | | 2 | | | |
| 666252297 | | | | | | | | | | | | | | | 1 | | | | | | | | | | | | |
| 666555064 | | | | | | | | | | | | | | | | | | | | | | | 1 | | | 1 | |
| 666575508 | | | | | | | | | | | | | | | | | | | | | | | 1 | 8 | | | |
| 666618877 | | | | | | | | | | | | 2 | | | | | | | | | | | | | | | |
| 666916874 | | | | | | | | | | | | | | | | 1 | | | | | | | | | | | |
| 666942629 | | | | | | | | | | | | | | | | | | | | | | | | | 1 | 1 | |
| 667266529 | | | | | | | | | | | | | | 1 | | | | | | | | | | | | | |
| 667486204 | | | | | | | | | | | | | | | 1 | | | | | | | | | | | | |
| 667599306 | | | | | | | | | | | | | | | | | | | | | | | 2 | | | | |
| 667603691 | | | | | | | | | | | | | | | | | | | | | | | | | | 1 | |
| 667620012 | | | | | | | | | | | | | | | | | | | | | | | | | 7 | 2 | |
| 667642283 | | | | | | | | | | | | 3 | | | | | | | | | | | | | | | |
| 667711365 | | | | | | | | | | | | | | 1 | | | | | | | | | | | | | |
| 667750417 | | | | | | | | | | | | | | | | | | | | | | | | 1 | | | |
| 667965556 | | | | | | | | | | | 1 | | | | | | | | | | | | | | | | |
| 668056654 | | | | | | | | | | | | 2 | | | | | | | | | | | | | | | |
| 668065234 | | | | | | | | | | | 7 | 4 | 1 | | | | | | | | | | | | | | |
| 668311132 | | | | | | | | | | | 1 | | | | | | | | | | | | | | | | |
| 668340913 | | | | 1 | | | | | | | | | | | 1 | 5 | | | | | | 1 | | 1 | | | |
| 668384542 | | | | | | | | | | | 28 | 30 | | 1 | | | | | | | | | | | | | 7 |
| 668397992 | | | | | | | | | | | | | | | | | | | | 1 | | | | | | | |
| 668652267 | | | | | | | | | | | 1 | | | | | | | | | | | | | | | | |
| 668660617 | | | | | | | | | | | | | 1 | | | | | | | | | | | | | | |
| 668742424 | | | | | | | | | | | 1 | | | | | | | | | | | | | | | | |
| 669066952 | | | | | | | | | | | | 2 | | | | | | | | | | | | | | | |
| 669075559 | | | | | | | | | | | 2 | 1 | 1 | | | | | | | | | | | | | | |
| 669355635 | | | | | | | | | | | | 1 | | | | | | | | | | | | | | | |
| 669403115 | | | | | | | | | | | | | | | | | | | | | | | | | | | 2 |
| 669421910 | | | | | | | | | | | 1 | | 1 | | | | | | | | | | | | | | 1 |
| 669675454 | | | | | | | | | | | 4 | | | | | | | | | | | | | 2 | | | |
| 669683969 | | | | | | | | | | | | | | | | | | | | | | | | 2 | 1 | | |
| 669697117 | | | | | | | | | | | | | | | | | | | | | | | | | | 1 | |
| 669739832 | | | | | | | | | | | | 3 | | 1 | | | | | | | | | | | | | |
| 669744300 | | | | | | | | | | | 20 | 9 | 2 | | | | | | | | | | | | | | |
| 670020922 | | | | | | | | | | | 1 | 1 | | | | | | | | | | | | | | | |
| 670024927 | | | | | | | | | | | | | | | | | | | | | | 1 | | | 1 | 1 | |
| 670072514 | | | | | | | | | | | | | | 1 | | | | | | | | | | | | | |
| 670077134 | | | | | | | | | | | | | 2 | 2 | | | | | | | | | | | | | |
| 670094261 | | | | | | | | | | | 6 | 2 | 2 | | | | | | | | | | | | | | 3 |
| 670111641 | | | | | | | | | | | | 2 | | | | | | | | | | | | | | | |
| 670353202 | | | | | | | | | | | | 1 | | | | | | | | | | | | | | | |

| | | | | | | | | | | | | | | | | | | |
|---|---|---|---|---|---|---|---|---|---|---|---|---|---|---|---|---|---|---|
| 670357457 | | | | | | | | 5 | | | | | | | | | | |
| 670370149 | | | | | | | 4 | | | | | | | | 1 | | | |
| 670370239 | | | | | | | | | 2 | 1 | | | | | | | | |
| 670370808 | | | | | | | 2 | 3 | | | | | | | 2 | | | |
| 670430840 | | | | | | | | 1 | | | | | | | | | | |
| 670430945 | | | | | | | 5 | 8 | | | | | | | | | | |
| 670431030 | | | | | | | 26 | 46 | 1 | 2 | | | | | | | | 3 |
| 670435190 | | | | | | | 1 | 1 | | | | | | | | | | 1 |
| 670435566 | | | | | | | | 2 | | | | | | | | | | |
| 670448503 | | | | | | | | 1 | | | | | | | | | | |
| 670448622 | | | | | | | | 1 | | | | | | | | | | |
| 670711621 | | | | | | | | | | | | | | | 2 | | 1 | |
| 670772147 | | | | | | | 40 | 25 | | | | | | | | | | 4 |
| 670776253 | | | | | | | 2 | 5 | | | | | | | | | | 3 |
| 670776480 | | | | | | | 4 | 6 | | | | | | | | | | 9 |
| 670883838 | | | | | | | 14 | 8 | | | | | | | | | | |
| 671234167 | | | | | | | | 1 | | | | | | | | | | |
| 671393987 | | | | | | | | | | | | | | | 1 | | | |
| 671401894 | | | | | | | | | | | | | | | | | 2 | |
| 671449467 | | | | | | | 1 | 4 | | | | | | | | | | 2 |
| 672248510 | | | | | | | | | | | | | 1 | | | | | |
| 672490308 | | | | | | | 5 | 2 | | | | | | | | | | |
| 672835284 | | | | | | | 1 | 4 | | | | | | | | | | |
| 672960748 | | | | | | | | | | | | | | | 1 | 1 | | |
| 673072855 | | | | | | | 1 | | | | | | | | | | | |
| 673189141 | | | | | | | | | | | | | | | | | | 1 |
| 673513595 | | | | | 1 | | | | | | | | | | | | | |
| 673746277 | | | | | | | | | | | | | | | 2 | | | |
| 673758836 | | | | | | | | 1 | | | | | | | | | | |
| 673867858 | | | | | | | 7 | 6 | | | | | | | | | | |
| 674108632 | | | | | | | 24 | 26 | | | | | | | | | | |
| 674427795 | | | | | | | | 1 | | | | | | | | | | |
| 675490252 | | | | | | | | | | | | | | | 2 | | | |
| 675767414 | | | | | | | | | | | | | | | 1 | | | |
| 675940037 | | | | | | | | | | | | | | | | | | 1 |
| 676262737 | | | | | | | | | | | | | | | | | | 1 |
| 694463250 | | | | | | | | | | | | 4 | | | | | | |
| 694471913 | | | | | | | | | | | 1 | 2 | | | | | | |
| 694472338 | | | | | | | | | | | 6 | 2 | | | | | | |
| 694813336 | | | | | | | | | | | 3 | 2 | | | | | | |
| 694813676 | | | | | | | | | | | 7 | 5 | | | | | | |
| 694818168 | | | | | | | | | | | | 1 | | | | | | |
| 695124651 | 1 | | | 8 | | | | 1 | 1 | 7 | | | | | | | | |
| 695128320 | | | | | | | 1 | | 2 | 1 | | 1 | | | | | | |
| 695128641 | | | | | | | | | | | | | | 2 | | | | |
| 695128703 | | | | | | | | | | | | | | 2 | | | | |
| 695132361 | | | | | | | | | 1 | | 1 | | | | | | | |
| 695154095 | | | | | | | | | | | 3 | | | | | | | |
| 695154667 | | | | | | | | | | | 3 | | | | | | | |
| 695172463 | | | | | | | | | | | 1 | 1 | | | | | | |
| 695465316 | | | | | | | | | | | | 1 | | | | | | |
| 695469100 | | | | | | | | | 1 | | | | 3 | 2 | | | | |
| 695508439 | | | | | | | | | | | | | | | 1 | | | |
| 695798021 | | | | | | | | | | | 1 | | | | | | | |
| 695875649 | | | | | | | | | | | | 1 | | | | | | |
| 696215925 | | | | | | | | | | | | | 1 | | | | | |
| 696216553 | | | | | | | | | | | | | 1 | | | | | |
| 696489026 | | | | | | | | | | | 1 | 1 | | | | | | |
| 696829800 | | | | | | | | | | | | 1 | | | | | 1 | |
| 696846594 | | | | | 2 | 2 | | | | 2 | 2 | 16 | | 1 | | | | |
| 696850811 | | 3 | | | | | | | | | 2 | | | | | | | |
| 696859236 | | | | | | | | | 1 | | | | | | | | | |
| 697252464 | | | | | | | | | | | | 1 | 1 | | | | | |
| 697265869 | | | | | | | | | | 2 | 1 | | | | 1 | | | |
| 697269889 | | | | | | | | | | | | | | | 2 | | | |
| 697295304 | | | | | | | | | | | | | | | | 2 | | |
| 697645622 | 4 | 5 | 3 | | | | | 2 | | 1 | | | 7 | 8 | | 2 | | |
| 697886966 | | | | | | | 2 | 5 | | | | | | | | | | |
| 697925511 | | | | | | | | | | | | | 2 | 2 | | | | |
| 697985761 | | | | | | | | | | | | | | | 1 | | | |
| 698072467 | | | | | | | | | 1 | | | | | | | | | |
| 698224251 | | | | | | | | 2 | | | | | | | | 1 | | |
| 698314499 | | | | | | | 4 | 8 | | | | | | | | | | |
| 698331117 | | | | | | | | | | | | | | | 1 | | | |
| 698418398 | | | | | | | 1 | | | | | | | | | | | |
| 698418640 | | | | | | | 2 | 1 | | | | | | | | | | |
| 698677285 | | | | | | | | | | | | | | | | 4 | | |
| 698742094 | | | | | | | | | 1 | | | | | | | | | |
| 698970725 | | | | | | | | | | | 3 | 3 | | | | | | |
| 698996246 | | | | | | | 6 | 13 | | | | | | | | | | |

| | | | | | | | | | | | | | | | | | | | | | | | | | | | | |
|---|---|---|---|---|---|---|---|---|---|---|---|---|---|---|---|---|---|---|---|---|---|---|---|---|---|---|---|---|
| 699031185 | | | | | 3 | | 1 | 1 | 1 | 2 | | | | | | | | | | | | | 1 | | | | | |
| 699078755 | | | | | | | | | | | | | | | | | | | | | | | | | | | | 3 |
| 699363923 | | | | | | | | | | | | 16 | 16 | | | | | | | | | | | | 1 | | | |
| 699687305 | | | | | | | | | | | | 18 | 22 | 2 | 2 | | | | | | | | | | | | 1 | 2 |
| 699695617 | | | | | | | | | | | | 2 | 4 | | 1 | | | | | | | | | | | | | |
| 700002191 | | | | | | | | | | | | | | | | | | | | | | | | 2 | | | | |
| 700028161 | | | | | | | | | | | | | | | | | | | | | | | | | 2 | | | |
| 700037217 | | | | | | | | | | | | 14 | 11 | 1 | | | | | | | | | | | | | | |
| 700437926 | | | | | | | | | | | | 1 | | | | | | | | | | | | | | | | 2 |
| 700731556 | | | | | | | | | | | | | | | | | | | | | | | | | 1 | | | |
| 700779022 | | | | | | | | | | | | | | | | | | | | | | | | | | | | 6 |
| 700801298 | | | | | | | | | | | | 6 | 7 | | | | | | | | | | | | | | | |
| 700805759 | | | | | | | | | | | | 6 | | | | | | | | | | | | | | | | |
| 701051650 | | | | | | | | | | | | 3 | | | | | | | | | | | | | | | | |
| 701055731 | | | | | | | | | | | | | | | | | | | | | | | | | 8 | | 5 | |
| 701077130 | | | | | | | | | | | | | | | 4 | | | | | | | | | | | | | |
| 701120315 | | | | | | | | | | | | | 1 | | | | | | | | | | | | | | | |
| 701151126 | | | | | | | | | | | | 4 | 1 | | | | | | | | | | | | | | | |
| 701409339 | | | | | | | | | | | | | | | | | | | | | | | | | | | 1 | |
| 701469455 | | | | | | | | | | | | | | 2 | | | | | | | | | | | | | | |
| 701474029 | | | | | | | | | | | | 1 | | | | | | | | | | | | | | | | |
| 701742479 | | | | | | | | | | | | 1 | | | | | | | | | | | | | | | | |
| 701746442 | | | | | | | | | | | | 1 | | | | | | | | | | | | | | | | |
| 701746512 | | | | | | | | | | | | 3 | | | | | | | | | | | | | | | | |
| 701750547 | | | | | | | | | | | | | | | | | | | | | | | | | | 3 | 1 | |
| 701759740 | | | | | | | | | | | | | | 1 | 1 | | | | | | | | | | | | | |
| 701806850 | | | | | | | | | | | | | 1 | | | | | | | | | | | | | | | |
| 702152113 | | | | | | | | | | | | 25 | 39 | | | | | | | | | | | | | | | |
| 702760942 | | | | | | | | | | | | 2 | | | | | | | | | | | | | | | | |
| 703179782 | | | | | | | | | | | | 1 | | | | | | | | | | | | | | | | |
| 703779386 | | | | | | | | | | | | 4 | 8 | | | | | | | | | | | | | | | |
| 703866010 | | | | | | | | | | | | | | | | | | | | | | | | | | | | 1 |
| 704453006 | | | | | | | | | | | | 3 | 2 | | | | | | | | | | | | | | | |
| 704487840 | | | | | | | | | 3 | | | | | | | | | | | | | | | | 1 | | | |
| 704552421 | | | | | | | | | | | | | 1 | | | | | | | | | | | | | | | |
| 704880200 | | | | | | | | | | | | 1 | 1 | | | | | | | | | | | | | | | |
| 704898746 | | | | | | | | | | | | | 1 | | | | | | | | | | | | | | | |
| 705229969 | | | | | | | | | | | | | 3 | | | | | | | | | | | | | | | |
| 705243675 | | | | | | | | | | | | | 3 | | | | | | | | | | | | | | | |
| 705938591 | | | | | | | | | | | | | 1 | | | | | | | | | | | | | | | |
| 706831482 | | | | | | | | | | | | | | | | 1 | | | | | | | | | | | | |
| 722471825 | | | | | | | | | | | | | | | 1 | | | | | | | | | | | | | |
| 725498630 | | | | | | | | | | | | | | | | 1 | 1 | | | | | | | | | | | |
| 725817857 | | | | | | | | | | | | | | 1 | | | | | | | | | | | | | | |
| 725839388 | | | | | | | | | | | | | | | | | 1 | | | | | | | | | | | |
| 726171930 | | | | | | | | | | | | | | | | 2 | 2 | | | | | | | | | | | |
| 726184776 | | | | | | | | | | | | | | | | 3 | | | | | | | | | | | | |
| 726207450 | | | | | | | | | | | | | | | | | 1 | | | | | | | | | | | |
| 726867105 | | | | | | | | | | | | | | | | 2 | 2 | | | | | | | | | | | |
| 727182822 | | | | | | | | | | | | | | | | | | | | | | | | 1 | | | | |
| 727212462 | | | | | | | | | | | | | | | | 2 | | | | | | | | | | | | |
| 727255451 | | | | | | | | | | | | | | | | 1 | | | | | | | | | | | | |
| 727260771 | | | | | | | | | | | | | 1 | | | | | | | | | | | | | | | |
| 727280915 | | | | | | | | | 2 | | 1 | | | | | | | | | | | | | 1 | | | | |
| 727285326 | | | | | | | | | | | | | | 1 | 1 | | | | | | | | | | | | | |
| 727929407 | | | | | | | | | | | | | | 1 | | | | | | | | | | | | | | |
| 728205616 | | | | | | | | | | | | | | | | | 1 | | | | | | | | | | | |
| 728292005 | | | | | | | | | | | | | | | | | | | | | | | | | 1 | | | |
| 728342932 | | | | | | | | | | | | | | | | | | | | | | | | | 2 | | | |
| 728633073 | | | | | | | | | | | | | | | | 1 | | | | | | | | | | | | |
| 729137627 | | | | | | | | | | | | | | | | | | | | | | | | | | | 1 | |
| 729444443 | | | | | | | | | | | | | 1 | | | | | | | | | | | | | | | |
| 729698667 | | | | | | | | | | | | 2 | 11 | | | | | | | | | | | | | | | |
| 729711951 | | | | | | | | | | | | | | | | | | | | | | | | | 5 | | | |
| 729715904 | | | | | | | | | | | | | | | | | | | | | | | | | | 2 | | |
| 729716051 | | | | | | | | | | | | | | | | | | | | | | | | | | | 1 | |
| 729781112 | | | | | | | | | | | | | | | | | | | | | | | | | | | | 6 |
| 729785519 | | | | | | | | | | | | | 2 | | | | | | | | | | | | | | | |
| 729790291 | | | | | | | | | | | | 1 | 1 | | | | | | | | | | | | | | | |
| 729954382 | | | | | | | | | | | | | | | | 2 | | | | | | | | | | | | |
| 730104740 | | | | | | | | | | | | | | 1 | | | | | | | | | | | | | | |
| 730118077 | | | | | | | | | | | | 5 | 2 | | | | | | | | | | | | | | | 6 |
| 730130818 | | | | | | | | | | | | 2 | 1 | | | | | | | | | | | | | | | |
| 730131007 | | | | | | | | | | | | 15 | 5 | | | | | | | | | | | | | | | |
| 730398148 | | | | | | | | | | | | 3 | 3 | | | | | | | | | | | | | | | 1 |
| 730398229 | | | | | | | | | | | | 3 | 11 | | | | | | | | | | | | | | | |
| 730475736 | | | | | | | | | | | | 1 | | 4 | | | | | | | | | | | | | | |
| 730717696 | | | | | | | | | | | | 1 | | | | | | | | | | | | | | | | |
| 730721558 | | | | | | | | | | | | 2 | 5 | | | | | | | | | | | | | | | |
| 730743732 | | | | | | | | | | | | | 1 | | | | | | | | | | | | | | | |

| | | | | | | | | | | | | | | | | | | | | |
|---|---|---|---|---|---|---|---|---|---|---|---|---|---|---|---|---|---|---|---|---|
| 730791480 | | | | | | | | | | | | | | | | | | | | 1 |
| 730804352 | | | | | 1 | | | | | | | | | | | | | | | |
| 731062953 | | | | 3 | 12 | | | | | | | | | | | | | | | |
| 731140650 | | | | 11 | | | | | | | | | | | | | | | | |
| 731140809 | | | | 36 | 18 | | | | | | | | | | | | | | | |
| 731145160 | 3 | 1 | | 13 | 10 | | | | | | | | | | | | | | | |
| 731145304 | | | | 4 | 3 | | | | | | | | | | | | | | | |
| 731482010 | | | | 14 | 5 | | | | | | | | | | | | | | | |
| 731494823 | | | | 10 | 1 | | | | | | | | | | | | | | | |
| 731831926 | | | | | 1 | | | | | | | | | | | | | | | |
| 732077752 | | | | 1 | | | | | | | | | | | | | | | | |
| 732086360 | | | | | 2 | | | | | | | | | | | | | | | |
| 732155427 | | | | 1 | | | | | | | | | | | | | | | | 1 |
| 732431788 | | | | 4 | 1 | | | | | | | | | | | | | | | |
| 732496591 | | | | 1 | | | | | | | | | | | | | | | | 9 |
| 732777416 | | | | 4 | | | | | | | | | | | | | | | | |
| 733182574 | 2 | 7 | | | | 1 | 1 | | | | | | | | | | | | | |
| 733192083 | | | | 2 | 3 | | | | | | | | | | | | | | | |
| 733873838 | | | | 1 | | | | | | | | | | | | | | | | |
| 734327429 | | | | 2 | 7 | | | | | | | | | | | | | | | |
| 734547253 | | | | | 1 | | | | | | | | | | | | | | | |
| 734901302 | | | | 1 | | | | | | | | | | | | | | | | |
| 735816081 | | | | | | | | | | | | | | | | | 2 | | | |
| 737189212 | | | | | | | | | | | | | | | 1 | | | | | |
| 738155917 | | | | | | | | | | | | | | | | | 1 | | | |
| 739567785 | | | | | | | | | | | | | | | | 1 | | | | |
| 756502912 | | | | | | | 2 | 3 | | | | | | | | | | | | |
| 756623488 | | | | | 1 | | | | | | | | | | | | | | | |
| 756870190 | | | | | | | | 1 | | | | | | | | | | | | |
| 756874510 | | | | | | | | 3 | 2 | | | | | | | | | | | |
| 757215581 | | | | | | | | 1 | | | | | | | | | | | | |
| 757556799 | | | | | | | | 2 | | | | | | | | | 1 | | | |
| 758320421 | | | | | | | | | | | | | | | | | 4 | | | |
| 758636869 | | | | | | | | 1 | 3 | | | | | | | | | | | |
| 758903321 | | | | | | | | | 1 | | | | | | | | | | | |
| 759136185 | | | | 1 | | | | | | | | | | | | | | | | |
| 759632919 | | | | | | | 1 | 2 | | | | | | | | | | | | |
| 759633968 | | | | | | | | | | | | | | | | | 1 | | | |
| 759792147 | | | | | | 1 | | | | | | | | | | | | | | |
| 759810119 | | | | | | | | | 1 | | | | | | | 1 | | | | |
| 760133472 | | | | | | 1 | | | | | | | | | | | | | | |
| 760293208 | | | | | | | | | | | | | | | | 4 | | | | |
| 760470777 | | | | | | | | | | | | | | | | | | | | 2 |
| 760487313 | | | | 6 | 1 | | | | | | | | | | | | | | | |
| 760690116 | | | | | | | | | | | | | | | | | 2 | | | |
| 760821030 | | | | 9 | 5 | | | | | | | | | | | | | | | |
| 760828908 | | | | 3 | 2 | | | | | | | | | | | | | | | |
| 761013909 | | | | | | | | | | | | | | | 2 | | | | | |
| 761083880 | | | | 5 | | | | | | | | | | | 1 | | | | | |
| 761088041 | | | | | 1 | | | | | | | | | | | | | | | |
| 761165432 | | | | 17 | 7 | 4 | 1 | | | | | | | | | | | | | |
| 761174130 | | | | 1 | | | | | | | | | | | | | | | | |
| 761441765 | | | | | | | 1 | | | | | | | | | | | | | |
| 761489398 | | | | | 2 | | | | | | | | | | | | | | | |
| 761744194 | | | | | | | | | | | | | | | | | 1 | | | |
| 761813205 | | | | | | | | | | | | | | 1 | | | | | 2 | 1 |
| 761838951 | | | | | 3 | | | | | | | | | | | | | | | |
| 761843188 | | | | 3 | | | | | | | | | | | | | | | | |
| 762119200 | | | | | 2 | | | | | | | | | | | | | | | |
| 762176004 | | | | 4 | | | | | | | | | | | | | | | | |
| 762184774 | | | | | 1 | | | | | | | | | | | | | | | |
| 762525980 | | | | 19 | 11 | | | | | | | | | | | | | | | |
| 762526201 | | | | 3 | | | | | | | | | | | | | | | | |
| 762534597 | | | | 1 | | | | | | | | | | | | | | | | |
| 762810507 | | | | | | 1 | | | | | | | | | | | | | | |
| 762857726 | | | | 22 | 8 | | | | | | | | | | | | | | | |
| 762857949 | | | | 16 | 6 | | | | | | | | | | | | | | | |
| 763544185 | | | | 6 | 3 | | | | | | | | | | | | | | | |
| 764144161 | | | | 5 | 1 | | | | | | | | | | | | | | | |
| 764148834 | | | | 13 | 8 | | | | | | | | | | | | | | | |
| 764222141 | | | | | 1 | | | | | | | | | | | | | | | |
| 764843337 | | | | | | | | | | | | | | | | | 3 | | | |
| 765245188 | | | | | 1 | | | | | | | | | | | | | | | |
| 765590602 | | | | 4 | 3 | | | | | | | | | | | | | | | |
| 765931172 | | | | 5 | 6 | | | | | | | | | | | | | | | |
| 766385396 | | | | 1 | | | | | | | | | | | | | | | | |
| 767313008 | | | | | | | | | | | | | | | | | 1 | | | |
| 767455273 | | | | | | | | | | | | | | | | | | | 1 | |
| 767878191 | | | | | | | | | | | | | | | | 1 | | | | |
| 768582563 | | | | | | | | | | | | | | | | | 3 | | | |

| | | | | | | | | | | | | | | | | | | | | | | | | | | | | |
|---|---|---|---|---|---|---|---|---|---|---|---|---|---|---|---|---|---|---|---|---|---|---|---|---|---|---|---|---|
| 769544445 | | | | | | | | | | | | | | | | | | | | | | | 63 | 46 | | | | |
| 770257411 | | | | | | | | | | | | | 1 | | | | | | | | | | | | | | | |
| 787882939 | | 1 | | | | | | | | | | | | 1 | | | | | | | | | | 1 | | | | |
| 787883174 | | | | | | | | | | | | | | | | | | | | | | | | | 1 | 2 | | |
| 787883482 | | | | | | | | | | | | | | | | | | | | | | | | | 2 | 1 | | |
| 787908929 | | | | | | | | | | | | | | | | 3 | 1 | | | | | | | | | | | |
| 787909611 | | | | | | | | | | | | | | | | | | | | | | | | | 1 | | | |
| 789330613 | | | | | | | | | | | | | | | | | | | | | | | 2 | | | 1 | | |
| 789726348 | | | | | | | | | | | | | | | | | | | | | | | | | 2 | | | |
| 789851418 | | | | | | | | | | | | | | | | | | | | 2 | | | | | | | | |
| 789990540 | | | | | | | | | | | | | | 1 | | | | | | | | | | | | | 1 | |
| 790417325 | | | | | | | | | | | | 1 | | 1 | | | | | | | | | | | | | | |
| 790508166 | | | | | | | | | | | | | | 1 | | | | | | | | | | | | | | |
| 790762435 | | | | | | | | | | | | | | | | | | | | | | | | | | | 1 | |
| 791371842 | | | | | | | | | | | | | | | | | | | | | | | | | | 1 | | |
| 791527493 | | | | | | | | | | | | | | 7 | 1 | | | | | | | | | | | | | |
| 791699510 | | | | | | | | | | | | | | | | | | | | | | | 2 | | | | | |
| 791738711 | | | | | | | | | | | | | | | 1 | | | 1 | | | | | | | | | | |
| 791876556 | | | | | | | | | | | | 3 | 4 | | | | | | | | | | | | | | | |
| 792040520 | | | | | | | | | | | | | | | | | | | | | | | | 1 | | | | |
| 792040687 | | | | | | | | | | | | | | | | | | | | | | | 1 | | | | | |
| 792040691 | | | | | | | | | | | | | | | | | | | | | | | 1 | | | | | |
| 792100709 | | | | | | | | | | | | 1 | | | | | | | | | | | | | | | | |
| 792386006 | | | | | | | | | | | | | | | | | | | | | | | 1 | 6 | | | | |
| 792692885 | | | | | | | | | | 1 | | | | | | | 5 | | | | | | | | | | 1 | |
| 792856887 | | | | | | | | | | | | | 1 | | | | | | | | | | | | | | | |
| 792891477 | | | | | | | | | | | | 20 | 6 | | | | | | | | | | | | | | | |
| 793879702 | | | | | | | | | | | | | | 3 | 5 | | | 8 | 6 | 1 | 1 | 4 | | | | | | |
| 793892682 | | | | | | | | | | | | 6 | | | | | | | | | | | | | | | | |
| 793901460 | | | | | | | | | | | | | 1 | | | | | | | | | | | | | | | |
| 794246765 | | | | | | | | | | | | 4 | 3 | | | | | | | | | | | | | | | |
| 794574805 | | | | | | | | | | | | | 1 | | | | | | | | | | | | | | | |
| 794591963 | | | | | | | | | | 2 | | 11 | 7 | | | | | | | | | | | | | | | |
| 794919689 | | | | | | | | | | | | 4 | 3 | | | | | | | | | | | | | | | |
| 794920152 | | | | | | | | | | | | 6 | 9 | | | | | | | | | | | | | | | |
| 795187525 | | | | | | | | | | | | | | | | | | | | | | | | | | | 1 | |
| 795546494 | 4 | 1 | 2 | 2 | | | | | | | | | | | 1 | | 1 | | | | | | | 2 | | | | |
| 795943400 | | | | | | | | | | | | 14 | 14 | | | | | | | | | | | | | | | |
| 796393522 | | | | | | | | | | | | 3 | 7 | | | | | | | | | | | | | | | |
| 796730640 | | | | | | | | | | | | 1 | | | | | | | | | | | | | | | | |
| 797079495 | | | | | | | | | | | | 1 | | | | | | | | | | | | | | | | |
| 801257348 | | | | | | | | | | | | | | | | | | | | | | | | | 2 | | | |
| 818240386 | 2 | 1 | 4 | 3 | | | | | | | | | | | | | | | | | | | | | | | | |
| 818930866 | | | | | | | | | | | | | | | | 1 | | | | | | | | | | | | |
| 820015203 | | | | | | | | | | | | | | | | | 1 | | | | | | | | | | | |
| 821007911 | | | | | | | | | | | | | | 1 | 2 | | | | | | | | | | | | | |
| 821340054 | | | | | | | | | | | | | | | | | 1 | | | | | | | | | | | |
| 821621397 | | | | | | | | | 1 | | | | | | | | | | | | | | | | | | | |
| 821702605 | | | | | | | | | | | | | 1 | | | | | | | | | | | | | | | |
| 821780859 | | | | | | | | | | | | | | | 1 | | | | | | | | 1 | 3 | 2 | 2 | | |
| 822376419 | | | | | | | | | | | | | | | | | | | | | | | | | | | 1 | |
| 822380754 | | | | | | | | | | | | 5 | 3 | | | | | | | | | | | | | 1 | | |
| 822562470 | | | | | | | | | | | | | | | | | | | | | | | | | | | | 3 |
| 822885728 | | | | | | | | | | | | | 1 | | | | | | | | | | | | | | | |
| 823063281 | | | | | | | | | | | | | | | | 5 | 1 | | | | | | | | | | | |
| 823230871 | | | | | | | | | | | | | 1 | | | | | | | | | | | | | | | |
| 823240287 | | | | | | | | | | | | | 3 | | | | | | | | | | | | | | | |
| 823580438 | | | | | | | | | | | | | | 2 | | | | | | | | | | | | | | |
| 823584732 | | | | | | | | | | | | 1 | 1 | | | | | | | | | | | | | | | |
| 823590011 | | | | | | | | | | | | 1 | 4 | | | | | | | | | | | | | | | |
| 823852745 | 6 | | 7 | 3 | | | | | | | | | | | 5 | | | | | | | | | 1 | 1 | 2 | 1 | |
| 823917726 | | | | | | | | | | | | 3 | 1 | | | | | | | | | | | | | | | |
| 823938775 | | | | | | | | | | | | | | | | | | | | | | | | | | | | 1 |
| 824115747 | | | | | | | | | | | | 2 | 1 | 1 | 1 | | | | | | | | | | | | 1 | |
| 824279766 | | | | | | | | | | 2 | | | | | | | | | | | | | | | | | | |
| 824940802 | | | | | | | | | | | | 1 | | | | | | | | | | | | | | | | |
| 825277806 | | | | | | | | | | | | 5 | 7 | | | | | | | | | | | | | | | |
| 825278007 | | | | | | | | | | | | 1 | 1 | | | | | | | | | | | | | | | |
| 827590855 | | | | | | | | | | | | 1 | | | | | | | | | | | | | | | | |
| 829141094 | | | | | | | | 5 | | | | | | 1 | | 7 | 3 | | | | | | | | 1 | | | |
| 829155062 | | | | | | | | | | | | | | | | | | | | | | | | 3 | | | | |
| 849270788 | | | | | | | | | | | | | | 1 | 2 | 1 | 1 | | 1 | | | | 3 | | 1 | | | |
| 849270890 | 1 | | | 1 | | | | | | | | | | | | | | | | | | | | | | | | |
| 849607776 | | | | | | | | | | | | | | 1 | | | | | | | | | | | | | | |
| 849612438 | 1 | | 3 | 2 | | | | | | | | | | | | | | | | | | | | | | | | |
| 850294342 | | | | | | | | | | | | | | | | 2 | | | | | | | | | | | | |
| 850717220 | | | | | | | | | | | | | | | | | 1 | | | | | | | | | | | |
| 851459972 | | | | | | | | | | | | | | | | 2 | 2 | | | | | | | | | | | |
| 852422640 | | | | | | | | | | | | | | | | | | | | | | | | | 1 | | | |
| 852733214 | | | | | | | | | | | | 2 | | | | | | | | | | | | | | | | |

| | | | | | | | | | | | | | | | | | | | | | | | | | | | | |
|---|---|---|---|---|---|---|---|---|---|---|---|---|---|---|---|---|---|---|---|---|---|---|---|---|---|---|---|---|
| 853376855 | | | | | | | | | | | | | | | | | 1 | | | | | | | | | | | |
| 853967794 | | | | | | | | | | | | 1 | | | | | | | | | | | | | | | | 3 |
| 854611780 | | | | | | | | | | | | 1 | 3 | | | | | | | | | | | | | | | |
| 854620103 | | | | | | | | | | | | 2 | 2 | | | | | | | | | | | | | | | |
| 854636911 | | | | | | | | | | | | 1 | 2 | | | | | | | | | | | | | | | |
| 854953009 | | | | | | | | | | | | 1 | | | | | | | | | | | | | | | | |
| 854974323 | | | | | | | | | | | | | | | | | | | | | | | | | | | | 1 |
| 855630800 | | | | | | | | | | | | 10 | 7 | | | | | | | | | | | | | | | |
| 855638407 | | | | | | | | | | | | 1 | | | | | | | | | | | | | | | | |
| 856338646 | | | | | | | | | | | | | 1 | | | | | | | | | | | | | | | |
| 856886459 | | | | | | | | | | | | | | | | | | | | | | | | | 1 | | | |
| 858237673 | | | | | | | | | | | | | | | | | | | | | | | | | | | 1 | |
| 859006021 | | | | | | | | | | | | | | 2 | | | | | | | | | | | | | | |
| 859265651 | | | | | | | | | | | | | | | | | | | | | | | | | 1 | | | |
| 859339195 | | | | | | | | | | | | | | | | | | | | 1 | | 1 | | | | | | |
| 859554255 | | | | | | | | | | | | | | 3 | | | | | | | | | | | | | | 1 |
| 859977121 | | | | | | | | | | | | | | | | | | | | | | | | | 1 | | | |
| 861233275 | | | | | | | | | | | | | | | | | | | | | | | 1 | | | | | |
| 861233292 | | | | | | | | | | | | | | | | | | | | | | | 1 | | | | | |
| 861578690 | | | | | | | | | | | | | | | | | | | | | | | | 1 | | | | |
| 880305613 | | | 2 | 1 | | | | | | | | | | | | | | | | | | | | | | | | |
| 880987657 | | | | | | | | | | | | | | | | | 1 | | | | | | | | | | | |
| 881005404 | | | | | | | | | | | | | | | | | 1 | | | | | | | | | | | |
| 881329648 | | | | | | | | | | | | | | | | | 1 | | | | | | | | | | | |
| 882383361 | | | | | | | | | | | | | | | | | 1 | | | | | | | | | | | |
| 884140963 | | | | | | | | | | | | | | 7 | 2 | | | | | | | | | 2 | 1 | | | |
| 884484964 | | | | | | | | | | | | | | | | | | | | | | | | | 3 | | | |
| 884921013 | | | | | | | | | | | | 1 | | | | | | | | | | | | | | | | 1 |
| 885119984 | | | | | | | | | | | | | | | 1 | | | | | | | | | | | | | |
| 885257860 | | | | | | | | | | | | 7 | 16 | 1 | 6 | | | | | | | | | | | | | |
| 885262087 | | | | | | | | | 4 | | | | | | | | | | | | | | | | | | | |
| 885599043 | | | | | | | | | | | | | | | | | | | | | | | | 1 | | | | |
| 885654338 | | | | | | | | | | | | 1 | 4 | | | | | | | | | | | | | | | |
| 886008733 | | | | | | | | | | | | | | | | | | | | | | | | | | | | 1 |
| 886328403 | | | | | | | | | | | | 13 | 20 | | | | | | | | | | | | | | | |
| 886367305 | | | | | | | | | | | | | 1 | | | | | | | | | | | | | | | |
| 886643729 | | | | | | | | | | | | | | | | | | | | | | | | | | | | 1 |
| 887930118 | | | | | | | | | | | | | | 3 | | | | | | | | | | | | | | |
| 891733363 | | | | | | | | | 1 | | | | | | | | | | | | | | | | | | | |
| 891927856 | | | | | | | | | | | | | | | | | | | | | | | 2 | | | | | |
| 911332304 | | | | | 2 | | | 2 | | | | | | 1 | 3 | | 1 | | | | | | | | | | | |
| 911681886 | | | | | | | | | | | | | | | | | | | | | | | 1 | | | | | |
| 912601209 | | | | | | | | | | | | | | | | 5 | | | | | | | | | | | | |
| 914492942 | | | | | | | | | | | | | 1 | 3 | | | | | | | | | 1 | 2 | | | | |
| 914834542 | | | | | | | | | | | | | | | | | | | | | | | | | 1 | | | |
| 915511932 | | | | | | | | | | | | | | | 1 | | | | | | | | | | | | | |
| 915610561 | | | | | | | | | | | | | | | | | | | | | | | | | 1 | | | |
| 915852772 | | | | | | | | | | | | | | | | | | | | | | | 1 | | | | | |
| 916655865 | | | | | | | | | | | | | | | | | | | | | | | | | | | | 2 |
| 917656984 | | | | | | | | | | | | | | | | | | | | | | | | 2 | | | | |
| 919168397 | | | | | | | | | | | | 4 | 2 | | | | | | | | | | | | | | | |
| 921935093 | | | | | | | | | | | | 1 | | | | | | | | | | | | | | | | |
| 924723528 | | | | | | | | | | | | | | | | 1 | | | | | | | | | | | | |
| 943420912 | | | | | | | | 1 | | | | | | | 1 | | | | | | | | | | | | | |
| 943822621 | | | | | | | | 1 | | | | | | | | | | | | | | | | | | | | |
| 944379777 | | | | | | | | | | | | | | | | | | | | | | | | | 1 | | | |
| 945354221 | | | | | | | | | | | | | 2 | | | | | | | | | | | | | | | |
| 945483923 | | | | | | | | | | | | | | | 1 | | | | | | | | | | | | | |
| 945527072 | | | | | | | | | | | | | | 4 | 5 | | | | | | | | 2 | | 3 | | | |
| 945869060 | | | | | | | | | | | | | | | | | | | | | | | 1 | | | | | |
| 946204816 | | | | | | | | | | | | | | | | | | | | | | | | | | | 1 | |
| 946546270 | | | | | | | | | | | | | | 1 | | | | | | | | | | | | 1 | | |
| 946550009 | | | | | | | | | | | | | | | 2 | | | | | | | | | | 1 | | | |
| 946978006 | | | | | | | | | | | | | | | | | | | | | | | | | | | | 3 |
| 947258925 | | | | | | | | | | | | | | | | | | | | | | | 2 | | | | | |
| 947391929 | | | | | | | | | | | | | 1 | | | | | | | | | | | | | | | |
| 947898091 | | | | | | | | | | | | 1 | | | | | | | | | | | | | | | | |
| 948363764 | | | | 1 | | | | | | | | 2 | 4 | 4 | 5 | | | | | | | | | 1 | | | | |
| 949045811 | 1 | 3 | | | | | | | | | | | | 2 | | | 1 | | | | | | 1 | | 1 | | | |
| 950591319 | | | | | | | | | | | | 2 | 7 | | | | | | | | | | | | 1 | | | |
| 951738965 | 4 | 6 | 5 | 5 | | | | | | | | 1 | | | 2 | 2 | | | | | | | | | 2 | 1 | 4 | |
| 952727922 | | | | | | | | | | | | | | | | | 1 | | | | | | | | | | | |
| 952792408 | | | | | | | | | | 1 | 5 | 1 | 2 | 10 | 5 | | | | | | | | | | | | | |
| 954679716 | | | | | | | | | | | | | | | | | | | | | | | 2 | | | | | |
| 973786403 | | | | | | | | | | | | | | | | 1 | | | | | | | | | | | | |
| 974771466 | | | | | | | | | | | | | | | | 1 | | | | | | | | | | | | |
| 974839580 | | | | | | | | | | | | | | | | | 1 | | | | | | | | | | | |
| 974843801 | | | | | | | | | | | | | | | 1 | | | | | | | | | | | | | |
| 974843875 | | | | | | | | 2 | | | | | | | | | 1 | | | | | | | | | | | |
| 974843957 | | | | | | | | | | | | | | | 1 | | | | | | | | | | | | | |

| | | | | | | | | | | | | | | | | | | | | | | | | | | | | |
|---|---|---|---|---|---|---|---|---|---|---|---|---|---|---|---|---|---|---|---|---|---|---|---|---|---|---|---|---|
| 975197713 | | | | | | | | | | | | | | | | | | | | | | | 2 | | | | | |
| 976903657 | | | | | | | | | | | | | | | | | | | | | | | 1 | | | | | |
| 977002744 | | | | | | | | | | | | | | | | | | | | | | | | | 1 | | | |
| 977710985 | | | | | | | | | | | | | | | | | | | | | | | 1 | | | | | |
| 977926158 | | | | | | | | | | | | | | | | | | | | | | | | | 1 | | | |
| 977995787 | 2 | 2 | | | | | | | | | | | | | | | 4 | | | | | | | 2 | | | | |
| 978051730 | | | | | | | | | | | | | | | | | | | | | | | | 1 | | | | |
| 978056035 | | | | | | | | | | | | 1 | | | | | | | | | | | | | | | | |
| 978778113 | | | | | | | | | | | | 4 | 2 | | | | | | | | | | | | | | | 1 |
| 979065964 | | | | | | | | | | | | | | | | | | | | | | | | 1 | | | | |
| 982994156 | | | | | | | | | | | | | | | | | | | | | | | | 1 | | | | |
| 983146199 | | | | | | | | | | | | | | | | | | 1 | 1 | | | | | | | | | |
| 984008765 | | | | | | | | | | | | | | 1 | | | | | | | | | | | | | | |
| 987424042 | | | | | | | | | | | | | | | | | | | | | | | | 1 | | | | |
| 987759904 | | | | | | | | | | | | | | | | | | | | | | | 74 | 39 | | | | |
| 1004436788 | | | | | | | | 1 | | | | | | | 1 | | 1 | | | | | | | | | | | |
| 1005174948 | | | | | | | | | | | | | | | | | | | | | | | | | | 1 | 1 | |
| 1005520576 | | | | | | | | | | | | | | 1 | | | | | | | | | | | | | | |
| 1006539760 | | | | | | | | | | | | | | | 1 | | 1 | | | | | | | | | | | |
| 1007087260 | | | | | | | | | | | | | | | | | | | | | | | | | | | | 1 |
| 1007402796 | | | | | | | | | | | | | | | 3 | | | | | | | | | | | | | |
| 1009578838 | | | | | | | | | | | 5 | | | | | | | | | | | | | | | | | |
| 1009984360 | | | | | | | | | | | | | | | | 1 | | | | | | | | | | | | |
| 1010157364 | | | | | | | | | | | | 1 | | | | | | | | | | | | | | | | |
| 1010731508 | | | | | | | | | | | | | | | | | | | | | | | | 1 | | | | |
| 1015049194 | | | | | | | | | | | | | | | | 1 | | | | | | | 1 | | | | | |
| 1015385741 | | | | | | | | | | | | | | | | | | | | | | | | 1 | | | | |
| 1035459511 | | | | | | | | | | | | | | | | 1 | | | | | | | | | | | | |
| 1036550962 | | | | | | | | | | | | | | | | | | | | | | | 2 | | 15 | 1 | 3 | |
| 1036637638 | | | | | | | | | | | | | | 2 | | | | | | | | | 1 | 5 | | | | |
| 1042495376 | | | | | | | | | | | | | | | | | | | | | | | | 1 | | | | |
| 1046039966 | | | | | | | | | | | | | | | | | | | | | | | 1 | | | | | |
| 1046039987 | | | | | | | | | | | | | | | | | | | | | | | 3 | | | | | |
| 1046044924 | | | | | | | | | | | | | | | | | | | | | | | 2 | | | | | |
| 1046385193 | | | | | | | | | | | | | | | | | | | | | | | 2 | | | | | |
| 1046385221 | | | | | | | | | | | | | | | | | | | | | | | 1 | | | | | |
| 1046726043 | | | | | | | | | | | | | | | | | | | | | | | 1 | | | | | |
| 1046726261 | | | | | | | | | | | | | | | | | | | | | | | 31 | 27 | | | | |
| 1046726280 | | | | | | | | | | | | | | | | | | | | | | | 2 | | | | | |
| 1046735096 | | | | | | | | | | | | | | | | | | | | | | | 3 | | | | | |
| 1047075623 | | | | | | | | | | | | | | | | | | | | | | | | 1 | | | | |
| 1047421689 | | | | | | | | | | | | | | | | | | | | | | | | 2 | | | | |
| 1047426385 | | | | | | | | | | | | | | | | | | | | | | | 47 | 19 | | | | |
| 1066511487 | | | | | | | | | | | | | | | 1 | 2 | 1 | | | | | | | 3 | | | | |
| 1067923228 | | | | | | | | | | | | | | | 1 | | | | | | | | | | | | | |
| 1072063538 | | | | | | | | | | | | 10 | 25 | 12 | 6 | 1 | 2 | | | | | | | | 1 | 2 | | |
| 1076724912 | | | | | | | | | | | | | | | | | | | | | | | 2 | 2 | | | | |
| 1076729226 | | | | | | | | | | | | | | | | | | | | | | | 2 | | | | | |
| 1076739012 | | | | | | | | | | | | | | | | | | | | | | | 1 | | | | | |
| 1077420145 | | | | | | | | | | | | | | | | | | | | | | | | 1 | | | | |
| 1077424204 | | | | | | | | | | | | | | | | | | | | | | | | 1 | | | | |
| 1077428662 | | | | | | | | | | | | | | | | | | | | | | | | 1 | | | | |
| 1077459077 | | | | | | | | | | | | | | | | | | | | | | | 19 | 20 | | | | |
| 1077459090 | | | | | | | | | | | | | | | | | | | | | | | 29 | 16 | | | | |
| 1077752379 | | | | | | | | | | | | | | | | | | | | | | | 4 | 9 | | | | |
| 1077756583 | | | | | | | | | | | | | | | | | | | | | | | 1 | | | | | |
| 1077756731 | | | | | | | | | | | | | | | | | | | | | | | 1 | 2 | | | | |
| 1077761302 | | | | | | | | | | | | | | | | | | | | | | | | 2 | | | | |
| 1078102419 | | | | | | | | | | | | | | | | | | | | | | | 2 | | | | | |
| 1078102468 | | | | | | | | | | | | | | | | | | | | | | | | 5 | | | | |
| 1078447707 | | | | | | | | | | | | | | | | | | | | | | | | 2 | | | | |
| 1078788519 | | | | | | | | | | | | | | | | | | | | | | | | 1 | | | | |
| 1095780246 | | | | | | | | | | | | | | | 1 | | | | | | | | | | | | | |
| 1096850735 | | | | | | | | | | | | | | 5 | 15 | | | | | | | | | | | | | |
| 1096859379 | | | | | | | | | | | | | 1 | | | | | | | | | | | | | | | |
| 1096859621 | | | | | | | 1 | 4 | | | | | | 1 | 2 | 1 | | | | | | | | | | | | |
| 1096894260 | | | | | | | | | | | | | | | | 1 | | | | | | | | | | | | |
| 1096963197 | | | | | | | | | | | | | | | | | | | | | | | | | 1 | | | |
| 1097240204 | | | | | | | | | | | | | | | 1 | | | | | | | | | | | | | |
| 1099389583 | | | | | | | | | | | | | | | 1 | | | | | | | | | | 1 | | | |
| 1099808012 | | | | | | | | | | | | | 1 | | | | | | | | | | | | | | | |
| 1100381898 | | | | | | | | | | | | 2 | | | | | | | | | | | | | | | | |
| 1104004164 | | | | | | | | | | | | | | | | | | | | | | | 1 | 3 | | | | |
| 1107423142 | | | | | | | | | | | | | | | | | | | | | | | 1 | | | | | |
| 1107764206 | | | | | | | | | | | | | | | | | | | | | | | 1 | | | | | |
| 1108109580 | | | | | | | | | | | | | | | | | | | | | | | 1 | | | | | |
| 1108109684 | | | | | | | | | | | | | | | | | | | | | | | 2 | | | | | |
| 1108166986 | | | | | | | | | | | | | | | | | | | | | | | 1 | | | | | |
| 1108459071 | | | | | | | | | | | | | | | | | | | | | | | 3 | 5 | | | | |
| 1108485790 | | | | | | | | | | | | | | | | | | | | | | | 1 | | | | | |

| | | | | | | | | | | | | | | | | | | |
|---|---|---|---|---|---|---|---|---|---|---|---|---|---|---|---|---|---|---|
| 1108791991 | | | | | | | | | | | | | | | 2 | | | |
| 1109133043 | | | | | | | | | | | | | | | 1 | | | |
| 1109141617 | | | | | | | | | | | | | | | | 1 | | |
| 1109141661 | | | | | | | | | | | | | | | | 1 | | |
| 1109149659 | | | | | | | | | | | | | | | | 1 | | |
| 1109478369 | | | | | | | | | | | | | | | 5 | | | |
| 1109832699 | | | | | | | | | | | | | | | 1 | | | |
| 1110164795 | | | | | | | | | | | | | | | | 2 | | |
| 1110868515 | | | | | | | | | | | | | | | 1 | | | |
| 1127886064 | | | | | | | | | | | | 5 | 3 | | | | | |
| 1127886209 | | | | | | | | | | | | | 1 | | | | | |
| 1127894310 | | | | | | | | | | | 1 | | | | | | | |
| 1127894824 | | | | | | | | | | | | 1 | | | | | | |
| 1127902989 | | | | | | | | | | 2 | 1 | | | | | | | |
| 1128235633 | | | | | | | | | | | | 3 | 1 | | | | | |
| 1128274593 | | | | | | | | | | 4 | 7 | 8 | 7 | | | | | |
| 1129306180 | | | | | | | | | | | | 3 | | | | | | |
| 1137590785 | | | | | | | 2 | | | | | | | | | | | |
| 1137767646 | | | | | | | | | | | | | | | | 1 | | |
| 1137767724 | | | | | | | | | | | | | | | | 1 | | |
| 1138453934 | | | | | | | | | | | | | | | | 1 | | |
| 1138458189 | | | | | | | | | | | | | | | | 1 | | |
| 1138462526 | | | | | | | | | | | | | | | 1 | | | |
| 1138807874 | | | | | | | | | | | | | | | | 1 | | |
| 1139144718 | | | | | | | | | | | | | | | 1 | 1 | | |
| 1140012085 | | | | | | | | | | | | | | 1 | | | | |
| 1140168071 | | | | | | | | | | | | | | | 1 | | | |
| 1140172133 | | | | | | | | | | | | | | | | 1 | | |
| 1140172174 | | | | | | | | | | | | | | | | 1 | | |
| 1140188920 | | | | | | | | | | | | | | | 1 | | | |
| 1140193366 | | | | | | | | | | | | | | | | 1 | | |
| 1140862679 | | | | | | | | | | | | | | | 10 | 2 | | |
| 1141213129 | | | | | | | | | | | | | | | 3 | | | |
| 1158631810 | | | | 1 | | | | | | | | | | | | | | |
| 1158657609 | | | | | | | | | | 1 | 1 | | | | | | | |
| 1158925051 | | | | | | | | | | 1 | | | | | | | | |
| 1158933766 | | 3 | | | | | | | | | | | | | | | | |
| 1158994954 | | | | | 1 | 2 | | | | | 2 | | | | | | | |
| 1159262177 | | | | | | | | | | | 1 | 3 | | | | | | |
| 1159611828 | | | | | | | | | | 1 | | | | | | | | |
| 1164131530 | | | | | | | | | | | | | | | 1 | | | |
| 1168625046 | | | | | | | | | | | | 1 | | | | | | |
| 1168798447 | | | | | | | | | | | | | | | | 1 | | |
| 1169359181 | | | | | | | | 1 | | | | | | | | | | |
| 1169576040 | | | | | | | | | | | | | | | 1 | | | |
| 1169898618 | | | | | | | | | | | | | | | 17 | 3 | | |
| 1170326295 | | | | | | | | | | | | 2 | 2 | | | | | |
| 1170667130 | | | | | | | | | | | | 1 | | | | | | |
| 1170857299 | | | | | | | | | | | | | | | | 2 | | |
| 1171207089 | | | | | | | | | | | | | | | 1 | | | |
| 1171207096 | | | | | | | | | | | | | | | 1 | | | |
| 1171548200 | | | | | | | | | | | | | | | | 1 | | |
| 1171621682 | | | | | | | | | | 1 | | | | | | | | |
| 1171690432 | | | | | | | | | | | | | | | | | 1 | |
| 1171910380 | | | | | | | | | | | | | | | 1 | | | |
| 1172713521 | | | | | | | | | | | | | 1 | | | | | |
| 1189619192 | | | | | | | | | | | | | 3 | | | | | |
| 1189641371 | | | | | | | | | | | 1 | | | | | | | |
| 1189964433 | | | | | | | | | | | | | 1 | | | | | |
| 1189973148 | | | | | | | | | | 4 | 3 | | | | | | | |
| 1190344036 | | | | | | 4 | | | | | | | | | | | | |
| 1190668746 | | | | | | | | | | | | | | | | | 2 | |
| 1199491728 | | | | | | | | | | | | | | | | 1 | | |
| 1199491747 | | | | | | | | | | | | | | | | 1 | | |
| 1199837203 | | | | | | | | | | | | | | | 1 | | | |
| 1199837219 | | | | | | | | | | | | | | | 1 | | | |
| 1200619473 | | | | | | | | | | | | | | | 1 | | | |
| 1200873417 | | | | | | | | | | | | | | | 2 | | | |
| 1220321245 | 2 | | 1 | | | | | | | | | | | | | | | |
| 1220434040 | | | | | | | | 3 | 1 | | | | | | | | | |
| 1225640135 | | | | | | | | | | | | | | | 2 | | | |
| 1251041030 | | | | | | | | | | 1 | | | | | | | | |
| 1252366645 | | | | | | | | | | 2 | 1 | | | | | | | |
| 1252409535 | | | | | | 1 | | | | | | | | | | | | |
| 1255289329 | | | | | | | | | | | | 1 | | | | | | |
| 1256619187 | | | | | | | | | | | | | | | | | | 2 |
| 1260879578 | | | | | | | | | | | | | 1 | | | | | |
| 1265037070 | | | | | | | | | 1 | | | | | | | | | |
| 1282097987 | | | 1 | 1 | | | | | | | 2 | | | | | | | |
| 1282413298 | | | | | | | | | | 2 | | | | | | | | |

| | | | | | | | | | | | | | | | | | | | |
|---|---|---|---|---|---|---|---|---|---|---|---|---|---|---|---|---|---|---|---|
| 1282754143 | | | | | | | | 1 | | | | | | | | | | | |
| 1283401111 | | | | | | | | | | | 1 | | | | | | | | |
| 1283406432 | | | | | | | | 1 | | | 1 | 1 | 2 | | | | | | |
| 1284117939 | | | | | | | | | | | | | 2 | | | | | | |
| 1288172070 | | | | | | | | | | | | | 1 | | | | | | |
| 1298804357 | | | | | | | | | | | | | | | | 2 | | | |
| 1313115480 | | | | | | | | 2 | | | | | | | | | | | |
| 1313538112 | | | | | | | | | | | 1 | | | | | | | | |
| 1320177531 | | | | | | | | | | | 1 | 1 | 44 | 41 | | | | | |
| 1344823436 | | | | | | | | 4 | | | | | | | | | | | |
| 1345087352 | | | | | | | | | | | | | 5 | | | | | | |
| 1345134192 | | | | | | | | | | | | | | 4 | | | | | |
| 1346814892 | | | | | | | | 1 | | | 7 | 10 | 38 | 47 | | | | | |
| 1348510522 | | | | | | | | | | | 1 | | | | | | | | |
| 1378212488 | | | | | 1 | | | | | | 1 | 1 | | | | | | | |
| 1378251040 | | | | | | | | | | | 1 | | | | | | | | |
| 1407554101 | | | | | | | | | | | | | | 4 | | | | | |
| 1418618235 | | | | | | | | | 5 | | | | | | | | | | |
| 1439262842 | | | | | | | | | | | 1 | | | | | | | | |
| 1439621300 | | | | | | | | 1 | | | | | | | | | | | |
| 1469623821 | | | | | | | | | | | | | | 4 | | | | | |
| 1481736987 | | | | | | | | | | | | 1 | | | 2 | | | 1 | |
| 1500663102 | | | | | | | | | | | | | | 2 | | | | | |
| 1501332510 | | | | | | | | | | | | 2 | | | | | | | |
| 1502700841 | | | | | | 1 | | | | 1 | 6 | 9 | | | | | | | |
| 1511240113 | | | | | | | | | | | | | | | | 1 | | | |
| 1532030136 | | | | | | | | | | | | | 3 | | | | | | |
| 1535902995 | | | | | | | | 1 | | | | | | | | | | | |
| 1562673627 | | | | | | | | 1 | | | | | | | | | | | |
| 1562737357 | | | | | | | | | | | 2 | | | | | | | | |
| 1563419167 | | | | | | | | 1 | | | | | | | | | | | |
| 1563929167 | | | | | | | 2 | | | | 1 | | | | | | | | |
| 1564399630 | | | | | | 1 | | | | | 10 | 9 | | | | | | | |
| 1570684665 | | | | | | | | | | | | | 1 | | | | | | |
| 1591701396 | | | | | | | | 1 | | | | | | | | | | | |
| 1658885859 | | | | | | | | | | | | | | 3 | | | | | |
| 1664980698 | 4 | | 5 | | | | | | | | | 1 | | | | | | | |
| 1670597454 | | | | 1 | | | | | | | | | | | | | | | |
| 1687555356 | | | | | | | | | | | | | 4 | | | | | | |
| 1689242749 | | | | | | | | | | | | 1 | | | | | | | |
| 1692278031 | | | | | | | | | | | | | | | | 1 | | | |
| 1702306047 | | | | | | | | | | | | | | 1 | | | | | |
| 1714639709 | | | | | | | | 1 | | | | | | | | | | | |
| 1745821751 | 1 | | | | | | | | | | | | | | | | | | |
| 1746373639 | | | | | | 1 | | | | | | | | | | | | | |
| 1749628974 | | | | 1 | | | | 1 | | | | 2 | | | | | | | |
| 1749974233 | | | | | | | | | | | | | | 4 | | | | | |
| 1750306408 | | | | | | | | | | | | 2 | | | | | | | |
| 1781000673 | | | | | | | | | | | | 1 | 1 | | | | | | |
| 1781000944 | | | | | | | | | | | | 4 | | | | | | | |
| 1788089103 | | | | | | 1 | | | | | | | | | | | | | |
| 1812035520 | | | | | | | | | | | | | 6 | | | | | | |
| 1839146351 | | | | | 1 | | | | | | | | | | | | | | |
| 1858901026 | | | | | | | | | | | | | 5 | 3 | | | | 1 | |
| 1874782886 | | | | | | | | | | | 1 | | | | | | | | |
| 1880183883 | | | | | | 1 | | | | | | | | | | | | | |
| 1881193508 | | | | | 2 | | | | | | | | | | | | | | |
| 1881401277 | | | | | | | | | | | | | | 1 | | | | | |
| 1881876433 | | | | | | | | 2 | | | | | | | | | | | |
| 1884625521 | | | | | | 3 | 3 | | | | 1 | | | | | | | | |
| 1899778504 | | | | | | | 1 | | | | | | | | | | | | |
| 1910528711 | | | | | | 1 | | | | | | | | | | | | | |
| 1912436017 | | | | | | | | | | | | | | | | | | | 1 |
| 1912789965 | | | | | | | | | | | | | | 1 | | | | | |
| 1913252043 | | | | | | | | 1 | | | | | | | | | | | |
| 1932479780 | | | | | | | | 1 | | | | | | | | | | | |
| 1935843067 | | | | 1 | | | | | | | | | | | | | | | |
| 1943812176 | | | | | | | | | | | | | 1 | 1 | | | | | |
| 1948107641 | | | | | | | | | | | | | 1 | 1 | | | | | |
| 1975187675 | | | | | | | | | | | | | 1 | | | | | | |
| 1975347348 | 9 | 1 | 5 | | | | | | | | | 1 | | | | | | | |
| 1975878958 | | | | | | | | | | | | | 3 | 1 | | | | | |
| 1999609010 | | | | | | | | | | | 1 | 1 | 2 | | | | | | |
| 1999945988 | | | | | | | | | | | | | 2 | | | | | | |
| 2003662505 | | | | | | | | | | | 1 | 5 | | | | | | | |
| 2007068523 | 5 | 3 | | 5 | | | | | | | | 3 | | | | | 1 | | |
| 2025584558 | | | | | | 1 | 1 | | | | | | | | | | | | |
| 2039096834 | | | | | | | | 2 | | | | | | | | | | | |
| 2039100722 | | | | | | 1 | | 2 | | | | | | | | | | | |
| 2062020024 | | | | | | | | 1 | | | | 1 | | | | | | | |

| | | | | | | | | | | | | | | | | | | | | | | | | | | | | |
|---|---|---|---|---|---|---|---|---|---|---|---|---|---|---|---|---|---|---|---|---|---|---|---|---|---|---|---|---|
| 2065745704 | | | | | 1 | | 1 | 1 | | | | | | 1 | | | | | | | | | | | | | | |
| 2067748663 | | | | | | | | | | | | | | | 1 | | | | | | | | | | | | | |
| 2067757054 | | | | | | | | 1 | | | | | | | | | | | | | | | | | | | | |
| 2068801704 | | 6 | 3 | 3 | | | | | | | | | | | 1 | | | | | | | | 3 | | | | | |
| 2069117386 | | | | | | | | 1 | | | | | | | | | | | | | | | | | | | | |
| 2070131720 | | | | | | | | | | | | | | | 1 | | | | | | | | | | | | | |
| 2094397733 | | | | | | | | | | | | | | | | | | | | | | | | 2 | | | | |
| 2097462638 | | | | | 1 | | | 2 | | | | | | | | | | | | | | | | | | | | |
| 2098455028 | | | | | | | | 1 | | | | | | | | | | | | | | | | | | | | |
| 2119349345 | | | | | | 1 | | | | | | | | | | | | | | | | | | | | | | |
| 2128838418 | | 1 | | | | | | | | | | | | | 2 | | | | | | | | | | | | | |
| 2129486034 | | | | | | | | 2 | | | | | | | | | | | | | | | | | | | | |
| 2129490083 | | | | | | | | 3 | | | | | | | | | | | | | | | | | | | | |
| 2130509519 | | | | | | | | 2 | | | | | | | | | | | | | | | | | | | | |
| 2151748489 | | | | | | 1 | | | | | | | | | | | | | | | | | | | | | | |
| 2152426324 | | | | | | 1 | | | | | | | | | | | | | | | | | | | | | | |
| 2152426357 | | | | | | 1 | | | | | | | | | | | | | | | | | | | | | | |
| 2152426385 | | | | | | 2 | | | | | | | | | | | | | | | | | | | | | | |
| 2154459819 | | | | | | 1 | | | | | | | | | | | | | | | | | | | | | | |
| 2155473983 | | | | | | 1 | | | | | | | | | | | | | | | | | | | | | | |
| 2162242957 | | | | | | | | | | | | | | | 1 | | | | | | | | | | | | | |
| 2191893077 | | | | | | | | | | | | | | | 2 | | | | | | | | | | | | | |
| 2191931219 | | | | | | | | | | | | | | 2 | 1 | | 1 | | | | | | | | | | | |
| 2221273723 | | | | | | | | 1 | | | | | | | | | | | | | | | | | | | | |
| 2222931956 | | | | | | | | | | | | | | | | | 1 | | | | | | | | | | | |
| 2253651262 | | | | | | | | | | | | | | | | 1 | | | | | | | | | | | | |
| 2254644527 | | | | | | | | 3 | | | | | | | | | | | | | | | | | | | | |
| 2289223313 | | | | | | | | | | | | | | | | | | | | | | 1 | 2 | | | | | |
| 2315060417 | | | | | | | | 4 | | | | | | | | | | | | | | | | | | | | |
| 5812979048 | | | | | | | | 6 | | | | | | | 1 | | | | | | | | | | | | | |
| 5812979938 | | | | | | | | | | | | | | | | | | | | | | | | | 1 | | | |
| 5812979953 | | | | | | | | | | | | | | | | | | | | | | | | 2 | | | | |
| 5812980006 | | | | | | | | | | | | | | | | 20 | 17 | | | | | | | 4 | | | 1 | |
| 5812980010 | 3 | | 2 | 4 | | | | | | | | | | 12 | 6 | 22 | 11 | | | | | | | | | | | |
| 5812980022 | | | | | | | | | | | | | | | | | | | | | | | | 1 | | | | |
| 5812980047 | | 1 | | | | | | | | | | | | 1 | | 1 | | | | | | | | | | | | |
| 5812980079 | | | | | | | | | 14 | 3 | 5 | | | | | | | | | | | | | | | | | |
| 5812980142 | | | | | | | | | 3 | | | | 1 | | | | | | | | | | | | | | 1 | |
| 5812980164 | | | 1 | | | | | | | | | | | | | | | | | | | | | | | | | |
| 5812980250 | 2 | 10 | 8 | 1 | | | | | | | 1 | | | | 5 | 1 | 4 | | | 1 | | | 1 | 9 | 1 | 1 | 1 | |
| 5812980257 | | | | | | | | | | | | 1 | | | | | | | | | | | | | | | | |
| 5812980282 | | | | | | | | | | | | | | | | 6 | 1 | | | | | | | | | | | |
| 5812980289 | | | | | | | | | | | | | | | | 44 | 65 | | | | | | | | | 1 | | |
| 5812980326 | | | | | | | | | | | | | | | | | | | | | | | | | 1 | | | |
| 5812980473 | | | | | | | | | | | | | | | | | | | | | | | 3 | 4 | | | | |
| 5812980483 | | | | | | | | | | | | | | | | | | | | | | | 1 | | | | | |
| 5812980497 | | | | | | | | | | | | | | | | | | | | | | | 1 | | | | | |
| 5812980501 | | | | | | | | | | | | | | | | | | | | | | 1 | | 1 | | | | |
| 5812980662 | | | | | | | | | | | | | | | 1 | | | | | | | | | | | | | |
| 5812980710 | | | | | | | | | | | | 2 | | 7 | 2 | | | | | | | | | | | | | |
| 5812980732 | | | | | | | | | | | | | | | | | | | | | | | | | 14 | | | 5 |
| 5812980884 | | | | | | | | | | | 1 | 1 | | 2 | 3 | | | | | | | | 1 | 2 | | 1 | | |
| 5812980938 | | | | | | | | | | | | | | | | | 1 | | | | | | | | | | | |
| 5812980982 | 2 | | | | | | | | | | | | | | | | | | | | | | | | | | | |
| 5812980984 | 1 | | | | | | | | 2 | 1 | 1 | | | 1 | 4 | | | 3 | 4 | 2 | 1 | 1 | 31 | 24 | | | | |
| 5812981000 | | | | | | | | | | | | | | | 1 | | | | | | | | | | | | | |
| 5812981016 | | | | | | | | | | | | | | | | 1 | | | | | | | | | | | | |
| 5812981194 | | | | | | | | | | | | | 1 | | 1 | | | 2 | 3 | | | | | | | | | |
| 5812981228 | | | | | | | | | | | | | 5 | | | | | | | | | | | | 1 | | | |
| 5812981237 | | | | | | | | | | | | | 1 | | | | | | | | | | | | | 1 | | |
| 5812981249 | | | | | | | | | | | | 2 | 5 | 2 | | | | | | | | | | | | | | |
| 5812981341 | | | | | | | | | | | | | | | | | 1 | | | | | | | | | | | |
| 5812981460 | | | | | | | | | | | | 1 | | 1 | 1 | 5 | 1 | | | | | | | | | | | |
| 5812981506 | | | | | | | | | | | | 4 | 2 | 2 | | | | | | | | | | | | 1 | | |
| 5812981517 | | | | | | | | | | | | 11 | 10 | | | | | | | | | | | | 3 | | | |
| 5812981715 | | | 1 | | | | | | | | | | | | | | | | | | | | | 2 | | | | |
| 5812981858 | | | | | | | | | | | | | | | | 2 | | | | | | | | | | | | |
| 5812981862 | | | | | | | | | | 1 | 1 | | | | | | | | | | | | | | | | | |
| 5812982247 | | | | | | | | | | | | | | 2 | | | | | | | | | | | | | | |
| 5812982278 | | | | | | | | | | | | | | 2 | | | | | | | | | | | | | | |
| 5812982324 | | | | | | | | | | | | 1 | | | | | | | | | | | | | | | 2 | |
| 5812982632 | | | | | | | | | | 1 | | | | | | | | | | | | | | 1 | | | 1 | |
| 5812982656 | | | | | | | | | | | | | | | | 1 | | | | | | | | | | | | |
| 5812983220 | | | | | | | | | 4 | 4 | 2 | | | 1 | 1 | | | | | | 1 | | 1 | | | | | |
| 5812983240 | | | | | | | | | | | | 10 | 6 | | | | | | | | | | | | | | | |
| 5812983519 | | | | | | | | | | | | | | | | | | | | | | | 2 | | | | | |
| 5812983533 | | | | | | | | | | | | | | | | | | | | | | | | | 1 | | | |
| 5812986565 | | | | | | | | | | | | | | | | | | | | | | | | 1 | | | | |
| 5812986870 | | | | | | | | | | | | | | | | | | | | | | | 1 | | | | | |
| 5812986876 | | | | | | | | | | | | | | | | | | | | | | | 21 | 4 | | | | |

| | | | | | | | | | | | | | | | | | | | | | | | | | | | | |
|---|---|---|---|---|---|---|---|---|---|---|---|---|---|---|---|---|---|---|---|---|---|---|---|---|---|---|---|---|
| 5812987236 | | | | | | | | | | | | | | | | | | | | | | | | 3 | | | | |
| 5812992942 | | | | | | | | | | | | | | | | | | | | | | | 1 | | | | | |
| 5812995141 | | | | | | | | | | | | | | | | 47 | 29 | | | | | | | | | | | |
| 5812995298 | | | | | | | | | | | | | | | | | 1 | | | 1 | | | | 1 | | | | |
| 5812996678 | | | | | | | | 1 | | | | | | 1 | 1 | | | | | | | | | | | | | |
| 5812996681 | | | | | | 1 | | | | | | | | | | | | | | | | | | | | | | |
| 5812996748 | | | | | | | | | 1 | | | | | | | | | | | | | | 1 | | | 1 | | |
| 5812997016 | | | | | | 1 | | | | | | | | | | | | | | | | | | | | | | |
| 5813000357 | | | | | | | | 1 | | | | | | | | | | | | | | | | | | | | |
| 5813000891 | | | | | 2 | | | | | | | | | | | | | | | | | | | | | | | |
| 5813001741 | | | | | 2 | 1 | | 2 | | | | | | 2 | | | | | | | | | | | | | | |
| 5813002592 | | | | | | | | | | | | | | | | | | | | 1 | | 1 | | | | | | |
| 5813002721 | | | | | | | | 3 | | | | | | | | | | | | | | | | | | | | |
| 5813002722 | | | | | | | | 1 | | | | | | | | | | | | | | | | | | | | |
| 5813002810 | | | | | | | | | | | | | | | | | | | 1 | | | | | | 1 | 2 | 1 | |
| 5813002812 | | | | | | | | | | | | | | | | | | | | | | 1 | | | | | | |
| 5813002813 | | | | | | | | | | | | 1 | | | 3 | | | 8 | 5 | 5 | 3 | 2 | | | | | | |
| 5813002840 | | | | | | | | | | | | | | 1 | | | | | | | | | | | | | | |
| 5813007108 | | | | | | | | | 11 | 7 | 3 | 1 | 5 | 1 | 2 | | | | | | | | | | 1 | 2 | | |
| 5813007116 | | | | | | | | | | | | 5 | | | | | | | | | | | | | | | | |
| 5813008796 | | | | | | | | | | | | | | | | | | 1 | | | | 1 | 13 | 4 | | | | |
| 5813008848 | | | | | | | | | | | | | | | | | | | | | | | 5 | 3 | | | | |
| 5813008911 | | | | | | | | | 4 | 5 | 7 | | 1 | | | | | | | | | | | | | | | |
| 5813008919 | | | | | | | | | | | | | | | | | | | | | | | 1 | | | | | |
| 5813008922 | | | | | | | | | | 1 | | | | | | | | | | | | | | | | | | |
| 5813009260 | | | | | | | | | | | | | | | | | | | | | | | 4 | 3 | | | | |
| 5813009280 | | | | | | | | | | | | | | | | | 1 | | | | | | | | | | | |
| 5813009350 | | | | | | | | | | | | 1 | | | | | | | 1 | | | | | | | | | |
| 5813009352 | | | | | | | | | | 1 | | | | | | | | | | | | | | | | | | |
| 5813009354 | | | | | | | | | | | | | | | | | | | | | | | 3 | | | | | |
| 5813009378 | 5 | 12 | 4 | 7 | | | | | | | | | | 1 | | | 2 | | | | | | 2 | | | | | |
| 5813009416 | | | | | | | | | | | | | | | | | | | | | | | | | 1 | | | |
| 5813009449 | | | | | | | | | | | | | | | | | | | | | | | | | 1 | | | |
| 5813009486 | | | | | | | | | 1 | | | | | | | | | | | | | | | 3 | | | | |
| 5813009521 | | | | | | | | | | | | | | | | | | | | | | | | 3 | | | | |
| 5813009554 | | | | | | | | | 2 | | 2 | | | | | | | | | | | | | | | | | |
| 5813009589 | | | | | | | | | | | | | | | | | | | | | | | | 1 | | | | |
| 5813009651 | | | | | | | | | | | 2 | | | | | | | | | | | | | | | | | |
| 5813009723 | | | | | | | | | | | | | | 1 | | | | | | | | | | | | | | |
| 5813009745 | | | | | | | | | | | | | | | | | | | | | | | 1 | | | | | |
| 5813009749 | | | | | | | | | | | | | | | | | | | | | | | 1 | | | | | |
| 5813009781 | | | | | | | | | | | | | | | | | | | | | | | | 1 | | | | |
| 5813009845 | | | | | | | | | | | 1 | 1 | | 2 | 1 | | | 4 | 6 | 7 | 4 | 9 | | 1 | 2 | | 3 | |
| 5813009865 | | | | | | | | | | | | | | | | | | | | | | | | 1 | | | | |
| 5813009881 | | | | | | | | | 21 | 5 | 9 | | | | | | | | | | 1 | | | | | | | |
| 5813009882 | | | | | | | | | 9 | 6 | 2 | | | | | | | | | | | | | | | | | |
| 5813009946 | | | | | | | | | 11 | 6 | 7 | | | | | | | | | | | | | | 4 | | | |
| 5813009950 | | | | | | | | | 1 | | 1 | | | | | | | | | | | | | | | | | |
| 5813009972 | | | | | | | | | | 1 | | | | | | | | | | | | | | | | | | |
| 5813009988 | | | | | | | | | | | | 3 | 1 | | | | | | | | | | | | | | | |
| 5813010120 | | | | | | | | | | | 1 | | | 2 | | | | | | | | | 1 | | | 4 | 2 | |
| 5813010121 | | | | | | | | | | | | | | | | | | | | | | | | | 2 | 3 | 7 | |
| 5813010135 | | | | | | | | | | | | 1 | | | | | | | | | | | | 5 | | | | |
| 5813010139 | | | | 1 | | | | | | | | | | | | | 1 | | | | | | | | | | | |
| 5813010153 | | | | | | | | | | | | | | 10 | 9 | | | | 1 | 2 | 1 | 1 | 3 | 3 | | 1 | 1 | |
| 5813010376 | | | | | | | | | | | | | | | 10 | 9 | | | | 1 | 3 | 2 | 10 | 2 | 14 | | 2 | |
| 5813010377 | | | | | | | | | | | | | 1 | 3 | 12 | | 2 | 1 | 4 | 2 | 3 | 5 | 21 | 12 | | | | |
| 5813010435 | | | | | | | | | | | | | | | | | | 4 | | | | 1 | 5 | 4 | 1 | 8 | 5 | |
| 5813010506 | | | | | | | | | | | | | | | | | | 2 | | | | | | | | | | |
| 5813010682 | | | | | | | | | | | | | | | | | | | 2 | 3 | | 2 | 2 | 5 | | | | |
| 5813010683 | | 1 | | 1 | | | | | | | | | 1 | 9 | 3 | | | 1 | 2 | 2 | | 1 | 1 | 2 | 2 | 12 | 3 | |
| 5813010721 | | | | | | | | | | | | | | | | | | | | | 2 | | | | | | | |
| 5813010722 | | | | | | | | | | | | | | | | | | 1 | | 1 | | | | | | | | |
| 5813010729 | | | | | | | | | | | | | | 1 | | | | | | | | | | | | | | |
| 5813010730 | | | | | | | | | | | | | | 1 | | | | | | | | | | | | | | |
| 5813010738 | | | | | | | | | | | | | | | | | | 1 | | | | 1 | | | | | | |
| 5813010828 | | | | | | | | | 1 | | | | | | | | | | | | | | | | | | | |
| 5813010938 | | | | | | | | | | | | 5 | 7 | | | | | | | | | | | | | | | |
| 5813010996 | | | | | | | | | | | | | | | | | | | | 4 | | | | | | | | |
| 5813011015 | | | | | | | | | | | | | | 3 | | | | | | | | | | 2 | | | 1 | |
| 5813011024 | | | | | | | | | | | | | | | | | | | | | | | | | 1 | | | |
| 5813011133 | | | | | | | | | | | | | | | | | | | | | | | | | 1 | | | |
| 5813011165 | | | | | | | | | | | | | | 4 | 1 | | | | | | | | | | | | | |
| 5813011306 | | | | | | | | | | | | | | | | | | | | | | 1 | | | | | | |
| 5813011308 | | | | | | | | | | | | | | 1 | | | | | | | | 1 | | | | | | |
| 5813011327 | | | | | | | | | | | | | | | | | | | | 6 | | | | | | | | |
| 5813011383 | | | | | | | | | | | | 1 | 4 | 1 | | | | | | | | 1 | | | | | | |
| 5813011384 | | | | | | | | | | | | | | | | | | | | | 1 | | | | | | | |
| 5813011391 | | | | | | | | | | | | | | 1 | | | | | | | | | | | | | | |
| 5813011407 | | | | | | | | | | | | 2 | | | | | | | | | | | | | | | | 1 |

| | | | | | | | | | | | | | | | | | | | | | | | | | | | | |
|---|---|---|---|---|---|---|---|---|---|---|---|---|---|---|---|---|---|---|---|---|---|---|---|---|---|---|---|---|
| 5813011443 | | | | | | | | | | | | | | | | | | | | | | | | | | | | 8 |
| 5813011483 | | | | | | | | | | | | 2 | | | | | | | | | | | | | | | | |
| 5813011493 | | | | | | | | | | | | | | | | | | | | | | | | | | | | 1 |
| 5813011660 | | | | | | | | 3 | | | | | | 2 | 4 | | 1 | | | | | | | 1 | | | | |
| 5813011683 | | | | | 1 | | 1 | 1 | | | | | | | 1 | 1 | | | | | | | | | | | | |
| 5813011742 | | | | | | | | | | | | | 1 | | | | | | | | | | | | | | | |
| 5813012113 | | | | | | | | | | | | 8 | 3 | 2 | | | | | | | | | | | | | | |
| 5813012172 | | | | | | | | | | | | | 1 | | | | | | | | | | | | | | | |
| 5813012188 | | | | | | | | | | | | 2 | | | | | | | | | | | | | | | | |
| 5813012196 | | | | | | | | | | | | | | 3 | 1 | | | | | | | | | | | | | |
| 5813012366 | 2 | 1 | | | | | | | | | | 1 | 2 | 4 | 1 | 23 | 18 | | | | | | | 1 | | | | |
| 5813012529 | | | | | | | | | | | | | | 1 | 2 | | | | | | | | | | | | | |
| 5813012781 | | | | | | | | | | | | | | | | 2 | | | | | | | | | | | | |
| 5813012786 | | | 1 | | | | | | | | | | | | | | | | | | | | 5 | 4 | | 1 | 4 | |
| 5813012837 | | | | | | | | | | | | | | 1 | | | | | | | | | | | | | | |
| 5813012838 | | | | | | | | | | | | | 1 | | | | | | | | | | | | | | | 1 |
| 5813013091 | | | | | | | | | | | | | | | | | 1 | | | | | | | | | | | |
| 5813013185 | | | | | | | | | | | | | 1 | | | | | | | | | | | | | | | |
| 5813013436 | | | | | | | | | | | | | | | | | | | | | | | | 1 | | | | |
| 5813013547 | | | | | | | | | | | | 17 | 6 | | | | | | | | | | | | | | | |
| 5813013843 | | | | | | | | | | | | 8 | 3 | 5 | | | | | | | | | | | | | | |
| 5813014028 | | | | | | | | | | | | | | | | | | | | | | | 2 | | | | | |
| 5813014029 | | | | | | | | | | | | | | | | | | | | | | | 2 | 4 | | | | |
| 5813014166 | | | | | | | | | | | | | | | | 1 | | | | | | | | | | | 1 | |
| 5813014341 | | | | | | | | | | | | 1 | | | | | | | | | | | | | | | | |
| 5813014572 | | | | | | | | | | | | | | | | | 1 | | | | | | | | | | | |
| 5813014907 | | | | | | | | 1 | | | | | | 2 | 1 | | | | | | | | | | | | | |
| 5813014911 | | | | | | | | | | | | | | | | | | | | | | | | 1 | | | | |
| 5813016373 | | | | | | | | | | | | | | | | | | | | | | | | 1 | | | | |
| 5813017614 | | | | | | | | | | | | | | | | | 1 | | | | | | | | | | | |
| 5813019517 | | | | | | | | | | | | | 1 | | | | | | | | | | | | 12 | | | |
| 5813019526 | | | | | | | | | | 2 | | | | | | | | | | | | | | | | | | |
| 5813019568 | | | | | | | | | 2 | | 2 | | | | | | | | | | | | | | | | | |
| 5813019732 | | | | | | | | | | | | | | | | | | | | | | | | | 17 | | | |
| 5813019745 | | | | | | | | | | | | | | | | | | | | | | | | | | | | 1 |
| 5813019868 | | | | | | | | | | | | 1 | | 1 | | 2 | | | | | | | 1 | 2 | | | | |
| 5813020136 | | | | | | | | | | | | | | | | 2 | | | | | | | | | | | | |
| 5813020143 | | | | | | | | | | | | | | | | | 1 | | | | | | | | | | | |
| 5813020347 | | | | | | | | | | | | 7 | 1 | | | | | | | | | | | | | | | 3 |
| 5813020673 | | | | | | | | | | | | | | | | 1 | | | | | | | | | 1 | | | |
| 5813020684 | | | | | | | | | | | | | | | | | | | | | | | | | | | 1 | |
| 5813020696 | | | | | | | | | 5 | 1 | 3 | 3 | 9 | 12 | 11 | | | | | | | | 6 | | | | 1 | |
| 5813020698 | | | | | | | | | | | | | | | | | | | | | | | 6 | | | | | |
| 5813020750 | | | | | | | | 1 | | | | | | 1 | | 1 | | | | | | | | | | | | |
| 5813020795 | | | | | | | | | | | | | | | 1 | | | | | | | | | | | | | |
| 5813020800 | | | | | | | | | | | | | | | | | | | | | | | | | | | 1 | |
| 5813020809 | | | | | | | | | | | | | | | | | | | | | | | | 1 | | | | |
| 5813020812 | | | | | | | | | | | | | | | | | | | | | | | 3 | 4 | 1 | | | |
| 5813020846 | | | | | | | | | | | | | | | | | 1 | | | | | | 1 | | | | | |
| 5813020850 | | | | | | | | | | | 5 | | | 1 | | | | | | | | | | | | | | |
| 5813020864 | | | | | | | | | | | | | | 8 | | | | | | | | 1 | 2 | | | | | |
| 5813020969 | | | | | | | | | | 3 | 4 | | | | | | | | | | | | | | | | | |
| 5813020993 | | | | | | | | | | | 2 | | | | | | | | | | | | | | | | | |
| 5813020996 | | | | | | | | | | | | | | | | | | | | | | | | | 3 | | | 5 |
| 5813021005 | | | | | | | | | | | | | | | | | | | | | | | 1 | | | | | |
| 5813021094 | | | | | | | | | | | | | 1 | | | 1 | 1 | | | 2 | 1 | | | | 2 | | | |
| 5813021192 | | | | | | | | | | | | 1 | | 21 | 25 | | | 6 | 13 | 8 | 2 | 5 | 4 | 5 | 1 | 2 | 2 | |
| 5813021196 | | | | | | | | | | | | | | | 1 | 3 | 1 | | | | | | | | | | | |
| 5813021202 | | | | | 1 | | 1 | | | | | | | | | | | | | | | | | | | | | |
| 5813021217 | | | | | | | 2 | | | | | | | | | 4 | | | | | | | | | | | | |
| 5813021221 | | | | | | | | | | | | | | 1 | | | 1 | | | | | | | | | | | |
| 5813021225 | | | | | | | | | | | | 8 | 4 | | | | | | | | | | | | | 1 | | |
| 5813021480 | | | | | | | | | | | | | | | 3 | | | | | | | | | | | | | |
| 5813021485 | | | | | | | | | | | | | | | 1 | 1 | | | | | | | | | | | | |
| 5813021535 | | | | | | | | | | | | | 1 | | | | | | | | | | 1 | | | | | |
| 5813021564 | | | | | | | | | | | | 6 | 10 | 1 | | | | | | | | | | 1 | 13 | 3 | 2 | |
| 5813021566 | | | | | | | | | | | | 2 | 4 | 2 | | | | | | | | | | | | | | |
| 5813021602 | | | | | | | | | | | | | 1 | | | | | | | | | | | | 11 | 1 | | |
| 5813021615 | | 1 | | | | | | 3 | | | | | | 1 | | 5 | 3 | | | | | | 5 | 1 | | | | |
| 5813021666 | | | | | | | | | | | | | | 40 | 55 | 2 | | | | | | | 1 | | | | | |
| 5813021671 | | | | | | | | | | | | | | | | | | | | | | | 1 | | | | | |
| 5813021689 | | | | | | | | | | | | | | | | | | | 1 | | | | | | | | | |
| 5813021788 | | | | | | | | | | | | | | | | | 1 | | | | | | | | | | | |
| 5813021874 | | | | | | | | | | | | | | 1 | | | | | | | | | | | | | | |
| 5813021882 | | | | | | | | | | | | | | 1 | | | 1 | | | | | | | | | | | |
| 5813021904 | | | | | | | | | | | | | | | | | 1 | | | | | | | | | | | |
| 5813021922 | | | | | | | | | | | | | | 2 | 2 | | 5 | | | | | | | | | | | |
| 5813021943 | | | | | | | | | | | | | | | 3 | 1 | 2 | | | | | | | | | | | |
| 5813021948 | | | | | | | | | | | | | | 3 | 1 | 18 | 26 | | | | | | | | | | | |
| 5813022037 | | | | | | | | | | | | | | | | 1 | 1 | | | | | | | | | | | |

| | | | | | | | | | | | | | | | | | | | | | | | | | | | | |
|---|---|---|---|---|---|---|---|---|---|---|---|---|---|---|---|---|---|---|---|---|---|---|---|---|---|---|---|---|
| 5813022150 | | | | | | | | | | | | | | 1 | | 1 | 1 | | | | | | | | | | | |
| 5813022160 | | | | | | | | | | | | | | | | | | | | | | | | 4 | | | | |
| 5813022182 | | | | | | | | | | | | | | 1 | | | | | | | | | | | | | | |
| 5813022192 | | | | | | | | | | | | | | 2 | 1 | | | | | | | | | | | | | |
| 5813022272 | | | | | | | | 2 | | | | | | | | | | | | | | | | | | | | |
| 5813022274 | | | | | | | | | | | | | | 3 | 3 | 10 | | | | | | | | | | | | |
| 5813022453 | | | | | | | | | | | | | | | | 1 | 4 | | | | | | | | | | | |
| 5813022753 | | | | | 1 | | | | | | | | | 20 | 27 | 38 | 34 | | | | | | | | | | | |
| 5813022839 | | | | | | | | | | | | | | 2 | 11 | | | | | | | | | | | | | |
| 5813023097 | | | | | | | | 1 | | | | | | 3 | 7 | | | | | | | | | | | | | |
| 5813024125 | | | | | 1 | | | | | | | | | | | | | | | | | | | | | | | |
| 5813024215 | | | | | 1 | | | | | | | | | | | | | | | | | | | | | | | |
| 5813024789 | | | | | 1 | | | | | | | | | 1 | | | | | | | | | | | | | | |
| 5813024792 | | | | | | | | 3 | | | | | | | | | | | | | | | | | | | | |
| 5813025330 | | | | | | | 2 | 2 | | | | | | | | | | | | | | | | | | | | |
| 5813025489 | | | | | | | | | | | | | | | | | | | 1 | | | 1 | | | | | | |
| 5813026364 | | | | | | | | | | | | | | | 2 | | | | | 2 | 1 | | 29 | 16 | | | | |
| 5813026366 | | | | | | | | | | | | | | | | | | | | | | | 1 | 1 | | | | |
| 5813026404 | | | | | | | | | | | | | | | 2 | | | | | | | | | | | | | |
| 5813026579 | | | | | | | | | | | | 4 | | 2 | 4 | | | | | | | | | 1 | | | | |
| 5813026589 | | | | | | | | | 1 | | | 15 | 10 | | | | | 1 | | | | | | | 18 | 18 | 10 | |
| 5813026613 | | | | | | | | | 1 | 1 | | | 2 | | 1 | | 1 | | | | | | 1 | | 3 | 42 | 40 | |
| 5813026646 | | | | | | | | | | | | | | 7 | | | | | | | | | | | | | | |
| 5813026661 | | | | | | | | | | | | | | | | | | | | | | | | | | 3 | 1 | |
| 5813026705 | | | | | | | | | | | | | | | | | | | | | | | | | 2 | | | |
| 5813026773 | | | | | 1 | 1 | | | | | | | | 1 | | | | | | | | | | | | | | |
| 5813026780 | | | | | | | | | 1 | | 2 | | 2 | 3 | 3 | | | | | | | 1 | | | | | | |
| 5813026785 | | | | | | | | | | | | | | | | | | | | | | | | | | | | 1 |
| 5813027028 | | | | | | 1 | | 3 | | | | | | | 1 | 2 | | | | | | | | | | | | |
| 5813027244 | | | | | | | | | | | | | | | | | 2 | | | | | | | | | | | |
| 5813027248 | | | | | | | | | | | | | | | | | 3 | | | | | | | | | | | |
| 5813032640 | | | | | | | | | | 4 | 3 | | 1 | | | | | | | | | | | | | | | |
| 5813032847 | | | | | | | | | | | | | | | | | | | | | | | | | 2 | | | |
| 5813032876 | | | | | | | | | | | | | | | | | 2 | | | | | | | | | | | |
| 5813032924 | 1 | | 2 | 1 | | | | | | | | | | | | | 1 | | | | | | | | | | | |
| 5813035070 | | | | | | | | | | | | 1 | 4 | | | | | | | | | | | | | | | 11 |
| 5813035111 | | | | | | | | | | | | | | 1 | 1 | | | | | | | | | | | | | |
| 5813035148 | | | | | | | | | | | | | 2 | 2 | 5 | | | 3 | 1 | 1 | | | | | | | | |
| 5813035236 | | | | | | | | | | | 1 | | | | | | | | | | | | | | 2 | | | |
| 5813035328 | | | | | | | | | | | | | | 2 | | | | 1 | | | | | | | | | | |
| 5813039659 | | | | | | | | | | | | | | | 1 | | | | | | | | | | | | | |
| 5813039662 | | | | | | | | | | | | 1 | 2 | | 2 | | | 4 | | 4 | 1 | | | | | | | 3 |
| 5813039678 | | | | | | | | | | | | 2 | | | 1 | | | 1 | 1 | | | 4 | | | | | | |
| 5813039679 | | | | | | | | | 1 | | | | | | | | | | | | | | | | | | | |
| 5813039857 | | | | | | | | | | | 2 | | | | | | | | | | | | | | | | | |
| 5813039859 | | | | | | | | | 1 | 1 | 2 | | | | | | | | | | | | | | | | | |
| 5813039860 | | | | | | | | | 1 | | | | | | | | | | | | | | | 2 | | 2 | | |
| 5813039902 | | | | | | | | | 12 | 2 | 3 | 3 | | | 1 | | | | 1 | | | | | | | | | |
| 5813039910 | | | | | | | | | 4 | 20 | 33 | | | | | | | | | | | | | | | | | |
| 5813039936 | | | | | | | | | | | | | | | | | | 2 | 1 | 3 | 3 | 2 | | | | | | |
| 5813039993 | | | | | | | | | | | | | | 3 | | | | | | | | | | | | | | |
| 5813040119 | | | | | | | | | | | | | | | | | 1 | | | | | | | | 1 | | 1 | |
| 5813040216 | | | | | | | | | | | | | | | | | | | | | | | 1 | | | | | |
| 5813040494 | | | | | | 4 | 1 | 1 | | | | | | | 2 | | | | | | | | | | | | | |
| 5813040711 | | | | | | | | | | | | | | | | | | | 1 | | | | | | | | | |
| 5813040712 | | | | | | | | | | | | 36 | 25 | 1 | 1 | | | | | | | | | | | | | 5 |
| 5813040731 | | | | | | | | | | | | | | 1 | | | | | | | | | 1 | | | | | |
| 5813040744 | | 1 | | 1 | | | | | | | | 1 | | 18 | 26 | 60 | 49 | | | | | | | 1 | | | | |
| 5813041149 | | | | | | | | | | | | | 3 | | | | | | | | | | | | | | | |
| 5813041180 | | | | | | | | | | | | | | | | | | | | | | | 3 | 5 | | | | |
| 5813044812 | | | | | | | | | | | | | | | 1 | | | | | | | | | | | | | |
| 5813046221 | | | | | | 1 | | | | | | | | | | | | | | | | | | | | | | |
| 5813046658 | | | | | | | | | | | | | | 1 | 1 | | | 1 | | | | | | | | | | |
| 5813046674 | | | | | | | | | | | | | | | | | | | 1 | | | | | | | | | |
| 5813046951 | | | | | | | | | | | 2 | 1 | | | | | | | | | | | | | | | | |
| 5813047053 | 1 | | 1 | | | | | | | | | | | | | 2 | | | | | | | 1 | | 2 | | 1 | |
| 5813047065 | | | 1 | | | | | | | | | | | | | | | | | | | | 1 | | | | | |
| 5813047101 | | | | | | | | | | | | | 1 | 1 | | | | | | | | | 1 | | | | | |
| 5813047131 | | | | | | | | | | | | | | | | | | | | | | | | | | | | 3 |
| 5813047134 | | | | | | | | | | | | | | | | | | | | 1 | | 1 | | | | | | |
| 5813047183 | | | | | | | | | | | | | | 9 | 6 | | | 2 | 9 | 1 | 1 | 5 | | 1 | | 1 | | |
| 5813047189 | | | | | | | | | | | | | | | | | 1 | | | | | | | | | | | |
| 5813047284 | | | | | | | | | | | | | | | 1 | | | | | | | 2 | | | | 4 | | |
| 5813047586 | 57 | 84 | 69 | 64 | | | | | | | | | | | | | | | | | | 1 | | 1 | | | | |
| 5813047601 | | | | | | | | | | | | 2 | 2 | | 1 | | | | | | | | | | | | | |
| 5813047683 | | | | | | | | | | | | | | | | 1 | | | | | | | | | | | | |
| 5813047713 | | | | | | | | | 7 | 1 | 3 | | | | | | | | | | | | | | | | | |
| 5813047739 | | | | | | | | | | | | 3 | 4 | 1 | | | | 2 | 1 | 1 | | 2 | | 1 | 11 | 68 | 55 | 6 |
| 5813047772 | | | | | | | | | | | | | | | | | | | | | | | 4 | 2 | | | | |
| 5813047774 | | | | | | | | | | | | | | 2 | 1 | 3 | 3 | | | | | | | | | | | |

| | | | | | | | | | | | | | | | | | | | | | | | | | | | | |
|---|---|---|---|---|---|---|---|---|---|---|---|---|---|---|---|---|---|---|---|---|---|---|---|---|---|---|---|---|
| 5813048437 | | | | | | | | | | | | 13 | 5 | | | | | | | | | | | | | | | |
| 5813048451 | | | | | | | | | | | | | | | | | | | | | | | 1 | | | | | |
| 5813049102 | | | | | | | | | | | | | | | | | | | | | | | 1 | 2 | 5 | | | |
| 5813049136 | | | | | | | | | | | | | | | | | | | | | | | | | 1 | | | |
| 5813049344 | | | | | | | | | | | | | | 3 | | | | | | | | | | | | | | |
| 5813049438 | | | | | | | | | | | | 2 | | | 1 | | | 1 | 1 | | 1 | | | | | | | 1 |
| 5813049907 | | | | | | | | | 7 | 2 | 7 | | | | | | | | | | | | | 1 | | | | |
| 5813050292 | | | | | | 1 | | | | | | | | | | | | | | | | | | | | | | |
| 5813053831 | | | | | | | | | | | | | | | | | | | | | | | | 2 | | | | |
| 5813053907 | | | | | | | | | | | | | | | 3 | | | | | | | | | | | | | |
| 5813054835 | | | | | | | | | | | | | | | 1 | | | | | | | | | | | | | |
| 5813055709 | | | | | | | | | | | | | | | | | | | | | | | | | 1 | | | |
| 5813055759 | 1 | 1 | 3 | 10 | | | | | | | | | 1 | | 2 | | | | | | | | 1 | 7 | 1 | | | |
| 5813055761 | | | | | | | | | | | 1 | | | | | | | | | | | | | | | | | |
| 5813055766 | | | | | | | | | 3 | | | | | | | | | | | | | | | | | | | |
| 5813055768 | | | | | | | | | 1 | | | | | | | | | | | | | | | | | | | |
| 5813055773 | | | | | | | | | 1 | | 4 | | | | | | | | | | | | | | | | | |
| 5813055790 | 1 | 1 | 2 | | | | | | | | | | | | 7 | | 1 | 2 | | 8 | 2 | 3 | 1 | 6 | | | | |
| 5813055817 | | | | | | | | | | 1 | 1 | 1 | | 3 | | | | | | | | | | | | | | |
| 5813055822 | | | | | | | | | | | | | 1 | 1 | | | | | | | | | | | | | | |
| 5813055865 | | | | | | | | | | | | | | | | | 1 | | | | | | | | | | | |
| 5813055932 | | | | | | | | | | | | | | | | | | | | | | | | | 1 | | | |
| 5813055949 | | | | | | | | | | | 1 | | | | | | | | | | | | 1 | | 6 | 1 | 1 | |
| 5813056062 | | | | | | | | | | | | | | 9 | 4 | | | 4 | 5 | 3 | 2 | 8 | | | | | | |
| 5813056071 | | | | | | | | | | | | | 6 | | | | | | | | | | | | | | | |
| 5813056168 | | | | | | | | | | | | | 2 | 5 | 1 | | | | | | | | | | | | | |
| 5813056278 | | | | | | | | | | | | | | | | 1 | | | | | | | | | | | | |
| 5813056759 | | | | | 3 | | | | | | | | | | | | | | | | | | | | | | | |
| 5813056887 | | | | | | | | | 1 | | 1 | 2 | 2 | 4 | 8 | | | | | | | | | | 10 | | | |
| 5813056896 | | | | | | | | | | | | 1 | | | | | | | | | | | 1 | | | | | |
| 5813056917 | | | | | | | | | | | | | 5 | 31 | 18 | | 1 | 9 | 6 | 5 | 2 | 8 | 3 | 4 | | 2 | | |
| 5813056920 | | | | | | | | | | | | | | | | | | | | | | | 1 | | | | | |
| 5813056932 | | | | | | | | | | | | 1 | 2 | | | | | | | | | | | | | | | |
| 5813057153 | 2 | | 2 | | | | | | | | | | | 19 | 32 | 4 | 7 | | | | 1 | | 1 | 1 | | | | |
| 5813057162 | | | | | | | | | | | | | | | | | | | | | | | | | 3 | | | |
| 5813057615 | | | | | | | | | | | | | | | 3 | 1 | 2 | | | | | | | | | | | |
| 5813058028 | | | | | | | | | | | | | | | | 2 | 4 | | | | | | | | | | | |
| 5813058085 | | | | | | | | | | | | | | | | | 1 | | | | | | | | | | | |
| 5813058097 | | | | | | | | | | | | 5 | 3 | | | | | | | | | | | | | | | |
| 5813058222 | | | | | | | | | | | | | 2 | | | | | | | | | | | | | | | |
| 5813058431 | | | | | | | | 4 | | | | | | | 1 | | 1 | | | | | | | | | | | |
| 5813060697 | | | | | | | | | | | | | | | | | | | | | | | | | | | | 1 |
| 5813060768 | | | | | | | | | | | | | | | | | | | | | | | 3 | 6 | | | | |
| 5813061249 | | | | | | | | | | | | | 4 | | | | | | | | | | | | | | | |
| 5813061260 | | | | | | | | | | | | | | | | | | | | | | | | | 1 | | | |
| 5813061690 | | | | | | | | | | | | | | | | | | | | | | | | | | 1 | | |
| 5813062858 | | | | | | | | | | | | | | | | | | | | | | | | | 1 | | | |
| 5813062922 | | | | | | | | 1 | | | | | | | | | | | | | | | | | | | | |
| 5813063150 | | | | | | | | | | | | | | 2 | 2 | | | | | | | | | | | | | |
| 5813063151 | | | | | | | | | 2 | | | | | | 2 | | | | | | | | 1 | | | | | |
| 5813063216 | | | | | | | | | | | | | | | | | | | | | | | | | | 1 | | |
| 5813063227 | | | | | | 1 | | 3 | | | | | | | | | | | | | | | | | | | | |
| 5813063239 | | | | | | | | | | | | | | | | | 1 | | | | | | | | | | | |
| 5813063829 | | | | | | | | | | | | | 1 | | 1 | 8 | 3 | | | | | | | | | | | |
| 5813064332 | | | | | | | | | | | | 8 | 7 | 1 | | | | | | | | | | | | | | |
| 5813064789 | | | | | | | | | 3 | 3 | | 2 | | | 1 | | | | | | | | 2 | | | | | |
| 5813064828 | | | | | | | | | 2 | 1 | 5 | | | | | | | | | | | | | | | | | |
| 5813067334 | | 2 | | 4 | | | | | | | | | | 3 | 2 | | | 3 | 3 | 6 | 4 | | 2 | 1 | | | | |
| 5813067390 | | | | | | | | | 2 | 1 | | | | | | | | | | | | | | | | | | |
| 5813067452 | | | | | | | | | | | 1 | | | | | | | | | | | | | | | | | |
| 5813067463 | | | | | | | | | 14 | 3 | | | | 2 | 3 | | | | | | 1 | | | | | | | |
| 5813067479 | | | | | | | | | | 1 | | | | | | | | | | | | | | | | | | |
| 5813067666 | | | | | | | | | | | 1 | | | | 1 | | | | | | | | | | | | | |
| 5813067725 | | | | | | | | | | | | | | | | | | | | | | | | 4 | | | | |
| 5813067799 | | | | | | | | | | | | | | | | | 1 | | | | | | | | | | | |
| 5813068296 | | | | | | | | | | | | | | 3 | 1 | | | 2 | | | | | | | | | | |
| 5813068467 | | | | | | | | | 2 | | | | | | | | | | | | | | | | | | | |
| 5813068563 | | | | | | | | | | | | 9 | 9 | | | | | | | | | | | | | | | |
| 5813068595 | | | | | | | | | | | | | 1 | | | | | | | | | | | | | | | |
| 5813068828 | | | | | 1 | 1 | | 1 | | | | | | | | | | | | | | | | | | | | |
| 5813068964 | | | | | | | | | | | | | | | | | | | | | | | 1 | | | | | |
| 5813068976 | | | | | | 1 | | | | | | | | | | | | | | | | | | | | | | |
| 5813068984 | | | | | | | | | | | | | | | | 1 | | | | | | | | | | | | |
| 5813069072 | | | | | | | | | 2 | | 1 | | | | | | | | | | | | | | | | | |
| 5813069078 | | | | | | | | | | | | | | | | | | 1 | | | | | | | | | | |
| 5813069193 | | | | | | | | 1 | | | | | | | 1 | | | | | | | | | | | | | |
| 5813069255 | | | | | 1 | | | | | | | | | | | | | | | | | | | | | | | |
| 5813069498 | | | | | | | | | | 1 | | | | | | | | | | | | | | | | | | |
| 5813069570 | | | | | 2 | | | | | | | | | | | | | | | | | | | | | | | |
| 5813069630 | | | | | | | | | | | 2 | | | | | | | | | | | | | | | | | |

| | | | | | | | | | | | | | | | | | | | | | | | | | | | | |
|---|---|---|---|---|---|---|---|---|---|---|---|---|---|---|---|---|---|---|---|---|---|---|---|---|---|---|---|---|
| 5813069636 | | | | | | | | | | | | | | 1 | | | | | | | | | | | | | | |
| 5813069648 | | | | | | | | | | | | | 1 | | 40 | 81 | 75 | 15 | 37 | 30 | 34 | 30 | 4 | 11 | | | | |
| 5813069844 | | | | | | | | | | | | | | 2 | 2 | | | | | | | | | | | | | |
| 5813069859 | | | | | | | | | | | | 1 | 2 | | | | | 1 | | | | | 1 | 2 | 23 | 1 | | 2 |
| 5813070017 | | | | | | | | | | | | | | 5 | 3 | 1 | | | | | | | | | | | | |
| 5813070991 | | | | | | | | | | | | | | | | | | | | | | | | | | | | 1 |
| 5813071288 | | | | | | | | | 2 | 3 | 2 | | | 8 | 1 | | | | | | | | 2 | 3 | | 1 | | |
| 5813071319 | 1 | | 1 | | | | | | | | | | | | 1 | 1 | | 2 | 1 | | | 1 | 5 | | | 1 | | |
| 5813071320 | | | | | | | | | | | | | | | | | | | | | | | 1 | | | | | |
| 5813075619 | | | | | | | | | | | | | 2 | | | | | | | | | | | | | | | |
| 5813075715 | | | | | 2 | 3 | | 13 | | | | | | 1 | 1 | | | | | | | | | | | | | |
| 5813075772 | | | | | | | | | | | | | | | | | | | | | | | | | | | | 1 |
| 5813075909 | | | | | | | | | | | | | | | | | | | | | | | 3 | | | | | |
| 5813075931 | | | | | | | | | | | | | | | | | | | | | | | | | 1 | | | |
| 5813075932 | | | | | | | | | | | | 2 | 5 | | | | | | | | | | | | | | | |
| 5813076017 | | | | | | | | | | | | | | | | | | | | | | | 2 | | | | | |
| 5813076981 | | | | | | | | | | | | | | | | 2 | | | | | | | | | | | | |
| 5813077405 | | | | | | | | | | | | | | 2 | 1 | | | | 1 | | | | | | | | | |
| 5813077469 | | | | | | | | | | | | | | | | | | | | | | | | | | | | 1 |
| 5813077488 | | | | | | | | | | | | 1 | | | | | | | | | | | | | | | | 8 |
| 5813077562 | | | | | | | | | 1 | | 1 | | 1 | | | | 1 | | | | | | 4 | | | 1 | | |
| 5813077567 | | | | | | | | | | | | | | | | | | | | | | | 1 | 1 | | | | |
| 5813077788 | | | | | | | | | | | | | | | | | | | | | | | | 1 | | | | |
| 5813077891 | | | | | | | | | 1 | 6 | 4 | | | | | | | | | | | | | | | | | |
| 5813078095 | | | | | | | | | | | | | | | 1 | | 3 | | | | | | | | | | | |
| 5813078300 | | | | | | | | | 2 | | | | | | | | | | | | | | 2 | | | | | |
| 5813078542 | | | | | | | | | | | | | | | | | | | | | | | 2 | | | | | |
| 5813078557 | 4 | 6 | 1 | | | | | 1 | | | | | | 2 | 2 | 18 | 27 | | 1 | | | | 1 | | | | 2 | |
| 5813078563 | | | | | | | | | | | | | | | | | | | | | | | | | 1 | | | |
| 5813078603 | | | | | | | | | | | | 2 | | | | | | | | | | | | | | | | |
| 5813078679 | | | | | | | | | | | | | | | | 1 | 1 | | | | | | | | | | | |
| 5813079014 | | | | | | | | | | | | | | | | | | | | | | | 2 | | | | | |
| 5813079452 | | | | | | | | | | | | | | | | | | | | | | | | | 1 | | | |
| 5813081849 | | | | | | | | | | | | | 5 | | | | | | | | | | | | | | | |
| 5813082948 | | | | | | | | | | | | | | | | | | | | | | | 1 | | | | | |
| 5813083733 | | | | | | | | | 1 | | | | | | | | | | | | | | | | | | | |
| 5813087531 | | | | | | | | | | | | 2 | 3 | 18 | 20 | | | 2 | 11 | 6 | 5 | 13 | 12 | 15 | 5 | 1 | 2 | |
| 5813087560 | | | | | | | | | | | | | | | | | | | | | | | | | | | | 1 |
| 5813087939 | | | | | | | | | | | 4 | | | | | | | | | | | | | | | | | |
| 5813087991 | | | | | | | | | | | | 1 | | | | | | | | | | | | | | | | |
| 5813088412 | | | | | | | | | | | | | | | 1 | | | 8 | 1 | 4 | 3 | 3 | | 2 | | | | |
| 5813088525 | | | | | | | | | | | | 19 | 17 | 1 | | | | | | | | | | | | | | |
| 5813089132 | | | | | | | | | | | | | | | | | | | 3 | | 2 | 1 | | | | | | |
| 5813091698 | | | | | | | | | | | | | | | | | | | | | | | 1 | | 1 | | | |
| 5813092319 | | | | | | | | | 17 | 7 | 15 | 1 | 1 | 1 | 4 | | | | | | | | | | 1 | 1 | | |
| 5813094916 | | | | | | | | | | | | | | 1 | | | | 3 | 3 | | 4 | 1 | | | | | 1 | |
| 5813096638 | | | | | | | | | | | | | | 3 | | | | | | | | | | | | | | |
| 5813097458 | | | | | | 1 | | 3 | | | | | | | | | | | | | | | | | | | | |
| 5813097632 | | | | | | | | | | | | | | | | | | | | | | | | | | 1 | | |
| 5813097957 | | | | | | | | | | | | | | 10 | 9 | | | | | | | | | | | | | |
| 5813098375 | | | | | | | | | | | | | | 2 | | | | | | | | | | | | | | |
| 5813099605 | | | | | | | | | | | | | | | | | | | | | | | | 1 | 2 | | | |
| 5813108084 | | | | | | | | | | | | | | | | 5 | 5 | | | | | | | | | | | |
| 5813109064 | | | | | | | | | | | | 8 | 10 | | | | | | | | | | | | | | | |
| 5813109120 | | | | | | | | | | | | | | 6 | | | | | | | | | | | | | | |
| 5813109159 | | | | | | | | | | | | | 1 | | | | | | | | | | | | | | | |
| 5813111989 | | | | | | | | | | | | 34 | 22 | | | | | | | | | | | | | | | |
| 5813113543 | | | | | | | | | | | | | | | | | | | | | | | 1 | | | | | |
| 5813115583 | | | | | | | | | | | | | | | | | 1 | | | | | | | | | | | |
| 5813124162 | | | | | | | | | | | | 1 | 3 | | | | | | | | | | | | | | | |
| 5813129114 | | | | | | | | | | | | | | | | | | | | | | | | | | | | 1 |
| 5813129572 | | | | | | | | | | | | | | | 1 | | 1 | | | | | | | | | 1 | | |
| 5813129573 | | | | | | | | | 2 | 2 | 3 | | | | | | | | | | | | | 1 | | | | |
| 5813130028 | | | | | | | | | | | | 3 | 9 | 4 | 6 | | | 2 | | | | | | | | 1 | | |
| 5813132515 | | | | | | | | | | | | | 3 | | | | | | | | | | | | | | | |
| 5813133506 | | | | | | | | | | | | 4 | | | | | | | | | | | | | | | | |
| 5813133974 | | | | | | | | | | | | | | | | | | | | | 1 | 1 | | | | | | |
| 5813134082 | | | | | | | | | | | | | 3 | | | | | | | | | | | | | | | |
| 5813134294 | | | | | | | | | | | | | | | | | | | | | | | | | | | | 9 |
| 5901193006 | | | | | | | | | | | | | | | | | | 10 | 1 | 4 | 3 | 3 | | | | | 2 | |
| 5901193416 | | | | | | | | | | | | | | | | | | | | | | | 1 | | | | | |
| 5901193482 | | | | | | | | | | | | | | | | | | | | | | | 2 | 3 | | | | |
| 5901195323 | | | | | | | | | | | | | | | 3 | | | | | | | | | | | | | |
| 5901195361 | | | | | | | | | | | | | | | | | | | | | | | | | 1 | | | |
| 5901195498 | | | | | | | | | 7 | 5 | 4 | | | | | | | | | | | | 2 | | | | | |
| 5901196159 | | | | | | | | | | | | | | | | | | | | | | | | | 1 | | | |
| 5901196628 | | | | | | | | | | | | 20 | 33 | 1 | 1 | | | | | | | | | | 9 | | | |
| 5901197274 | | | | | | | | | | | | | 2 | | | | | | | | | | | | | | | |
| 5901197487 | | | | | | | | | | | | | | | | | | | | | | | | | | | | 1 |
| 5901198030 | | | | | | | | | 1 | | | | | 2 | | | | | | | | | | | | | | |

| | | | | | | | | | | | | | | | | | | | | | | | | | | | | |
|---|---|---|---|---|---|---|---|---|---|---|---|---|---|---|---|---|---|---|---|---|---|---|---|---|---|---|---|---|
| 5901203310 | | | | | | | | | | | | | | | | | | | | | | | | | | | | 1 |
| 5901203987 | | | | | | | | | | | | | | | | | | | | | | | | | 1 | | | |
| 5901206990 | | | | | | | | | | | | | | 1 | | | | | | | | | | | | | | |
| 5901207157 | | | | | | | | | 1 | 5 | 2 | | | | | | | | | | | | | | | | | |
| 5901211896 | | | | | | | | | | | | | | | | | 5 | | | | | | | | | | | |
| 5901212649 | | | | | | | | | 2 | 1 | | | | | | | | | | | | | | | | | | |
| 5901215966 | | | | | | | | | | | | 1 | | 3 | 1 | | | 5 | 3 | 6 | 4 | 6 | | | | | | 5 |
| 5901220093 | | | | | | | | | | | | | | | | | | | | | | | | | 1 | | | |
| 5901220179 | | | | | | | | | | | | | | | 1 | | | | | | | | | | | | | |
| 5901221363 | | | | | | | | | | | | 1 | | | | | | | | | | | 2 | 1 | | | | |
| 5901222742 | | | | | 1 | | | 8 | | | | | | | 1 | | | | | | | | | | | | | |
| 5901227104 | | | | | | | | | | | | | | 1 | | | | | | | | | | | | | | |
| 5901232053 | | | | | | | | | | | | 12 | 9 | 9 | 3 | | | | | | | | | | 3 | 1 | 2 | 4 |
| 6400000773 | | | | | | | | | | | | | 3 | | | | | | | | | | | | 1 | | | |
| 7112612987 | | | | | | | | | | | | | | | 1 | 1 | | | | | | | | | | | | |
| 7112613371 | | | | | | | | | | | | | 1 | | | | | | | | | | | | | | | 1 |
| 7112613943 | | | | | | | | | | | | | | | 1 | | | | | | | | | | | | | |
| 7112614253 | | | | | | | | | | | | | | | | 7 | | | | | | | | | | | | |
| 7112614490 | | | | | | | | 1 | | | | | | 2 | 1 | | | | | | | | | | | | | |
| 7112615473 | | | | | | | | | | | | | | | 1 | | 2 | | | | | | | | | | | |
| 7112615863 | | 2 | 1 | | | | | | | | | | | | | | | | | | | | | | | | | |
| 7112616299 | | | | | | | | | | | | | | | 2 | | | | | | | | | | | | | |
| 7112617046 | | | | | | | | | | | | | | | | | | | | | | 1 | | | | | | |
| 7112622017 | | | | | | | | | | | | | | 1 | 2 | | | 1 | 1 | | | 2 | | | | | 1 | |
| 7112622236 | | | | | | | | | | | | | | | 1 | | | | | 1 | | | | | | | | |
| 7112625263 | | | | | | | | | | | | | | | | | | | | | | | | | | | | 2 |
| 7112625363 | | | 3 | | | | | | | | | | | | 1 | | | | | | | | | | | | | |
